# Natural variation of the streptococcal Group A carbohydrate biosynthesis genes impacts host-pathogen interaction

**DOI:** 10.1101/2024.11.04.621835

**Authors:** Kim Schipper, Sara M. Tamminga, Nicholas Murner, Matthew Davies, Paul Berkhout, Debra E. Bessen, Astrid Hendriks, Natalia Korotkova, Yvonne Pannekoek, Nina M. van Sorge

**Author notes:** Corresponding author: Prof. N.M. van Sorge, PhD, Amsterdam UMC location University of Amsterdam, Department of Medical Microbiology and Infection Prevention, Meibergdreef 9, IWO building IA3.211, 1105 AZ Amsterdam, the Netherlands. E. Authors contributed equally (Kim Schipper, Sara M. Tamminga).

## Abstract

*Streptococcus pyogenes* (*S. pyogenes*) is a leading cause of infection-related mortality in humans globally. The characteristic cell wall-anchored Group A Carbohydrate (GAC) is expressed by all *S. pyogenes* strains and consists of a polyrhamnose backbone with alternating *N*-acetylglucosamine (GlcNAc) side chains, of which 25% are decorated with glycerol phosphate (GroP). The genes in the *gacA-L* cluster are critical for GAC biosynthesis with *gacH-L* being responsible for the characteristic GlcNAc-GroP decoration, which confers the agglutination in rapid test diagnostic assays and contributes to *S. pyogenes* pathogenicity. Historical research papers described *S. pyogenes* isolates, so-called A-variant strains, that lost the characteristic GlcNAc side chain following serial animal passage. Genomic analysis of a single viable historic parent/A-variant strain pair revealed a premature inactivating stop codon in *gacI*, explaining the described loss of the GlcNAc side chain. Subsequently, we analyzed the genetic variation of the 12 *gacA-L* genes in a collection of 2,021 *S. pyogenes* genome sequences. Although all *gac* genes (*gacA-L*) displayed genetic variation, we only identified 26 isolates (1.3%) with a premature stop codon in one of the *gac* genes. Twelve out of 26 (46%) isolates contained a premature stop codon in *gacH*, which encodes the enzyme responsible for the GroP modification. To study the functional consequences of the different premature stop codons for GacH function, we plasmid-expressed three *gacH* variants in a *S. pyogenes gacH*-deficient strain. Cell wall analysis confirmed GacH loss-of-function for the studied *gacH* variants through the significant reduction of GAC GroP, complete resistance to killing by the human bactericidal enzyme group IIA-secreted phospholipase, and susceptibility to zinc toxicity. Overall, our data provide a comprehensive overview of the genetic variation of the *gacA-L* cluster in a global population of *S. pyogenes* strains and the functional consequences of rare inactivating mutations in *gacH* for host interaction.

## Introduction

*Streptococcus pyogenes* (*S. pyogenes*) is a human-restricted pathogen, responsible for hundreds of millions of infections globally each year (2, 3). *S. pyogenes* causes a wide spectrum of clinical manifestations ranging from pharyngitis and impetigo to invasive infections *e.g.* puerperal sepsis, necrotizing fasciitis, and streptococcal toxic shock syndrome. Moreover, repeated infections can result in the development of post-infectious sequelae such as glomerulonephritis and rheumatic heart disease. Together, infections by this single pathogen result in an estimated 500,000 deaths annually worldwide, with resource-limited areas and Indigenous populations being affected disproportionally (2–4). Developing new strategies for effective treatment and prevention of *S. pyogenes* infection and their complications remains a critical public health priority, as safe and effective vaccines are not yet available.

An important feature of the cell-wall of *S. pyogenes* is the Group A Carbohydrate (GAC)(5). The GAC is expressed by all *S. pyogenes* strains, which has resulted in application of GAC-reactive rapid diagnostic test kits for streptococcal group identification. The GAC glycopolymers comprise up to half of the cell wall mass and are composed of a linear polyrhamnose chain decorated with *N*-acetylglucosamine (GlcNAc) side chains (6, 7). Recently, researchers showed that 25% of GlcNAc side chains are further modified with negatively-charged glycerol phosphate (GroP) moieties (8). GAC is important for the structural integrity of the bacterial cell wall, and biosynthesis of the polyrhamnose backbone is essential for the viability of *S. pyogenes* (6, 9–11). Removal of the GlcNAc-GroP side chain resulted in increased *in vitro* killing by human whole blood, neutrophils, and platelet releasate and attenuated virulence in murine and rabbit infection models (12), whereas removal of just the GroP moiety resulted in resistance against human cationic antimicrobial proteins, human bactericidal enzyme group IIA-secreted phospholipase (hGIIa), lysozyme and histones, while increasing susceptibility to zinc (8, 13).

The 12-gene *gac* cluster is crucial for GAC biosynthesis and exhibits limited genetic variation across the *S. pyogenes* population (12, 14, 15). The first seven genes of the cluster (*gacABCDEFG*) encode proteins catalyzing the biosynthesis and transport of the polyrhamnose backbone (9, 12, 16). GacIJKL, encoded by *gacIJKL*, are involved in the decoration of the polyrhamnose backbone with the GlcNAc side chain. Finally, GacH is the GroP transferase enzyme that cleaves membrane phosphatidylglycerol to release and attach GroP to C-6 of the GlcNAc side chains (8). Interestingly, historic research reported isolates, referred to as “A-variants”, in which GAC lost its characteristic GlcNAc side chain after serial passage in mice and rabbits (17, 18). However, the underlying mechanisms responsible for the loss of the GAC GlcNAc side chain were never reported. Additionally, this A-variant phenotype was not detected among a stock collection of human isolates (18).

With the current availability of whole genome sequences, we aimed to re-examine the concept that strains with a non-canonical GAC may arise in humans. To this end, we first analyzed the genomic sequences of a historic *S. pyogenes* A-variant/parent strain pair to pinpoint the genetic alteration that could explain the loss of the GlcNAc side chain in the evolved A-variant strain. Additionally, we analyzed the genetic variation of *gacA-L* in a collection of 2,044 *S. pyogenes* genomes and identified potential inactivating mutations in *gacH*. We investigated the impact of these mutations on GacH function by complementing a *gacH*-deficient strain with plasmid-expressed truncated *gacH* variants and subsequent detection of GroP by biochemical, phenotypical, and functional analysis.

## Methods

### Plasmids, bacterial strains and culture conditions

All plasmids and *S. pyogenes* strains used in this study for wet-lab experiments are listed in Table S1. *Escherichia coli* (*E. coli*) was grown in Lysogeny Broth (LB, Oxoid) medium or LB agar plates, supplemented with 500 µg/ml erythromycin at 37°C. *S. pyogenes* strains were grown on Todd Hewitt supplemented with 0.5 % yeast (THY, Oxoid) agar plates or in THY broth supplemented with 5 µg/ml erythromycin for the plasmid-complemented strains. No antibiotics were added for wild type strains and the *gacH* knockout. Overnight cultures were grown at 37 °C, without CO_2_, subcultured the next day in fresh THY and grown to mid-exponential growth phase, corresponding to an optical density at 600 nm (OD_600_) of 0.4.

### Antibody ab9191 binding assay

*S. pyogenes* strains were grown to midlog phase in THY (OD_600_ 0.4), centrifuged, resuspended in PBS containing 0.1% Bovine serum albumin (BSA; Sigma) and incubated with 1 µg/ml ab9191, a goat polyclonal Group A carbohydrate antibody (Abcam), for 20 min at 4 °C. After washing, bacteria were resuspended in PBS 0.1% BSA containing Protein G Alexa fluor 488 conjugate (1:1,000, Thermo Fisher Scientific P11065). Fluorescence was analyzed by FACSCanto II Flow Cytometer (BD Bioscience). Per sample, 10,000 gated events were collected and fluorescence was expressed as the geometric mean.

### Whole genome sequence comparison of D315 and D315/87/3

The whole genome sequences of an A-variant/wild type strain pair available from the historical Lancefield streptococcal collection were compared to determine genomic changes underlying the loss of GlcNAc side chain in GAC. The A-variant strain D315/87/3 (*emm*58; (1)) was sequenced by the Genomics Technologies Facility in Lausanne, Switzerland using NovaSeq6000. The raw reads have been uploaded to SRA under the project number <u>PRJNA1234579</u>. Raw reads were processed with Trimmomatic v.039 (19) to remove adaptor sequences and bases of insufficient quality with parameters ILLUMINACLIP:TruSeq3-SE:2:30:10 LEADING:3 TRAILING:3 SLIDINGWINDOW:4:15 MINLEN:36. Trimmed reads were then *de novo* assembled into scaffolds with SPAdes v3.15.5 (20) and assembly quality was assessed with QUAST v5.0.2. Single nucleotide polymorphism (SNP) differences between parent strain D315 (also referred to as NCTC10876; GenBank accession: LS483360) and D315/87/3 were detected with Snippy v4.6.0 (21)(https://github.com/tseemann/snippy) with default parameters.

### Group A *streptococcus* analysis of *gac* gene cluster

Whole genome sequences (Illumina short-read) of 2,044 *S. pyogenes* isolates were uploaded to the open-access PubMLST database (www.pubmlst.org) (14, 22). *gacA*-*gacL* were annotated and screened for genetic variation, where allele 1 was assigned to reference strain MGAS5005 (23). Strains of which one of the *gac* genes was truncated because it was located at the end of a contig (n = 23) were excluded from the analysis. The sequence of allele 1 was blasted against the genomes of strains without an allele number using the PubMLST database to identify sequences with a premature stop codon. Default BLAST settings were used: BLASTN word size:11; BLASTN scoring: reward: 2; penalty: -3; gap open: 5; gap extend: 2; Hits per isolate: 1; Flanking length (bp): 0.

A 14,416 bp region containing each *gac* gene, as well as the intergenic regions upstream of *gacA*, *gacB* and *gacI*, was detected in each isolate with blastn (24) against reference genome MGAS5005 (accession number: CP000017) with default parameters. Alignment start and end region and subject strand information were used to extract sequences with bedtools getfasta (25), reverse complementing when located on the antisense strand. A multiple sequence alignment was constructed with mafft (26) and input into snp-sites to determine polymorphic regions (27). The resulting variant call format file was annotated with snpEff (28) and its pre-built database Streptococcus_pyogenes_mgas5005 to determine synonymous and non-synonymous SNPs of coding regions.

### Generation of complemented *S. pyogenes* Δ*gacH* strains with *gacH* variant

All primers used in this study are listed in Table S2. Expression vector pDCerm_*gacH* (*gacH* allele 1) was isolated from *E. coli* MC1061. The plasmid was digested with EcoRI and BglII. For complementation of the *S. pyogenes* Δ*gacH* knockout strain with the different *gacH* variants, different strategies were used. To create *gacH* complementation plasmids that contained a premature stop codon at nucleotide position 928 and 937, sequences were codon optimized and gblocks were ordered. The gblocks were amplified with primer STOP309AAF/ STOP309AAR or STOP31**2**AAF/STOP31**2**AAR, digested with EcoRI and BglII and ligated into EcoRI/BglII-digested pDCerm. Correct insertion was confirmed by Sanger sequencing with primers pDCermF, pDCermR, *gacH*309_checkF1, *gacH*309_checkR1 or pDCermF, pDCermR, STOP312checkF, STOP312checkR. For complementation with *gacH* containing a premature stop codon at nucleotide position 2320, a gblock could not be generated, even after codon optimization. Therefore, wild type *S. pyogenes* strain 20162146, which was identified to contain this *gacH* stop codon, was obtained from the Centers for Disease Control (CDC, Atlanta, GO, USA). The *gacH* gene was amplified with primers *gacH*EcoRIF and *gacH*BglIIR, followed by a similar procedure as described for the gblocks. The correct insert was confirmed with primers *gacH*check1, *gacH*check2, *gacH*check3, pDCermF and pDCermR. Plasmids with the correct inserts were transformed to *S. pyogenes* 5448Δ*gacH*.

### Measurement of relative phosphate concentrations on GAC

*S. pyogenes* cell wall was isolated from late exponential phase cultures (OD_600_ ∼0.8) by the SDS-boiling procedure as previously described (13, 29). Purified cell wall samples were lyophilized and stored at −20°C before the analysis. GAC was released from cell wall preparations by mild acid hydrolysis as previously described (13). After hydrolysis, samples were purified by running over a PD-10 desalting column (VWR, 17-0851-01), with de-ionized water as the exchange buffer. Phosphate was released from GAC as outlined (8). Briefly, soluble GAC was incubated with 2 N HCl at 100 °C for 2 h to cleave GroP. Samples were neutralized with NaOH in the presence of 62.5 mM HEPES pH 7.5. To release phosphate from GroP, samples (100 μL) were incubated with 2 μL of 1 U/μL alkaline phosphatase (New England Biolabs; quick CIP) in alkaline phosphatase buffer (New England Biolabs; rCutSmart buffer) at 37°C overnight. Phosphate concentrations were measured using the malachite green method. The reactions were diluted to 160 μL with water and 100 μL was transferred to a flat-bottom 96-well culture plate. Malachite Green reagent (0.2 mL) was added and the absorbance at 620 nm was read after 10 min at room temperature. Malachite Green reagent contained one volume 4.2% ammonium molybdate tetrahydrate (by weight) in 4 M HCl, 3 volumes 0.045% malachite green (by weight) in water and 0.01% Tween 20. Phosphate concentrations were determined using a phosphate standard curve. Concentrations of rhamnose (Rha) in GAC were measured by an anthrone assay as previously described (13). The concentrations of phosphate were normalized to total Rha content and presented as a percentage of the ratios in the WT strain.

### Human group IIA-secreted phospholipase A2 killing assay

hGIIa killing was determined as described before (30). Briefly, *S. pyogenes* strains were grown to midlog phase (OD_600_ 0.4), diluted 1,000-fold in HEPES solution (20 mM HEPES, 2 mM Ca^2+^, 1% BSA (pH 7.4)) with or without 0.5 µg/ml hGIIa. Samples were incubated for 2 hours at 37°C and viability was assessed using serial dilutions plating on blood agar plates. The next day, colony forming units (CFU) were counted. Percentage survival was calculated as follows:. 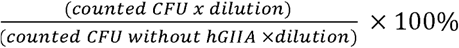

### Zinc susceptibility assay

*S.pyogenes* strains were grown to midlog phase in THY, cultures were diluted in fresh THY to OD_600_ 0.1. An equal volume of 2.5 mM ZnSO_4_ in THY was added in a 96 well plate (technical duplicates) and cultures were incubated at 37°C. After 20 hours, serial dilutions were plated for CFU determination.

### Statistical analysis

Flow cytometry data were analyzed using FlowJo 10 (FlowJo). Data were analyzed using GraphPad Prism 10.2.0 (GraphPad software). Statistical significance was tested using an unpaired t test or a one-way ANOVA followed by a Dunnett’s test for multiple comparison. The *P* values are depicted in the figures or mentioned in the caption, and *P*<0.05 was considered significant.

## Results

We obtained a viable parent strain (D315) and animal-passaged A-variant strain (D315/87/3) from the historical Lancefield streptococcal collection (Table S1). The A-variant phenotype of strain D315/87/3 was confirmed by absence of StrepTex latex agglutination (not shown) and loss of binding of a GAC-GlcNAc reactive polyclonal antibody (Figure 1, Supplementary Figure S1). Next, we compared whole genome sequences of these strains to pinpoint the underlying genetic defect. We identified 141 SNPs, of which 115 were identified in coding regions and were disruptive, i.e. premature stop codon or switch to non-synonymous or frameshift mutations (Supplementary Table S3). Within the *gac* cluster, we identified a nucleotide substitution C382T in the *gacI* sequence of D315/87/3 (A-variant). GacI is the glycosyltransferase encoded by *gacI*, that is critical for expression of the GAC GlcNAc side chain (31, 32). This mutation resulted in a premature stop codon at nt position 382 leading to a truncated protein of 127 amino acids instead of 231, thereby reducing GacI size by 55%, which likely results in a non-functional protein. Therefore, the identified *gacI* mutation likely explains the absence of GlcNAc in the historic mouse-passaged A-variant D315/87/3.

**Figure 1.**
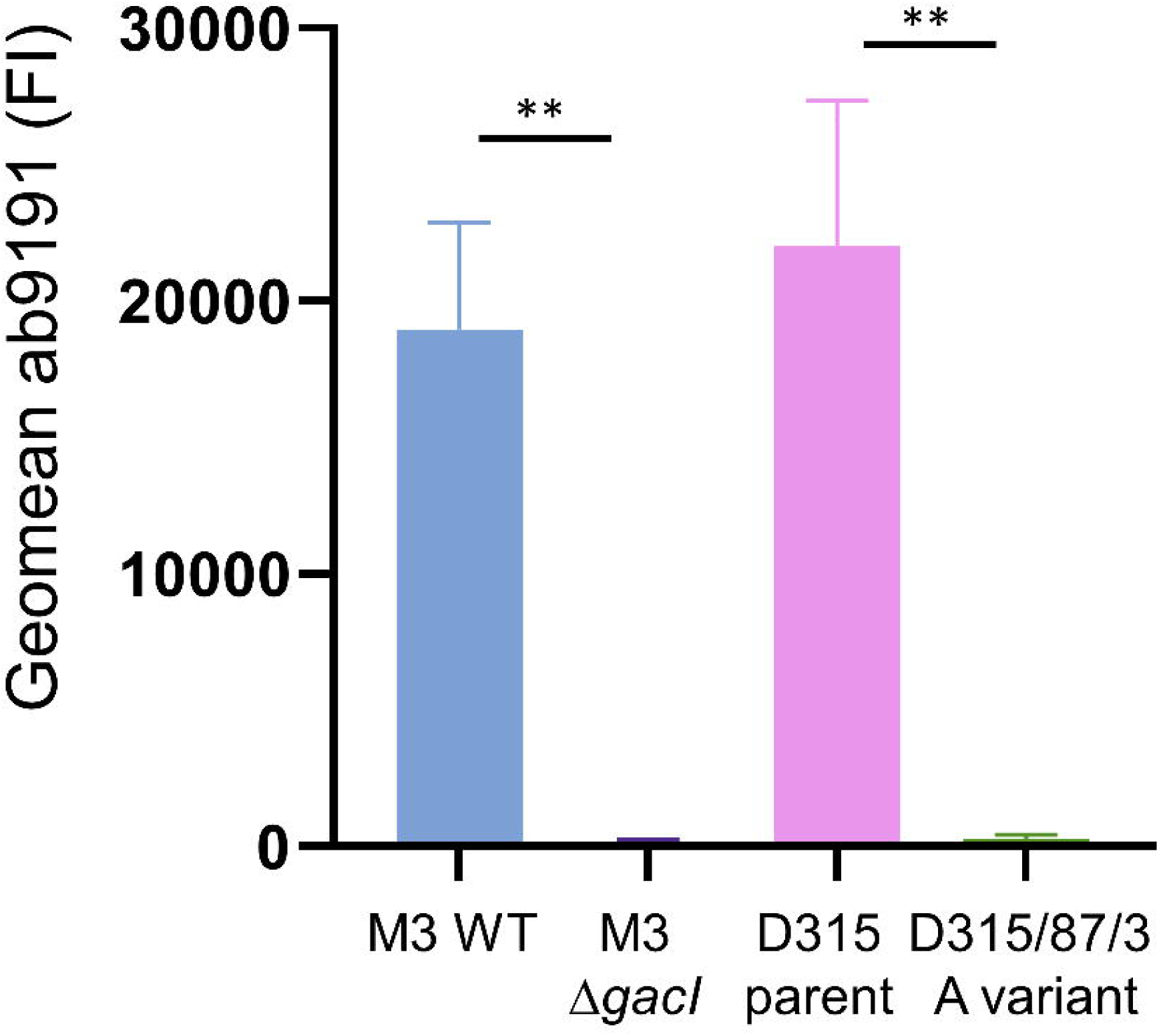
Historical A-variant lacks GlcNAc side chain. Binding of goat polyclonal *Streptococcus pyogenes* Group A carbohydrate antibody ab9191 (1 µg/ml) to wild type (WT) *S. pyogenes* M3, an isogenic *gacI* mutant, the historic A variant (D315/87/3) and its parent strain (D315). Data are depicted as geometric mean fluorescence intensity (FI) of three individually displayed biological replicates (mean + standard deviation). *P* values were calculated by two unpaired t tests; ** *P*<0.01.

To comprehensively analyze the genetic variation in *gacA-L*, we analyzed allelic variation and presence of specific mutations in these genes in 2,021 *S. pyogenes* genomes (Supplementary Table S4) (14). The number of nucleotide alleles varied between 33 (*gacJ*) and 284 (*gacH*) (Table 1). The 14,416 bp region, spanning the *gac* operon and the 137 bp upstream intergenic regions, contained 1,629 single nucleotide polymorphic (SNP) across 1,568 unique *gac* positions with 61 being polymorphic (Supplementary Table S5). Across non-M1 strains, an average of 57 SNPs per *gac* cluster per strain were detected relative to the MGAS5005 reference genome; 796 out of 1,629 (48.9 %) SNPs were non-synonymous; 745 out of 629 (45.7%) SNPs were synonymous and 89/1629 (5.5%) were identified in intergenic regions (Figure 2; Supplementary Table S5). Two promoters for the *gac* gene cluster have been described and confirmed through RNA-sequencing, i.e. one upstream of *gac*A (nt 604738..604788 in MGAS5005) and one upstream of *gac*B (nt 605767-605817 genome MGAS5005 (33). We observed 11 different mutations in the promoter region of *gac*A in 35 (1.7%) out of 2,021 isolates. Only one of these mutations (isolate K41948) was in the predicted -10 region (ATGAAA→ATGAAG). Two mutations were found in the spacer region (isolates NGAS130 and Bra36) and none were found in the -35 region. For the promotor upstream of *gac*B, 10 different mutations in 88 (4.4%) out of 2021 strains were identified. None of these mutations were found in the -10, spacer nor -35 predicted regions. Overall and as expected, the *gac* gene cluster was genetically highly conserved in the *S. pyogenes* population.

**Figure 2.**
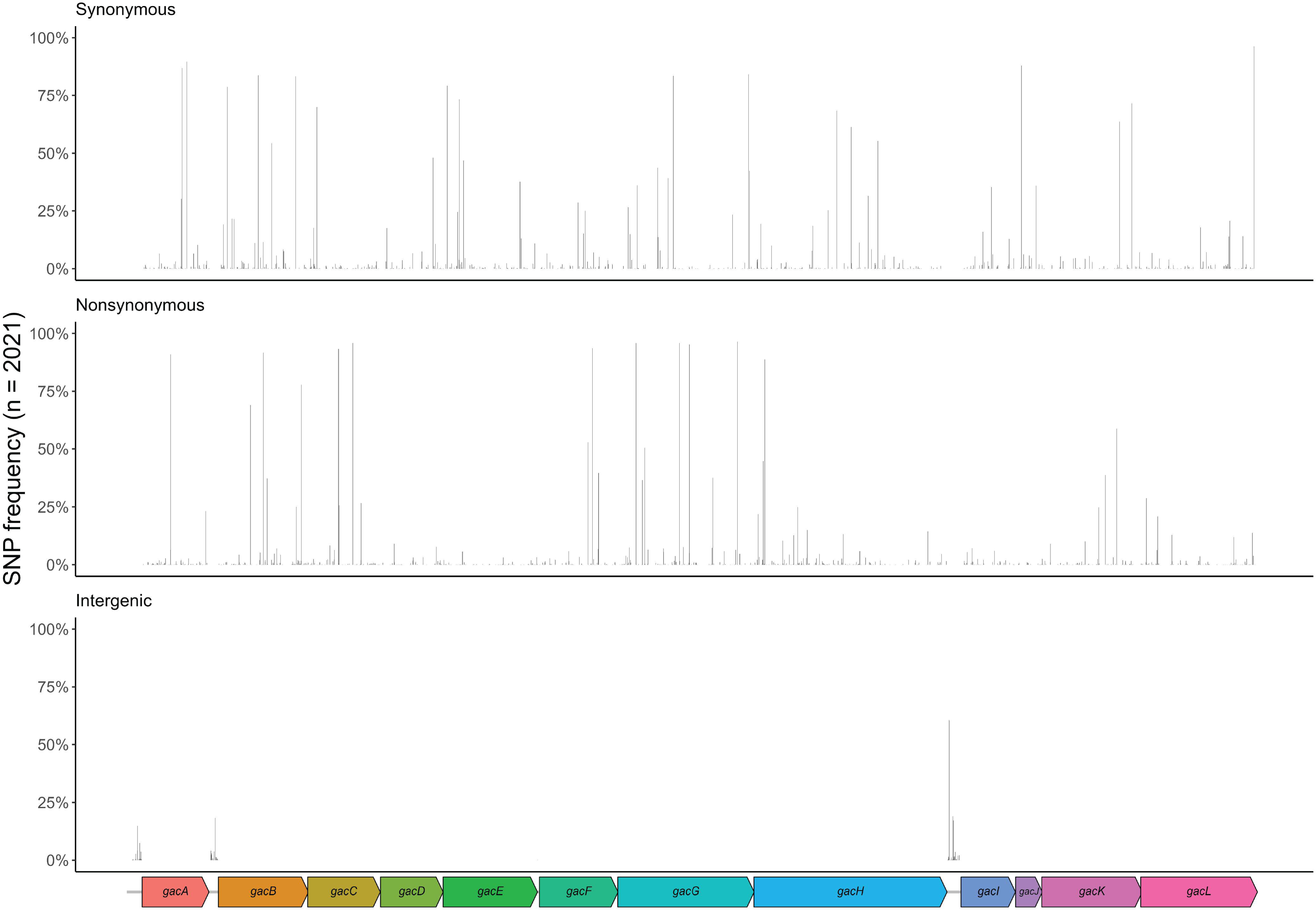
Single nucleotide polymorphic sites of the *gac* operon across 2,021 *S. pyogenes* genome sequences. The 14,416 bp, 12-gene *gac* operon including 137 bp upstream, intergenic regions of reference genome MGAS5005 (accession: CP000017) are represented at the bottom. Relative positions of 1,629 SNPs at 1,568 polymorphic sites are represented by bars across the x-axis, with their height representing its frequency across 2,021 *S. pyogenes* genome sequences. SNPs are categorized into three groups relative to MGAS5005: synonymous mutations resulting in no amino acid change (upper graph); nonsynonymous mutations resulting in an amino acid change (middle graph); and SNPs detected in intergenic regions (bottom graph).

**Table 1.**
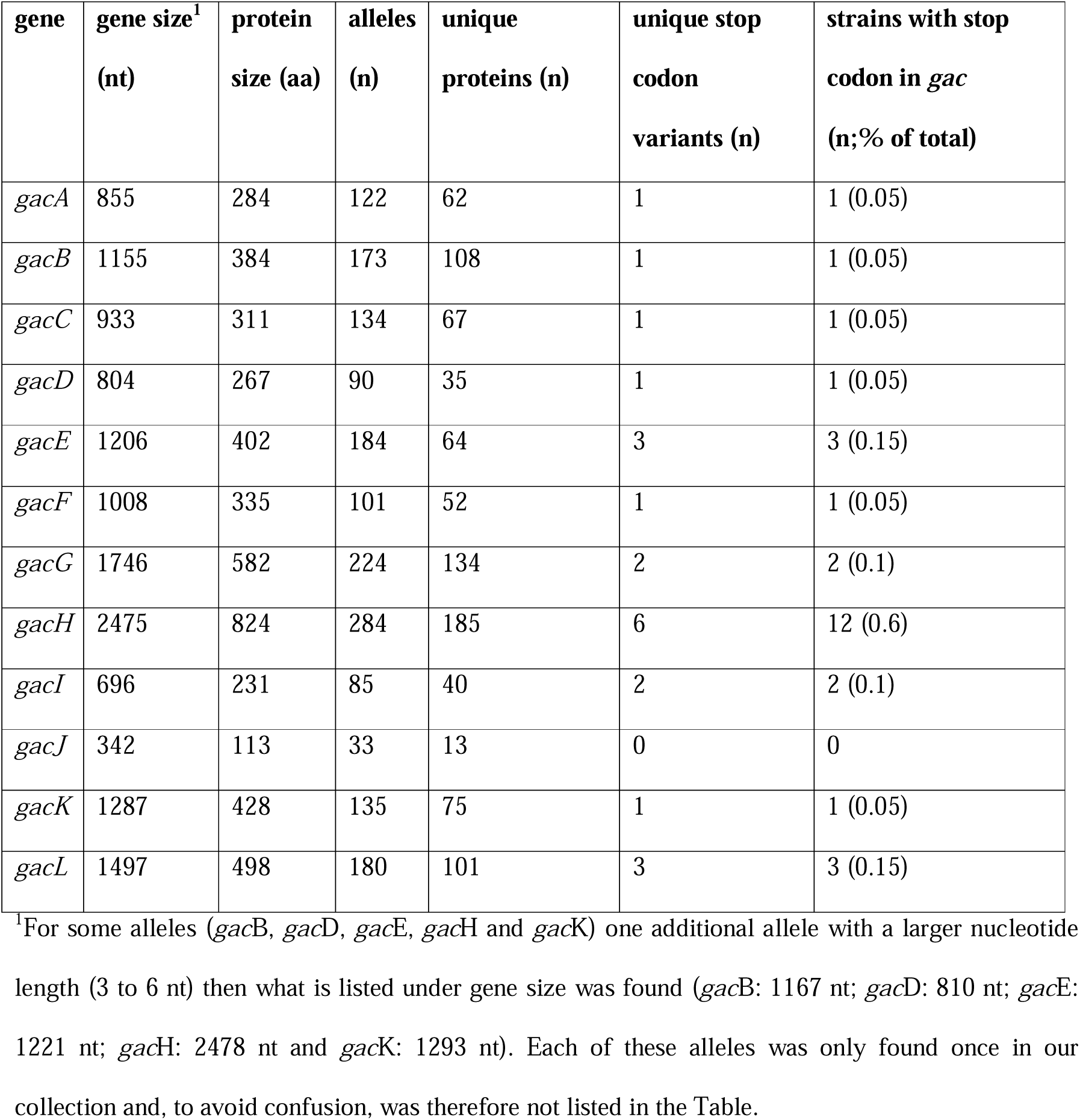
Overview of genetic variation of *gacA-L* in the 2,021 *S. pyogenes* collection. Shaded rows indicate genes implicated in polyrhamnose biosynthesis, white rows indicate genes critical for formation of the GlcNAc-GroP side chain. Allele sequences of the *gac* genes can be downloaded from PubMLST (<u>Download alleles</u>).

Both the number of alleles and unique proteins of each *gac* gene correlated strongly with gene size (Figure 3A, B). *gacH* displayed the highest number of unique protein sequences (Table 1; Figure 3B). Interestingly, for all *gac* genes except *gacJ*, we identified variants that contained a premature stop codon (Table 2; supplementary Table S5). Overall, we identified 26 isolates (1.3% of all analyzed strains) that contained a premature stop codon in one or two of the *gac* genes. Three isolates (NS534, MTB313 and STAB901) were observed to contain a nucleotide deletion in *gacI* at position 449 resulting in a premature stop codon. However, Sanger sequencing *gacI* of STAB901 (34) did not confirm this nucleotide deletion and likely resulted from a sequencing error in a poly(A)-tract of 8 nucleotides. Therefore, we did not include this particular *gacI* ‘stop codon’ in Table 2. In addition, we observed a deletion in *gacA* at position 414 resulting in a premature stop codon at nucleotide position 502 in isolate Manfredo (Table 2). We confirmed the presence of this mutation by Sanger sequencing. Although this deletion predicts a truncated protein consisting of only 167 of the 284 amino acids (59%)(9), this strain is still positive in the rapid test agglutination assay (data not snown), suggesting a functional GacA. None of the other premature stop codons were analyzed functionally due to lack of the specific strains.

**Figure 3.**
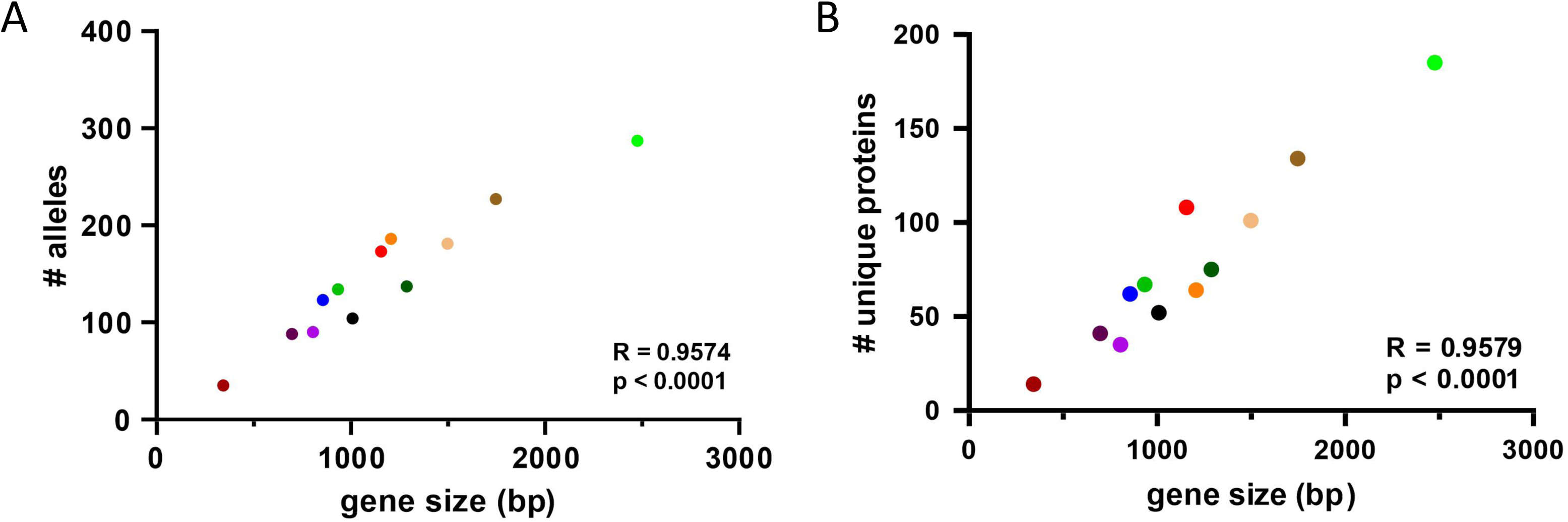
Unique allelic and protein variants of *gacA-L* in 2,021 *S. pyogenes* isolates. The number of (A) unique alleles in and (B) unique protein sequences encoded by each of the *gacA-L* genes and correlation with gene size.

**Table 2.**
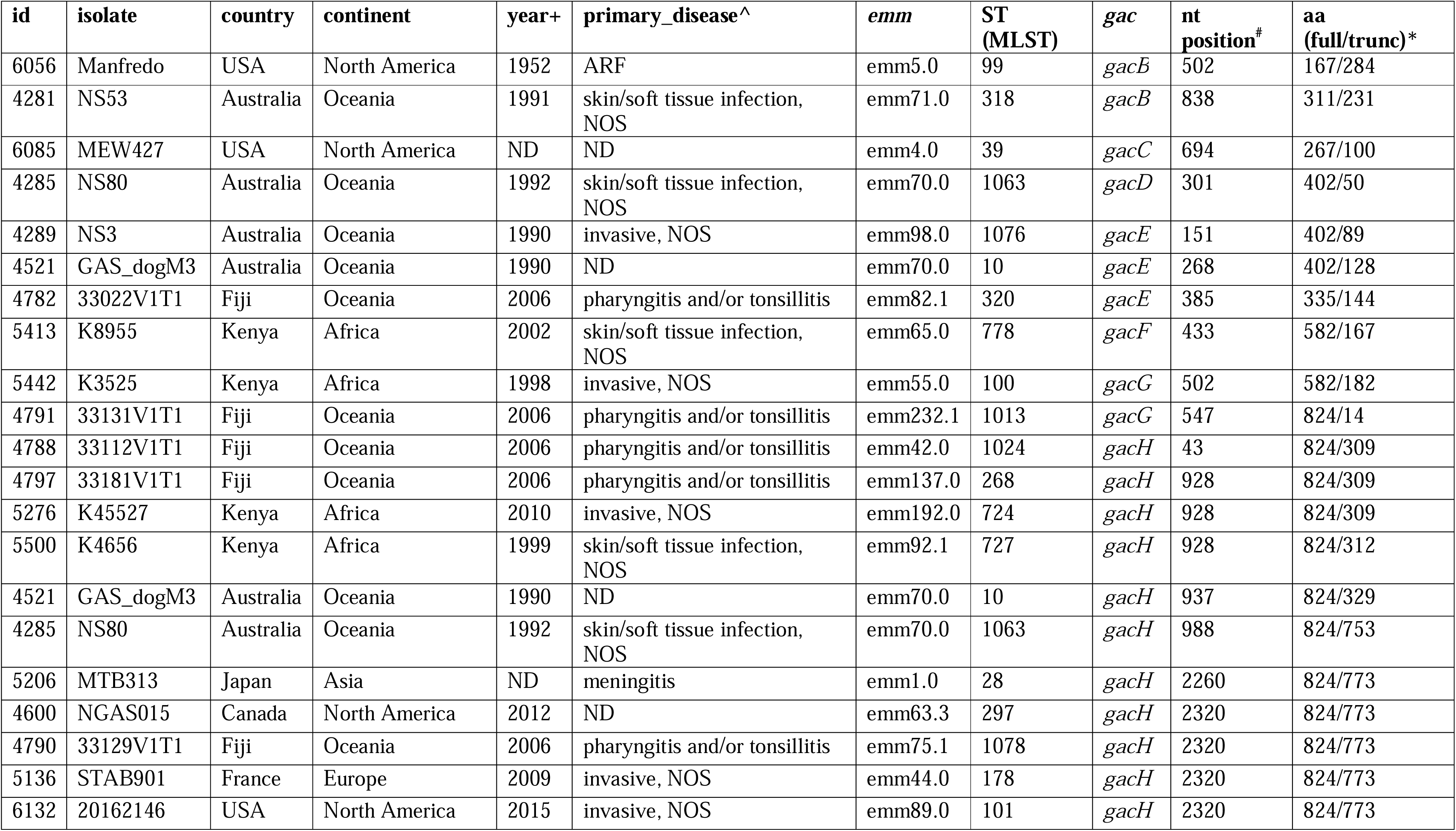
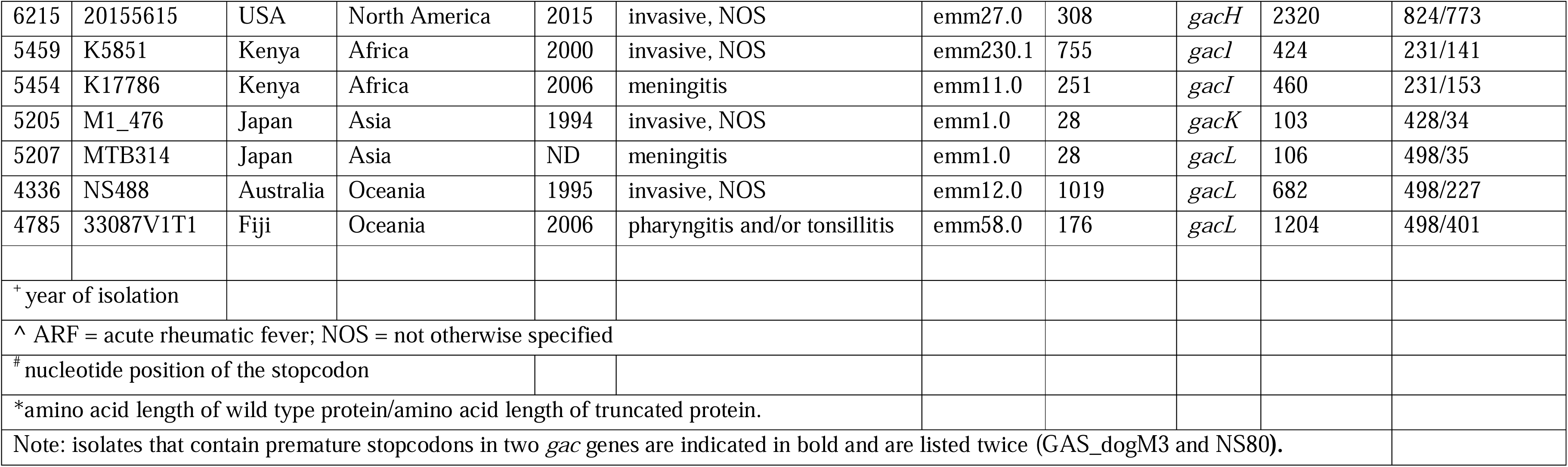
Overview of *S. pyogenes* isolates (n=26) with stop codons in one or two *gac* genes.

| id | isolate | country | continent | year+ | primary_disease^ | emm | ST (MLST) | *gac* | nt position<sup>#</sup> | aa (full/trunc)* |
| --- | --- | --- | --- | --- | --- | --- | --- | --- | --- | --- |
| 6056 | Manfredo | USA | North America | 1952 | ARF | emm5.0 | 99 | *gacB* | 502 | 167/284 |
| 4281 | NS53 | Australia | Oceania | 1991 | skin/soft tissue infection, NOS | emm71.0 | 318 | *gacB* | 838 | 311/231 |
| 6085 | MEW427 | USA | North America | ND | ND | emm4.0 | 39 | *gacC* | 694 | 267/100 |
| 4285 | NS80 | Australia | Oceania | 1992 | skin/soft tissue infection, NOS | emm70.0 | 1063 | *gacD* | 301 | 402/50 |
| 4289 | NS3 | Australia | Oceania | 1990 | invasive, NOS | emm98.0 | 1076 | *gacE* | 151 | 402/89 |
| 4521 | GAS_dogM3 | Australia | Oceania | 1990 | ND | emm70.0 | 10 | *gacE* | 268 | 402/128 |
| 4782 | 33022V1T1 | Fiji | Oceania | 2006 | pharyngitis and/or tonsillitis | emm82.1 | 320 | *gacE* | 385 | 335/144 |
| 5413 | K8955 | Kenya | Africa | 2002 | skin/soft tissue infection, NOS | emm65.0 | 778 | *gacF* | 433 | 582/167 |
| 5442 | K3525 | Kenya | Africa | 1998 | invasive, NOS | emm55.0 | 100 | *gacG* | 502 | 582/182 |
| 4791 | 33131V1T1 | Fiji | Oceania | 2006 | pharyngitis and/or tonsillitis | emm232.1 | 1013 | *gacG* | 547 | 824/14 |
| 4788 | 33112V1T1 | Fiji | Oceania | 2006 | pharyngitis and/or tonsillitis | emm42.0 | 1024 | *gacH* | 43 | 824/309 |
| 4797 | 33181V1T1 | Fiji | Oceania | 2006 | pharyngitis and/or tonsillitis | emm137.0 | 268 | *gacH* | 928 | 824/309 |
| 5276 | K45527 | Kenya | Africa | 2010 | invasive, NOS | emm192.0 | 724 | *gacH* | 928 | 824/309 |
| 5500 | K4656 | Kenya | Africa | 1999 | skin/soft tissue infection, NOS | emm92.1 | 727 | *gacH* | 928 | 824/312 |
| 4521 | GAS_dogM3 | Australia | Oceania | 1990 | ND | emm70.0 | 10 | *gacH* | 937 | 824/329 |
| 4285 | NS80 | Australia | Oceania | 1992 | skin/soft tissue infection, NOS | emm70.0 | 1063 | *gacH* | 988 | 824/753 |
| 5206 | MTB313 | Japan | Asia | ND | meningitis | emm1.0 | 28 | *gacH* | 2260 | 824/773 |
| 4600 | NGAS015 | Canada | North America | 2012 | ND | emm63.3 | 297 | *gacH* | 2320 | 824/773 |
| 4790 | 33129V1T1 | Fiji | Oceania | 2006 | pharyngitis and/or tonsillitis | emm75.1 | 1078 | *gacH* | 2320 | 824/773 |
| 5136 | STAB901 | France | Europe | 2009 | invasive, NOS | emm44.0 | 178 | *gacH* | 2320 | 824/773 |
| 6132 | 20162146 | USA | North America | 2015 | invasive, NOS | emm89.0 | 101 | *gacH* | 2320 | 824/773 |
| 6215 | 20155615 | USA | North America | 2015 | invasive, NOS | emm27.0 | 308 | *gacH* | 2320 | 824/773 |
| 5459 | K5851 | Kenya | Africa | 2000 | invasive, NOS | emm230.1 | 755 | *gacI* | 424 | 231/141 |
| 5454 | K17786 | Kenya | Africa | 2006 | meningitis | emm11.0 | 251 | *gacI* | 460 | 231/153 |
| 5205 | M1_476 | Japan | Asia | 1994 | invasive, NOS | emm1.0 | 28 | *gacK* | 103 | 428/34 |
| 5207 | MTB314 | Japan | Asia | ND | meningitis | emm1.0 | 28 | *gacL* | 106 | 498/35 |
| 4336 | NS488 | Australia | Oceania | 1995 | invasive, NOS | emm12.0 | 1019 | *gacL* | 682 | 498/227 |
| 4785 | 33087V1T1 | Fiji | Oceania | 2006 | pharyngitis and/or tonsillitis | emm58.0 | 176 | *gacL* | 1204 | 498/401 |
| <sup>+</sup> year of isolation |  |  |  |  |  |  |  |  |  |  |
| ^ ARF = acute rheumatic fever; NOS = not otherwise specified |  |  |  |  |  |  |  |  |  |  |
| # nucleotide position of the stopcodon |  |  |  |  |  |  |  |  |  |  |
| *amino acid length of wild type protein/amino acid length of truncated protein. |  |  |  |  |  |  |  |  |  |  |
| Note: isolates that contain premature stopcodons in two *gac* genes are indicated in bold and are listed twice (GAS_dogM3 and NS80). |  |  |  |  |  |  |  |  |  |  |

For 12 (46%) of the 26 strains, a premature stop codon was found in *gacH* (Table 2). These isolates did not cluster based on *emm* typing nor on multi locus sequence typing (MLST). Furthermore, these strains were isolated from different disease manifestations (pharyngitis n=3, skin/soft tissue infection n=2, invasive infection n= 4, meningitis n=1, unknown n=2) and originated from different continents (Oceania n=5, Africa n=2, Asia n=1, North America n=3 and Europe n=1).

GacH modifies the GAC GlcNAc side chain with GroP (8) and consists of a transmembrane domain (Figure 4; amino acids 1 – 395; green) and a catalytic domain on the extracellular site of the membrane (Figure 4; amino acids 444 – 822, red). The premature stop codons were located at nucleotide position 43 (n=1), 928 (n=3), 937 (n=1), 988 (n=1), 2260 (n=1) and 2320 (n=5), resulting in truncated proteins of 14, 309, 312, 329, 753 and 773 amino acids long (Figure 4). We also checked for the presence of mutations that flank the catalytic site as well as the sites that are important for binding ligand and substrate based on structural information in Edgar et al. 2019. We did not identify any amino acid substitutions in these positions indicating that these sites are 100% conserved.

**Figure 4.**
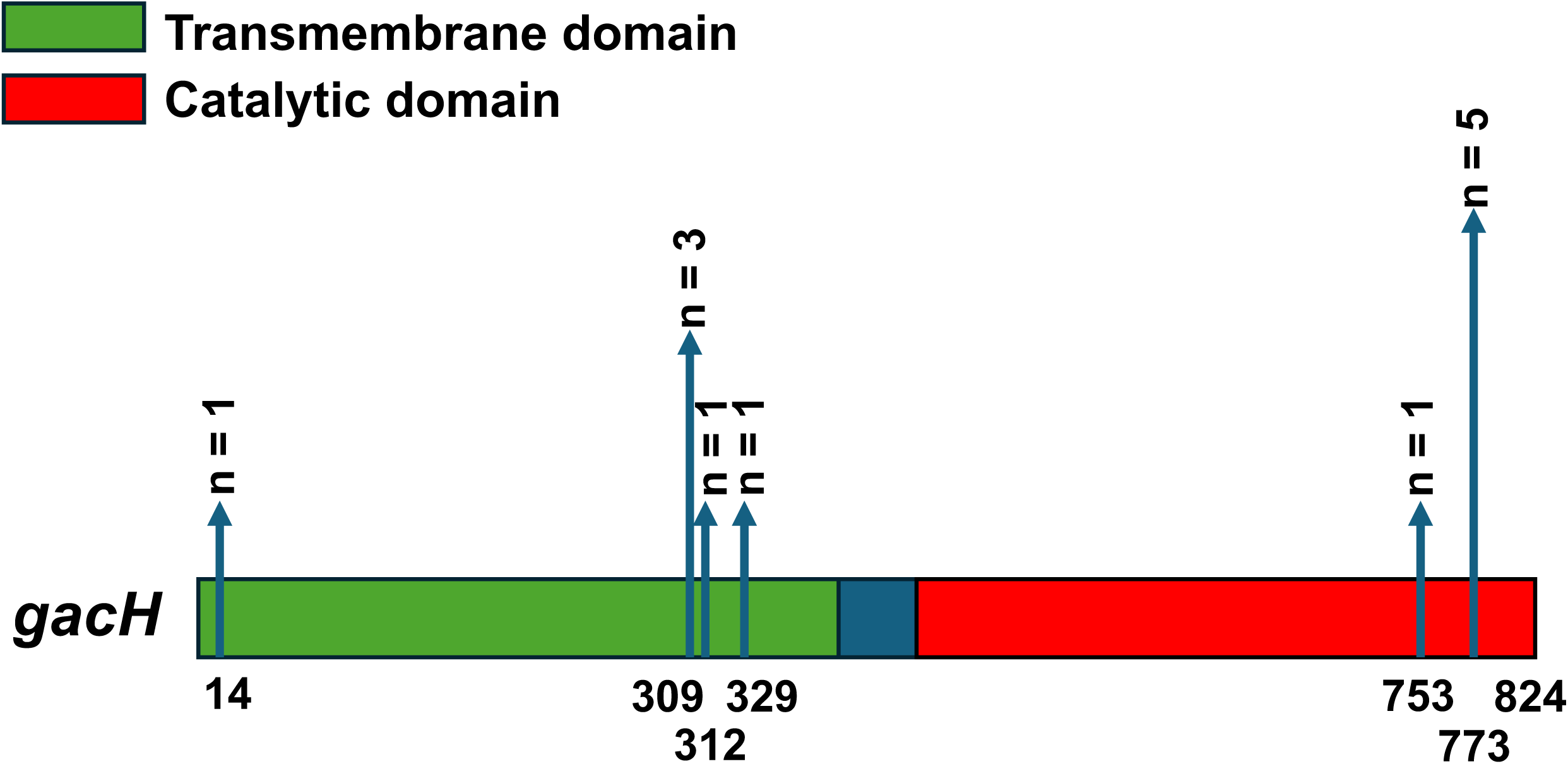
*In silico* analysis of premature stop codons in *gacH*. Scaled 2D representation of *gacH* (2,475 bp, 824 amino acids), containing a transmembrane domain (amino acids 1-395; green) and a catalytic domain (amino acids 444-822; red). Premature stop codons were identified in 12 isolates and are indicated by the vertical blue arrows that show the amino acid position and frequency.

To study the functional consequences of the premature stop codons in *gacH*, we expressed three *gacH* variants (stop codon at nt position 928, 937, and 2320) on a plasmid in a *S. pyogenes* mutant lacking *gacH* (5448Δ*gacH*). Additionally, we included a wild-type strain (20162146) that contained the naturally-occurring premature stop codon in *gacH* at amino acid position 773. To determine the presence of GroP on the GAC GlcNAc side chain, we measured the phosphate content in the isolated GAC in the different *S. pyogenes* strains (8). As expected, GAC isolated from 5448Δ*gacH* contained a significantly reduced amount of phosphate compared to WT 5448 GAC (Figure 5A). This phenotype could be restored by complementation with a plasmid containing WT *gacH*, but not with *gacH* variants encoding GacH variants truncated from amino acid position positions 309, 312 or 773 (Figure 5A). Similarly, the WT strain 20162146, which naturally acquired the premature stop codon at position 2320 in *gacH*, showed strongly reduced levels of phosphate in the isolated GAC (Figure 5A).

**Figure 5.**
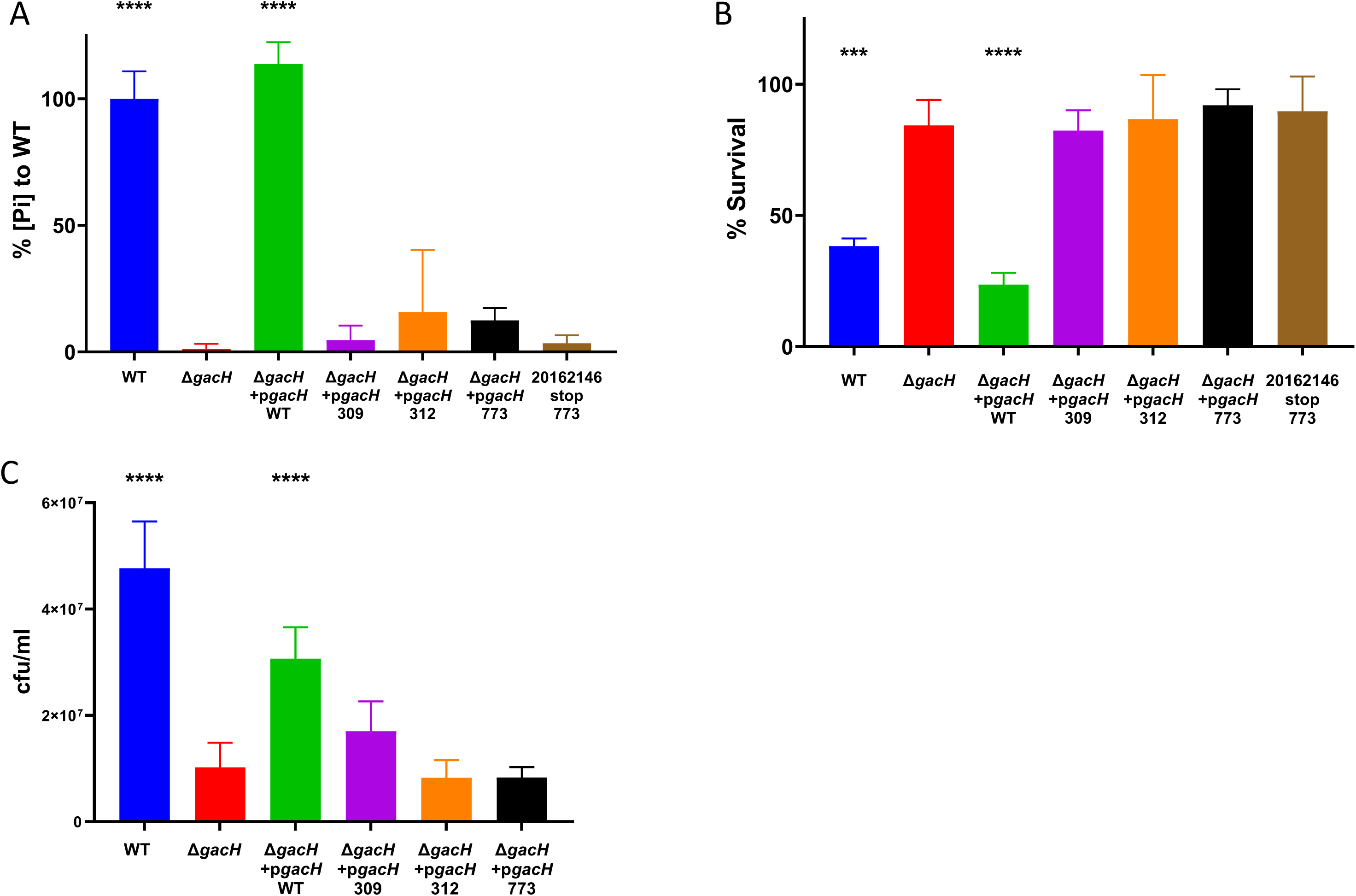
Biochemical and functional analysis of *gacH* variants with premature stop codon in *S. pyogenes*. (A) Analysis of phosphate content in GAC isolated from WT 5448 *S. pyogenes*, 5448Δ*gacH*, and 5448Δ*gacH* complemented with plasmid-expressed WT *gacH*, premature stop codon *gacH* (on positions 309, 312 or 773) or WT *S. pyogenes* that naturally acquired a premature stop codon in *gacH* (strain 20162146). The concentrations of phosphate are relative to *S. pyogenes* WT 5448. Bars and error bars represent the mean relative phosphate concentrations measured in three biological replicates and the standard deviation, respectively. Survival of all strains mentioned in (A) after exposure to (B) 0.5 μg/mL recombinant hGIIA or (C) 1.25 mM Zn2+. Bars and error bars represent the mean percentage survival and standard deviation, respectively (n=3 biological replicates). *P* values were calculated by one-way analysis of variance (ANOVA) comparing all strains to 5448Δ*gacH*; *** *P*<0.001, **** *P*<0.0001.

The presence of GroP confers susceptibility of *S. pyogenes* to the bactericidal enzyme hGIIa and resistance to zinc toxicity (8). To confirm the functional implications of GroP loss in the complemented strains expressing *gacH* variants, we determined bacterial survival of these *gacH* variants in a hGIIa-killing assay and zinc susceptibility assay. Similar to the 5448Δ*gacH* mutant, all premature stop codon variants displayed resistance towards the bactericidal enzyme hGIIa and increased susceptibility to zinc compared to the isolate that plasmid-expressed wild-type *gacH* (Figure 5B, C). Overall, both the biochemical and functional assay confirmed that *gacH* variants with an early stop codon in the gene resulting in truncated proteins of 309 and 312 amino acids long and even a premature stop codon close to the C-terminus (resulting in a protein of 773 amino acids instead of 824) were defective in modifying of GAC with GroP.

## Discussion

GAC is a major and characteristic cell wall component of *S. pyogenes* and plays important roles in bacterial physiology and pathogenesis. Functional and structural analysis of the GAC has been performed for a few well-known *S. pyogenes* laboratory strains, where deletion of *gacI* results in reduced survival in human blood and animal models of systemic infection (15, 32). Furthermore, removal of *gacH* renders *S. pyogenes* susceptible to zinc and resistant to the host cationic antimicrobial peptides including hGIIA (8). Here, we identified that a mutation in *gacI*, resulting in a premature stop codon, likely underlies the historical A-variant phenotype that has lost the characteristic GAC GlcNAc side chain after frequent animal passage. Furthermore, by analyzing 2,021 *S. pyogenes* genomes, we identified a small number of clinical *S. pyogenes* isolates that contain a *gacH* allele with a premature stop codon. By expressing these allelic variants in a *gacH*-deficient strain, we demonstrated that these genetic variants are severely attenuated in their enzymatic activity, resulting in loss of the GroP side chain and acquiring resistance to hGIIA but susceptibility to zinc.

In this study, we analyzed a geographically and clinically diverse collection of *S. pyogenes* genome sequences, comprising 150 different *emm* types and 484 MLST types (14), to obtain a more comprehensive overview of the variability of the *gac* genes across the *S. pyogenes* population. We uploaded the genome sequences to PubMLST, which comprises an important freely-accessible tool for *S. pyogenes* research on the presence and genetic variation of specific genes at the population level (22). Our results thereby expand observations from previous work, where variation in *gac* genes was analyzed in 520 of these 2,044 *S. pyogenes* strains (15, 35). In these studies, the average number of SNPs was about 2-fold lower in the *gac* gene cluster (1 bp per 260 bp) compared to the core genome (1 bp per 133 bp) of the analyzed strains, suggesting negative selection. Negative selection was also implied by the ratio of nonsynonymous to synonymous SNPs (dN/dS) of the *gac* operon, which was 0.24 (14). In line with these studies, SNP analysis of the strain collection analyzed here observed an average of 57 SNPs, equivalent to 1 polymorphism per 260 bp, per *gac* cluster per genome. Although we did not carry out additional dN/dS analyses, our data also strongly suggest that sequence variation of the *gac* gene cluster is subject to negative selection, confirming the biological significance of the encoded biosynthesis machinery and GAC itself.

Similar to our study, the study of Henningham et al (15) also reported the existence of strains with premature stop codons in the *gac* genes although these variants were functionally confirmed. In the expanded data set, we identified 10 isolates with premature stop codons in genes *gacA-gacG*, which are critical for biosynthesis of the polyrhamnose backbone. In addition, 16 isolates were identified that contained a premature stop codon in genes that are critical for decoration of polyrhamnose with GlcNAc-GroP (*gacH*-*gacL*). Twelve (75%) of these 16 isolates contained a premature stop codon in *gacH* had a unique MLST profile, and, with one exception, a unique *emm* type.

We aimed to assess the functional consequences of some *gac* stop codons. Indeed, strain Manfredo contained a premature stop codon in *gacA* but still agglutinated in the diagnostic latex agglutination test, suggestion a functional GacA protein. However, for a mutation in *gacI* that introduced a premature stop codon, we could not confirm a defect in GAC GlcNAc expression. Upon reexamination of the *gacI* sequence by Sanger sequencing, we were not able to confirm the mutation. This is likely related to the presence of a poly-A stretch, which could easily result in a sequencing error. Therefore, we advise caution when interpreting the presence of premature stop codons from whole genome sequence data without functional validation.

We further analyzed the *gacH* premature stop codons for functional consequences to GAC biosynthesis. In five of the 12 isolates, the premature stop codon resulted in a GacH protein of 773 amino acids. Three of these strains were isolated from different pathologies and different continents, suggesting the acquisition of this premature stop codon evolved independently. Despite preserving ∼94% of the protein, the stop codon at position 2320 resulted in the loss of GroP, suggesting that the last 50 amino acids of the C-terminus of GacH are crucial for its enzymatic activity.

In addition to gene sequence variation, environmental conditions may also affect *gacI* and *gacH* expression or enzymatic activity. Historically, it was reported that GlcNAc is present in a 1:2 ratio to the rhamnose backbone (6, 31, 36). Furthermore, approximately 25% of GlcNAc residues contains a GroP group (8). Possibly, the enzymes implicated in biosynthesis of the GlcNAc-GroP epitope exhibit different expression levels or activities under different environmental conditions. Whether and how the amount of GlcNAc and GroP present on GAC is regulated remains to be determined.

In conclusion, the *gacA-L* gene cluster is highly conserved in presence and genetic sequence. We showed that there are few exceptions in which *gac* genes are present but contained a premature stop codon. For three of these *gacH* stop codon variants, we confirmed that the GAC-GroP modification is lost, which resulted in increased resistance to hGIIA. The high conservation of *gac* genes and sequence in the *S. pyogenes* population highlights the essential nature of this molecule for streptococcal survival in the human host. Nevertheless, expression could vary due to transcriptional or post-transcriptional regulation. Understanding these regulatory mechanisms would provide insight into disease pathogenesis given the importance of the GlcNAc-GroP of GAC for host immune interaction

## Abbreviations

S. pyogenes: Streptococcus pyogenes
GAC: group A carbohydrate
GlcNAc: N-acetylglucosamine
GroP: glycerol phosphate
hGIIa: human group IIA-secreted phospholipase A2.

## Data summary

All *S. pyogenes* genome sequences used for this analysis are available within the publication by Davies *et al.* (2019), ‘Atlas of group A streptococcal vaccine candidates compiled using large-scale comparative genomics’ Nature Genetics, 51(6):1035-43

The raw reads for A-variant strain D315/87/3 (*emm*58; (1)) are available in SRA under the project number <u>PRJNA1234579</u>. The complete genome sequence of *S. pyogenes* strain D315 (also referred to as NCTC10876) is available through GenBank accession: LS483360.

## References

1. Scott JR, Pulliam WM, Hollingshead SK, Fischetti VA. Relationship of M protein genes in group A streptococci. Proc Natl Acad Sci U S A. 1985;82(6):1822–6.

2. Amy Sims Sanyahumbi SC, Rosemary Wyber, and Jonathan R. Carapetis. Global Disease Burden of Group A Streptococcus2016.

3. Carapetis JR, Steer AC, Mulholland EK, Weber M. The global burden of group A streptococcal diseases. Lancet Infect Dis. 2005;5(11):685–94.

4. Walker MJ, Barnett TC, McArthur JD, Cole JN, Gillen CM, Henningham A, et al. Disease manifestations and pathogenic mechanisms of Group A Streptococcus. Clin Microbiol Rev. 2014;27(2):264–301.

5. Lancefield RC. The Antigenic Complex of Streptococcus Haemolyticus: II. Chemical and Immunological Properties of the Protein. J Exp Med. 1928;47(3):469–80.

6. McCarty M. The lysis of group A hemolytic streptococci by extracellular enzymes of *Streptomyces albus.* II. Nature of the cellular substrate attacked by the lytic enzymes. J Exp Med. 1952;96(5):569–80.

7. Uwe C. Kreis VV, B.Mario Pinto. Application of two-dimensional NMR spectroscopy and molecular dynamics simulations to the conformational analysis of oligosaccharides corresponding to the cell-wall polysaccharide of Streptococcus group A. International Journal of Biological Macromolecules. 1995:117–30.

8. Edgar RJ, van Hensbergen VP, Ruda A, Turner AG, Deng P, Le Breton Y, et al. Discovery of glycerol phosphate modification on streptococcal rhamnose polysaccharides. Nat Chem Biol. 2019;15(5):463–71.

9. van der Beek SL, Le Breton Y, Ferenbach AT, Chapman RN, van Aalten DM, Navratilova I, et al. GacA is essential for Group A Streptococcus and defines a new class of monomeric dTDP-4-dehydrorhamnose reductases (RmlD). Mol Microbiol. 2015;98(5):946–62.

10. van der Beek SL, Zorzoli A, Canak E, Chapman RN, Lucas K, Meyer BH, et al. Streptococcal dTDP-L-rhamnose biosynthesis enzymes: functional characterization and lead compound identification. Mol Microbiol. 2019;111(4):951–64.

11. Nina J. Gao ERL, Victor Nizet. Immunobiology of the Classical Lancefield Group A Streptococcal Carbohydrate Antigen. 2021.

12. van Sorge NM, Cole JN, Kuipers K, Henningham A, Aziz RK, Kasirer-Friede A, et al. The classical lancefield antigen of group A Streptococcus is a virulence determinant with implications for vaccine design. Cell Host Microbe. 2014;15(6):729–40.

13. Rush JS, Parajuli P, Ruda A, Li J, Pohane AA, Zamakhaeva S, et al. PplD is a de-N-acetylase of the cell wall linkage unit of streptococcal rhamnopolysaccharides. Nat Commun. 2022;13(1):590.

14. Davies MR, McIntyre L, Mutreja A, Lacey JA, Lees JA, Towers RJ, et al. Atlas of group A streptococcal vaccine candidates compiled using large-scale comparative genomics. Nat Genet. 2019;51(6):1035–43.

15. Henningham A, Davies MR, Uchiyama S, van Sorge NM, Lund S, Chen KT, et al. Virulence Role of the GlcNAc Side Chain of the Lancefield Cell Wall Carbohydrate Antigen in Non-M1-Serotype Group A Streptococcus. mBio. 2018;9(1).

16. Zorzoli A, Meyer BH, Adair E, Torgov VI, Veselovsky VV, Danilov LL, et al. Group A, B, C, and G Streptococcus Lancefield antigen biosynthesis is initiated by a conserved alpha-d-GlcNAc-beta-1,4-l-rhamnosyltransferase. J Biol Chem. 2019;294(42):15237–56.

17. Wilson AT. Loss of Group Carbohydrate During Mouse Passages of a Group A Hemolytic Streptococcus. J Exp Med. 1945.

18. McCarty M, Lancefield R.C. . Variation in the group-specific carbohydrate of group A streptococci. I. Immunochemical studies on the carbohydrates of variant strains. J Exp Med. 1955.

19. Bolger AM, Lohse M, Usadel B. Trimmomatic: a flexible trimmer for Illumina sequence data. Bioinformatics. 2014;30(15):2114–20.

20. Bankevich A, Nurk S, Antipov D, Gurevich AA, Dvorkin M, Kulikov AS, et al. SPAdes: a new genome assembly algorithm and its applications to single-cell sequencing. J Comput Biol. 2012;19(5):455–77.

21. Seeman T. Snippy: fast bacterial variant calling from NGS reads [Internet]. 2015.

22. Jolley KA, Bray JE, Maiden MCJ. Open-access bacterial population genomics: BIGSdb software, the PubMLST.org website and their applications. Wellcome Open Res. 2018;3:124.

23. Paul Sumby SFP, Andres G. Madrigal, Kent D. Barbian, Kimmo Virtaneva, Stacy M. Ricklefs,, Daniel E. Sturdevant MRG, Jaana Vuopio-Varkila, Nancy P. Hoe, and James M. Musser. Evolutionary Origin and Emergence of a Highly Successful Clone of Serotype M1 Group A Streptococcus Involved Multiple Horizontal Gene Transfer Events. J Infect Dis. 2005;192(5):771–82.

24. Camacho C, Coulouris G, Avagyan V, Ma N, Papadopoulos J, Bealer K, et al. BLAST+: architecture and applications. BMC Bioinformatics. 2009;10(1):421.

25. Quinlan AR, Hall IM. BEDTools: a flexible suite of utilities for comparing genomic features. Bioinformatics. 2010;26(6):841–2.

26. Katoh K, Standley DM. MAFFT Multiple Sequence Alignment Software Version 7: Improvements in Performance and Usability. Molecular Biology and Evolution. 2013;30(4):772–80.

27. Page AJ, Taylor B, Delaney AJ, Soares J, Seemann T, Keane JA, et al. SNP-sites: rapid efficient extraction of SNPs from multi-FASTA alignments. Microb Genom. 2016;2(4):e000056.

28. Cingolani P, Platts A, Wang LL, Coon M, Nguyen T, Wang L, et al. A program for annotating and predicting the effects of single nucleotide polymorphisms, SnpEff. Fly. 2012;6(2):80–92.

29. Bui NK, Eberhardt A, Vollmer D, Kern T, Bougault C, Tomasz A, et al. Isolation and analysis of cell wall components from *Streptococcus pneumonia*e. Anal Biochem. 2012;421(2):657–66.

30. van Hensbergen VP, Movert E, de Maat V, Luchtenborg C, Le Breton Y, Lambeau G, et al. Streptococcal Lancefield polysaccharides are critical cell wall determinants for human Group IIA secreted phospholipase A2 to exert its bactericidal effects. PLoS Pathog. 2018;14(10):e1007348.

31. Rush JS, Edgar RJ, Deng P, Chen J, Zhu H, van Sorge NM, et al. The molecular mechanism of N-acetylglucosamine side-chain attachment to the Lancefield group A carbohydrate in *Streptococcus pyogenes*. J Biol Chem. 2017;292(47):19441–57.

32. van Sorge NM, Cole JN, Kuipers K, Henningham A, Aziz RK, Kasirer-Friede A, et al. The classical lancefield antigen of Group A Streptococcus is a virulence determinant with implications for vaccine design. Cell Host Microbe. 2014;15(6):729–40.

33. Rosinski-Chupin I, Sauvage E, Fouet A, Poyart C, Glaser P. Conserved and specific features of *Streptococcus pyogenes* and *Streptococcus agalactiae* transcriptional landscapes. BMC Genomics. 2019;20(1):236.

34. Soriano N, Vincent P, Piau C, Moullec S, Gautier P, Lagente V, et al. Complete Genome Sequence of *Streptococcus pyogenes* M/emm44 Strain STAB901, Isolated in a Clonal Outbreak in French Brittany. Genome Announc. 2014;2(6).

35. Seale AC, Davies MR, Anampiu K, Morpeth SC, Nyongesa S, Mwarumba S, et al. Invasive Group A Streptococcus Infection among Children, Rural Kenya. Emerg Infect Dis. 2016;22(2):224–32.

36. McCarty M. Variation in the group-specific carbohydrate of group A streptococci. II. Studies on the chemical basis for serological specificity of the carbohydrates. J Exp Med. 1956;104(5):629–43.

