## Supplemental information for "Natural variation of the streptococcal Group A carbohydrate biosynthesis genes impacts host-pathogen interaction"

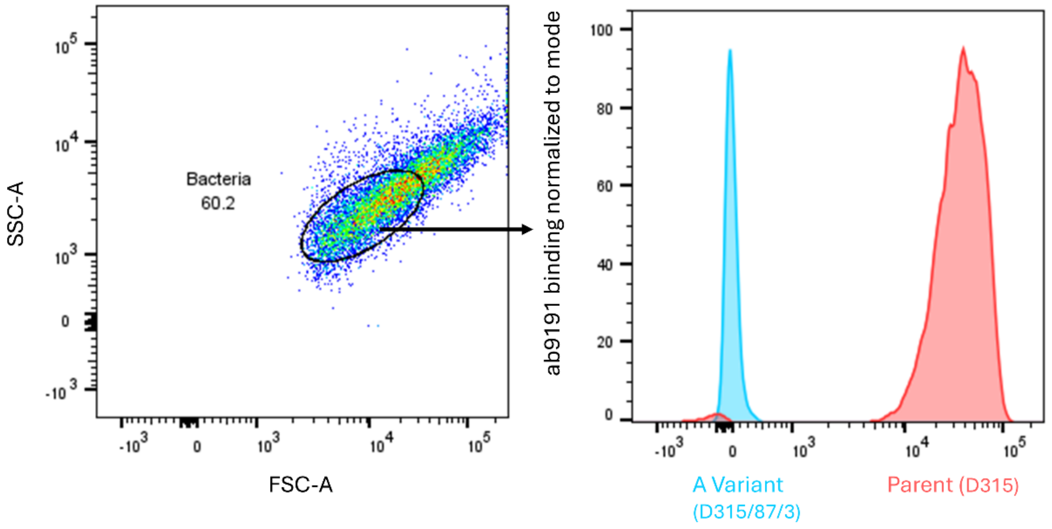


**Figure S1 Gating strategy for ab9191 binding to *S. pyogenes* parent and A-variant strain**

Representable forward- and side scatter with bacteria gate (left), and histograms of ab9191 (protein G-AF488) binding to A-variant (D315/87/3, animal-passaged; turquoise) strain and the parent (D315, wild-type; red) strain (right).

**Table S1.** Bacterial strains and plasmids used in this study.

| Strain | Source | Information |
|---|---|---|
| *S. pyogenes* (M3) | (1) | |
| *S. pyogenes* (M3) Δ*gacI* | (1) | |
| *S. pyogenes* D315 (M58) (NCTC10876; R67/3884) | (2, 3). Obtained from UK Health Security Agency | Wild type |
| *S. pyogenes* D315/87/3 | (4). Provided by Dr. D. Bessen, New York Medical College | 'A variant', passaged in mice |
| *E. coli* MC1061 pDCerm_*gacH* | (5) | |
| *S. pyogenes* 5448 | (5) | |
| *S. pyogenes* 5448 Δ*gacH* | (5) | |
| *S. pyogenes* 5448 Δ*gacH* + pDCerm_*gacH* | (5) | *gacH* from 5448 |
| *S. pyogenes* 5448 Δ*gacH* + pDCerm_*gacH*_STOP309 | This study | |
| *S. pyogenes* 5448 Δ*gacH* + pDCerm_*gacH*_STOP312 | This study | |
| *S. pyogenes* 5448 Δ*gacH* + pDCerm_*gacH*_STOP773 | This study | *gacH* from 20162146 |
| *S. pyogenes* 20162146 | CDC, (6) | Wildtype but *gacH* stop codon at position 773 |

**Table S2.** Primers used in this study

| Primer | Sequence (5' to 3') |
|---|---|
| STOP309AAF | CCGGAATTCATGATTAAAGATACTTTCCTTAAGACTAAC |
| STOP309AAR | GAAGATCTTCAGCGTGAGATTTTAAAGAAGGTCTTGTG |
| gacH309_checkF1 | CTTAGTTGCCTATTTCGCAATCG |
| gacH309_checkR1 | CTCTCAGACGGATTGATCAATG |
| STOP312AAF | CCGGAATTCATGATTAAGGACACATTTTTAAAAACCAATTG |
| STOP312AAR | GAAGATCTTCAGCGTGAGATTTTAAAGAAGGTCTTGTG |
| STOP312checkF | GTGGCCTACTTTGCTATTG |
| STOP312checkR | GATCACGCTAAAGAATTGAG |
| gacHEcoRIF | CCGGAATTCATGATTAAAGACACATTTTTAAAAACCAAT |
| gacHBglIIR | GAAGATCTTTAACGTGATATCTTAAAAAAAGTTTTGTGT |
| gacHcheck1 | CTGAATTAGTAACAGCAGGTAATAG |
| gacHcheck2 | GCTAGGTACAAATCTTTAGC |
| gacHcheck3 | GGTAGCTTGGTATCTACATG |
| pDCermF | GTACCGTTACTTATGAGCAAG |
| pDCermR | CACGGCATAAATCGCTCAG |

#### References


1. Henningham A, Davies MR, Uchiyama S, van Sorge NM, Lund S, Chen KT, et al. Virulence Role of the GlcNAc Side Chain of the Lancefield Cell Wall Carbohydrate Antigen in Non-M1-Serotype Group A Streptococcus. mBio. 2018;9(1).

2. Parker MT, Bassett DC, Maxted WR, Arneaud JD. Acute glomerulonephritis in Trinidad: serological typing of group A streptococci. J Hyg (Lond). 1968;66(4):657-75.

3. Fox EN. M proteins of group A streptococci. Bacteriol Rev. 1974;38(1):57-86.

4. Scott JR, Pulliam WM, Hollingshead SK, Fischetti VA. Relationship of M protein genes in group A streptococci. Proc Natl Acad Sci U S A. 1985;82(6):1822-6.

5. Edgar RJ, van Hensbergen VP, Ruda A, Turner AG, Deng P, Le Breton Y, et al. Discovery of glycerol phosphate modification on streptococcal rhamnose polysaccharides. Nat Chem Biol. 2019;15(5):463-71.

6. Chochua S, Metcalf BJ, Li Z, Rivers J, Mathis S, Jackson D, et al. Population and Whole Genome Sequence Based Characterization of Invasive Group A Streptococci Recovered in the United States during 2015. mBio. 2017;8(5).

**Supplementary Table S3. Overview of genetic changes between parent strain D315 and A-variant derivative D315/87/3**

| CHROM | POS | TYPE | REF | ALT | FTYPE | STRAND | EFFECT | Nucleotide substitution | Amino acid substitution | LOCUS_TAG | GENE | PRODUCT |
|---|---|---|---|---|---|---|---|---|---|---|---|---|
| LS483360 | 6371 | snp | T | G | CDS | + | missense_variant | c.1366T>G | p.Leu456Val | NCTC10876_00006 | mfd | transcription-repair coupling factor |
| LS483360 | 6585 | snp | C | A | CDS | + | stop_gained | c.1580C>A | p.Ser527* | NCTC10876_00006 | mfd | transcription-repair coupling factor |
| LS483360 | 45225 | snp | C | A | CDS | + | missense_variant | c.1216C>A | p.Leu406Ile | NCTC10876_00059 | purH | IMP cyclohydrolase / Phosphoribosylaminoimidazolecarboxamide formyltransferase |
| LS483360 | 49840 | snp | C | A | CDS | + | stop_gained | c.828C>A | p.Cys276* | NCTC10876_00063 | purK | phosphoribosylaminoimidazole carboxylase ATPase subunit |
| LS483360 | 59414 | del | AAACCAGCTACTTGGATTG | A | CDS | + | disruptive_inframe_deletion | c.947_964delAACCAGCTACTTGGATTG | p.Lys316_Ala322delinsThr | NCTC10876_00072 | adhE | Alcohol dehydrogenase |
| LS483360 | 59466 | snp | C | A | CDS | + | missense_variant | c.998C>A | p.Thr333Asn | NCTC10876_00072 | adhE | Alcohol dehydrogenase |
| LS483360 | 59485 | snp | A | T | CDS | + | missense_variant | c.1017A>T | p.Glu339Asp | NCTC10876_00072 | adhE | Alcohol dehydrogenase |
| LS483360 | 136470 | snp | A | G | CDS | - | missense_variant | c.644T>C | p.Val215Ala | NCTC10876_00164 | | sensory box protein |
| LS483360 | 138536 | snp | C | T | CDS | + | missense_variant | c.848C>T | p.Thr283Ile | NCTC10876_00166 | | Transcriptional regulator pfoR |
| LS483360 | 209033 | snp | T | G | CDS | + | missense_variant | c.24T>G | p.Phe8Leu | NCTC10876_00235 | radA | DNA repair protein RadA |
| LS483360 | 209059 | snp | C | A | CDS | + | stop_gained | c.50C>A | p.Ser17* | NCTC10876_00235 | radA | DNA repair protein RadA |
| LS483360 | 260275 | ins | A | AT | Intergenic | | | | | | | |
| LS483360 | 278065 | snp | A | C | CDS | + | missense_variant | c.208A>C | p.Ile70Leu | NCTC10876_00299 | tig | trigger factor |
| LS483360 | 278079 | snp | A | C | CDS | + | missense_variant | c.222A>C | p.Glu74Asp | NCTC10876_00299 | tig | trigger factor |
| LS483360 | 278318 | snp | T | G | CDS | + | missense_variant | c.461T>G | p.Ile154Ser | NCTC10876_00299 | tig | trigger factor |
| LS483360 | 309642 | snp | C | A | CDS | - | missense_variant | c.15G>T | p.Gln5His | NCTC10876_00332 | | hypothetical cytosolic protein |
| LS483360 | 311526 | snp | A | C | CDS | - | missense_variant | c.1451T>G | p.Leu484Arg | NCTC10876_00335 | pepX | xaa-Pro dipeptidyl-peptidase |
| LS483360 | 335469 | snp | T | C | CDS | + | synonymous_variant | c.75T>C | p.Ala25Ala | NCTC10876_00357 | ssb_2 | single strand binding protein |
| LS483360 | 352450 | snp | C | A | Intergenic | | | | | | | |
| LS483360 | 380388 | snp | A | C | CDS | + | missense_variant | c.204A>C | p.Gln68His | NCTC10876_00392 | yheD | cof-like hydrolase family protein |
| LS483360 | 381408 | snp | A | C | CDS | + | missense_variant | c.1224A>C | p.Glu408Asp | NCTC10876_00392 | yheD | cof-like hydrolase family protein |
| LS483360 | 383894 | snp | T | G | CDS | + | stop_gained | c.84T>G | p.Tyr28* | NCTC10876_00395 | codY | transcriptional repressor |
| LS483360 | 412882 | snp | G | C | CDS | - | missense_variant | c.906C>G | p.Asp302Glu | NCTC10876_00428 | manL | PTS system mannose-specific transporter subunit IIAB |
| LS483360 | 412894 | snp | T | G | CDS | - | missense_variant | c.894A>C | p.Glu298Asp | NCTC10876_00428 | manL | PTS system mannose-specific transporter subunit IIAB |
| LS483360 | 412925 | del | GATAAAACGTTGTTAACCATTGTTTTAC | G | CDS | - | disruptive_inframe_deletion | c.836_862delGTAAAACAATGGTTAACAACGTTTTAT | p.Gly279_Ser288delinsAla | NCTC10876_00428 | manL | PTS system mannose-specific transporter subunit IIAB |
| LS483360 | 413008 | snp | A | C | CDS | - | synonymous_variant | c.780T>G | p.Val260Val | NCTC10876_00428 | manL | PTS system mannose-specific transporter subunit IIAB |
| LS483360 | 425415 | snp | A | C | CDS | + | stop_lost&splice_region_variant | c.1146A>C | p.Ter382Tyrext*? | NCTC10876_00442 | nusA | transcription elongation factor |
| LS483360 | 425667 | snp | C | T | CDS | + | synonymous_variant | c.237C>T | p.Tyr79Tyr | NCTC10876_00443 | | putative cytoplasmic protein |
| LS483360 | 426783 | del | GCAAGA | G | CDS | + | frameshift_variant | c.748_752delCAAGA | p.Gln250fs | NCTC10876_00445 | infB | translation initiation factor IF-2 |
| LS483360 | 453667 | snp | A | C | CDS | + | missense_variant | c.1094A>C | p.Tyr365Ser | NCTC10876_00474 | | Sodium-dependent phosphate transporter |
| LS483360 | 491565 | snp | T | G | CDS | + | synonymous_variant | c.666T>G | p.Pro222Pro | NCTC10876_00507 | pepC | aminopeptidase C |
| LS483360 | 503847 | snp | C | A | CDS | - | missense_variant | c.1171G>T | p.Val391Leu | NCTC10876_00520 | atoB | acetyl-CoA acetyltransferase |
| LS483360 | 503862 | snp | C | A | CDS | - | stop_gained | c.1156G>T | p.Gly386* | NCTC10876_00520 | atoB | acetyl-CoA acetyltransferase |
| LS483360 | 503876 | snp | A | C | CDS | - | missense_variant | c.1142T>G | p.Ile381Ser | NCTC10876_00520 | atoB | acetyl-CoA acetyltransferase |
| LS483360 | 522886 | snp | A | C | CDS | + | missense_variant | c.488A>C | p.Gln163Pro | NCTC10876_00538 | | helicase conserved C-terminal domain protein |
| LS483360 | 589308 | complex | TT | GG | CDS | + | missense_variant | c.382_383delTTinsGG | p.Leu128Gly | NCTC10876_00601 | | arginine/ornithine antiporter |
| LS483360 | 594572 | snp | A | T | CDS | + | missense_variant | c.407A>T | p.Asn136Ile | NCTC10876_00605 | rsmD | methyltransferase |
| LS483360 | 596827 | snp | T | G | CDS | + | missense_variant | c.213T>G | p.Asn71Lys | NCTC10876_00608 | rbsR | LacI family transcriptional regulator |
| LS483360 | 596835 | snp | T | G | CDS | + | missense_variant | c.221T>G | p.Phe74Cys | NCTC10876_00608 | rbsR | LacI family transcriptional regulator |
| LS483360 | 596855 | snp | G | T | CDS | + | missense_variant | c.241G>T | p.Ala81Ser | NCTC10876_00608 | rbsR | LacI family transcriptional regulator |
| LS483360 | 612145 | snp | G | T | CDS | + | missense_variant | c.1279G>T | p.Asp427Tyr | NCTC10876_00623 | ftsZ | cell division protein FtsZ |
| LS483360 | 618463 | snp | A | C | Intergenic | | | | | | | |
| LS483360 | 632871 | snp | A | T | CDS | + | missense_variant | c.545A>T | p.Glu182Val | NCTC10876_00644 | recN | DNA repair protein recN |
| LS483360 | 648033 | snp | A | C | CDS | + | missense_variant | c.719A>C | p.Gln240Pro | NCTC10876_00658 | murF | UDP-N-acetylmuramoyl-tripeptide--D-alanyl-D- alanine ligase |
| LS483360 | 656645 | snp | T | G | CDS | + | missense_variant | c.134T>G | p.Phe45Cys | NCTC10876_00665 | plsC | 1-acyl-sn-glycerol-3-phosphate acyltransferase |
| LS483360 | 659854 | snp | G | T | CDS | + | missense_variant | c.1750G>T | p.Gly584Cys | NCTC10876_00667 | comEC | DNA internalization-related competence protein ComEC/Rec2 |
| LS483360 | 664616 | del | TTTTCAAAGGCAATCCGCTGATAGTCTTTC | T | CDS | - | frameshift_variant | c.33_61delGAAAGACTATCAGCGGATTGCCTTTGAAA | p.Lys12fs | NCTC10876_00672 | | hypothetical cytosolic protein |
| LS483360 | 690332 | snp | T | A | CDS | + | missense_variant | c.839T>A | p.Val280Glu | NCTC10876_00696 | deadD | polysaccharide deacetylase family protein |
| LS483360 | 690337 | snp | A | C | CDS | + | missense_variant | c.844A>C | p.Asn282His | NCTC10876_00696 | deadD | polysaccharide deacetylase family protein |
| LS483360 | 694166 | complex | CAAT | GCAG | CDS | + | missense_variant | c.129_132delCAATinsGCAG | p.Asn44Gln | NCTC10876_00701 | msrC | GAF domain-containing protein |
| LS483360 | 694190 | snp | C | G | CDS | + | stop_gained | c.153C>G | p.Tyr51* | NCTC10876_00701 | msrC | GAF domain-containing protein |
| LS483360 | 694209 | complex | TTA | GTC | CDS | + | missense_variant | c.172_174delTTAinsGTC | p.Leu58Val | NCTC10876_00701 | msrC | GAF domain-containing protein |
| LS483360 | 694222 | snp | C | A | CDS | + | missense_variant | c.185C>A | p.Pro62His | NCTC10876_00701 | msrC | GAF domain-containing protein |
| LS483360 | 694234 | snp | G | T | CDS | + | missense_variant | c.197G>T | p.Arg66Met | NCTC10876_00701 | msrC | GAF domain-containing protein |
| LS483360 | 694368 | snp | A | T | CDS | + | missense_variant | c.331A>T | p.Ile111Phe | NCTC10876_00701 | msrC | GAF domain-containing protein |
| LS483360 | 694381 | snp | T | G | CDS | + | missense_variant | c.344T>G | p.Met115Arg | NCTC10876_00701 | msrC | GAF domain-containing protein |
| LS483360 | 707059 | snp | G | T | CDS | + | missense_variant | c.553G>T | p.Ala185Ser | NCTC10876_00710 | map | methionine aminopeptidase Map |
| LS483360 | 761506 | snp | G | T | CDS | - | missense_variant | c.95C>A | p.Pro32Gln | NCTC10876_00758 | | phage protein |
| LS483360 | 765067 | snp | T | G | CDS | - | missense_variant | c.658A>C | p.Lys220Gln | NCTC10876_00765 | | Uncharacterised protein |
| LS483360 | 765127 | snp | C | A | CDS | - | missense_variant | c.598G>T | p.Asp200Tyr | NCTC10876_00765 | | Uncharacterised protein |
| LS483360 | 765158 | snp | A | T | CDS | - | synonymous_variant | c.567T>A | p.Pro189Pro | NCTC10876_00765 | | Uncharacterised protein |
| LS483360 | 783367 | complex | AGGGAAAG | CGGTAAAC | CDS | + | missense_variant | c.327_334delAGGGAAAGinsCGGTAAAC | p.GluGlyLysAsp109AspGlyLysHis | NCTC10876_00797 | | phage major capsid protein |
| LS483360 | 783387 | snp | T | G | CDS | + | missense_variant | c.347T>G | p.Ile116Ser | NCTC10876_00797 | | phage major capsid protein |
| LS483360 | 799782 | snp | T | A | CDS | - | missense_variant | c.453A>T | p.Lys151Asn | NCTC10876_00815 | speC_1 | exotoxin type C |
| LS483360 | 825197 | snp | A | C | CDS | + | missense_variant | c.470A>C | p.Asn157Thr | NCTC10876_00840 | | response regulator |
| LS483360 | 825217 | snp | A | C | CDS | + | missense_variant | c.490A>C | p.Asn164His | NCTC10876_00840 | | response regulator |
| LS483360 | 851134 | ins | C | CA | CDS | + | frameshift_variant | c.486dupA | p.Gln163fs | NCTC10876_00867 | rsmC | 16S rRNA methyltransferase |
| LS483360 | 852521 | snp | G | A | CDS | + | missense_variant | c.136G>A | p.Gly46Ser | NCTC10876_00869 | tmpC | nucleoside-binding protein |
| LS483360 | 853529 | snp | T | A | Intergenic | | | | | | | |
| LS483360 | 909790 | snp | G | T | CDS | - | missense_variant | c.644C>A | p.Ser215Tyr | NCTC10876_00925 | topA | DNA topoisomerase I |
| LS483360 | 912031 | complex | TT | GA | CDS | - | missense_variant | c.202_203delAAinsTC | p.Lys68Ser | NCTC10876_00927 | rnhB | ribonuclease HII |
| LS483360 | 923872 | ins | C | CT | CDS | - | frameshift_variant | c.1052dupA | p.Thr352fs | NCTC10876_00940 | msbA | ABC transporter |
| LS483360 | 930148 | snp | T | C | CDS | - | missense_variant | c.398A>G | p.Lys133Arg | NCTC10876_00946 | rimN | Sua5/YciO/YrdC/YwlC family protein |
| LS483360 | 930282 | snp | A | C | CDS | - | synonymous_variant | c.264T>G | p.Ala88Ala | NCTC10876_00946 | rimN | Sua5/YciO/YrdC/YwlC family protein |
| LS483360 | 930317 | snp | A | T | CDS | - | missense_variant | c.229T>A | p.Phe77Ile | NCTC10876_00946 | rimN | Sua5/YciO/YrdC/YwlC family protein |
| LS483360 | 930326 | snp | A | C | CDS | - | missense_variant | c.220T>G | p.Tyr74Asp | NCTC10876_00946 | rimN | Sua5/YciO/YrdC/YwlC family protein |
| LS483360 | 936059 | snp | A | C | CDS | - | synonymous_variant | c.288T>G | p.Val96Val | NCTC10876_00953 | xpt | xanthine phosphoribosyltransferase |
| LS483360 | 937718 | snp | C | A | Intergenic | | | | | | | |
| LS483360 | 937907 | snp | T | A | Intergenic | | | | | | | |
| LS483360 | 937923 | snp | T | G | CDS | - | missense_variant | c.1516A>C | p.Ile506Leu | NCTC10876_00955 | opuCB | putative ABC transporter |
| LS483360 | 961165 | snp | T | G | CDS | + | missense_variant | c.1277T>G | p.Ile426Ser | NCTC10876_00977 | dcuS | Sensor kinase dpiB |
| LS483360 | 977466 | snp | T | G | CDS | - | missense_variant | c.182A>C | p.Glu61Ala | NCTC10876_00999 | srtF | lantibiotic ABC transporter ATP-binding protein |
| LS483360 | 988940 | snp | T | A | Intergenic | | | | | | | |

| | | | | | | | | | | | | |
|---|---|---|---|---|---|---|---|---|---|---|---|---|
| LS483360 | 988946 | complex | GAAT | CAAA | Intergenic | | | | | | | |
| LS483360 | 999963 | snp | T | G | CDS | - | missense_variant | c.1073A>C | p.Lys358Thr | NCTC10876_01023 | yehU | two-component sensor kinase |
| LS483360 | 1025744 | del | TTTCA | T | CDS | - | frameshift_variant | c.51_54delTGAA | p.Glu18fs | NCTC10876_01044 | | dihydrolipoamide dehydrogenase |
| LS483360 | 1033389 | snp | G | C | CDS | - | synonymous_variant | c.1530C>G | p.Gly510Gly | NCTC10876_01051 | | Zn-dependent hydrolase |
| LS483360 | 1059087 | del | TTCTGACATATTAA | T | CDS | - | bidirectional_gene_fusion | n.1059100_1059088delTTAATATGTCAGA | | NCTC10876_01082 | | phage protein |
| LS483360 | 1073224 | snp | C | A | CDS | - | missense_variant | c.307G>T | p.Ala103Ser | NCTC10876_01109 | | phage protein |
| LS483360 | 1073289 | snp | T | G | CDS | - | missense_variant | c.242A>C | p.Tyr81Ser | NCTC10876_01109 | | phage protein |
| LS483360 | 1073297 | snp | A | C | CDS | - | synonymous_variant | c.234T>G | p.Arg78Arg | NCTC10876_01109 | | phage protein |
| LS483360 | 1081070 | snp | T | G | CDS | - | missense_variant | c.68A>C | p.Lys23Thr | NCTC10876_01121 | rmlA1 | glucose-1-phosphate thymidyl transferase |
| LS483360 | 1086941 | snp | T | C | CDS | - | missense_variant | c.1232A>G | p.His411Arg | NCTC10876_01128 | recJ | single-stranded-DNA-specific exonuclease RecJ |
| LS483360 | 1111135 | snp | G | T | CDS | - | synonymous_variant | c.579C>A | p.Val193Val | NCTC10876_01153 | amiC | amidase |
| LS483360 | 1140514 | snp | T | C | Intergenic | | | | | | | |
| LS483360 | 1150889 | snp | G | T | Intergenic | | | | | | | |
| LS483360 | 1151554 | del | GAGGA | G | CDS | - | frameshift_variant | c.1325_1328delTCCT | p.Ile442fs | NCTC10876_01194 | fruA | PTS system fructose-specific transporter subunit IIABC |
| LS483360 | 1155810 | snp | A | C | CDS | - | missense_variant | c.941T>G | p.Ile314Arg | NCTC10876_01198 | | regulatory protein |
| LS483360 | 1181380 | snp | C | A | CDS | - | missense_variant | c.130G>T | p.Gly44Trp | NCTC10876_01220 | oxyR | LysR family transcriptional regulator |
| **LS483360** | **1204515** | **snp** | **G** | **A** | **CDS** | **-** | **stop_gained** | **c.382C>T** | **p.Arg128*** | **NCTC10876_01248** | **gacI** | **glycosyl transferase 2 family protein** |
| LS483360 | 1217554 | snp | A | C | CDS | - | missense_variant | c.1471T>G | p.Leu491Val | NCTC10876_01259 | dnaG_2 | DNA primase |
| LS483360 | 1247252 | snp | A | C | CDS | - | missense_variant | c.80T>G | p.Val27Gly | NCTC10876_01286 | | bacteriophage Gp15 family protein |
| LS483360 | 1282848 | snp | G | T | CDS | - | stop_gained | c.1383C>A | p.Tyr461* | NCTC10876_01345 | atpA | F0F1 ATP synthase subunit alpha |
| LS483360 | 1282869 | snp | C | A | CDS | - | missense_variant | c.1362G>T | p.Leu454Phe | NCTC10876_01345 | atpA | F0F1 ATP synthase subunit alpha |
| LS483360 | 1282881 | snp | T | A | CDS | - | synonymous_variant | c.1350A>T | p.Val450Val | NCTC10876_01345 | atpA | F0F1 ATP synthase subunit alpha |
| LS483360 | 1325779 | snp | T | C | CDS | - | missense_variant | c.611A>G | p.Glu204Gly | NCTC10876_01378 | agaS | tagatose-6-phosphate ketose/aldose isomerase |
| LS483360 | 1327354 | snp | C | G | CDS | - | missense_variant | c.1471G>C | p.Glu491Gln | NCTC10876_01380 | znuA | high-affinity zinc uptake system protein |
| LS483360 | 1327374 | snp | C | G | CDS | - | missense_variant | c.1451G>C | p.Ser484Thr | NCTC10876_01380 | znuA | high-affinity zinc uptake system protein |
| LS483360 | 1368803 | snp | T | G | CDS | - | missense_variant | c.583A>C | p.Ile195Leu | NCTC10876_01415 | ftsW | cell division protein |
| LS483360 | 1369087 | snp | T | C | CDS | - | missense_variant | c.299A>G | p.Lys100Arg | NCTC10876_01415 | ftsW | cell division protein |
| LS483360 | 1385438 | snp | T | G | CDS | + | synonymous_variant | c.210T>G | p.Ala70Ala | NCTC10876_01431 | | membrane protein |
| LS483360 | 1409807 | snp | A | T | CDS | - | missense_variant | c.2046T>A | p.Ser682Arg | NCTC10876_01454 | mcdM | putative lantibiotic modifying protein |
| LS483360 | 1421524 | snp | A | T | CDS | - | missense_variant | c.7T>A | p.Ser3Thr | NCTC10876_01468 | | huntington interacting protein HYPE |
| LS483360 | 1452470 | ins | A | AT | CDS | - | frameshift_variant | c.41dupA | p.Asn14fs | NCTC10876_01499 | drrA_3 | antibiotic transport system ATP-binding protein |
| LS483360 | 1458388 | snp | T | G | CDS | - | missense_variant | c.89A>C | p.Asn30Thr | NCTC10876_01504 | ccpA | Catabolite control protein A |
| LS483360 | 1491975 | snp | T | G | CDS | - | missense_variant | c.145A>C | p.Thr49Pro | NCTC10876_01545 | frr | ribosome recycling factor |
| LS483360 | 1491986 | snp | T | G | CDS | - | missense_variant | c.134A>C | p.Tyr45Ser | NCTC10876_01545 | frr | ribosome recycling factor |
| LS483360 | 1495349 | del | ATCCGCTTCAGCT | A | CDS | - | disruptive_inframe_deletion | c.2130_2141delAGCTGAAGCGGA | p.Gln710_Asp714delinsHis | NCTC10876_01550 | ftsK_1 | cell division protein FtsK |
| LS483360 | 1512152 | snp | A | C | CDS | - | missense_variant | c.74T>G | p.Met25Arg | NCTC10876_01568 | | hypothetical membrane associated protein |
| LS483360 | 1514458 | snp | T | G | Intergenic | | | | | | | |
| LS483360 | 1530113 | snp | A | C | CDS | - | missense_variant | c.1022T>G | p.Ile341Ser | NCTC10876_01579 | | L-lactate oxidase |
| LS483360 | 1550073 | snp | G | T | CDS | - | missense_variant | c.1193C>A | p.Pro398Gln | NCTC10876_01601 | hlyX | hemolysin |
| LS483360 | 1561392 | snp | A | C | Intergenic | | | | | | | |
| LS483360 | 1579923 | snp | G | A | CDS | - | missense_variant | c.47C>T | p.Pro16Leu | NCTC10876_01634 | csrS | HAMP domain-containing protein |
| LS483360 | 1611322 | snp | T | G | CDS | + | missense_variant | c.2699T>G | p.Leu900Trp | NCTC10876_01665 | hsdR | type I site-specific deoxyribonuclease, HsdR family protein |
| LS483360 | 1612597 | del | ATTTACAAGGTGGCCAAGGAAATTTATCTGGT | A | CDS | + | frameshift_variant | c.985_1015delTTACAAGGTGGCCAAGGAAATTTATCTGGTT | p.Gln330fs | NCTC10876_01666 | hsdS | Type I restriction-modification system specificity subunit |
| LS483360 | 1614774 | del | AAGCT | A | CDS | - | frameshift_variant | c.7_10delAGCT | p.Ser3fs | NCTC10876_01668 | | Uncharacterised protein |
| LS483360 | 1621385 | snp | A | G | CDS | - | synonymous_variant | c.543T>C | p.Ser181Ser | NCTC10876_01674 | | salivaricin A modification enzyme |
| LS483360 | 1630018 | snp | A | C | CDS | + | missense_variant | c.432A>C | p.Leu144Phe | NCTC10876_01683 | lacR_2 | Lactose phosphotransferase system repressor |
| LS483360 | 1631598 | snp | C | A | CDS | + | missense_variant | c.301C>A | p.Pro101Thr | NCTC10876_01685 | | Toxin to DNA-damage-inducible protein J |
| LS483360 | 1639544 | snp | A | C | CDS | + | missense_variant | c.110A>C | p.Lys37Thr | NCTC10876_01694 | | integrase |
| LS483360 | 1647726 | snp | A | G | CDS | - | missense_variant | c.166T>C | p.Tyr56His | NCTC10876_01705 | cysS | cysteinyl-tRNA synthetase |
| LS483360 | 1654902 | snp | T | G | CDS | - | missense_variant | c.1958A>C | p.Asp653Ala | NCTC10876_01713 | ulaC_2 | PTS system mannitol (cryptic)-specific transporter subunit IIA |
| LS483360 | 1654971 | complex | ATCAA | CTCAC | CDS | - | missense_variant | c.1885_1889delTTGATinsGTGAG | p.LeuIle629ValSer | NCTC10876_01713 | ulaC_2 | PTS system mannitol (cryptic)-specific transporter subunit IIA |
| LS483360 | 1679350 | snp | T | A | CDS | + | missense_variant | c.287T>A | p.Met96Lys | NCTC10876_01730 | skc | streptokinase |
| LS483360 | 1714385 | snp | T | G | CDS | - | missense_variant | c.556A>C | p.Ile186Leu | NCTC10876_01759 | emm49 | Antiphagocytic M protein |
| LS483360 | 1814605 | snp | G | T | CDS | - | missense_variant | c.38C>A | p.Pro13His | NCTC10876_01863 | | tetratricopeptide repeat family protein |
| LS483360 | 1814621 | snp | G | T | CDS | - | missense_variant | c.22C>A | p.Gln8Lys | NCTC10876_01863 | | tetratricopeptide repeat family protein |
| LS483360 | 1814646 | snp | A | T | Intergenic | | | | | | | |
| LS483360 | 1854714 | snp | C | A | CDS | + | missense_variant | c.31C>A | p.Leu11Met | NCTC10876_01902 | cadD | cadmium resistance protein |
| LS483360 | 1897438 | snp | A | C | CDS | + | missense_variant | c.465A>C | p.Glu155Asp | NCTC10876_01943 | yheS_2 | ABC transporter ATP-binding protein |

**Supplementary Table S4: Metadata of 2,021 S. pyogenes isolates screened in this study**

| Isolate | Country | Continent | Year | Primary_disease^ | *Emm* | ST (MLST) | *gacA* # | *gacB* | *gacC* | *gacD* | *gacE* | *gacF* | *gacG* | *gacH* | *gacI* | *gacJ* | *gacK* | *gacL* |
|---|---|---|---|---|---|---|---|---|---|---|---|---|---|---|---|---|---|---|
| NS2403 | Australia | Oceania | 2000 | APSGN | emm3.22 | 335 | 9 | 9 | 8 | 8 | 8 | 10 | 9 | 10 | 5 | 5 | 9 | 9 |
| NS2413 | Australia | Oceania | 2000 | skin/soft tissue infection, NOS | emm85.0 | 109 | 10 | 10 | 4 | 9 | 9 | 11 | 10 | 11 | 7 | 3 | 10 | 10 |
| NS2985 | Australia | Oceania | 2001 | APSGN | emm105.0 | 954 | 6 | 6 | 6 | 6 | 5 | 12 | 6 | 12 | 5 | 4 | 6 | 11 |
| NS4729 | Australia | Oceania | 2005 | APSGN | emm55.0 | 100 | 11 | 11 | 7 | 10 | 10 | 13 | 11 | 13 | 8 | 3 | 11 | 12 |
| NS4972 | Australia | Oceania | 2005 | APSGN | emm55.0 | 100 | 11 | 11 | 7 | 10 | 10 | 13 | 11 | 13 | 8 | 3 | 11 | 12 |
| NS81 | Australia | Oceania | 1992 | APSGN | emm55.0 | 100 | 11 | 11 | 7 | 10 | 10 | 13 | 11 | 13 | 8 | 3 | 11 | 12 |
| NS2188 | Australia | Oceania | 2000 | APSGN | emm49.3 | 190 | 4 | 8 | 4 | 11 | 11 | 14 | 12 | 14 | 9 | 2 | 12 | 13 |
| GAS253 | Australia | Oceania | ND | ND | emm19.7 | 370 | 12 | 12 | 4 | 12 | 12 | 13 | 13 | 15 | 8 | 3 | 13 | 14 |
| NS5268 | Australia | Oceania | 2005 | APSGN | emm55.0 | 100 | 11 | 11 | 7 | 10 | 10 | 13 | 11 | 13 | 8 | 3 | 11 | 12 |
| NS426 | Australia | Oceania | 1995 | APSGN | emm239.0 | 92 | 13 | 1 | 1 | 1 | 1 | 15 | 14 | 16 | 5 | 6 | 14 | 15 |
| NS100 | Australia | Oceania | 1992 | APSGN | emm68.2 | 298 | 14 | 13 | 9 | 8 | 13 | 13 | 15 | 17 | 10 | 2 | 15 | 16 |
| NS6902 | Australia | Oceania | ND | APSGN | emm49.0 | 371 | 4 | 8 | 4 | 13 | 14 | 14 | 16 | 18 | 9 | 2 | 12 | 13 |
| NS3965 | Australia | Oceania | ND | skin/soft tissue infection, NOS | emm49.3 | 190 | 4 | 8 | 4 | 14 | 11 | 14 | 12 | 19 | 9 | 2 | 12 | 17 |
| Dog48 | Australia | Oceania | 1990 | ND | emm91.0 | 12 | 11 | 5 | 10 | 15 | 15 | 16 | 17 | 13 | 1 | 2 | 16 | 18 |
| NS27 | Australia | Oceania | 1991 | APSGN | emm91.0 | 12 | 11 | 5 | 10 | 15 | 15 | 16 | 17 | 13 | 1 | 2 | 16 | 18 |
| NS38 | Australia | Oceania | 1991 | APSGN | emm57.2 | 33 | 11 | 14 | 11 | 8 | 16 | 17 | 18 | 20 | 11 | 7 | 17 | 19 |
| NS5658 | Australia | Oceania | 2005 | APSGN | emm55.0 | 100 | 11 | 11 | 7 | 10 | 10 | 13 | 11 | 13 | 8 | 3 | 11 | 12 |
| NS76 | Australia | Oceania | 1992 | APSGN | emm98.1 | 205 | 15 | 15 | 10 | 1 | 10 | 18 | 19 | 21 | 12 | 2 | 6 | 20 |
| NS48 | Australia | Oceania | 1991 | APSGN | emm57.2 | 33 | 11 | 14 | 11 | 8 | 16 | 17 | 18 | 20 | 11 | 7 | 17 | 19 |
| SDog | Australia | Oceania | 1990 | ND | emm70.0 | 10 | 16 | 16 | 3 | 16 | 17 | 7 | 20 | 22 | 1 | 2 | 18 | 3 |
| NS22 | Australia | Oceania | 1991 | APSGN | emm57.2 | 33 | 11 | 14 | 11 | 8 | 16 | 17 | 18 | 20 | 11 | 7 | 17 | 19 |
| YARB5 | Australia | Oceania | 2010 | skin/soft tissue infection, NOS | emm52.1 | 180 | 15 | 17 | 10 | 17 | 18 | 18 | 21 | 23 | 12 | 8 | 19 | 21 |
| NS2283 | Australia | Oceania | 2000 | APSGN | emm3.22 | 335 | 9 | 9 | 8 | 8 | 8 | 10 | 9 | 10 | 5 | 5 | 9 | 9 |
| NS2299 | Australia | Oceania | 2000 | APSGN | emm3.22 | 335 | 9 | 9 | 8 | 8 | 8 | 10 | 9 | 10 | 5 | 5 | 9 | 9 |
| NS2361 | Australia | Oceania | 2000 | APSGN | emm70.0 | 10 | 16 | 16 | 3 | 16 | 17 | 7 | 20 | 22 | 1 | 2 | 18 | 3 |
| YARC4 | Australia | Oceania | 2010 | skin/soft tissue infection, NOS | emm92.0 | 82 | 17 | 4 | 3 | 18 | 19 | 19 | 22 | 24 | 13 | 2 | 20 | 22 |
| YARG2 | Australia | Oceania | 2010 | skin/soft tissue infection, NOS | emm11.0 | ND | 1 | 6 | 12 | 1 | 20 | 20 | 13 | 25 | 8 | 3 | 21 | 23 |
| YARG4 | Australia | Oceania | 2010 | skin/soft tissue infection, NOS | emm124.0 | 544 | 18 | 18 | 4 | 16 | 21 | 21 | 23 | 26 | 8 | 9 | 21 | 23 |
| NS53 | Australia | Oceania | 1991 | skin/soft tissue infection, NOS | emm71.0 | 318 | 19 | [S] | 13 | 19 | 2 | 22 | 3 | 27 | 1 | 1 | 4 | 24 |
| NS564 | Australia | Oceania | 1995 | invasive, NOS | emm44.0 | 178 | 5 | 7 | 7 | 3 | 2 | 3 | 24 | 8 | 14 | 3 | 7 | 7 |
| NS696 | Australia | Oceania | 1996 | pharyngitis and/or tonsillitis | emm1.0 | 1014 | 1 | 1 | 1 | 1 | 1 | 1 | 1 | 1 | 1 | 1 | 1 | 1 |
| NS730 | Australia | Oceania | 1996 | skin/soft tissue infection, NOS | emm90.5 | 1083 | 20 | 20 | 14 | 19 | 2 | 23 | 25 | 28 | 15 | 3 | 22 | 25 |
| NS80 | Australia | Oceania | 1992 | skin/soft tissue infection, NOS | emm70.0 | 1063 | 16 | 16 | 3 | [S] | 17 | 7 | 20 | [S] | 1 | 2 | 18 | 3 |
| NS803 | Australia | Oceania | 1996 | invasive, NOS | emm97.1 | 216 | 21 | 21 | 15 | 8 | 22 | 24 | 26 | 30 | 8 | 10 | 12 | 26 |
| NS83 | Australia | Oceania | 1992 | invasive, NOS | emm232.1 | 288 | 18 | 18 | 16 | 16 | 21 | 25 | 27 | 26 | 16 | 3 | 21 | 23 |
| NS931 | Australia | Oceania | 1996 | skin/soft tissue infection, NOS | emm65.0 | 1015 | 22 | 22 | 17 | 21 | 23 | 26 | 28 | 31 | 1 | 1 | 1 | 27 |
| NS3 | Australia | Oceania | 1990 | invasive, NOS | emm98.0 | 1076 | 15 | 15 | 10 | 1 | [S] | 18 | 19 | 21 | 12 | 2 | 6 | 20 |
| NS101 | Australia | Oceania | 1992 | skin/soft tissue infection, NOS | emm110.0 | 628 | 14 | 23 | 18 | 6 | 25 | 27 | 29 | 32 | 1 | 2 | 23 | 28 |
| NS236 | Australia | Oceania | 1994 | skin/soft tissue infection, NOS | emm77.0 | 63 | 23 | 24 | 19 | 1 | 26 | 28 | 30 | 33 | 8 | 3 | 24 | 29 |
| NS240 | Australia | Oceania | 1995 | skin/soft tissue infection, NOS | ND | 166 | 14 | 25 | 3 | 22 | 27 | 13 | 26 | 29 | 8 | 3 | 25 | 30 |
| NS50.1 | Australia | Oceania | 1991 | skin/soft tissue infection, NOS | emm108.0 | 304 | 11 | 5 | 10 | 20 | 15 | 16 | 31 | 13 | 1 | 2 | 16 | 31 |
| NS1033 | Australia | Oceania | 1997 | ND | emm230.0 | 205 | 1 | 15 | 10 | 1 | 24 | 18 | 19 | 21 | 12 | 2 | 6 | 20 |
| NS1122 | Australia | Oceania | 1997 | pharyngitis and/or tonsillitis | emm63.3 | 297 | 24 | 19 | 3 | 23 | 28 | 7 | 32 | 34 | 17 | 11 | 26 | 32 |
| NS1210 | Australia | Oceania | 1997 | invasive, NOS | emm75.0 | 632 | 25 | 26 | 20 | 24 | 29 | 29 | 33 | 35 | 15 | 3 | 12 | 33 |
| NS13 | Australia | Oceania | 1991 | invasive, NOS | emm53.0 | 299 | 11 | 5 | 10 | 20 | 15 | 16 | 17 | 13 | 1 | 2 | 18 | 3 |

| | | | | | | | | | | | | | | | | | | |
|---|---|---|---|---|---|---|---|---|---|---|---|---|---|---|---|---|---|---|
| NS192 | Australia | Oceania | 1994 | skin/soft tissue infection, NOS | emm106.0 | 338 | 26 | 27 | 21 | 25 | 30 | 19 | 34 | 36 | 15 | 3 | 27 | 34 |
| NS195 | Australia | Oceania | 1994 | skin/soft tissue infection, NOS | ND | 370 | 12 | 12 | 4 | 12 | 12 | 13 | 13 | 15 | 8 | 3 | 13 | 14 |
| NS20 | Australia | Oceania | 1991 | skin/soft tissue infection, NOS | emm75.1 | 607 | 27 | 28 | 22 | 26 | 31 | 13 | 35 | 37 | 18 | 3 | 28 | 30 |
| NS216 | Australia | Oceania | 1994 | invasive, NOS | emm229.0 | 614 | 28 | 29 | 23 | 1 | 32 | 30 | 36 | 38 | 8 | 5 | 29 | 35 |
| NS1 | Australia | Oceania | 1990 | skin/soft tissue infection, NOS | emm100.0 | 119 | 11 | 11 | 7 | 10 | 10 | 7 | 37 | 39 | 19 | 2 | 30 | 31 |
| NS1132 | Australia | Oceania | 1997 | skin/soft tissue infection, NOS | emm25.1 | 192 | 14 | 30 | 24 | 2 | 33 | 13 | 13 | 40 | 20 | 3 | 31 | 36 |
| NS1133 | Australia | Oceania | 1997 | invasive, NOS | emm101.0 | 182 | 15 | 31 | 10 | 20 | 15 | 31 | 38 | 41 | 1 | 3 | 32 | 37 |
| NS1140 | Australia | Oceania | 1997 | skin/soft tissue infection, NOS | emm57.0 | 348 | 29 | 32 | 7 | 1 | 34 | 32 | 39 | 42 | 1 | 2 | 23 | 3 |
| NS10 | Australia | Oceania | 1990 | APSGN | emm53.0 | 299 | 11 | 5 | 10 | 20 | 15 | 16 | 17 | 13 | 1 | 2 | 18 | 3 |
| NS1036 | Australia | Oceania | 1997 | ND | emm110.0 | 179 | 14 | 33 | 18 | 6 | 25 | 27 | 40 | 43 | 1 | 2 | 18 | 38 |
| NS1045 | Australia | Oceania | 1997 | ND | emm60.7 | 193 | 14 | 34 | 25 | 27 | 2 | 33 | 41 | 44 | 1 | 2 | 33 | 39 |
| NS1053 | Australia | Oceania | 1997 | invasive, NOS | emm60.4 | 193 | 14 | 34 | 25 | 27 | 2 | 33 | 41 | 44 | 1 | 2 | 33 | 39 |
| NS1096 | Australia | Oceania | 1997 | skin/soft tissue infection, NOS | emm25.0 | 191 | 30 | 35 | 26 | 28 | 35 | 34 | 42 | 45 | 11 | 2 | 12 | 40 |
| NS1107 | Australia | Oceania | 1997 | skin/soft tissue infection, NOS | emm76.4 | 631 | 31 | 36 | 27 | 1 | 36 | 35 | 4 | 46 | 21 | 1 | 4 | 41 |
| NS1017 | Australia | Oceania | 1997 | skin/soft tissue infection, NOS | emm90.5 | 184 | 20 | 20 | 14 | 19 | 2 | 23 | 25 | 28 | 15 | 3 | 22 | 25 |
| NS1120 | Australia | Oceania | 1997 | skin/soft tissue infection, NOS | emm77.0 | 1016 | 23 | 24 | 19 | 1 | 26 | 28 | 30 | 33 | 8 | 3 | 24 | 29 |
| NS196 | Australia | Oceania | 1994 | ARF | emm58.0 | 176 | 14 | 13 | 10 | 29 | 13 | 24 | 15 | 47 | 10 | 2 | 34 | 31 |
| NS204 | Australia | Oceania | 1994 | invasive, NOS | emm2.0 | 55 | 32 | 26 | 28 | 24 | 29 | 13 | 43 | 48 | 15 | 3 | 12 | 42 |
| NS210 | Australia | Oceania | 1994 | invasive, NOS | emm22.0 | 202 | 33 | 11 | 7 | 10 | 35 | 36 | 44 | 49 | 19 | 2 | 30 | 43 |
| NS1216 | Australia | Oceania | 1997 | invasive, NOS | emm109.1 | 633 | 8 | 4 | 29 | 30 | 19 | 37 | 45 | 50 | 15 | 3 | 35 | 34 |
| NS125 | Australia | Oceania | 1993 | invasive, NOS | emm95.0 | 604 | 11 | 32 | 30 | 31 | 37 | 7 | 46 | 51 | 1 | 2 | 6 | 44 |
| NS14 | Australia | Oceania | 1991 | skin/soft tissue infection, NOS | emm102.2 | 185 | 14 | 37 | 31 | 3 | 2 | 38 | 47 | 52 | 1 | 2 | 18 | 25 |
| NS176 | Australia | Oceania | 1994 | ND | emm207.1 | 332 | 11 | 38 | 32 | 16 | 38 | 18 | 48 | 53 | 1 | 2 | 6 | 20 |
| NS178 | Australia | Oceania | 1994 | skin/soft tissue infection, NOS | emm54.1 | 302 | 11 | 5 | 10 | 20 | 15 | 16 | 17 | 13 | 1 | 2 | 16 | 31 |
| NS179 | Australia | Oceania | 1994 | skin/soft tissue infection, NOS | emm9.0 | 204 | 34 | 39 | 33 | 32 | 2 | 1 | 49 | 54 | 5 | 3 | 11 | 45 |
| NS190 | Australia | Oceania | 1994 | invasive, NOS | emm92.0 | 312 | 35 | 40 | 34 | 33 | 39 | 39 | 22 | 24 | 13 | 2 | 20 | 22 |
| NS223 | Australia | Oceania | 1994 | skin/soft tissue infection, NOS | emm91.0 | 12 | 11 | 5 | 10 | 15 | 15 | 16 | 17 | 13 | 1 | 2 | 16 | 18 |
| NS282 | Australia | Oceania | 1995 | skin/soft tissue infection, NOS | emm207.1 | 332 | 11 | 38 | 32 | 16 | 38 | 18 | 48 | 55 | 1 | 2 | 6 | 20 |
| NS297 | Australia | Oceania | 1995 | ND | emm11.0 | ND | 1 | 6 | 12 | 1 | 20 | 20 | 13 | 25 | 8 | 3 | 21 | 23 |
| NS32 | Australia | Oceania | 1991 | invasive, NOS | emm101.0 | 182 | 15 | 31 | 10 | 20 | 40 | 31 | 38 | 41 | 1 | 3 | 32 | 37 |
| NS225 | Australia | Oceania | 1994 | invasive, NOS | emm99.0 | 141 | 14 | 41 | 35 | 1 | 20 | 40 | 26 | 56 | 5 | 4 | 20 | 46 |
| NS226 | Australia | Oceania | 1994 | invasive, NOS | emm4.5 | 1082 | 36 | 42 | 36 | 34 | 41 | 7 | 50 | 57 | 18 | 3 | 36 | 47 |
| NS235 | Australia | Oceania | 1994 | skin/soft tissue infection, NOS | emm24.1 | 582 | 14 | 43 | 33 | 1 | 42 | 13 | 32 | 58 | 22 | 8 | 37 | 48 |
| NS239 | Australia | Oceania | 1995 | invasive, NOS | emm207.1 | 610 | 6 | 6 | 6 | 6 | 5 | 6 | 6 | 7 | 5 | 4 | 6 | 6 |
| NS244 | Australia | Oceania | 1995 | ND | emm4.5 | 1017 | 36 | 42 | 36 | 34 | 41 | 7 | 50 | 57 | 18 | 3 | 36 | 47 |
| NS25 | Australia | Oceania | 1991 | invasive, NOS | emm55.0 | 1018 | 11 | 11 | 7 | 10 | 10 | 13 | 11 | 13 | 8 | 3 | 11 | 12 |
| NS253 | Australia | Oceania | 1995 | invasive, NOS | emm52.1 | 180 | 15 | 17 | 10 | 17 | 18 | 18 | 21 | 23 | 12 | 8 | 19 | 21 |
| NS265 | Australia | Oceania | 1995 | ND | emm56.0 | 115 | 14 | 44 | 24 | 2 | 12 | 41 | 51 | 22 | 23 | 1 | 6 | 49 |
| NS488 | Australia | Oceania | 1995 | invasive, NOS | emm12.0 | 1019 | 14 | 3 | 3 | 3 | 2 | 3 | 52 | 59 | 24 | 12 | 38 | [S] |
| NS578 | Australia | Oceania | 1995 | ND | emm232.0 | 110 | 18 | 18 | 16 | 16 | 21 | 42 | 27 | 26 | 25 | 3 | 21 | 51 |
| NS581 | Australia | Oceania | 1995 | invasive, NOS | emm42.1 | 80 | 37 | 32 | 37 | 35 | 43 | 43 | 44 | 60 | 26 | 2 | 39 | 52 |
| NS59 | Australia | Oceania | 1992 | ND | emm53.0 | 299 | 11 | 5 | 10 | 20 | 15 | 16 | 17 | 13 | 1 | 2 | 18 | 3 |
| NS495 | Australia | Oceania | 1995 | skin/soft tissue infection, NOS | emm90.5 | 184 | 20 | 20 | 14 | 19 | 2 | 23 | 25 | 28 | 15 | 3 | 22 | 25 |
| NS50 | Australia | Oceania | 1991 | invasive, NOS | emm218.1 | 292 | 38 | 45 | 3 | 16 | 44 | 44 | 53 | 61 | 27 | 2 | 12 | 3 |
| NS506 | Australia | Oceania | 1995 | invasive, NOS | emm14.4 | 118 | 11 | 5 | 38 | 8 | 15 | 16 | 17 | 13 | 1 | 2 | 40 | 53 |
| NS516 | Australia | Oceania | 1995 | ND | emm110.0 | 596 | 14 | 23 | 18 | 6 | 25 | 27 | 40 | 43 | 1 | 2 | 18 | 28 |
| NS534 | Australia | Oceania | 1995 | invasive, NOS | emm232.0 | 110 | 18 | 18 | 16 | 16 | 21 | 42 | 27 | 26 | | 3 | 21 | 51 |
| NS539 | Australia | Oceania | 1995 | skin/soft tissue infection, NOS | emm76.6 | 291 | 39 | 46 | 39 | 36 | 45 | 45 | 54 | 62 | 2 | 2 | 41 | 54 |

| | | | | | | | | | | | | | | | | | | |
|---|---|---|---|---|---|---|---|---|---|---|---|---|---|---|---|---|---|---|
| NS567 | Australia | Oceania | 1995 | skin/soft tissue infection, NOS | emm193.0 | 108 | 14 | 47 | 40 | 1 | 20 | 2 | 55 | 63 | 8 | 2 | 42 | 55 |
| NS611 | Australia | Oceania | 1995 | invasive, NOS | emm15.1 | 629 | 14 | 48 | 41 | 2 | 2 | 2 | 56 | 13 | 8 | 3 | 43 | 56 |
| NS733 | Australia | Oceania | 1996 | invasive, NOS | emm90.5 | 184 | 20 | 20 | 14 | 19 | 2 | 23 | 25 | 28 | 15 | 3 | 22 | 25 |
| NS736 | Australia | Oceania | 1996 | ND | emm221.0 | 113 | 11 | 49 | 2 | 37 | 2 | 32 | 57 | 42 | 1 | 2 | 23 | 57 |
| NS755 | Australia | Oceania | 1996 | invasive, NOS | emm90.5 | 184 | 20 | 20 | 14 | 19 | 2 | 23 | 25 | 28 | 15 | 3 | 22 | 25 |
| NS671 | Australia | Oceania | 1995 | invasive, NOS | emm81.3 | 183 | 8 | 4 | 4 | 4 | 3 | 13 | 4 | 42 | 21 | 1 | 4 | 4 |
| NS672 | Australia | Oceania | 1995 | invasive, NOS | emm28.0 | 52 | 3 | 3 | 3 | 3 | 2 | 3 | 3 | 4 | 1 | 1 | 44 | 3 |
| NS684 | Australia | Oceania | 1996 | skin/soft tissue infection, NOS | emm81.2 | 290 | 24 | 50 | 25 | 27 | 2 | 33 | 58 | 64 | 18 | 3 | 28 | 30 |
| NS687 | Australia | Oceania | 1996 | ND | emm58.0 | 176 | 14 | 13 | 10 | 29 | 13 | 13 | 15 | 47 | 10 | 2 | 34 | 31 |
| NS691 | Australia | Oceania | 1996 | skin/soft tissue infection, NOS | emm65.0 | 111 | 22 | 22 | 17 | 38 | 23 | 43 | 28 | 65 | 1 | 1 | 1 | 27 |
| NS702 | Australia | Oceania | 1996 | ARF | emm65.0 | 111 | 22 | 22 | 17 | 38 | 23 | 43 | 28 | 65 | 1 | 1 | 1 | 27 |
| NS718 | Australia | Oceania | 1996 | skin/soft tissue infection, NOS | emm92.0 | 312 | 35 | 40 | 34 | 33 | 39 | 39 | 22 | 24 | 13 | 2 | 20 | 22 |
| NS731 | Australia | Oceania | 1996 | invasive, NOS | emm90.5 | 184 | 20 | 20 | 14 | 19 | 2 | 23 | 25 | 28 | 15 | 3 | 22 | 25 |
| NS5530 | Australia | Oceania | 1993 | ND | emm232.1 | 288 | 18 | 18 | 16 | 16 | 21 | 25 | 27 | 26 | 16 | 3 | 21 | 23 |
| NS804 | Australia | Oceania | 1996 | invasive, NOS | emm97.1 | 216 | 21 | 21 | 15 | 8 | 46 | 24 | 26 | 30 | 8 | 10 | 12 | 26 |
| NS833 | Australia | Oceania | 1996 | ND | emm221.0 | 113 | 11 | 49 | 2 | 37 | 2 | 32 | 57 | 42 | 1 | 2 | 23 | 57 |
| NS836 | Australia | Oceania | 1996 | ND | emm221.0 | 113 | 11 | 49 | 2 | 37 | 2 | 32 | 57 | 42 | 1 | 2 | 23 | 57 |
| NS930 | Australia | Oceania | 1996 | pharyngitis and/or tonsillitis | emm65.0 | 111 | 22 | 22 | 17 | 38 | 23 | 43 | 28 | 65 | 1 | 1 | 1 | 27 |
| NS5341 | Australia | Oceania | 1991 | invasive, NOS | emm85.0 | 109 | 10 | 10 | 4 | 9 | 9 | 11 | 10 | 11 | 7 | 3 | 10 | 10 |
| NS5347 | Australia | Oceania | 1991 | skin/soft tissue infection, NOS | emm4.5 | 289 | 14 | 51 | 42 | 39 | 47 | 33 | 59 | 66 | 8 | 13 | 45 | 58 |
| NS35 | Australia | Oceania | 1991 | invasive, NOS | emm58.0 | 176 | 14 | 13 | 10 | 29 | 13 | 13 | 15 | 47 | 10 | 2 | 34 | 31 |
| NS473 | Australia | Oceania | 1995 | invasive, NOS | emm232.0 | 110 | 18 | 18 | 16 | 16 | 21 | 42 | 27 | 26 | 25 | 3 | 21 | 51 |
| NS474 | Australia | Oceania | 1995 | invasive, NOS | emm58.0 | 176 | 14 | 13 | 10 | 29 | 13 | 24 | 15 | 47 | 10 | 2 | 34 | 31 |
| NS476 | Australia | Oceania | 1995 | invasive, NOS | ND | 1084 | 16 | 16 | 3 | 16 | 17 | 46 | 20 | 22 | 1 | 2 | 46 | 3 |
| NS351 | Australia | Oceania | 1995 | ND | emm58.0 | 176 | 14 | 13 | 10 | 29 | 13 | 24 | 15 | 47 | 10 | 2 | 34 | 31 |
| NS365 | Australia | Oceania | 1995 | invasive, NOS | emm58.0 | 176 | 14 | 13 | 10 | 29 | 13 | 13 | 15 | 47 | 10 | 2 | 34 | 31 |
| NS414 | Australia | Oceania | 1995 | skin/soft tissue infection, NOS | emm11.0 | ND | 1 | 6 | 12 | 1 | 20 | 20 | 13 | 25 | 8 | 3 | 21 | 23 |
| NS415 | Australia | Oceania | 1995 | skin/soft tissue infection, NOS | emm4.5 | 177 | 36 | 42 | 36 | 34 | 41 | 7 | 50 | 57 | 18 | 3 | 36 | 47 |
| NS43 | Australia | Oceania | 1991 | pharyngitis and/or tonsillitis | emm25.0 | 600 | 30 | 11 | 7 | 40 | 48 | 31 | 60 | 67 | 1 | 1 | 47 | 40 |
| NS436 | Australia | Oceania | 1995 | ND | emm11.0 | ND | 1 | 6 | 12 | 1 | 20 | 20 | 13 | 25 | 8 | 3 | 21 | 23 |
| NS452 | Australia | Oceania | 1995 | skin/soft tissue infection, NOS | emm25.0 | 191 | 30 | 35 | 26 | 28 | 35 | 34 | 42 | 45 | 11 | 2 | 12 | 40 |
| NS455 | Australia | Oceania | 1995 | invasive, NOS | emm52.1 | 180 | 15 | 17 | 10 | 17 | 18 | 18 | 21 | 23 | 12 | 8 | 19 | 21 |
| NS6 | Australia | Oceania | 1990 | invasive, NOS | emm123.0 | 325 | 6 | 6 | 6 | 6 | 5 | 6 | 6 | 7 | 5 | 4 | 6 | 6 |
| NS8 | Australia | Oceania | 1990 | skin/soft tissue infection, NOS | emm85.0 | 109 | 10 | 10 | 4 | 9 | 9 | 11 | 10 | 11 | 7 | 3 | 10 | 10 |
| NS16 | Australia | Oceania | 1991 | ND | emm77.0 | 166 | 14 | 25 | 3 | 22 | 27 | 13 | 26 | 29 | 8 | 3 | 25 | 30 |
| NS205 | Australia | Oceania | 1994 | invasive, NOS | emm56.1 | 115 | 14 | 44 | 24 | 2 | 12 | 41 | 51 | 22 | 23 | 1 | 6 | 49 |
| NS232 | Australia | Oceania | 1994 | pharyngitis and/or tonsillitis | emm49.3 | 190 | 4 | 8 | 4 | 11 | 11 | 14 | 12 | 14 | 9 | 2 | 12 | 13 |
| NS234 | Australia | Oceania | 1994 | pharyngitis and/or tonsillitis | emm42.0 | 80 | 37 | 32 | 37 | 35 | 43 | 43 | 44 | 60 | 26 | 2 | 39 | 52 |
| NS237 | Australia | Oceania | 1994 | ND | emm97.0 | 203 | 11 | 49 | 2 | 37 | 2 | 32 | 61 | 42 | 1 | 2 | 23 | 57 |
| NS343 | Australia | Oceania | 1995 | skin/soft tissue infection, NOS | emm225.0 | 48 | 28 | 33 | 18 | 6 | 49 | 6 | 62 | 68 | 29 | 2 | 48 | 59 |
| NS501 | Australia | Oceania | 1995 | invasive, NOS | emm14.4 | 118 | 11 | 5 | 38 | 8 | 15 | 16 | 17 | 13 | 1 | 2 | 40 | 53 |
| NS514 | Australia | Oceania | 1995 | ND | emm73.0 | 331 | 14 | 52 | 3 | 41 | 50 | 13 | 3 | 69 | 7 | 2 | 49 | 60 |
| NS678 | Australia | Oceania | 1995 | pharyngitis and/or tonsillitis | emm4.0 | 39 | 4 | 8 | 4 | 7 | 51 | 9 | 63 | 9 | 7 | 2 | 8 | 8 |
| NS909 | Australia | Oceania | 1996 | invasive, NOS | emm81.0 | 117 | 40 | 53 | 43 | 42 | 52 | 47 | 64 | 70 | 30 | 14 | 30 | 61 |
| NS1043 | Australia | Oceania | 1997 | skin/soft tissue infection, NOS | emm222.0 | 116 | 21 | 54 | 44 | 1 | 35 | 24 | 26 | 71 | 8 | 15 | 12 | 62 |
| NS1051 | Australia | Oceania | 1997 | invasive, NOS | emm114.2 | 189 | 41 | 28 | 40 | 43 | 53 | 40 | 65 | 72 | 18 | 3 | 50 | 63 |
| NS1185 | Australia | Oceania | 1997 | invasive, NOS | emm80.2 | 10 | 16 | 16 | 3 | 16 | 17 | 46 | 20 | 22 | 1 | 2 | 46 | 3 |
| NS1196 | Australia | Oceania | 1997 | skin/soft tissue infection, NOS | emm60.4 | 193 | 14 | 34 | 25 | 27 | 2 | 33 | 41 | 44 | 1 | 2 | 33 | 39 |

| | | | | | | | | | | | | | | | | | | |
|---|---|---|---|---|---|---|---|---|---|---|---|---|---|---|---|---|---|---|
| NS1353 | Australia | Oceania | 1998 | skin/soft tissue infection, NOS | emm233.1 | 206 | 11 | 49 | 2 | 37 | 54 | 32 | 61 | 42 | 31 | 1 | 51 | 25 |
| NS1425 | Australia | Oceania | 1998 | skin/soft tissue infection, NOS | emm13.0 | 286 | 14 | 13 | 10 | 8 | 13 | 13 | 15 | 17 | 10 | 2 | 15 | 16 |
| NS1438 | Australia | Oceania | 1997 | skin/soft tissue infection, NOS | emm71.2 | 318 | 19 | 20 | 13 | 19 | 2 | 22 | 3 | 27 | 1 | 1 | 4 | 24 |
| NS1498 | Australia | Oceania | 1998 | skin/soft tissue infection, NOS | emm217.0 | 594 | 42 | 55 | 3 | 6 | 5 | 48 | 66 | 73 | 1 | 2 | 52 | 11 |
| NS1691 | Australia | Oceania | 1999 | skin/soft tissue infection, NOS | emm75.3 | 590 | 10 | 56 | 45 | 6 | 55 | 2 | 67 | 74 | 1 | 2 | 30 | 31 |
| NS1932 | Australia | Oceania | 1999 | skin/soft tissue infection, NOS | emm22.0 | 175 | 33 | 11 | 7 | 10 | 35 | 36 | 44 | 49 | 19 | 2 | 30 | 64 |
| NS1944 | Australia | Oceania | 1999 | skin/soft tissue infection, NOS | emm18.12 | 535 | 43 | 57 | 2 | 2 | 2 | 2 | 67 | 75 | 8 | 3 | 53 | 2 |
| NS1951 | Australia | Oceania | 1999 | pharyngitis and/or tonsillitis | emm90.2 | 418 | 14 | 13 | 10 | 8 | 13 | 13 | 15 | 17 | 10 | 2 | 15 | 16 |
| NS1980 | Australia | Oceania | 1999 | skin/soft tissue infection, NOS | emm77.0 | 572 | 14 | 25 | 3 | 39 | 56 | 13 | 68 | 76 | 32 | 3 | 54 | 65 |
| NS1984 | Australia | Oceania | 1999 | skin/soft tissue infection, NOS | emm82.0 | 26 | 14 | 58 | 46 | 1 | 6 | 18 | 69 | 77 | 1 | 1 | 1 | 3 |
| NS1988 | Australia | Oceania | 1999 | ND | emm9.0 | 603 | 44 | 39 | 33 | 32 | 2 | 1 | 49 | 54 | 5 | 3 | 11 | 45 |
| NS1999 | Australia | Oceania | 1999 | skin/soft tissue infection, NOS | emm75.3 | 590 | 10 | 56 | 45 | 6 | 55 | 2 | 67 | 74 | 1 | 2 | 30 | 31 |
| NS2000 | Australia | Oceania | 1999 | skin/soft tissue infection, NOS | emm77.0 | 63 | 23 | 24 | 19 | 1 | 26 | 28 | 30 | 33 | 8 | 3 | 24 | 29 |
| NS2012 | Australia | Oceania | 1999 | skin/soft tissue infection, NOS | emm99.5 | 593 | 11 | 32 | 47 | 1 | 57 | 32 | 61 | 32 | 1 | 2 | 55 | 66 |
| NS2023 | Australia | Oceania | 1999 | pharyngitis and/or tonsillitis | emm22.0 | 46 | 7 | 5 | 5 | 5 | 4 | 5 | 5 | 6 | 4 | 3 | 5 | 5 |
| NS2128 | Australia | Oceania | 2000 | ND | emm192.0 | 612 | 11 | 5 | 10 | 44 | 57 | 49 | 70 | 42 | 1 | 2 | 23 | 31 |
| NS2231 | Australia | Oceania | 2000 | skin/soft tissue infection, NOS | emm49.3 | 190 | 4 | 8 | 4 | 11 | 11 | 14 | 12 | 14 | 9 | 2 | 12 | 13 |
| NS2416 | Australia | Oceania | 2000 | skin/soft tissue infection, NOS | emm104.0 | 597 | 45 | 34 | 48 | 8 | 58 | 50 | 71 | 78 | 8 | 3 | 31 | 67 |
| NS2509 | Australia | Oceania | 2000 | skin/soft tissue infection, NOS | emm85.0 | 1020 | 10 | 59 | 4 | 9 | 9 | 51 | 72 | 11 | 7 | 3 | 10 | 68 |
| NS2663 | Australia | Oceania | 2001 | skin/soft tissue infection, NOS | emm53.0 | 363 | 46 | 32 | 7 | 1 | 6 | 7 | 73 | 79 | 1 | 16 | 56 | 31 |
| NS2664 | Australia | Oceania | 2001 | skin/soft tissue infection, NOS | emm89.14 | 812 | 14 | 49 | 20 | 16 | 59 | 52 | 74 | 80 | 33 | 11 | 57 | 25 |
| NS2671 | Australia | Oceania | 2001 | skin/soft tissue infection, NOS | emm102.7 | 287 | 1 | 60 | 7 | 45 | 60 | 19 | 75 | 79 | 1 | 2 | 18 | 69 |
| NS2676 | Australia | Oceania | 2001 | skin/soft tissue infection, NOS | emm232.0 | 185 | 14 | 37 | 31 | 3 | 2 | 38 | 47 | 81 | 1 | 2 | 58 | 25 |
| NS2679 | Australia | Oceania | 1990 | skin/soft tissue infection, NOS | emm74.0 | 120 | 28 | 13 | 3 | 41 | 50 | 53 | 76 | 82 | 11 | 4 | 59 | 70 |
| NS2763 | Australia | Oceania | 2001 | invasive, NOS | emm70.0 | 10 | 16 | 16 | 3 | 16 | 17 | 7 | 20 | 22 | 1 | 2 | 18 | 3 |
| NS2811 | Australia | Oceania | 2001 | skin/soft tissue infection, NOS | emm9.0 | 204 | 34 | 39 | 33 | 32 | 2 | 1 | 49 | 54 | 5 | 3 | 11 | 45 |
| NS3003 | Australia | Oceania | 2001 | skin/soft tissue infection, NOS | emm225.0 | 48 | 28 | 33 | 18 | 6 | 49 | 6 | 62 | 68 | 29 | 2 | 48 | 59 |
| NS3275 | Australia | Oceania | 2002 | skin/soft tissue infection, NOS | emm8.0 | 602 | 18 | 18 | 7 | 10 | 10 | 7 | 77 | 39 | 19 | 2 | 30 | 31 |
| NS3309 | Australia | Oceania | 2002 | invasive, NOS | emm86.1 | 4 | 47 | 5 | 10 | 46 | 15 | 16 | 17 | 13 | 1 | 7 | 60 | 71 |
| NS3332 | Australia | Oceania | 2002 | skin/soft tissue infection, NOS | emm108.0 | 304 | 48 | 5 | 10 | 20 | 15 | 16 | 31 | 13 | 1 | 2 | 16 | 31 |
| NS3335 | Australia | Oceania | 2002 | skin/soft tissue infection, NOS | emm39.4 | 268 | 1 | 60 | 49 | 45 | 61 | 19 | 13 | 83 | 34 | 3 | 61 | 29 |
| NS3473 | Australia | Oceania | 2002 | ND | emm82.0 | 26 | 14 | 58 | 46 | 1 | 6 | 18 | 69 | 77 | 1 | 1 | 1 | 3 |
| NS4505 | Australia | Oceania | 2004 | skin/soft tissue infection, NOS | emm110.0 | 179 | 14 | 33 | 18 | 6 | 25 | 27 | 40 | 43 | 1 | 2 | 18 | 38 |
| NS4517 | Australia | Oceania | 2004 | skin/soft tissue infection, NOS | emm53.0 | 11 | 49 | 5 | 10 | 8 | 15 | 54 | 17 | 84 | 1 | 2 | 6 | 20 |
| NS4518 | Australia | Oceania | 2004 | ND | emm12.1 | 36 | 14 | 3 | 3 | 3 | 62 | 3 | 52 | 59 | 24 | 12 | 38 | 3 |
| NS4618 | Australia | Oceania | 2004 | ND | emm76.6 | 291 | 39 | 46 | 39 | 36 | 45 | 45 | 54 | 85 | 2 | 2 | 62 | 54 |
| NS4624 | Australia | Oceania | 2004 | skin/soft tissue infection, NOS | emm19.4 | 616 | 50 | 32 | 10 | 1 | 6 | 55 | 78 | 13 | 1 | 2 | 6 | 31 |
| NS4697 | Australia | Oceania | 2005 | ND | emm1.0 | 28 | 1 | 1 | 1 | 1 | 1 | 1 | 1 | 1 | 1 | 1 | 1 | 1 |
| NS4706 | Australia | Oceania | 2005 | ND | emm13.0 | 286 | 14 | 13 | 10 | 8 | 13 | 13 | 15 | 17 | 10 | 2 | 15 | 16 |
| NS4716 | Australia | Oceania | 2005 | ND | emm91.1 | 12 | 11 | 5 | 10 | 15 | 15 | 16 | 17 | 13 | 1 | 2 | 16 | 18 |
| NS4743 | Australia | Oceania | 2005 | ND | emm58.8 | 549 | 51 | 61 | 50 | 47 | 63 | 56 | 79 | 86 | 35 | 3 | 35 | 72 |
| NS4766 | Australia | Oceania | 2005 | skin/soft tissue infection, NOS | emm183.2 | 586 | 11 | 62 | 3 | 22 | 27 | 13 | 26 | 29 | 8 | 3 | 25 | 30 |
| NS3523 | Australia | Oceania | 2002 | ND | emm197.0 | 615 | 11 | 63 | 32 | 24 | 64 | 57 | 80 | 42 | 15 | 3 | 23 | 57 |
| NS4797 | Australia | Oceania | 2005 | skin/soft tissue infection, NOS | emm100.0 | 119 | 52 | 11 | 7 | 10 | 10 | 7 | 37 | 39 | 19 | 2 | 30 | 31 |
| NS4800 | Australia | Oceania | 2005 | skin/soft tissue infection, NOS | emm97.1 | 216 | 21 | 21 | 15 | 8 | 46 | 24 | 26 | 30 | 8 | 10 | 12 | 26 |
| NS4804 | Australia | Oceania | 2005 | skin/soft tissue infection, NOS | emm81.2 | 608 | 11 | 11 | 7 | 48 | 65 | 13 | 81 | 79 | 21 | 1 | 4 | 4 |
| NS4883 | Australia | Oceania | 2005 | skin/soft tissue infection, NOS | emm103.0 | 611 | 14 | 13 | 3 | 1 | 66 | 13 | 82 | 87 | 1 | 2 | 49 | 4 |
| NS4982 | Australia | Oceania | 2005 | skin/soft tissue infection, NOS | emm42.0 | 80 | 37 | 32 | 37 | 35 | 43 | 43 | 44 | 60 | 26 | 2 | 39 | 52 |

| | | | | | | | | | | | | | | | | | | |
|---|---|---|---|---|---|---|---|---|---|---|---|---|---|---|---|---|---|---|
| NS4984 | Australia | Oceania | 2005 | skin/soft tissue infection, NOS | emm124.0 | 544 | 18 | 18 | 4 | 16 | 21 | 21 | 23 | 26 | 8 | 9 | 21 | 23 |
| NS4985 | Australia | Oceania | 2005 | skin/soft tissue infection, NOS | emm14.4 | 118 | 11 | 5 | 38 | 8 | 15 | 16 | 17 | 13 | 1 | 2 | 40 | 53 |
| NS5025 | Australia | Oceania | 2005 | ND | emm105.0 | 954 | 6 | 6 | 6 | 6 | 5 | 12 | 6 | 12 | 5 | 4 | 6 | 11 |
| NS5048 | Australia | Oceania | 2005 | ND | emm55.0 | 100 | 11 | 11 | 7 | 10 | 10 | 13 | 11 | 13 | 8 | 3 | 11 | 12 |
| NS5128 | Australia | Oceania | 2005 | ND | emm77.0 | 588 | 14 | 25 | 51 | 39 | 56 | 13 | 68 | 76 | 32 | 3 | 54 | 65 |
| NS3785 | Australia | Oceania | 2003 | ND | emm12.0 | 36 | 14 | 3 | 3 | 3 | 2 | 3 | 52 | 59 | 24 | 12 | 38 | 3 |
| NS5146 | Australia | Oceania | 2005 | ND | emm78.3 | 181 | 11 | 32 | 52 | 1 | 67 | 13 | 83 | 88 | 1 | 2 | 63 | 73 |
| NS5340 | Australia | Oceania | 1991 | skin/soft tissue infection, NOS | emm65.4 | 1021 | 53 | 64 | 53 | 35 | 68 | 43 | 44 | 89 | 2 | 2 | 39 | 2 |
| NS5378 | Australia | Oceania | 1991 | skin/soft tissue infection, NOS | emm124.0 | 544 | 18 | 18 | 4 | 16 | 69 | 21 | 23 | 26 | 8 | 3 | 21 | 23 |
| NS5427 | Australia | Oceania | 1991 | ND | emm63.0 | 598 | 5 | 65 | 34 | 27 | 39 | 13 | 84 | 90 | 13 | 2 | 20 | 22 |
| NS5434 | Australia | Oceania | 1991 | ND | emm57.2 | 33 | 11 | 14 | 11 | 8 | 16 | 17 | 18 | 20 | 11 | 7 | 17 | 19 |
| NS5496 | Australia | Oceania | 1992 | skin/soft tissue infection, NOS | emm77.0 | 994 | 20 | 66 | 40 | 38 | 70 | 6 | 25 | 91 | 18 | 3 | 22 | 25 |
| NS5539 | Australia | Oceania | 1993 | skin/soft tissue infection, NOS | emm33.0 | 1077 | 54 | 11 | 7 | 10 | 35 | 36 | 85 | 79 | 1 | 2 | 6 | 20 |
| NS5544 | Australia | Oceania | 1993 | skin/soft tissue infection, NOS | emm76.0 | 617 | 2 | 67 | 54 | 49 | 36 | 9 | 86 | 92 | 1 | 2 | 49 | 4 |
| NS5568 | Australia | Oceania | 1992 | skin/soft tissue infection, NOS | emm77.0 | 572 | 14 | 25 | 3 | 39 | 56 | 13 | 68 | 76 | 32 | 3 | 54 | 65 |
| NS5578 | Australia | Oceania | 1992 | skin/soft tissue infection, NOS | emm123.0 | 610 | 6 | 6 | 6 | 6 | 5 | 6 | 6 | 7 | 5 | 4 | 6 | 6 |
| NS3844 | Australia | Oceania | 2004 | ND | emm25.1 | 192 | 14 | 30 | 24 | 2 | 33 | 13 | 13 | 40 | 36 | 3 | 31 | 36 |
| NS5749 | Australia | Oceania | 1987 | skin/soft tissue infection, NOS | emm238.2 | 372 | 1 | 1 | 1 | 1 | 1 | 15 | 14 | 93 | 5 | 6 | 14 | 15 |
| NS5582 | Australia | Oceania | 2005 | ND | emm18.12 | 535 | 43 | 57 | 2 | 2 | 2 | 2 | 67 | 75 | 8 | 3 | 53 | 2 |
| NS5615 | Australia | Oceania | 2005 | skin/soft tissue infection, NOS | emm33.0 | 3 | 11 | 11 | 7 | 10 | 35 | 36 | 85 | 94 | 1 | 2 | 6 | 20 |
| NS5630 | Australia | Oceania | 2005 | skin/soft tissue infection, NOS | emm98.0 | 182 | 15 | 31 | 10 | 20 | 15 | 31 | 38 | 41 | 1 | 3 | 32 | 37 |
| NS5694 | Australia | Oceania | 2005 | skin/soft tissue infection, NOS | emm230.0 | 205 | 1 | 15 | 10 | 1 | 10 | 18 | 19 | 21 | 12 | 2 | 6 | 20 |
| NS5758 | Australia | Oceania | 2005 | skin/soft tissue infection, NOS | emm58.0 | 176 | 14 | 13 | 10 | 8 | 13 | 13 | 15 | 47 | 10 | 2 | 34 | 31 |
| NS5778 | Australia | Oceania | 2005 | skin/soft tissue infection, NOS | emm98.1 | 205 | 15 | 15 | 10 | 1 | 10 | 18 | 19 | 21 | 12 | 2 | 6 | 20 |
| NS5779 | Australia | Oceania | 2005 | skin/soft tissue infection, NOS | emm108.0 | 304 | 48 | 5 | 10 | 20 | 15 | 16 | 31 | 13 | 1 | 2 | 16 | 31 |
| NS3850 | Australia | Oceania | 2003 | ND | emm11.0 | ND | 1 | 6 | 12 | 1 | 20 | 20 | 13 | 25 | 8 | 3 | 21 | 23 |
| NS3870 | Australia | Oceania | 2004 | ND | emm75.1 | 609 | 27 | 28 | 55 | 50 | 71 | 45 | 35 | 37 | 18 | 3 | 28 | 30 |
| NS4147 | Australia | Oceania | 2004 | ND | emm58.0 | 176 | 14 | 13 | 10 | 29 | 13 | 13 | 15 | 47 | 10 | 2 | 34 | 31 |
| NS4150 | Australia | Oceania | 2004 | ND | emm22.0 | 46 | 7 | 5 | 5 | 5 | 4 | 5 | 5 | 6 | 4 | 3 | 5 | 5 |
| NS4155 | Australia | Oceania | 2004 | ND | emm71.0 | 318 | 19 | 20 | 13 | 19 | 2 | 22 | 3 | 27 | 1 | 1 | 4 | 24 |
| NS5782 | Australia | Oceania | 2005 | skin/soft tissue infection, NOS | emm89.0 | 101 | 2 | 2 | 2 | 2 | 2 | 2 | 2 | 2 | 2 | 2 | 2 | 2 |
| NS5789 | Australia | Oceania | 2005 | skin/soft tissue infection, NOS | emm4.0 | 39 | 4 | 8 | 4 | 7 | 7 | 9 | 8 | 9 | 37 | 2 | 8 | 8 |
| NS5958 | Australia | Oceania | 2005 | skin/soft tissue infection, NOS | emm56.0 | 115 | 14 | 44 | 24 | 2 | 12 | 41 | 51 | 22 | 23 | 1 | 6 | 49 |
| NS6033 | Australia | Oceania | 2006 | ND | emm87.0 | 541 | 55 | 68 | 10 | 1 | 72 | 58 | 87 | 95 | 1 | 2 | 52 | 74 |
| NS6078 | Australia | Oceania | 2006 | skin/soft tissue infection, NOS | ND | 92 | 1 | 1 | 1 | 1 | 1 | 15 | 14 | 96 | 5 | 6 | 14 | 15 |
| NS6087 | Australia | Oceania | 2006 | skin/soft tissue infection, NOS | emm114.2 | 140 | 11 | 69 | 7 | 10 | 73 | 7 | 88 | 97 | 15 | 3 | 64 | 34 |
| NS6142 | Australia | Oceania | 2006 | skin/soft tissue infection, NOS | emm183.2 | 166 | 14 | 25 | 3 | 22 | 27 | 13 | 26 | 29 | 8 | 3 | 25 | 30 |
| NS6185 | Australia | Oceania | 2006 | skin/soft tissue infection, NOS | emm15.1 | 872 | 14 | 48 | 41 | 2 | 2 | 2 | 56 | 13 | 8 | 3 | 43 | 56 |
| NS6187 | Australia | Oceania | 2006 | skin/soft tissue infection, NOS | emm13.0 | 286 | 14 | 13 | 10 | 8 | 13 | 13 | 15 | 17 | 10 | 2 | 15 | 16 |
| NS6221 | Australia | Oceania | 2006 | skin/soft tissue infection, NOS | emm100.5 | 119 | 11 | 11 | 7 | 10 | 10 | 7 | 37 | 39 | 19 | 2 | 30 | 31 |
| NS6263 | Australia | Oceania | 2006 | skin/soft tissue infection, NOS | emm232.0 | 185 | 14 | 37 | 31 | 3 | 2 | 38 | 47 | 81 | 1 | 2 | 58 | 25 |
| NS6441 | Australia | Oceania | 2006 | skin/soft tissue infection, NOS | emm70.0 | 10 | 16 | 16 | 3 | 16 | 17 | 7 | 20 | 22 | 1 | 2 | 18 | 3 |
| NS6474 | Australia | Oceania | 2006 | skin/soft tissue infection, NOS | emm44.0 | 178 | 5 | 7 | 7 | 3 | 2 | 3 | 24 | 8 | 14 | 3 | 7 | 7 |
| NS6531 | Australia | Oceania | 2006 | skin/soft tissue infection, NOS | emm91.0 | 12 | 11 | 5 | 10 | 15 | 15 | 16 | 17 | 13 | 1 | 2 | 16 | 18 |
| NS6570 | Australia | Oceania | 2007 | skin/soft tissue infection, NOS | emm82.1 | 591 | 56 | 70 | 4 | 1 | 20 | 2 | 89 | 98 | 38 | 2 | 65 | 3 |
| NS6595 | Australia | Oceania | 2007 | skin/soft tissue infection, NOS | emm123.0 | 610 | 6 | 6 | 6 | 6 | 5 | 6 | 6 | 7 | 5 | 4 | 6 | 6 |
| NS6648 | Australia | Oceania | 2007 | invasive, NOS | emm41.2 | 579 | 11 | 5 | 10 | 51 | 15 | 16 | 77 | 99 | 1 | 2 | 6 | 20 |
| NS6749 | Australia | Oceania | 2007 | skin/soft tissue infection, NOS | emm54.1 | 302 | 11 | 5 | 10 | 20 | 15 | 16 | 17 | 13 | 1 | 2 | 16 | 31 |

| | | | | | | | | | | | | | | | | | | |
|---|---|---|---|---|---|---|---|---|---|---|---|---|---|---|---|---|---|---|
| NS6770 | Australia | Oceania | 2007 | skin/soft tissue infection, NOS | emm11.0 | ND | 1 | 6 | 12 | 1 | 20 | 20 | 13 | 25 | 8 | 3 | 21 | 23 |
| NS6783 | Australia | Oceania | 2007 | skin/soft tissue infection, NOS | emm65.2 | 585 | 15 | 15 | 10 | 1 | 74 | 7 | 7 | 8 | 39 | 3 | 7 | 75 |
| NS6845 | Australia | Oceania | 2007 | skin/soft tissue infection, NOS | emm178.0 | 592 | 11 | 71 | 7 | 41 | 75 | 32 | 90 | 42 | 1 | 2 | 23 | 76 |
| NS6930 | Australia | Oceania | 2007 | skin/soft tissue infection, NOS | emm99.5 | 593 | 11 | 32 | 47 | 1 | 57 | 32 | 61 | 32 | 1 | 2 | 55 | 66 |
| NS6983 | Australia | Oceania | 2007 | skin/soft tissue infection, NOS | emm74.0 | 120 | 28 | 13 | 3 | 41 | 50 | 53 | 76 | 82 | 11 | 4 | 59 | 70 |
| NS6989 | Australia | Oceania | 2007 | skin/soft tissue infection, NOS | emm101.0 | 182 | 15 | 31 | 10 | 20 | 15 | 31 | 38 | 41 | 1 | 3 | 32 | 37 |
| NS7048 | Australia | Oceania | 2007 | skin/soft tissue infection, NOS | emm24.1 | 582 | 14 | 43 | 33 | 1 | 42 | 13 | 32 | 58 | 22 | 8 | 37 | 48 |
| NS7056 | Australia | Oceania | 2007 | skin/soft tissue infection, NOS | emm116.1 | 227 | 57 | 72 | 33 | 1 | 42 | 13 | 32 | 100 | 5 | 4 | 11 | 77 |
| NS7092 | Australia | Oceania | 1997 | skin/soft tissue infection, NOS | emm78.3 | 181 | 11 | 32 | 52 | 1 | 67 | 13 | 83 | 88 | 1 | 2 | 63 | 73 |
| NS7096 | Australia | Oceania | 1997 | skin/soft tissue infection, NOS | emm53.0 | 11 | 49 | 5 | 10 | 8 | 15 | 54 | 17 | 84 | 1 | 2 | 6 | 20 |
| NS7098 | Australia | Oceania | 1997 | skin/soft tissue infection, NOS | emm86.1 | 4 | 47 | 5 | 10 | 46 | 15 | 16 | 17 | 13 | 1 | 7 | 60 | 71 |
| NS7124 | Australia | Oceania | 1997 | ND | emm124.0 | 199 | 18 | 18 | 4 | 16 | 21 | 21 | 23 | 101 | 8 | 3 | 21 | 23 |
| NS7133 | Australia | Oceania | 1997 | ND | emm165.0 | 112 | 58 | 73 | 56 | 1 | 76 | 59 | 27 | 102 | 8 | 3 | 61 | 29 |
| NS7158 | Australia | Oceania | 1999 | skin/soft tissue infection, NOS | emm99.0 | 141 | 14 | 41 | 35 | 1 | 20 | 40 | 26 | 56 | 5 | 4 | 20 | 46 |
| NS7160 | Australia | Oceania | 1999 | ND | emm92.0 | 599 | 5 | 65 | 34 | 27 | 39 | 39 | 22 | 103 | 13 | 2 | 20 | 22 |
| NS7168 | Australia | Oceania | 1999 | skin/soft tissue infection, NOS | emm4.5 | 606 | 59 | 74 | 40 | 43 | 77 | 60 | 65 | 72 | 18 | 2 | 11 | 78 |
| NS7175 | Australia | Oceania | 1999 | skin/soft tissue infection, NOS | emm192.0 | 612 | 11 | 5 | 10 | 44 | 57 | 49 | 70 | 42 | 1 | 2 | 23 | 31 |
| NS7185 | Australia | Oceania | 1999 | ND | emm238.1 | 1022 | 60 | 1 | 1 | 1 | 1 | 15 | 14 | 104 | 1 | 2 | 30 | 15 |
| NS7249 | Australia | Oceania | 1999 | ND | emm75.3 | 590 | 10 | 56 | 45 | 6 | 55 | 2 | 67 | 74 | 1 | 2 | 30 | 31 |
| NS7259 | Australia | Oceania | 1999 | ND | emm192.0 | 612 | 11 | 5 | 10 | 44 | 57 | 49 | 70 | 42 | 1 | 2 | 23 | 31 |
| NS7970 | Australia | Oceania | 1995 | skin/soft tissue infection, NOS | emm41.2 | 296 | 11 | 71 | 7 | 1 | 78 | 32 | 91 | 42 | 1 | 2 | 23 | 57 |
| NS8042 | Australia | Oceania | 1995 | ND | emm89.0 | 142 | 61 | 8 | 3 | 11 | 79 | 14 | 92 | 18 | 9 | 2 | 12 | 13 |
| NS8145 | Australia | Oceania | 1995 | ND | emm22.0 | 175 | 33 | 11 | 7 | 10 | 35 | 36 | 44 | 49 | 19 | 2 | 30 | 43 |
| NS10015 | Australia | Oceania | 1987 | skin/soft tissue infection, NOS | emm97.0 | 197 | 11 | 5 | 10 | 1 | 57 | 32 | 93 | 105 | 1 | 2 | 23 | 79 |
| patientB | Australia | Oceania | 2010 | ND | emm1.0 | 28 | 1 | 1 | 1 | 1 | 1 | 1 | 1 | 1 | 1 | 1 | 1 | 1 |
| patientC | Australia | Oceania | 2010 | ND | emm77.0 | 1023 | 23 | 24 | 57 | 1 | 26 | 28 | 30 | 33 | 8 | 3 | 24 | 29 |
| patientG | Australia | Oceania | 2010 | ND | emm89.0 | 101 | 2 | 2 | 2 | 2 | 2 | 2 | 2 | 2 | 2 | 2 | 2 | 2 |
| patientH | Australia | Oceania | 2010 | invasive, NOS | emm75.0 | 150 | 25 | 26 | 20 | 24 | 29 | 29 | 33 | 48 | 15 | 3 | 12 | 33 |
| patientI | Australia | Oceania | 2010 | ND | emm89.0 | 101 | 2 | 2 | 2 | 2 | 2 | 2 | 2 | 2 | 2 | 2 | 2 | 2 |
| GAS_dog | Australia | Oceania | 1990 | ND | emm70.0 | 10 | 16 | 16 | 3 | 16 | 17 | 7 | 20 | 22 | 1 | 2 | 18 | 3 |
| GAS_dogM3 | Australia | Oceania | 1990 | ND | emm70.0 | 10 | 16 | 16 | 3 | 16 | [S] | 7 | 20 | [S] | 1 | 2 | 18 | 3 |
| GAS_dog48 | Australia | Oceania | 1990 | ND | emm91.0 | 12 | 11 | 5 | 10 | 15 | 15 | 16 | 17 | 13 | 1 | 2 | 16 | 18 |
| 1021-1 | Australia | Oceania | 2004 | skin/soft tissue infection, NOS | emm70.0 | 10 | 16 | 16 | 3 | 16 | 17 | 7 | 20 | 22 | 1 | 2 | 18 | 3 |
| 1024-1 | Australia | Oceania | 2004 | skin/soft tissue infection, NOS | emm53.0 | 11 | 49 | 5 | 10 | 8 | 15 | 54 | 17 | 84 | 1 | 2 | 6 | 20 |
| 2095-1 | Australia | Oceania | 2004 | skin/soft tissue infection, NOS | emm81.0 | 974 | 62 | 75 | 45 | 52 | 5 | 6 | 45 | 106 | 24 | 17 | 66 | 61 |
| 2098-1 | Australia | Oceania | 2004 | skin/soft tissue infection, NOS | emm58.0 | 176 | 14 | 13 | 10 | 29 | 13 | 13 | 15 | 47 | 10 | 2 | 34 | 31 |
| 2241-1 | Australia | Oceania | 2004 | skin/soft tissue infection, NOS | emm101.0 | 182 | 15 | 31 | 10 | 20 | 15 | 31 | 38 | 107 | 1 | 3 | 32 | 37 |
| 2102-1 | Australia | Oceania | 2004 | skin/soft tissue infection, NOS | emm108.0 | 304 | 48 | 5 | 10 | 20 | 15 | 16 | 31 | 13 | 1 | 2 | 16 | 31 |
| 2103-1 | Australia | Oceania | 2004 | skin/soft tissue infection, NOS | emm207.1 | 332 | 18 | 18 | 16 | 16 | 21 | 25 | 94 | 26 | 8 | 3 | 21 | 80 |
| 2235-1 | Australia | Oceania | 2004 | skin/soft tissue infection, NOS | emm44.0 | 641 | 5 | 7 | 7 | 1 | 81 | 13 | 26 | 108 | 8 | 18 | 67 | 30 |
| 2233-1 | Australia | Oceania | 2004 | skin/soft tissue infection, NOS | emm44.0 | 641 | 5 | 7 | 7 | 1 | 81 | 13 | 26 | 108 | 8 | 18 | 67 | 30 |
| 2088-1 | Australia | Oceania | 2004 | skin/soft tissue infection, NOS | emm230.0 | 205 | 1 | 15 | 10 | 1 | 10 | 18 | 19 | 21 | 12 | 2 | 6 | 81 |
| Bel007 | Belgium | Europe | 2004 | skin/soft tissue infection, NOS | emm184.0 | 174 | 63 | 45 | 3 | 16 | 82 | 13 | 20 | 109 | 1 | 2 | 11 | 78 |
| Bel006 | Belgium | Europe | 2004 | pharyngitis and/or tonsillitis | emm19.4 | 1007 | 50 | 32 | 10 | 1 | 6 | 55 | 78 | 13 | 1 | 19 | 6 | 31 |
| Bel005 | Belgium | Europe | 2004 | pharyngitis and/or tonsillitis | emm8.0 | 59 | 14 | 13 | 3 | 41 | 50 | 13 | 3 | 110 | 7 | 2 | 49 | 4 |
| Bel004 | Belgium | Europe | 2004 | pharyngitis and/or tonsillitis | emm232.0 | 187 | 11 | 32 | 58 | 16 | 38 | 18 | 77 | 111 | 11 | 7 | 60 | 71 |
| Bel003 | Belgium | Europe | 2004 | skin/soft tissue infection, NOS | emm209.0 | 573 | 64 | 76 | 4 | 53 | 21 | 61 | 23 | 26 | 8 | 3 | 21 | 23 |
| Bel002 | Belgium | Europe | 2004 | skin/soft tissue infection, NOS | emm186.2 | 147 | 11 | 11 | 7 | 1 | 83 | 31 | 77 | 79 | 1 | 1 | 1 | 3 |

| | | | | | | | | | | | | | | | | | | |
|---|---|---|---|---|---|---|---|---|---|---|---|---|---|---|---|---|---|---|
| Bel001 | Belgium | Europe | 2004 | pharyngitis and/or tonsillitis | emm75.0 | 150 | 25 | 26 | 20 | 24 | 29 | 29 | 33 | 112 | 15 | 3 | 12 | 33 |
| Bra017 | Brazil | South America | 2011 | scarlet fever | emm22.0 | 46 | 7 | 5 | 5 | 5 | 4 | 5 | 5 | 6 | 4 | 3 | 5 | 5 |
| Bra005 | Brazil | South America | 2013 | skin/soft tissue infection, NOS | emm88.2 | 397 | 14 | 33 | 59 | 6 | 25 | 27 | 40 | 43 | 1 | 2 | 18 | 28 |
| Bra040 | Brazil | South America | 2012 | invasive, NOS | emm9.0 | 75 | 34 | 77 | 60 | 32 | 2 | 1 | 95 | 113 | 5 | 3 | 11 | 45 |
| Bra028 | Brazil | South America | 2011 | scarlet fever | emm22.0 | 46 | 7 | 5 | 5 | 5 | 4 | 5 | 5 | 6 | 4 | 3 | 5 | 5 |
| Bra016 | Brazil | South America | 2012 | pharyngitis and/or tonsillitis | emm80.0 | 538 | 11 | 78 | 61 | 54 | 84 | 9 | 96 | 114 | 40 | 3 | 68 | 3 |
| Bra004 | Brazil | South America | 2012 | pharyngitis and/or tonsillitis | ND | 174 | 14 | 79 | 3 | 26 | 82 | 13 | 20 | 109 | 1 | 2 | 11 | 78 |
| Bra051 | Brazil | South America | 2005 | scarlet fever | emm12.0 | 36 | 14 | 3 | 3 | 3 | 2 | 3 | 52 | 59 | 24 | 12 | 38 | 3 |
| Bra039 | Brazil | South America | 2007 | invasive, NOS | emm6.4 | 37 | 65 | 80 | 62 | 55 | 85 | 62 | 97 | 115 | 11 | 20 | 69 | 82 |
| Bra027 | Brazil | South America | 2011 | scarlet fever | emm22.0 | 46 | 7 | 5 | 5 | 5 | 4 | 5 | 5 | 6 | 4 | 3 | 5 | 5 |
| Bra015 | Brazil | South America | 2007 | invasive, NOS | emm4.0 | 38 | 4 | 8 | 63 | 11 | 86 | 9 | 8 | 9 | 7 | 2 | 8 | 8 |
| Bra003 | Brazil | South America | 2009 | scarlet fever | emm28.0 | 626 | 3 | 3 | 3 | 3 | 87 | 3 | 98 | 4 | 1 | 1 | 3 | 3 |
| Bra050 | Brazil | South America | 2012 | scarlet fever | emm75.0 | 150 | 25 | 26 | 20 | 24 | 29 | 29 | 33 | 112 | 15 | 3 | 12 | 33 |
| Bra038 | Brazil | South America | 2014 | skin/soft tissue infection, NOS | emm119.1 | 1061 | 28 | 32 | 10 | 1 | 88 | 63 | 78 | 13 | 1 | 2 | 6 | 31 |
| Bra026 | Brazil | South America | 2005 | skin/soft tissue infection, NOS | emm15.1 | 999 | 66 | 81 | 7 | 1 | 89 | 64 | 99 | 79 | 1 | 1 | 70 | 3 |
| Bra014 | Brazil | South America | 2003 | invasive, NOS | emm92.0 | 82 | 17 | 4 | 3 | 18 | 19 | 19 | 22 | 24 | 13 | 2 | 20 | 22 |
| Bra002 | Brazil | South America | 2013 | scarlet fever | emm87.0 | 62 | 67 | 15 | 10 | 1 | 72 | 58 | 87 | 95 | 1 | 2 | 52 | 74 |
| Bra049 | Brazil | South America | 2004 | invasive, NOS | emm83.1 | 5 | 11 | 5 | 10 | 8 | 15 | 65 | 97 | 115 | 11 | 2 | 69 | 83 |
| Bra037 | Brazil | South America | 2013 | skin/soft tissue infection, NOS | emm63.0 | 1005 | 68 | 10 | 4 | 37 | 90 | 66 | 72 | 116 | 17 | 11 | 26 | 3 |
| Bra025 | Brazil | South America | 2003 | scarlet fever | emm48.1 | 161 | 69 | 51 | 42 | 39 | 91 | 7 | 100 | 117 | 41 | 2 | 71 | 84 |
| Bra013 | Brazil | South America | 2000 | ARF | emm22.0 | 46 | 7 | 5 | 5 | 5 | 4 | 5 | 5 | 6 | 4 | 3 | 5 | 5 |
| Bra001 | Brazil | South America | 2005 | scarlet fever | emm87.0 | 62 | 67 | 15 | 10 | 1 | 72 | 58 | 87 | 95 | 1 | 2 | 52 | 74 |
| Bra048 | Brazil | South America | 2006 | scarlet fever | emm1.0 | 28 | 1 | 1 | 1 | 1 | 1 | 1 | 1 | 1 | 1 | 1 | 1 | 1 |
| Bra036 | Brazil | South America | 2013 | pharyngitis and/or tonsillitis | ND | 53 | 70 | 4 | 3 | 18 | 19 | 19 | 101 | 118 | 42 | 2 | 72 | 3 |
| Bra024 | Brazil | South America | 2012 | pharyngitis and/or tonsillitis | emm81.0 | 624 | 5 | 53 | 43 | 42 | 52 | 47 | 64 | 119 | 30 | 14 | 30 | 61 |
| Bra012 | Brazil | South America | 2012 | skin/soft tissue infection, NOS | emm43.5 | 1006 | 11 | 5 | 10 | 8 | 15 | 16 | 77 | 120 | 1 | 2 | 6 | 20 |
| Bra047 | Brazil | South America | 2003 | scarlet fever | emm87.0 | 62 | 67 | 15 | 10 | 1 | 72 | 58 | 87 | 95 | 1 | 2 | 52 | 74 |
| Bra035 | Brazil | South America | 2008 | invasive, NOS | emm66.0 | 44 | 14 | 82 | 64 | 31 | 6 | 67 | 69 | 77 | 1 | 1 | 73 | 3 |
| Bra023 | Brazil | South America | 2000 | pharyngitis and/or tonsillitis | emm86.1 | 4 | 11 | 5 | 10 | 20 | 15 | 16 | 17 | 13 | 1 | 7 | 60 | 71 |
| Bra011 | Brazil | South America | 2011 | scarlet fever | emm12.0 | 36 | 14 | 3 | 3 | 3 | 2 | 3 | 52 | 59 | 24 | 12 | 38 | 3 |
| Bra046 | Brazil | South America | 2004 | APSGN | emm94.1 | 89 | 8 | 4 | 4 | 4 | 3 | 13 | 4 | 42 | 21 | 1 | 4 | 4 |
| Bra034 | Brazil | South America | 2009 | invasive, NOS | emm118.0 | 1075 | 14 | 83 | 3 | 3 | 92 | 7 | 25 | 121 | 36 | 4 | 74 | 25 |
| Bra022 | Brazil | South America | 2002 | pharyngitis and/or tonsillitis | emm59.0 | 172 | 71 | 84 | 24 | 2 | 93 | 68 | 102 | 122 | 18 | 3 | 75 | 85 |
| Bra010 | Brazil | South America | 2003 | pharyngitis and/or tonsillitis | emm64.3 | 1008 | 14 | 25 | 3 | 26 | 71 | 45 | 103 | 123 | 43 | 3 | 76 | 30 |
| Bra045 | Brazil | South America | 2000 | ARF | emm87.0 | 62 | 67 | 15 | 10 | 1 | 72 | 58 | 87 | 95 | 1 | 2 | 52 | 74 |
| Bra033 | Brazil | South America | 2007 | scarlet fever | emm4.0 | 1009 | 4 | 8 | 4 | 7 | 51 | 9 | 8 | 9 | 7 | 2 | 8 | 8 |
| Bra021 | Brazil | South America | 2013 | scarlet fever | emm22.0 | 46 | 7 | 5 | 5 | 5 | 4 | 5 | 5 | 6 | 4 | 3 | 5 | 5 |
| Bra009 | Brazil | South America | 2004 | scarlet fever | emm1.0 | 28 | 1 | 1 | 1 | 1 | 1 | 1 | 1 | 1 | 1 | 1 | 1 | 1 |
| Bra044 | Brazil | South America | 2011 | pharyngitis and/or tonsillitis | emm27.0 | 308 | 10 | 85 | 65 | 56 | 1 | 69 | 104 | 18 | 9 | 2 | 12 | 13 |
| Bra032 | Brazil | South America | 2012 | pharyngitis and/or tonsillitis | emm1.0 | 28 | 1 | 1 | 1 | 1 | 1 | 1 | 1 | 1 | 44 | 1 | 1 | 1 |
| Bra007 | Brazil | South America | 2011 | scarlet fever | emm101.1 | 11 | 11 | 86 | 10 | 20 | 15 | 16 | 17 | 13 | 1 | 2 | 16 | 31 |
| Bra043 | Brazil | South America | 2003 | skin/soft tissue infection, NOS | emm117.1 | 986 | 14 | 83 | 3 | 3 | 92 | 7 | 25 | 121 | 36 | 4 | 74 | 25 |
| Bra031 | Brazil | South America | 2013 | invasive, NOS | emm8.0 | 59 | 14 | 13 | 3 | 41 | 50 | 13 | 3 | 110 | 7 | 2 | 49 | 4 |
| Bra019 | Brazil | South America | 2007 | scarlet fever | emm1.0 | 28 | 1 | 1 | 1 | 1 | 1 | 1 | 1 | 1 | 44 | 1 | 1 | 1 |
| Bra008 | Brazil | South America | 2012 | ND | emm22.0 | 46 | 7 | 5 | 5 | 5 | 4 | 5 | 5 | 6 | 4 | 3 | 5 | 5 |
| Bra042 | Brazil | South America | 2007 | ARF | emm75.0 | 150 | 25 | 26 | 20 | 24 | 29 | 29 | 33 | 112 | 15 | 3 | 12 | 33 |
| Bra030 | Brazil | South America | 2005 | pharyngitis and/or tonsillitis | emm2.0 | 55 | 32 | 26 | 28 | 24 | 29 | 13 | 43 | 48 | 15 | 3 | 12 | 42 |
| Bra018 | Brazil | South America | 2002 | pharyngitis and/or tonsillitis | emm185.0 | 1010 | 18 | 18 | 66 | 1 | 10 | 7 | 105 | 124 | 1 | 1 | 77 | 86 |

| | | | | | | | | | | | | | | | | | | |
|---|---|---|---|---|---|---|---|---|---|---|---|---|---|---|---|---|---|---|
| Bra041 | Brazil | South America | 2014 | skin/soft tissue infection, NOS | emm53.0 | 11 | 11 | 5 | 10 | 8 | 15 | 54 | 17 | 84 | 1 | 2 | 6 | 20 |
| Bra029 | Brazil | South America | 2013 | invasive, NOS | emm34.0 | 14 | 28 | 87 | 10 | 1 | 6 | 63 | 78 | 126 | 1 | 2 | 6 | 31 |
| 5448 | Canada | North America | 1994 | necrotizing fasciitis | emm1.0 | 28 | 1 | 1 | 1 | 1 | 1 | 1 | 1 | 1 | 1 | 1 | 1 | 1 |
| MGAS15252 | Canada | North America | 2008 | ND | emm59.0 | 172 | 72 | 84 | 24 | 2 | 93 | 68 | 107 | 122 | 18 | 3 | 75 | 85 |
| NGAS322 | Canada | North America | ND | invasive, NOS | emm114.0 | 188 | 14 | 18 | 67 | 16 | 21 | 61 | 108 | 26 | 46 | 3 | 78 | 87 |
| NGAS596 | Canada | North America | ND | invasive, NOS | emm82.0 | 334 | 73 | 58 | 46 | 1 | 6 | 18 | 69 | 77 | 1 | 1 | 1 | 3 |
| NGAS638 | Canada | North America | ND | invasive, NOS | emm101.0 | 182 | 15 | 31 | 10 | 20 | 15 | 31 | 38 | 41 | 1 | 21 | 32 | 37 |
| NGAS743 | Canada | North America | ND | ND | emm87.0 | 62 | 67 | 15 | 10 | 1 | 72 | 58 | 87 | 95 | 1 | 2 | 52 | 74 |
| NGAS005 | Canada | North America | 2012 | ND | emm90.2 | 184 | 14 | 88 | 68 | 57 | 94 | 70 | 109 | 127 | 1 | 2 | 79 | 88 |
| NGAS008 | Canada | North America | 2012 | ND | emm122.2 | 1080 | 14 | 37 | 20 | 24 | 64 | 57 | 110 | 128 | 15 | 3 | 18 | 89 |
| NGAS015 | Canada | North America | 2012 | ND | emm63.3 | 297 | 24 | 19 | 3 | 23 | 28 | 7 | 32 | [S] | 17 | 11 | 26 | 32 |
| NGAS025 | Canada | North America | 2012 | ND | emm89.0 | 101 | 2 | 2 | 2 | 58 | 2 | 2 | 2 | 2 | 2 | 2 | 2 | 2 |
| NGAS026 | Canada | North America | 2012 | ND | emm89.0 | 101 | 2 | 2 | 2 | 2 | 2 | 2 | 2 | 2 | 2 | 2 | 2 | 2 |
| NGAS035 | Canada | North America | 2012 | ND | emm1.0 | 1064 | 1 | 1 | 1 | 1 | 1 | 1 | 1 | 1 | 1 | 1 | 1 | 1 |
| NGAS056 | Canada | North America | 2012 | ND | emm1.0 | 28 | 1 | 1 | 1 | 1 | 95 | 1 | 1 | 1 | 1 | 1 | 1 | 1 |
| NGAS057 | Canada | North America | 2012 | ND | emm88.3 | 971 | 14 | 89 | 69 | 35 | 96 | 2 | 67 | 74 | 1 | 2 | 6 | 31 |
| NGAS061 | Canada | North America | 2012 | ND | emm1.0 | 28 | 1 | 1 | 1 | 1 | 1 | 1 | 1 | 1 | 1 | 1 | 1 | 1 |
| NGAS070 | Canada | North America | 2012 | ND | emm89.0 | 101 | 2 | 2 | 2 | 2 | 97 | 2 | 2 | 2 | 2 | 2 | 2 | 2 |
| NGAS071 | Canada | North America | 2012 | ND | emm87.0 | 62 | 67 | 15 | 10 | 1 | 72 | 58 | 87 | 95 | 1 | 2 | 52 | 74 |
| NGAS075 | Canada | North America | 2012 | ND | emm9.0 | 75 | 34 | 77 | 60 | 32 | 2 | 1 | 95 | 113 | 5 | 3 | 11 | 45 |
| NGAS076 | Canada | North America | 2012 | ND | emm1.0 | 28 | 1 | 1 | 1 | 1 | 1 | 1 | 1 | 1 | 1 | 1 | 1 | 1 |
| NGAS078 | Canada | North America | 2012 | ND | emm1.0 | 28 | 1 | 1 | 1 | 1 | 1 | 1 | 1 | 1 | 1 | 1 | 1 | 1 |
| NGAS083 | Canada | North America | 2012 | ND | emm1.0 | 28 | 1 | 1 | 1 | 1 | 1 | 1 | 1 | 1 | 1 | 1 | 1 | 1 |
| NGAS084 | Canada | North America | 2012 | ND | emm28.0 | 52 | 3 | 3 | 3 | 3 | 2 | 3 | 3 | 4 | 1 | 1 | 3 | 3 |
| NGAS088 | Canada | North America | 2012 | ND | emm89.0 | 101 | 2 | 2 | 2 | 2 | 97 | 2 | 2 | 2 | 2 | 2 | 2 | 2 |
| NGAS094 | Canada | North America | 2012 | ND | emm3.1 | 15 | 15 | 15 | 10 | 1 | 10 | 32 | 111 | 129 | 1 | 2 | 6 | 31 |
| NGAS096 | Canada | North America | 2012 | ND | emm12.0 | 36 | 14 | 3 | 3 | 3 | 2 | 3 | 52 | 59 | 24 | 12 | 38 | 3 |
| NGAS100 | Canada | North America | 2012 | ND | emm3.1 | 15 | 15 | 15 | 10 | 1 | 10 | 32 | 111 | 129 | 1 | 2 | 6 | 31 |
| NGAS104 | Canada | North America | 2012 | ND | emm3.53 | 406 | 15 | 15 | 10 | 1 | 10 | 32 | 111 | 129 | 1 | 2 | 6 | 31 |
| NGAS113 | Canada | North America | 2012 | ND | emm122.0 | 200 | 11 | 13 | 3 | 41 | 50 | 13 | 112 | 42 | 31 | 1 | 51 | 4 |
| NGAS123 | Canada | North America | 2012 | ND | emm6.79 | 382 | 65 | 80 | 62 | 55 | 85 | 62 | 97 | 130 | 11 | 20 | 69 | 82 |
| NGAS128 | Canada | North America | 2012 | ND | emm14.3 | 84 | 11 | 5 | 10 | 8 | 15 | 16 | 113 | 13 | 1 | 2 | 6 | 20 |
| NGAS130 | Canada | North America | 2012 | ND | emm191.0 | 208 | 11 | 90 | 3 | 59 | 50 | 13 | 4 | 42 | 31 | 1 | 4 | 90 |
| NGAS145 | Canada | North America | 2012 | ND | emm12.0 | 36 | 14 | 3 | 3 | 3 | 2 | 3 | 52 | 59 | 24 | 12 | 38 | 3 |
| NGAS148 | Canada | North America | 2012 | ND | ND | 99 | 14 | 91 | 70 | 1 | 17 | 71 | 114 | 131 | 47 | 22 | 80 | 91 |
| NGAS151 | Canada | North America | 2012 | ND | emm6.0 | 382 | 65 | 80 | 62 | 55 | 85 | 62 | 97 | 130 | 11 | 20 | 69 | 82 |
| NGAS162 | Canada | North America | 2012 | ND | emm6.0 | 382 | 65 | 80 | 62 | 55 | 85 | 62 | 97 | 130 | 11 | 20 | 69 | 82 |
| NGAS169 | Canada | North America | 2012 | ND | emm77.0 | 63 | 23 | 24 | 19 | 1 | 26 | 28 | 30 | 33 | 8 | 3 | 24 | 29 |
| NGAS185 | Canada | North America | 2012 | ND | emm12.8 | 36 | 74 | 3 | 3 | 3 | 2 | 3 | 52 | 59 | 24 | 12 | 38 | 3 |
| NGAS197 | Canada | North America | 2012 | ND | emm89.0 | 101 | 2 | 2 | 2 | 2 | 97 | 2 | 2 | 2 | 2 | 2 | 2 | 2 |
| NGAS202 | Canada | North America | 2012 | ND | emm1.0 | 28 | 1 | 1 | 1 | 1 | 1 | 1 | 1 | 1 | 1 | 1 | 1 | 1 |
| NGAS209 | Canada | North America | 2012 | ND | emm18.0 | ND | 75 | 92 | 61 | 5 | 98 | 72 | 115 | 132 | 47 | 2 | 69 | 5 |
| NGAS226 | Canada | North America | 2012 | ND | emm2.0 | 55 | 32 | 26 | 28 | 24 | 29 | 13 | 43 | 48 | 15 | 3 | 12 | 42 |
| NGAS227 | Canada | North America | 2012 | ND | emm102.2 | 895 | 14 | 93 | 71 | 60 | 99 | 13 | 116 | 133 | 13 | 2 | 81 | 92 |
| NGAS228 | Canada | North America | 2012 | ND | emm4.0 | 39 | 4 | 8 | 4 | 7 | 51 | 9 | 8 | 9 | 7 | 2 | 8 | 8 |
| NGAS233 | Canada | North America | 2012 | ND | emm89.0 | 101 | 2 | 2 | 2 | 2 | 2 | 2 | 2 | 2 | 2 | 2 | 2 | 2 |
| NGAS240 | Canada | North America | 2012 | ND | emm82.0 | 334 | 73 | 58 | 46 | 1 | 6 | 18 | 69 | 77 | 1 | 1 | 1 | 3 |
| NGAS241 | Canada | North America | 2012 | ND | emm82.0 | 334 | 73 | 58 | 46 | 1 | 6 | 18 | 69 | 77 | 1 | 1 | 1 | 3 |

| | | | | | | | | | | | | | | | | | | |
|---|---|---|---|---|---|---|---|---|---|---|---|---|---|---|---|---|---|---|
| NGAS245 | Canada | North America | 2012 | ND | emm58.0 | 176 | 14 | 94 | 10 | 8 | 13 | 13 | 15 | 47 | 10 | 2 | 34 | 31 |
| NGAS253 | Canada | North America | 2012 | ND | emm83.1 | 853 | 11 | 5 | 10 | 8 | 15 | 73 | 97 | 115 | 11 | 2 | 82 | 83 |
| NGAS255 | Canada | North America | 2012 | ND | emm1.0 | 28 | 1 | 1 | 1 | 1 | 1 | 1 | 1 | 1 | 1 | 1 | 1 | 1 |
| NGAS260 | Canada | North America | 2012 | ND | emm12.8 | 36 | 74 | 3 | 3 | 3 | 2 | 3 | 52 | 59 | 24 | 12 | 38 | 3 |
| NGAS266 | Canada | North America | 2012 | ND | emm3.1 | 15 | 15 | 15 | 10 | 1 | 10 | 32 | 111 | 129 | 1 | 2 | 6 | 31 |
| NGAS282 | Canada | North America | 2012 | ND | emm89.0 | 101 | 2 | 2 | 2 | 2 | 2 | 2 | 2 | 2 | 2 | 2 | 2 | 2 |
| NGAS287 | Canada | North America | 2012 | ND | emm6.0 | 382 | 65 | 80 | 62 | 55 | 85 | 62 | 97 | 130 | 11 | 20 | 69 | 82 |
| NGAS289 | Canada | North America | 2012 | ND | emm6.0 | 382 | 65 | 80 | 62 | 55 | 85 | 62 | 97 | 130 | 11 | 20 | 69 | 82 |
| NGAS290 | Canada | North America | 2012 | ND | emm87.0 | 62 | 67 | 15 | 10 | 1 | 72 | 58 | 87 | 95 | 1 | 2 | 52 | 74 |
| NGAS291 | Canada | North America | 2012 | ND | emm87.0 | 62 | 67 | 15 | 10 | 1 | 72 | 58 | 87 | 95 | 1 | 2 | 52 | 74 |
| NGAS292 | Canada | North America | 2012 | ND | emm87.0 | 62 | 67 | 15 | 10 | 1 | 72 | 58 | 87 | 95 | 1 | 2 | 52 | 74 |
| NGAS297 | Canada | North America | 2012 | ND | emm1.0 | 28 | 1 | 1 | 1 | 1 | 1 | 1 | 1 | 134 | 1 | 1 | 1 | 1 |
| NGAS298 | Canada | North America | 2011 | invasive, NOS | emm87.0 | 62 | 67 | 15 | 10 | 1 | 72 | 58 | 87 | 95 | 1 | 2 | 52 | 74 |
| NGAS299 | Canada | North America | 2011 | invasive, NOS | emm87.0 | 62 | 67 | 15 | 10 | 1 | 72 | 58 | 87 | 95 | 1 | 2 | 52 | 74 |
| NGAS300 | Canada | North America | 2011 | invasive, NOS | emm83.1 | 5 | 11 | 5 | 10 | 8 | 15 | 65 | 97 | 115 | 11 | 2 | 69 | 83 |
| NGAS302 | Canada | North America | 2011 | invasive, NOS | emm28.0 | 52 | 3 | 95 | 3 | 3 | 2 | 3 | 117 | 4 | 1 | 1 | 3 | 3 |
| NGAS304 | Canada | North America | 2011 | invasive, NOS | emm28.0 | 52 | 3 | 3 | 3 | 3 | 2 | 3 | 3 | 4 | 1 | 1 | 3 | 3 |
| NGAS305 | Canada | North America | 2011 | invasive, NOS | emm1.0 | 28 | 1 | 1 | 1 | 1 | 1 | 1 | 1 | 1 | 1 | 1 | 1 | 1 |
| NGAS308 | Canada | North America | 2011 | invasive, NOS | emm18.0 | ND | 75 | 92 | 61 | 5 | 98 | 72 | 115 | 132 | 47 | 2 | 69 | 5 |
| NGAS312 | Canada | North America | 2012 | ND | emm80.0 | 538 | 11 | 78 | 61 | 54 | 84 | 9 | 96 | 114 | 40 | 3 | 68 | 3 |
| NGAS320 | Canada | North America | 2011 | invasive, NOS | emm83.1 | 853 | 11 | 5 | 10 | 8 | 15 | 73 | 97 | 115 | 11 | 2 | 82 | 83 |
| NGAS323 | Canada | North America | 2011 | ND | emm82.0 | 334 | 73 | 58 | 46 | 1 | 6 | 18 | 69 | 77 | 1 | 1 | 1 | 3 |
| NGAS325 | Canada | North America | 2011 | invasive, NOS | emm22.0 | 1069 | 76 | 5 | 5 | 5 | 4 | 5 | 5 | 6 | 4 | 3 | 5 | 5 |
| NGAS328 | Canada | North America | 2011 | ND | emm82.0 | 334 | 73 | 58 | 46 | 1 | 6 | 18 | 69 | 77 | 1 | 1 | 1 | 3 |
| NGAS332 | Canada | North America | 2011 | ND | emm82.0 | 334 | 73 | 58 | 46 | 1 | 6 | 18 | 69 | 77 | 1 | 1 | 1 | 3 |
| NGAS336 | Canada | North America | 2011 | invasive, NOS | emm1.0 | 28 | 1 | 1 | 1 | 1 | 1 | 1 | 1 | 134 | 1 | 1 | 1 | 1 |
| NGAS338 | Canada | North America | 2011 | invasive, NOS | emm82.0 | 334 | 73 | 58 | 46 | 1 | 6 | 18 | 69 | 77 | 1 | 1 | 1 | 3 |
| NGAS339 | Canada | North America | 2011 | ND | emm59.0 | 172 | 72 | 84 | 24 | 2 | 93 | 68 | 107 | 122 | 18 | 3 | 75 | 85 |
| NGAS340 | Canada | North America | 2011 | ND | emm118.0 | 167 | 11 | 32 | 52 | 1 | 67 | 13 | 118 | 135 | 1 | 23 | 83 | 93 |
| NGAS344 | Canada | North America | 2011 | invasive, NOS | emm75.0 | 49 | 25 | 26 | 20 | 24 | 29 | 74 | 33 | 136 | 15 | 3 | 12 | 33 |
| NGAS346 | Canada | North America | 2011 | ND | emm41.2 | 579 | 11 | 5 | 10 | 51 | 15 | 16 | 77 | 99 | 1 | 2 | 6 | 20 |
| NGAS347 | Canada | North America | 2011 | ND | emm82.0 | 334 | 73 | 58 | 46 | 1 | 6 | 18 | 69 | 77 | 1 | 1 | 1 | 3 |
| NGAS358 | Canada | North America | 2012 | ND | emm169.3 | 53 | 70 | 4 | 3 | 18 | 19 | 19 | 101 | 118 | 42 | 2 | 72 | 3 |
| NGAS364 | Canada | North America | 2012 | ND | emm77.0 | 63 | 23 | 24 | 57 | 1 | 26 | 28 | 30 | 33 | 8 | 3 | 24 | 29 |
| NGAS367 | Canada | North America | 2012 | ND | emm169.3 | 53 | 70 | 4 | 3 | 18 | 19 | 19 | 101 | 118 | 42 | 2 | 72 | 3 |
| NGAS372 | Canada | North America | 2012 | ND | emm92.0 | 82 | 17 | 4 | 3 | 18 | 19 | 19 | 22 | 24 | 13 | 2 | 20 | 22 |
| NGAS401 | Canada | North America | 2012 | ND | emm102.2 | 895 | 14 | 93 | 71 | 60 | 99 | 13 | 116 | 133 | 13 | 2 | 81 | 92 |
| NGAS408 | Canada | North America | 2012 | ND | emm1.0 | 28 | 1 | 1 | 1 | 1 | 1 | 1 | 1 | 1 | 1 | 1 | 1 | 1 |
| NGAS413 | Canada | North America | 2012 | ND | emm1.0 | 28 | 1 | 1 | 1 | 1 | 1 | 1 | 1 | 1 | 1 | 1 | 1 | 1 |
| NGAS419 | Canada | North America | 2012 | ND | emm3.1 | 15 | 15 | 15 | 10 | 1 | 10 | 32 | 111 | 129 | 1 | 2 | 6 | 31 |
| NGAS425 | Canada | North America | 2012 | ND | emm1.0 | 28 | 1 | 1 | 1 | 1 | 1 | 1 | 1 | 1 | 1 | 1 | 1 | 1 |
| NGAS426 | Canada | North America | 2012 | ND | emm3.1 | 15 | 15 | 15 | 10 | 1 | 10 | 32 | 111 | 129 | 1 | 2 | 6 | 31 |
| NGAS428 | Canada | North America | 2012 | ND | emm5.14 | 99 | 14 | 91 | 70 | 1 | 17 | 71 | 114 | 131 | 47 | 22 | 80 | 91 |
| NGAS438 | Canada | North America | 2012 | ND | emm73.0 | 331 | 14 | 96 | 3 | 41 | 50 | 13 | 3 | 137 | 48 | 2 | 49 | 94 |
| NGAS440 | Canada | North America | 2012 | ND | emm9.2 | 12 | 11 | 5 | 10 | 20 | 15 | 16 | 17 | 138 | 1 | 2 | 16 | 31 |
| NGAS447 | Canada | North America | 2012 | ND | emm12.0 | 36 | 14 | 3 | 3 | 3 | 2 | 3 | 52 | 59 | 24 | 12 | 38 | 3 |
| NGAS450 | Canada | North America | 2012 | ND | emm1.0 | 28 | 1 | 1 | 1 | 1 | 1 | 1 | 1 | 1 | 1 | 1 | 1 | 1 |
| NGAS465 | Canada | North America | 2012 | ND | emm29.1 | 12 | 11 | 5 | 10 | 20 | 15 | 16 | 17 | 138 | 1 | 2 | 16 | 31 |

| | | | | | | | | | | | | | | | | | | |
|---|---|---|---|---|---|---|---|---|---|---|---|---|---|---|---|---|---|---|
| NGAS473 | Canada | North America | 2012 | ND | emm82.0 | 36 | 14 | 3 | 3 | 3 | 2 | 3 | 52 | 59 | 24 | 12 | 38 | 3 |
| NGAS474 | Canada | North America | 2012 | ND | emm87.0 | 62 | 67 | 15 | 10 | 1 | 72 | 58 | 87 | 95 | 1 | 2 | 52 | 74 |
| NGAS495 | Canada | North America | 2012 | ND | emm82.0 | 334 | 73 | 58 | 46 | 1 | 6 | 18 | 69 | 77 | 1 | 1 | 1 | 3 |
| NGAS500 | Canada | North America | 2012 | ND | emm73.0 | 331 | 14 | 96 | 3 | 41 | 50 | 13 | 3 | 137 | 48 | 2 | 49 | 60 |
| NGAS501 | Canada | North America | 2012 | ND | emm82.0 | 334 | 73 | 58 | 46 | 1 | 6 | 18 | 69 | 77 | 1 | 1 | 1 | 3 |
| NGAS508 | Canada | North America | 2012 | ND | emm87.0 | 62 | 67 | 15 | 10 | 1 | 72 | 58 | 87 | 95 | 1 | 2 | 52 | 74 |
| NGAS510 | Canada | North America | 2012 | ND | emm9.2 | 891 | 34 | 77 | 60 | 32 | 2 | 1 | 95 | 113 | 5 | 3 | 84 | 45 |
| NGAS516 | Canada | North America | 2012 | ND | emm89.0 | 101 | 2 | 2 | 2 | 2 | 97 | 2 | 2 | 2 | 2 | 2 | 2 | 2 |
| NGAS520 | Canada | North America | 2012 | ND | emm83.1 | 5 | 11 | 5 | 10 | 8 | 15 | 65 | 97 | 115 | 11 | 2 | 69 | 83 |
| NGAS532 | Canada | North America | 2012 | ND | emm11.0 | ND | 78 | 97 | 40 | 61 | 100 | 20 | 120 | 139 | 8 | 3 | 85 | 95 |
| NGAS535 | Canada | North America | 2012 | ND | ND | 1081 | 14 | 49 | 33 | 32 | 2 | 1 | 121 | 140 | 5 | 4 | 86 | 96 |
| NGAS556 | Canada | North America | 2012 | ND | emm87.0 | 62 | 67 | 15 | 10 | 1 | 72 | 58 | 87 | 95 | 1 | 2 | 52 | 74 |
| NGAS565 | Canada | North America | 2012 | ND | emm87.0 | 62 | 67 | 15 | 10 | 1 | 72 | 58 | 87 | 95 | 1 | 2 | 52 | 74 |
| NGAS592 | Canada | North America | 2012 | ND | emm82.0 | 334 | 73 | 58 | 46 | 1 | 6 | 18 | 69 | 77 | 1 | 1 | 1 | 3 |
| NGAS594 | Canada | North America | 2012 | ND | emm82.0 | 334 | 73 | 58 | 46 | 1 | 6 | 18 | 69 | 77 | 1 | 1 | 1 | 3 |
| NGAS595 | Canada | North America | 2012 | ND | emm89.0 | 101 | 2 | 2 | 2 | 2 | 2 | 2 | 2 | 2 | 2 | 2 | 2 | 2 |
| NGAS597 | Canada | North America | 2012 | invasive, NOS | emm82.0 | 334 | 73 | 58 | 46 | 1 | 6 | 18 | 69 | 77 | 1 | 1 | 1 | 3 |
| NGAS599 | Canada | North America | 2012 | ND | emm114.1 | 188 | 14 | 18 | 67 | 16 | 21 | 61 | 108 | 26 | 46 | 3 | 78 | 87 |
| NGAS601 | Canada | North America | 2012 | ND | emm82.0 | 334 | 73 | 58 | 46 | 1 | 6 | 18 | 69 | 77 | 1 | 1 | 1 | 3 |
| NGAS603 | Canada | North America | 2012 | invasive, NOS | emm6.0 | 382 | 65 | 80 | 62 | 55 | 85 | 62 | 97 | 130 | 11 | 20 | 69 | 82 |
| NGAS604 | Canada | North America | 2012 | invasive, NOS | emm75.0 | 49 | 25 | 26 | 20 | 24 | 29 | 74 | 33 | 136 | 15 | 3 | 12 | 33 |
| NGAS605 | Canada | North America | 2012 | ND | emm82.0 | 334 | 73 | 58 | 46 | 1 | 6 | 18 | 69 | 77 | 1 | 1 | 1 | 3 |
| NGAS606 | Canada | North America | 2012 | invasive, NOS | emm80.0 | 538 | 11 | 78 | 61 | 54 | 84 | 9 | 96 | 114 | 40 | 3 | 68 | 3 |
| NGAS608 | Canada | North America | 2012 | ND | emm82.0 | 334 | 73 | 58 | 46 | 1 | 6 | 18 | 69 | 77 | 1 | 1 | 1 | 3 |
| NGAS609 | Canada | North America | 2012 | invasive, NOS | emm82.0 | 334 | 73 | 98 | 46 | 1 | 6 | 18 | 69 | 77 | 1 | 1 | 1 | 3 |
| NGAS610 | Canada | North America | 2012 | invasive, NOS | emm87.0 | 62 | 67 | 15 | 10 | 1 | 72 | 58 | 87 | 95 | 1 | 2 | 52 | 74 |
| NGAS612 | Canada | North America | 2012 | ND | emm87.0 | 62 | 67 | 15 | 10 | 1 | 72 | 58 | 87 | 95 | 1 | 2 | 52 | 74 |
| NGAS613 | Canada | North America | 2012 | invasive, NOS | emm82.0 | 334 | 73 | 58 | 46 | 1 | 6 | 18 | 69 | 77 | 1 | 1 | 1 | 3 |
| NGAS615 | Canada | North America | 2012 | ND | emm82.0 | 334 | 73 | 58 | 46 | 1 | 6 | 18 | 69 | 77 | 1 | 1 | 1 | 3 |
| NGAS616 | Canada | North America | 2012 | ND | emm22.0 | 1069 | 76 | 5 | 5 | 5 | 4 | 5 | 5 | 6 | 4 | 3 | 5 | 5 |
| NGAS618 | Canada | North America | 2012 | invasive, NOS | emm6.0 | 382 | 65 | 80 | 62 | 55 | 85 | 62 | 97 | 130 | 11 | 20 | 69 | 82 |
| NGAS623 | Canada | North America | 2012 | ND | emm1.0 | 28 | 1 | 1 | 1 | 1 | 1 | 1 | 1 | 1 | 1 | 1 | 1 | 1 |
| NGAS629 | Canada | North America | 2012 | ND | emm83.1 | 5 | 11 | 5 | 10 | 8 | 15 | 65 | 97 | 115 | 11 | 2 | 69 | 83 |
| NGAS630 | Canada | North America | 2012 | ND | emm87.0 | 62 | 67 | 15 | 10 | 1 | 72 | 58 | 87 | 95 | 1 | 2 | 52 | 74 |
| NGAS633 | Canada | North America | 2012 | ND | emm83.1 | 853 | 11 | 5 | 10 | 8 | 15 | 73 | 97 | 115 | 11 | 2 | 82 | 83 |
| NGAS634 | Canada | North America | 2012 | invasive, NOS | emm87.0 | 62 | 67 | 15 | 10 | 1 | 72 | 58 | 87 | 95 | 1 | 2 | 52 | 74 |
| NGAS639 | Canada | North America | 2012 | invasive, NOS | emm59.0 | 172 | 72 | 84 | 24 | 2 | 93 | 68 | 107 | 122 | 18 | 3 | 75 | 85 |
| NGAS640 | Canada | North America | 2012 | ND | emm11.0 | ND | 1 | 6 | 40 | 61 | 20 | 20 | 122 | 141 | 8 | 3 | 18 | 4 |
| NGAS641 | Canada | North America | 2012 | ND | emm114.0 | 188 | 14 | 18 | 67 | 16 | 21 | 61 | 108 | 26 | 46 | 3 | 78 | 87 |
| NGAS643 | Canada | North America | 2012 | ND | emm83.1 | 853 | 11 | 5 | 10 | 8 | 15 | 73 | 97 | 115 | 11 | 2 | 82 | 83 |
| NGAS650 | Canada | North America | 2012 | ND | emm3.6 | 315 | 15 | 15 | 10 | 1 | 10 | 32 | 111 | 129 | 1 | 2 | 6 | 31 |
| NGAS653 | Canada | North America | 2012 | ND | emm28.0 | 458 | 3 | 3 | 3 | 3 | 2 | 3 | 3 | 4 | 1 | 1 | 3 | 3 |
| NGAS655 | Canada | North America | 2012 | ND | emm83.1 | 853 | 11 | 5 | 10 | 8 | 15 | 73 | 97 | 115 | 11 | 2 | 82 | 83 |
| NGAS657 | Canada | North America | 2012 | ND | emm1.0 | 28 | 1 | 1 | 1 | 1 | 1 | 1 | 1 | 134 | 1 | 1 | 1 | 1 |
| NGAS660 | Canada | North America | 2012 | ND | emm89.0 | 101 | 2 | 2 | 2 | 2 | 97 | 2 | 2 | 2 | 2 | 2 | 2 | 2 |
| NGAS662 | Canada | North America | 2012 | ND | emm87.0 | 62 | 67 | 15 | 10 | 1 | 72 | 58 | 87 | 95 | 1 | 2 | 52 | 74 |
| NGAS663 | Canada | North America | 2012 | ND | emm89.0 | 101 | 2 | 2 | 2 | 2 | 97 | 2 | 2 | 2 | 2 | 2 | 2 | 2 |
| NGAS664 | Canada | North America | 2012 | ND | emm74.0 | 120 | 28 | 13 | 3 | 41 | 50 | 53 | 123 | 82 | 11 | 4 | 59 | 70 |

| | | | | | | | | | | | | | | | | | |
|---|---|---|---|---|---|---|---|---|---|---|---|---|---|---|---|---|---|
| NGAS665 | Canada | North America | 2012 | ND | emm87.0 | 62 | 67 | 15 | 10 | 1 | 72 | 58 | 87 | 95 | 1 | 2 | 52 | 74 |
| NGAS668 | Canada | North America | 2012 | ND | emm87.0 | 62 | 67 | 15 | 10 | 1 | 72 | 58 | 87 | 95 | 1 | 2 | 52 | 74 |
| NGAS669 | Canada | North America | 2012 | ND | emm87.0 | 62 | 67 | 15 | 10 | 1 | 72 | 58 | 87 | 95 | 1 | 2 | 52 | 74 |
| NGAS670 | Canada | North America | 2012 | ND | emm4.0 | 39 | 4 | 8 | 4 | 7 | 51 | 9 | 63 | 9 | 7 | 2 | 8 | 8 |
| NGAS671 | Canada | North America | 2012 | ND | emm2.0 | 55 | 32 | 26 | 28 | 24 | 29 | 13 | 43 | 48 | 15 | 3 | 12 | 42 |
| NGAS673 | Canada | North America | 2012 | ND | emm89.0 | 101 | 2 | 2 | 2 | 2 | 97 | 2 | 2 | 2 | 2 | 2 | 2 | 2 |
| NGAS675 | Canada | North America | 2012 | ND | emm4.0 | 39 | 4 | 8 | 4 | 7 | 51 | 9 | 63 | 9 | 7 | 2 | 8 | 8 |
| NGAS717 | Canada | North America | 2012 | ND | ND | 82 | 17 | 4 | 3 | 18 | 19 | 19 | 22 | 24 | 13 | 2 | 20 | 22 |
| NGAS719 | Canada | North America | 2012 | ND | emm90.5 | 144 | 79 | 20 | 14 | 32 | 2 | 23 | 25 | 57 | 18 | 3 | 22 | 25 |
| NGAS726 | Canada | North America | 2012 | ND | emm68.3 | 894 | 25 | 26 | 20 | 24 | 29 | 13 | 124 | 48 | 49 | 3 | 12 | 42 |
| NGAS729 | Canada | North America | 2012 | ND | emm94.0 | 89 | 8 | 4 | 4 | 4 | 3 | 13 | 4 | 42 | 21 | 24 | 4 | 4 |
| NGAS732 | Canada | North America | 2012 | ND | emm94.0 | 89 | 8 | 4 | 4 | 4 | 3 | 13 | 4 | 42 | 21 | 24 | 4 | 4 |
| NGAS737 | Canada | North America | 2012 | ND | emm4.0 | 39 | 4 | 8 | 4 | 7 | 51 | 9 | 8 | 9 | 7 | 2 | 8 | 8 |
| NGAS739 | Canada | North America | 2013 | invasive, NOS | emm87.0 | 62 | 67 | 15 | 10 | 1 | 72 | 58 | 87 | 95 | 1 | 2 | 52 | 74 |
| NGAS746 | Canada | North America | 2013 | invasive, NOS | emm4.0 | 39 | 4 | 8 | 4 | 7 | 51 | 9 | 8 | 9 | 7 | 2 | 8 | 8 |
| NGAS747 | Canada | North America | 2013 | invasive, NOS | emm87.0 | 62 | 67 | 15 | 10 | 1 | 72 | 58 | 87 | 95 | 1 | 2 | 52 | 74 |
| NGAS749 | Canada | North America | 2013 | invasive, NOS | emm12.0 | 36 | 14 | 3 | 3 | 3 | 2 | 3 | 52 | 59 | 24 | 12 | 38 | 3 |
| NGAS750 | Canada | North America | 2013 | invasive, NOS | emm1.0 | 28 | 1 | 1 | 1 | 1 | 1 | 1 | 1 | 134 | 1 | 1 | 1 | 1 |
| NGAS751 | Canada | North America | 2012 | ND | emm115.0 | 135 | 77 | 11 | 72 | 1 | 15 | 31 | 119 | 142 | 1 | 1 | 1 | 3 |
| NGAS755 | Canada | North America | 2013 | ND | emm101.0 | 182 | 15 | 31 | 10 | 20 | 15 | 31 | 38 | 41 | 1 | 21 | 32 | 37 |
| NGAS757 | Canada | North America | 2013 | ND | emm82.0 | 334 | 73 | 58 | 46 | 1 | 6 | 18 | 69 | 77 | 1 | 1 | 1 | 3 |
| NGAS758 | Canada | North America | 2013 | ND | emm4.0 | 39 | 4 | 8 | 4 | 7 | 51 | 9 | 8 | 9 | 7 | 2 | 8 | 8 |
| NGAS759 | Canada | North America | 2012 | ND | emm4.0 | 39 | 4 | 8 | 4 | 7 | 51 | 9 | 8 | 9 | 7 | 2 | 8 | 8 |
| NGAS760 | Canada | North America | 2012 | ND | emm4.0 | 39 | 4 | 8 | 4 | 7 | 51 | 9 | 63 | 9 | 7 | 2 | 8 | 97 |
| TJ11-001 | China | Asia | 2011 | scarlet fever | emm12.0 | 36 | 14 | 3 | 3 | 3 | 2 | 3 | 52 | 59 | 24 | 12 | 38 | 3 |
| AP1 | Czech Republic | Europe | ND | bacteremia | emm1.0 | 28 | 1 | 1 | 1 | 1 | 1 | 1 | 1 | 1 | 1 | 1 | 1 | 1 |
| 33022V1T1 | Fiji | Oceania | 2006 | pharyngitis and/or tonsillitis | emm82.1 | 320 | 56 | 70 | 4 | 1 | [S] | 2 | 89 | 98 | 38 | 2 | 65 | 3 |
| 33042V1T1 | Fiji | Oceania | 2006 | pharyngitis and/or tonsillitis | emm14.4 | 118 | 11 | 5 | 38 | 8 | 15 | 16 | 17 | 13 | 1 | 2 | 40 | 53 |
| 33076V1T1 | Fiji | Oceania | 2006 | pharyngitis and/or tonsillitis | emm232.1 | 1013 | 18 | 18 | 16 | 16 | 21 | 25 | 125 | 26 | 8 | 3 | 21 | 23 |
| 33087V1T1 | Fiji | Oceania | 2006 | pharyngitis and/or tonsillitis | emm58.0 | 176 | 14 | 13 | 73 | 29 | 80 | 75 | 126 | 47 | 10 | 2 | 34 | [S] |
| 33100V1T1 | Fiji | Oceania | 2006 | pharyngitis and/or tonsillitis | emm54.1 | 990 | 11 | 5 | 10 | 62 | 101 | 16 | 17 | 143 | 1 | 2 | 16 | 31 |
| 33101V1T1 | Fiji | Oceania | 2006 | pharyngitis and/or tonsillitis | emm33.0 | 1060 | 11 | 11 | 7 | 10 | 35 | 36 | 85 | 144 | 1 | 2 | 6 | 20 |
| 33112V1T1 | Fiji | Oceania | 2006 | pharyngitis and/or tonsillitis | emm42.0 | 1024 | 37 | 32 | 37 | 35 | 43 | 43 | 44 | [S] | 2 | 2 | 39 | 99 |
| 33124V1T1 | Fiji | Oceania | 2006 | pharyngitis and/or tonsillitis | emm65.4 | 129 | 11 | 66 | 40 | 35 | 102 | 76 | 44 | 145 | 1 | 2 | 87 | 31 |
| 33129V1T1 | Fiji | Oceania | 2006 | pharyngitis and/or tonsillitis | emm75.1 | 1078 | 27 | 28 | 22 | 26 | 31 | 77 | 35 | [S] | 18 | 3 | 28 | 30 |
| 33131V1T1 | Fiji | Oceania | 2006 | pharyngitis and/or tonsillitis | emm232.1 | 1013 | 18 | 18 | 16 | 16 | 21 | 25 | [S] | 26 | 8 | 3 | 21 | 23 |
| 33133V1T1 | Fiji | Oceania | 2006 | pharyngitis and/or tonsillitis | emm71.0 | 318 | 19 | 20 | 13 | 19 | 2 | 22 | 3 | 27 | 1 | 1 | 88 | 24 |
| 33140V1T1 | Fiji | Oceania | 2006 | pharyngitis and/or tonsillitis | emm86.2 | 963 | 11 | 32 | 74 | 20 | 15 | 16 | 128 | 13 | 1 | 2 | 6 | 100 |
| 33149V1T1 | Fiji | Oceania | 2006 | pharyngitis and/or tonsillitis | emm77.0 | 572 | 15 | 99 | 3 | 39 | 56 | 13 | 68 | 146 | 32 | 3 | 54 | 101 |
| 33167V1T1 | Fiji | Oceania | 2006 | pharyngitis and/or tonsillitis | emm230.0 | 10 | 16 | 100 | 75 | 16 | 17 | 7 | 20 | 22 | 1 | 2 | 18 | 3 |
| 33173V1T1 | Fiji | Oceania | 2006 | pharyngitis and/or tonsillitis | emm71.0 | 318 | 19 | 20 | 13 | 19 | 2 | 22 | 3 | 27 | 1 | 1 | 4 | 24 |
| 33181V1T1 | Fiji | Oceania | 2006 | pharyngitis and/or tonsillitis | emm137.0 | 268 | 1 | 60 | 49 | 45 | 103 | 19 | 13 | [S] | 34 | 3 | 61 | 29 |
| 33185V1T1 | Fiji | Oceania | 2006 | pharyngitis and/or tonsillitis | emm74.0 | 120 | 28 | 13 | 75 | 41 | 50 | 53 | 123 | 147 | 11 | 4 | 59 | 70 |
| 33193V1T1 | Fiji | Oceania | 2006 | pharyngitis and/or tonsillitis | emm25.1 | 192 | 14 | 101 | 24 | 2 | 33 | 13 | 13 | 40 | 36 | 3 | 31 | 36 |
| 33202V1T1 | Fiji | Oceania | 2006 | pharyngitis and/or tonsillitis | emm33.0 | 3 | 11 | 11 | 7 | 10 | 35 | 36 | 85 | 148 | 1 | 2 | 6 | 20 |
| 20004V1S1 | Fiji | Oceania | 2006 | ND | emm105.0 | 954 | 6 | 6 | 6 | 6 | 5 | 6 | 6 | 12 | 5 | 4 | 6 | 11 |
| 20028V1I1 | Fiji | Oceania | 2006 | invasive, NOS | emm81.3 | 995 | 8 | 4 | 4 | 4 | 3 | 13 | 4 | 42 | 21 | 1 | 4 | 4 |
| 20033V1I1 | Fiji | Oceania | 2006 | invasive, NOS | emm81.3 | 995 | 8 | 4 | 4 | 4 | 3 | 13 | 4 | 42 | 21 | 1 | 4 | 4 |

| | | | | | | | | | | | | | | | | | | |
|---|---|---|---|---|---|---|---|---|---|---|---|---|---|---|---|---|---|---|
| 20034V1I1 | Fiji | Oceania | 2006 | bacteremia | emm76.4 | 353 | 45 | 34 | 48 | 8 | 58 | 50 | 71 | 57 | 18 | 3 | 22 | 102 |
| 20051V1I1 | Fiji | Oceania | 2006 | invasive, NOS | emm76.4 | 353 | 45 | 34 | 48 | 8 | 58 | 50 | 71 | 57 | 18 | 3 | 22 | 102 |
| 20057V1I1 | Fiji | Oceania | 2006 | pneumonia | emm105.0 | 954 | 6 | 6 | 6 | 6 | 5 | 6 | 6 | 12 | 5 | 4 | 6 | 11 |
| 20058V1I1 | Fiji | Oceania | 2006 | invasive, NOS | emm18.22 | 535 | 14 | 102 | 2 | 2 | 2 | 78 | 67 | 75 | 8 | 3 | 89 | 2 |
| 20059V1I1 | Fiji | Oceania | 2006 | pneumonia | emm58.0 | 176 | 14 | 13 | 73 | 29 | 13 | 13 | 126 | 47 | 10 | 2 | 34 | 31 |
| 20061V1I1 | Fiji | Oceania | 2006 | invasive, NOS | emm76.4 | 353 | 45 | 34 | 48 | 8 | 58 | 50 | 71 | 57 | 18 | 3 | 22 | 102 |
| 20064V1I1 | Fiji | Oceania | 2006 | bacteremia | emm57.0 | 1025 | 80 | 9 | 8 | 1 | 32 | 79 | 129 | 149 | 8 | 2 | 23 | 103 |
| 20065V1I1 | Fiji | Oceania | 2006 | invasive, NOS | emm113.0 | 148 | 14 | 3 | 3 | 3 | 2 | 3 | 52 | 59 | 24 | 12 | 38 | 3 |
| 20006V1I1 | Fiji | Oceania | 2006 | invasive, NOS | emm11.0 | ND | 1 | 6 | 76 | 1 | 20 | 20 | 13 | 25 | 8 | 3 | 21 | 23 |
| 20066V1I1 | Fiji | Oceania | 2006 | arthritis | emm33.0 | 3 | 11 | 11 | 7 | 10 | 35 | 36 | 85 | 79 | 1 | 2 | 6 | 20 |
| 20068V1I1 | Fiji | Oceania | 2006 | bacteremia | emm123.0 | 325 | 6 | 6 | 6 | 6 | 5 | 6 | 6 | 7 | 5 | 4 | 6 | 6 |
| 20071V1I1 | Fiji | Oceania | 2006 | bacteremia | emm73.0 | 957 | 14 | 18 | 4 | 16 | 21 | 61 | 27 | 26 | 46 | 3 | 66 | 87 |
| 20077V1I1 | Fiji | Oceania | 2006 | invasive, NOS | emm60.7 | 193 | 14 | 34 | 25 | 27 | 2 | 33 | 41 | 44 | 1 | 2 | 52 | 39 |
| 20079V1I1 | Fiji | Oceania | 2006 | invasive, NOS | emm104.0 | 789 | 14 | 13 | 10 | 8 | 13 | 13 | 15 | 17 | 10 | 2 | 15 | 16 |
| 20081V1I1 | Fiji | Oceania | 2006 | invasive, NOS | emm53.0 | 11 | 11 | 5 | 10 | 8 | 15 | 54 | 17 | 84 | 1 | 2 | 6 | 20 |
| 20085V1I1 | Fiji | Oceania | 2006 | necrotizing fasciitis | emm11.0 | ND | 1 | 6 | 76 | 1 | 104 | 20 | 13 | 25 | 8 | 3 | 21 | 23 |
| 20086V1I1 | Fiji | Oceania | 2006 | invasive, NOS | emm33.0 | 3 | 11 | 11 | 7 | 10 | 35 | 36 | 85 | 79 | 1 | 2 | 6 | 20 |
| 20095V1I1 | Fiji | Oceania | 2006 | skin/soft tissue infection, NOS | emm73.0 | 957 | 14 | 18 | 4 | 16 | 21 | 61 | 27 | 26 | 46 | 3 | 66 | 87 |
| 20011V1S1 | Fiji | Oceania | 2006 | skin/soft tissue infection, NOS | emm68.0 | 993 | 81 | 103 | 33 | 63 | 105 | 7 | 130 | 150 | 34 | 3 | 90 | 104 |
| 20097V1I1 | Fiji | Oceania | 2006 | invasive, NOS | emm106.0 | 140 | 11 | 69 | 7 | 10 | 106 | 7 | 88 | 36 | 15 | 3 | 64 | 34 |
| 20098V1I1 | Fiji | Oceania | 2006 | bacteremia | emm106.0 | 140 | 11 | 69 | 7 | 10 | 106 | 7 | 88 | 36 | 15 | 3 | 64 | 34 |
| 20099V1I1 | Fiji | Oceania | 2006 | invasive, NOS | emm52.1 | 180 | 15 | 17 | 10 | 17 | 18 | 18 | 131 | 23 | 12 | 8 | 19 | 21 |
| 20109V1I1 | Fiji | Oceania | 2006 | necrotizing fasciitis | emm100.0 | 119 | 11 | 11 | 7 | 10 | 10 | 7 | 37 | 39 | 19 | 2 | 30 | 31 |
| 20110V1I1 | Fiji | Oceania | 2006 | bacteremia | emm76.4 | 353 | 45 | 34 | 48 | 8 | 58 | 50 | 71 | 57 | 18 | 3 | 22 | 102 |
| 20111V1I1 | Fiji | Oceania | 2006 | bacteremia | emm15.1 | 872 | 14 | 48 | 41 | 2 | 2 | 2 | 56 | 13 | 8 | 3 | 43 | 105 |
| 20113V1I1 | Fiji | Oceania | 2006 | invasive, NOS | emm100.0 | 119 | 11 | 11 | 7 | 10 | 10 | 7 | 37 | 39 | 19 | 2 | 30 | 31 |
| 20116V1I1 | Fiji | Oceania | 2006 | bacteremia | emm100.0 | 119 | 11 | 11 | 7 | 10 | 10 | 7 | 37 | 39 | 19 | 2 | 30 | 31 |
| 20123V1I1 | Fiji | Oceania | 2006 | bacteremia | emm100.0 | 119 | 11 | 11 | 7 | 10 | 10 | 7 | 37 | 39 | 19 | 2 | 30 | 31 |
| 20008V1I1 | Fiji | Oceania | 2006 | arthritis | emm33.0 | 3 | 11 | 11 | 7 | 10 | 35 | 36 | 85 | 79 | 1 | 2 | 6 | 20 |
| 20139V1I1 | Fiji | Oceania | 2006 | pneumonia | emm77.0 | 996 | 82 | 25 | 3 | 64 | 107 | 13 | 26 | 29 | 8 | 3 | 25 | 30 |
| 20148V1I1 | Fiji | Oceania | 2006 | bacteremia | emm106.0 | 140 | 11 | 69 | 7 | 10 | 106 | 7 | 88 | 36 | 15 | 3 | 64 | 34 |
| 20170V1I1 | Fiji | Oceania | 2006 | invasive, NOS | emm22.0 | 202 | 33 | 11 | 7 | 10 | 35 | 36 | 44 | 49 | 19 | 2 | 30 | 43 |
| 20174V1I1 | Fiji | Oceania | 2006 | arthritis | emm82.1 | 320 | 56 | 70 | 4 | 1 | 20 | 2 | 89 | 98 | 38 | 2 | 65 | 3 |
| 20178V1I1 | Fiji | Oceania | 2006 | bacteremia | emm11.0 | ND | 1 | 6 | 76 | 1 | 104 | 20 | 13 | 25 | 8 | 3 | 21 | 23 |
| 20183V1I1 | Fiji | Oceania | 2006 | bacteremia | emm57.0 | 1025 | 80 | 9 | 8 | 1 | 32 | 79 | 129 | 149 | 8 | 2 | 23 | 103 |
| 20184V1I1 | Fiji | Oceania | 2006 | necrotizing fasciitis | emm75.1 | 468 | 27 | 28 | 22 | 16 | 44 | 80 | 35 | 37 | 18 | 3 | 28 | 30 |
| 20232V1I1 | Fiji | Oceania | 2006 | invasive, NOS | emm101.0 | 182 | 15 | 31 | 10 | 20 | 15 | 31 | 38 | 41 | 1 | 3 | 32 | 37 |
| 20237V1I1 | Fiji | Oceania | 2006 | bacteremia | emm63.3 | 297 | 24 | 19 | 3 | 23 | 28 | 7 | 32 | 151 | 17 | 11 | 26 | 32 |
| 20242V1I1 | Fiji | Oceania | 2006 | bacteremia | emm19.4 | 616 | 50 | 32 | 10 | 1 | 6 | 55 | 78 | 13 | 1 | 2 | 6 | 106 |
| 20012V1I1 | Fiji | Oceania | 2006 | invasive, NOS | emm65.4 | 129 | 11 | 66 | 40 | 35 | 102 | 76 | 44 | 145 | 1 | 2 | 87 | 31 |
| 30083V1T1 | Fiji | Oceania | 2006 | pharyngitis and/or tonsillitis | emm86.2 | 963 | 11 | 32 | 10 | 20 | 15 | 16 | 128 | 13 | 1 | 2 | 6 | 100 |
| 30085V1T1 | Fiji | Oceania | 2006 | pharyngitis and/or tonsillitis | emm232.1 | 1013 | 18 | 18 | 16 | 16 | 21 | 25 | 125 | 26 | 8 | 3 | 21 | 23 |
| 30108V1T1 | Fiji | Oceania | 2006 | pharyngitis and/or tonsillitis | emm11.0 | ND | 1 | 6 | 76 | 1 | 20 | 20 | 13 | 25 | 8 | 3 | 21 | 23 |
| 30109V1T1 | Fiji | Oceania | 2006 | pharyngitis and/or tonsillitis | emm92.0 | 1026 | 5 | 65 | 34 | 27 | 39 | 39 | 22 | 152 | 13 | 2 | 20 | 22 |
| 30124V1T1 | Fiji | Oceania | 2006 | pharyngitis and/or tonsillitis | emm92.0 | 1026 | 5 | 65 | 34 | 27 | 39 | 39 | 22 | 152 | 13 | 2 | 20 | 22 |
| 30164V1T1 | Fiji | Oceania | 2006 | pharyngitis and/or tonsillitis | emm105.0 | 954 | 6 | 6 | 6 | 6 | 5 | 6 | 6 | 12 | 5 | 4 | 6 | 11 |
| 30178V1T1 | Fiji | Oceania | 2006 | pharyngitis and/or tonsillitis | emm123.0 | 325 | 6 | 6 | 6 | 6 | 5 | 6 | 6 | 7 | 5 | 4 | 6 | 6 |
| 30227V1T1 | Fiji | Oceania | 2006 | pharyngitis and/or tonsillitis | emm232.1 | 1013 | 18 | 18 | 16 | 16 | 21 | 25 | 125 | 26 | 8 | 3 | 21 | 23 |

| | | | | | | | | | | | | | | | | | | |
|---|---|---|---|---|---|---|---|---|---|---|---|---|---|---|---|---|---|---|
| 20018V1I1 | Fiji | Oceania | 2006 | necrotizing fasciitis | emm87.0 | 541 | 55 | 68 | 10 | 1 | 72 | 58 | 87 | 95 | 1 | 2 | 52 | 74 |
| 30237V1T1 | Fiji | Oceania | 2006 | pharyngitis and/or tonsillitis | emm86.2 | 963 | 11 | 32 | 10 | 20 | 15 | 16 | 128 | 13 | 1 | 2 | 6 | 100 |
| 30239V1T1 | Fiji | Oceania | 2006 | pharyngitis and/or tonsillitis | emm238.2 | 867 | 1 | 1 | 1 | 1 | 1 | 15 | 14 | 93 | 5 | 6 | 14 | 15 |
| 31002V1T1 | Fiji | Oceania | 2006 | pharyngitis and/or tonsillitis | emm56.0 | 115 | 14 | 44 | 24 | 2 | 12 | 41 | 51 | 153 | 23 | 1 | 6 | 49 |
| 31138V1T1 | Fiji | Oceania | 2006 | pharyngitis and/or tonsillitis | emm4.5 | 1027 | 14 | 51 | 42 | 39 | 47 | 33 | 132 | 79 | 8 | 3 | 91 | 107 |
| 31150V1T1 | Fiji | Oceania | 2006 | pharyngitis and/or tonsillitis | emm56.0 | 115 | 14 | 44 | 24 | 2 | 12 | 41 | 51 | 153 | 23 | 1 | 6 | 49 |
| 31168V1T1 | Fiji | Oceania | 2006 | pharyngitis and/or tonsillitis | emm101.0 | 182 | 15 | 31 | 10 | 20 | 15 | 31 | 38 | 41 | 1 | 3 | 32 | 37 |
| 20019V1I1 | Fiji | Oceania | 2006 | bacteremia | emm60.7 | 193 | 14 | 34 | 25 | 27 | 2 | 33 | 41 | 44 | 1 | 2 | 52 | 39 |
| 20025V1I1 | Fiji | Oceania | 2006 | arthritis | emm86.2 | 1028 | 11 | 32 | 10 | 20 | 15 | 16 | 128 | 13 | 1 | 2 | 6 | 100 |
| 31001V5S1 | Fiji | Oceania | 2006 | skin/soft tissue infection, NOS | emm11.0 | ND | 1 | 6 | 76 | 1 | 104 | 20 | 13 | 25 | 8 | 3 | 21 | 23 |
| 31002V5S1 | Fiji | Oceania | 2006 | skin/soft tissue infection, NOS | emm11.0 | ND | 1 | 6 | 76 | 1 | 20 | 20 | 13 | 25 | 8 | 3 | 21 | 23 |
| 31004V6S1 | Fiji | Oceania | 2006 | skin/soft tissue infection, NOS | emm49.0 | 433 | 4 | 8 | 4 | 11 | 14 | 14 | 16 | 18 | 9 | 2 | 12 | 13 |
| 31005V6S1 | Fiji | Oceania | 2006 | skin/soft tissue infection, NOS | emm39.4 | 268 | 1 | 60 | 49 | 45 | 61 | 19 | 13 | 83 | 34 | 3 | 61 | 29 |
| 31006V5S1 | Fiji | Oceania | 2006 | skin/soft tissue infection, NOS | emm103.0 | 327 | 14 | 13 | 3 | 1 | 66 | 13 | 82 | 87 | 1 | 2 | 49 | 4 |
| 20027V1I1 | Fiji | Oceania | 2006 | bacteremia | emm110.0 | 515 | 14 | 33 | 18 | 6 | 25 | 27 | 40 | 43 | 1 | 2 | 92 | 28 |
| 31009V5S1 | Fiji | Oceania | 2006 | skin/soft tissue infection, NOS | emm49.0 | 433 | 4 | 8 | 4 | 11 | 14 | 14 | 16 | 18 | 9 | 2 | 12 | 13 |
| 31022V5S1 | Fiji | Oceania | 2006 | skin/soft tissue infection, NOS | emm11.0 | ND | 1 | 6 | 76 | 1 | 20 | 20 | 13 | 25 | 8 | 3 | 21 | 23 |
| 31038V6S1 | Fiji | Oceania | 2006 | skin/soft tissue infection, NOS | emm103.0 | 327 | 14 | 13 | 3 | 1 | 66 | 13 | 82 | 87 | 1 | 2 | 49 | 4 |
| 31039V5S1 | Fiji | Oceania | 2006 | skin/soft tissue infection, NOS | emm49.0 | 433 | 4 | 8 | 4 | 11 | 14 | 14 | 16 | 18 | 9 | 2 | 12 | 13 |
| 31039V6S1 | Fiji | Oceania | 2006 | skin/soft tissue infection, NOS | emm93.4 | 814 | 28 | 104 | 10 | 1 | 108 | 32 | 61 | 42 | 50 | 2 | 93 | 69 |
| 31042V5S1 | Fiji | Oceania | 2006 | skin/soft tissue infection, NOS | emm56.0 | 115 | 14 | 44 | 24 | 2 | 12 | 41 | 51 | 153 | 23 | 1 | 6 | 49 |
| 31043V6S1 | Fiji | Oceania | 2006 | skin/soft tissue infection, NOS | emm56.0 | 115 | 14 | 44 | 24 | 2 | 12 | 41 | 51 | 153 | 23 | 1 | 6 | 49 |
| 31046V5S1 | Fiji | Oceania | 2006 | skin/soft tissue infection, NOS | emm82.1 | 320 | 56 | 70 | 4 | 1 | 20 | 2 | 89 | 98 | 38 | 2 | 65 | 3 |
| 31072V6S2 | Fiji | Oceania | 2006 | skin/soft tissue infection, NOS | emm33.0 | 3 | 11 | 11 | 7 | 10 | 35 | 36 | 85 | 79 | 1 | 2 | 6 | 20 |
| 31073V5S1 | Fiji | Oceania | 2006 | skin/soft tissue infection, NOS | emm76.4 | 353 | 45 | 34 | 48 | 8 | 58 | 50 | 71 | 57 | 18 | 3 | 22 | 102 |
| 31075V5S1 | Fiji | Oceania | 2006 | skin/soft tissue infection, NOS | emm56.0 | 115 | 14 | 44 | 24 | 2 | 12 | 41 | 51 | 153 | 23 | 1 | 6 | 49 |
| 31086V5S1 | Fiji | Oceania | 2006 | skin/soft tissue infection, NOS | emm65.4 | 129 | 11 | 66 | 40 | 35 | 68 | 76 | 44 | 145 | 1 | 2 | 87 | 31 |
| 31090V5S1 | Fiji | Oceania | 2006 | skin/soft tissue infection, NOS | emm11.0 | ND | 1 | 6 | 76 | 1 | 104 | 20 | 13 | 25 | 8 | 3 | 21 | 23 |
| 31097V5S1 | Fiji | Oceania | 2006 | skin/soft tissue infection, NOS | emm56.0 | 115 | 14 | 44 | 24 | 2 | 12 | 41 | 51 | 153 | 23 | 1 | 6 | 49 |
| 31107V5S1 | Fiji | Oceania | 2006 | skin/soft tissue infection, NOS | emm65.4 | 129 | 11 | 66 | 40 | 35 | 102 | 76 | 44 | 145 | 1 | 2 | 87 | 31 |
| 31108V5S1 | Fiji | Oceania | 2006 | skin/soft tissue infection, NOS | emm93.4 | 814 | 28 | 104 | 10 | 1 | 108 | 32 | 61 | 42 | 50 | 2 | 93 | 69 |
| 31113V5S1 | Fiji | Oceania | 2006 | skin/soft tissue infection, NOS | emm74.0 | 120 | 28 | 13 | 75 | 41 | 50 | 53 | 123 | 82 | 11 | 4 | 59 | 70 |
| 31114V5S1 | Fiji | Oceania | 2006 | skin/soft tissue infection, NOS | emm39.4 | 268 | 1 | 60 | 49 | 45 | 61 | 19 | 13 | 83 | 34 | 3 | 61 | 29 |
| 31049V5S1 | Fiji | Oceania | 2006 | skin/soft tissue infection, NOS | emm65.4 | 129 | 11 | 66 | 40 | 35 | 68 | 76 | 44 | 145 | 1 | 2 | 87 | 31 |
| 31114V6S1 | Fiji | Oceania | 2006 | skin/soft tissue infection, NOS | emm103.0 | 327 | 14 | 13 | 3 | 1 | 66 | 13 | 82 | 87 | 1 | 2 | 49 | 4 |
| 31126V5S1 | Fiji | Oceania | 2006 | skin/soft tissue infection, NOS | emm65.4 | 129 | 11 | 66 | 40 | 35 | 68 | 76 | 44 | 145 | 1 | 2 | 87 | 31 |
| 31128V6S1 | Fiji | Oceania | 2006 | skin/soft tissue infection, NOS | emm70.0 | 10 | 16 | 16 | 3 | 16 | 109 | 7 | 20 | 22 | 1 | 2 | 18 | 3 |
| 31134V5S1 | Fiji | Oceania | 2006 | skin/soft tissue infection, NOS | emm11.0 | ND | 1 | 6 | 76 | 1 | 104 | 20 | 13 | 25 | 8 | 3 | 21 | 23 |
| 31136V5S1 | Fiji | Oceania | 2006 | skin/soft tissue infection, NOS | emm56.0 | 115 | 14 | 44 | 24 | 2 | 12 | 41 | 51 | 153 | 23 | 1 | 6 | 49 |
| 31136V6S1 | Fiji | Oceania | 2006 | skin/soft tissue infection, NOS | emm39.4 | 268 | 1 | 60 | 49 | 45 | 61 | 19 | 13 | 83 | 34 | 3 | 61 | 29 |
| 31138V6S1 | Fiji | Oceania | 2006 | skin/soft tissue infection, NOS | emm39.4 | 268 | 1 | 60 | 49 | 45 | 61 | 19 | 13 | 83 | 34 | 3 | 61 | 29 |
| 31142V6S1 | Fiji | Oceania | 2006 | skin/soft tissue infection, NOS | emm93.4 | 814 | 28 | 104 | 10 | 1 | 108 | 32 | 61 | 42 | 50 | 2 | 93 | 69 |
| 31144V6S1 | Fiji | Oceania | 2006 | skin/soft tissue infection, NOS | emm105.0 | 954 | 6 | 6 | 6 | 6 | 5 | 6 | 6 | 12 | 5 | 4 | 6 | 11 |
| 31049V5S2 | Fiji | Oceania | 2006 | skin/soft tissue infection, NOS | emm70.0 | 10 | 16 | 16 | 3 | 16 | 17 | 7 | 20 | 22 | 1 | 2 | 18 | 3 |
| 31145V5S1 | Fiji | Oceania | 2006 | skin/soft tissue infection, NOS | emm93.4 | 814 | 28 | 104 | 10 | 1 | 108 | 32 | 61 | 42 | 50 | 2 | 93 | 69 |
| 31154V5S1 | Fiji | Oceania | 2006 | skin/soft tissue infection, NOS | emm4.5 | 1027 | 14 | 51 | 42 | 39 | 47 | 33 | 132 | 79 | 8 | 3 | 91 | 107 |
| 31155V6S1 | Fiji | Oceania | 2006 | skin/soft tissue infection, NOS | emm56.0 | 115 | 14 | 44 | 24 | 2 | 12 | 41 | 51 | 153 | 23 | 1 | 6 | 49 |
| 31158V5S1 | Fiji | Oceania | 2006 | skin/soft tissue infection, NOS | emm49.0 | 433 | 4 | 8 | 4 | 11 | 14 | 14 | 16 | 18 | 9 | 2 | 12 | 13 |

| | | | | | | | | | | | | | | | | | | |
|---|---|---|---|---|---|---|---|---|---|---|---|---|---|---|---|---|---|---|
| 31163V5S1 | Fiji | Oceania | 2006 | skin/soft tissue infection, NOS | emm70.0 | 10 | 16 | 16 | 3 | 16 | 109 | 7 | 20 | 22 | 1 | 2 | 18 | 3 |
| 31165V6S1 | Fiji | Oceania | 2006 | skin/soft tissue infection, NOS | emm39.4 | 268 | 1 | 60 | 49 | 45 | 61 | 19 | 13 | 83 | 34 | 3 | 61 | 29 |
| 31168V5S1 | Fiji | Oceania | 2006 | skin/soft tissue infection, NOS | emm39.4 | 268 | 1 | 60 | 49 | 45 | 61 | 19 | 13 | 83 | 34 | 3 | 61 | 29 |
| 31170V5S1 | Fiji | Oceania | 2006 | skin/soft tissue infection, NOS | emm93.4 | 814 | 28 | 104 | 10 | 1 | 108 | 32 | 61 | 42 | 50 | 2 | 93 | 69 |
| 41095V1S1 | Fiji | Oceania | 2006 | skin/soft tissue infection, NOS | emm63.3 | 297 | 24 | 19 | 3 | 23 | 28 | 7 | 32 | 151 | 17 | 11 | 26 | 32 |
| 41126V1S1 | Fiji | Oceania | 2006 | skin/soft tissue infection, NOS | emm11.0 | ND | 1 | 6 | 76 | 1 | 20 | 20 | 13 | 25 | 8 | 3 | 21 | 23 |
| 31050V6S1 | Fiji | Oceania | 2006 | skin/soft tissue infection, NOS | emm73.0 | 957 | 14 | 18 | 4 | 16 | 21 | 61 | 27 | 26 | 46 | 3 | 66 | 87 |
| 42003V1S1 | Fiji | Oceania | 2006 | skin/soft tissue infection, NOS | emm71.0 | 318 | 19 | 20 | 13 | 19 | 2 | 22 | 3 | 27 | 1 | 1 | 4 | 24 |
| 42023V1S1 | Fiji | Oceania | 2006 | skin/soft tissue infection, NOS | emm55.0 | 100 | 11 | 11 | 7 | 10 | 10 | 13 | 11 | 13 | 8 | 3 | 11 | 12 |
| 42024V1S1 | Fiji | Oceania | 2006 | skin/soft tissue infection, NOS | emm11.0 | ND | 1 | 6 | 76 | 1 | 104 | 20 | 13 | 25 | 8 | 3 | 21 | 23 |
| 42028V1S1 | Fiji | Oceania | 2006 | skin/soft tissue infection, NOS | emm100.0 | 119 | 11 | 11 | 7 | 10 | 10 | 7 | 37 | 39 | 19 | 2 | 30 | 31 |
| 42043V1S1 | Fiji | Oceania | 2006 | skin/soft tissue infection, NOS | emm74.0 | 120 | 28 | 13 | 75 | 41 | 50 | 53 | 123 | 82 | 11 | 4 | 59 | 70 |
| 42048V1S1 | Fiji | Oceania | 2006 | skin/soft tissue infection, NOS | emm81.3 | 995 | 8 | 4 | 4 | 4 | 3 | 13 | 4 | 42 | 21 | 1 | 4 | 4 |
| 42056V1S1a | Fiji | Oceania | 2006 | skin/soft tissue infection, NOS | emm25.0 | 191 | 30 | 35 | 26 | 28 | 35 | 34 | 42 | 45 | 11 | 2 | 12 | 40 |
| 42058V1S1 | Fiji | Oceania | 2006 | skin/soft tissue infection, NOS | emm74.1 | 468 | 27 | 28 | 22 | 26 | 31 | 77 | 35 | 37 | 18 | 3 | 28 | 30 |
| 31056V5S1 | Fiji | Oceania | 2006 | skin/soft tissue infection, NOS | emm11.0 | ND | 1 | 6 | 76 | 1 | 104 | 20 | 13 | 25 | 8 | 3 | 21 | 23 |
| 42063V1S1 | Fiji | Oceania | 2006 | skin/soft tissue infection, NOS | emm97.1 | 216 | 21 | 21 | 15 | 8 | 46 | 24 | 133 | 30 | 8 | 10 | 12 | 26 |
| 42070V1S1 | Fiji | Oceania | 2006 | skin/soft tissue infection, NOS | emm44.0 | 178 | 5 | 7 | 7 | 3 | 2 | 3 | 24 | 8 | 14 | 3 | 7 | 7 |
| 43078V1S1 | Fiji | Oceania | 2006 | skin/soft tissue infection, NOS | ND | 1029 | 15 | 15 | 10 | 1 | 110 | 7 | 134 | 22 | 23 | 1 | 6 | 108 |
| 43086V1S1 | Fiji | Oceania | 2006 | skin/soft tissue infection, NOS | emm11.0 | ND | 1 | 6 | 76 | 1 | 20 | 20 | 13 | 25 | 8 | 3 | 21 | 23 |
| 43110V1S1 | Fiji | Oceania | 2006 | skin/soft tissue infection, NOS | emm57.0 | 1030 | 80 | 9 | 8 | 1 | 32 | 79 | 129 | 149 | 8 | 2 | 23 | 103 |
| 44001V1S1 | Fiji | Oceania | 2006 | skin/soft tissue infection, NOS | emm123.0 | 325 | 6 | 6 | 6 | 6 | 5 | 6 | 6 | 7 | 5 | 4 | 6 | 6 |
| 44002V1S1 | Fiji | Oceania | 2006 | skin/soft tissue infection, NOS | emm103.0 | 327 | 14 | 13 | 3 | 1 | 66 | 13 | 82 | 87 | 1 | 2 | 49 | 4 |
| 44004V1S1 | Fiji | Oceania | 2006 | skin/soft tissue infection, NOS | emm82.1 | 591 | 56 | 70 | 4 | 1 | 20 | 2 | 135 | 98 | 38 | 2 | 65 | 3 |
| 44015V1S1 | Fiji | Oceania | 2006 | skin/soft tissue infection, NOS | emm81.3 | 995 | 8 | 4 | 4 | 4 | 3 | 13 | 4 | 42 | 21 | 1 | 4 | 4 |
| 31056V6S1 | Fiji | Oceania | 2006 | skin/soft tissue infection, NOS | emm11.0 | ND | 1 | 6 | 76 | 1 | 104 | 20 | 13 | 25 | 8 | 3 | 21 | 23 |
| 44031V1S1 | Fiji | Oceania | 2006 | skin/soft tissue infection, NOS | emm82.1 | 320 | 56 | 70 | 4 | 1 | 20 | 2 | 89 | 98 | 38 | 2 | 65 | 3 |
| 44037V1S1 | Fiji | Oceania | 2006 | skin/soft tissue infection, NOS | emm116.1 | 227 | 28 | 105 | 32 | 16 | 111 | 63 | 87 | 154 | 51 | 4 | 11 | 77 |
| 44052V1S1 | Fiji | Oceania | 2006 | skin/soft tissue infection, NOS | emm44.0 | 178 | 5 | 7 | 7 | 3 | 2 | 3 | 24 | 8 | 14 | 3 | 7 | 7 |
| 44079V1S1 | Fiji | Oceania | 2006 | skin/soft tissue infection, NOS | emm87.0 | 541 | 55 | 68 | 10 | 1 | 72 | 58 | 87 | 95 | 52 | 2 | 52 | 74 |
| 31004V2S1 | Fiji | Oceania | 2006 | skin/soft tissue infection, NOS | emm70.0 | 10 | 16 | 16 | 3 | 16 | 17 | 7 | 20 | 22 | 1 | 2 | 18 | 3 |
| 31006V2S1 | Fiji | Oceania | 2006 | skin/soft tissue infection, NOS | emm70.0 | 10 | 16 | 16 | 3 | 16 | 17 | 7 | 20 | 22 | 1 | 2 | 18 | 3 |
| 31012V2S1 | Fiji | Oceania | 2006 | skin/soft tissue infection, NOS | emm73.4 | 814 | 28 | 104 | 10 | 1 | 108 | 32 | 61 | 42 | 50 | 2 | 93 | 69 |
| 31059V6S1 | Fiji | Oceania | 2006 | skin/soft tissue infection, NOS | emm53.0 | 11 | 11 | 5 | 10 | 8 | 15 | 54 | 17 | 84 | 1 | 2 | 6 | 20 |
| 31026V2S1 | Fiji | Oceania | 2006 | skin/soft tissue infection, NOS | emm105.0 | 954 | 6 | 6 | 6 | 6 | 5 | 6 | 6 | 12 | 5 | 4 | 6 | 11 |
| 31027V2S1 | Fiji | Oceania | 2006 | skin/soft tissue infection, NOS | emm52.1 | 180 | 15 | 17 | 10 | 17 | 18 | 18 | 131 | 23 | 12 | 8 | 19 | 21 |
| 31028V2S1 | Fiji | Oceania | 2006 | skin/soft tissue infection, NOS | emm73.0 | 957 | 14 | 18 | 4 | 16 | 21 | 61 | 27 | 26 | 46 | 3 | 66 | 87 |
| 31041V2S1 | Fiji | Oceania | 2006 | skin/soft tissue infection, NOS | emm70.0 | 10 | 16 | 16 | 3 | 16 | 17 | 7 | 20 | 22 | 1 | 2 | 18 | 3 |
| 31056V2S1 | Fiji | Oceania | 2006 | skin/soft tissue infection, NOS | emm65.4 | 129 | 11 | 66 | 40 | 35 | 68 | 76 | 44 | 145 | 1 | 2 | 87 | 31 |
| 31067V2S1 | Fiji | Oceania | 2006 | skin/soft tissue infection, NOS | emm33.0 | 3 | 11 | 11 | 7 | 10 | 35 | 36 | 85 | 79 | 1 | 2 | 6 | 20 |
| 31069V2S1 | Fiji | Oceania | 2006 | skin/soft tissue infection, NOS | emm82.1 | 320 | 56 | 70 | 4 | 1 | 20 | 2 | 89 | 98 | 38 | 2 | 65 | 3 |
| 31076V2S1 | Fiji | Oceania | 2006 | skin/soft tissue infection, NOS | emm56.0 | 115 | 14 | 44 | 24 | 2 | 12 | 41 | 51 | 153 | 23 | 1 | 6 | 49 |
| 31062V6S1 | Fiji | Oceania | 2006 | skin/soft tissue infection, NOS | emm103.0 | 327 | 14 | 13 | 3 | 1 | 66 | 13 | 82 | 87 | 1 | 2 | 49 | 4 |
| 31082V2S1 | Fiji | Oceania | 2006 | skin/soft tissue infection, NOS | emm11.0 | ND | 1 | 6 | 76 | 1 | 104 | 20 | 13 | 25 | 8 | 3 | 21 | 23 |
| 31084V2S1 | Fiji | Oceania | 2006 | skin/soft tissue infection, NOS | emm70.0 | 10 | 16 | 16 | 3 | 16 | 17 | 7 | 20 | 22 | 1 | 2 | 18 | 3 |
| 31072V6S1 | Fiji | Oceania | 2006 | skin/soft tissue infection, NOS | emm11.0 | ND | 1 | 6 | 76 | 1 | 20 | 20 | 13 | 25 | 8 | 3 | 21 | 23 |
| 31011V3T1 | Fiji | Oceania | 2006 | pharyngitis and/or tonsillitis | emm65.4 | 129 | 11 | 66 | 40 | 35 | 68 | 76 | 44 | 145 | 1 | 2 | 87 | 31 |
| 31034V1S1 | Fiji | Oceania | 2006 | skin/soft tissue infection, NOS | emm105.0 | 954 | 6 | 6 | 6 | 6 | 5 | 6 | 6 | 12 | 5 | 4 | 6 | 11 |

| | | | | | | | | | | | | | | | | | | |
|---|---|---|---|---|---|---|---|---|---|---|---|---|---|---|---|---|---|---|
| 31036V1S1 | Fiji | Oceania | 2006 | skin/soft tissue infection, NOS | emm25.0 | 1032 | 30 | 35 | 26 | 28 | 35 | 34 | 42 | 45 | 11 | 2 | 12 | 40 |
| 31039V1S1 | Fiji | Oceania | 2006 | skin/soft tissue infection, NOS | emm25.0 | 1032 | 30 | 35 | 26 | 28 | 35 | 34 | 42 | 45 | 11 | 2 | 12 | 40 |
| 31043V1S1 | Fiji | Oceania | 2006 | skin/soft tissue infection, NOS | emm70.0 | 10 | 16 | 16 | 3 | 16 | 17 | 7 | 20 | 22 | 1 | 2 | 18 | 3 |
| 31063V1S1 | Fiji | Oceania | 2006 | skin/soft tissue infection, NOS | emm73.0 | 957 | 14 | 18 | 4 | 16 | 21 | 61 | 27 | 26 | 46 | 3 | 66 | 87 |
| 31065V1S1 | Fiji | Oceania | 2006 | skin/soft tissue infection, NOS | emm56.0 | 115 | 14 | 44 | 24 | 2 | 12 | 41 | 51 | 153 | 23 | 1 | 6 | 49 |
| 31068V1S1 | Fiji | Oceania | 2006 | skin/soft tissue infection, NOS | emm73.0 | 957 | 14 | 18 | 4 | 16 | 21 | 61 | 27 | 26 | 46 | 3 | 66 | 87 |
| 31078V1S1 | Fiji | Oceania | 2006 | skin/soft tissue infection, NOS | emm25.1 | 192 | 14 | 101 | 24 | 2 | 33 | 13 | 13 | 40 | 36 | 3 | 31 | 36 |
| 31080V1S1 | Fiji | Oceania | 2006 | skin/soft tissue infection, NOS | emm25.1 | 192 | 14 | 101 | 24 | 2 | 33 | 13 | 13 | 40 | 36 | 3 | 31 | 36 |
| 31013V2T1 | Fiji | Oceania | 2006 | pharyngitis and/or tonsillitis | emm39.4 | 268 | 1 | 60 | 49 | 45 | 61 | 19 | 13 | 83 | 34 | 3 | 61 | 29 |
| 31081V1S1 | Fiji | Oceania | 2006 | skin/soft tissue infection, NOS | emm73.0 | 957 | 14 | 18 | 4 | 16 | 21 | 61 | 27 | 26 | 46 | 3 | 66 | 87 |
| 31082V1S1 | Fiji | Oceania | 2006 | skin/soft tissue infection, NOS | emm73.0 | 957 | 14 | 18 | 4 | 16 | 21 | 61 | 27 | 26 | 46 | 3 | 66 | 87 |
| 31086V1S1 | Fiji | Oceania | 2006 | skin/soft tissue infection, NOS | emm73.0 | 957 | 14 | 18 | 4 | 16 | 21 | 61 | 27 | 26 | 46 | 3 | 66 | 87 |
| 31106V1S1 | Fiji | Oceania | 2006 | skin/soft tissue infection, NOS | emm25.0 | 1032 | 30 | 35 | 26 | 28 | 35 | 34 | 42 | 45 | 11 | 2 | 12 | 40 |
| 31111V1S1 | Fiji | Oceania | 2006 | skin/soft tissue infection, NOS | emm93.4 | 814 | 28 | 104 | 10 | 1 | 108 | 32 | 61 | 42 | 50 | 2 | 93 | 69 |
| 31114V1S1 | Fiji | Oceania | 2006 | skin/soft tissue infection, NOS | emm25.0 | 1032 | 30 | 35 | 26 | 28 | 35 | 34 | 42 | 45 | 11 | 2 | 12 | 40 |
| 31122V1S1 | Fiji | Oceania | 2006 | skin/soft tissue infection, NOS | emm207.1 | 332 | 11 | 38 | 32 | 16 | 38 | 18 | 48 | 53 | 1 | 2 | 6 | 20 |
| 31129V1S2 | Fiji | Oceania | 2006 | skin/soft tissue infection, NOS | emm73.0 | 957 | 14 | 18 | 4 | 16 | 21 | 61 | 27 | 26 | 46 | 3 | 66 | 87 |
| 31023V2T1 | Fiji | Oceania | 2006 | pharyngitis and/or tonsillitis | emm56.0 | 115 | 14 | 44 | 24 | 2 | 12 | 41 | 51 | 153 | 23 | 1 | 6 | 49 |
| 31132V1S1 | Fiji | Oceania | 2006 | skin/soft tissue infection, NOS | emm25.0 | 1032 | 30 | 35 | 26 | 28 | 35 | 34 | 42 | 45 | 11 | 2 | 12 | 40 |
| 31140V1S1 | Fiji | Oceania | 2006 | skin/soft tissue infection, NOS | emm98.1 | 205 | 15 | 15 | 10 | 1 | 10 | 18 | 19 | 21 | 12 | 2 | 6 | 20 |
| 31143V1S1 | Fiji | Oceania | 2006 | skin/soft tissue infection, NOS | emm71.0 | 318 | 19 | 20 | 13 | 19 | 2 | 22 | 3 | 27 | 1 | 1 | 4 | 24 |
| 31147V1S1 | Fiji | Oceania | 2006 | skin/soft tissue infection, NOS | emm25.0 | 1032 | 30 | 35 | 26 | 28 | 35 | 34 | 42 | 45 | 11 | 2 | 12 | 40 |
| 31150V1S1 | Fiji | Oceania | 2006 | skin/soft tissue infection, NOS | emm56.0 | 115 | 14 | 44 | 24 | 2 | 12 | 41 | 51 | 153 | 23 | 1 | 6 | 49 |
| 31158V1S1 | Fiji | Oceania | 2006 | skin/soft tissue infection, NOS | emm25.0 | 1032 | 30 | 35 | 26 | 28 | 35 | 34 | 42 | 45 | 11 | 2 | 12 | 40 |
| 31165V1S1 | Fiji | Oceania | 2006 | skin/soft tissue infection, NOS | emm25.0 | 1032 | 30 | 35 | 26 | 28 | 35 | 34 | 42 | 45 | 11 | 2 | 12 | 40 |
| 31167V1S1 | Fiji | Oceania | 2006 | skin/soft tissue infection, NOS | emm56.0 | 115 | 14 | 44 | 24 | 2 | 12 | 41 | 51 | 153 | 23 | 1 | 6 | 49 |
| 31175V1S1 | Fiji | Oceania | 2006 | skin/soft tissue infection, NOS | emm25.0 | 1032 | 30 | 35 | 26 | 28 | 35 | 34 | 42 | 45 | 11 | 2 | 12 | 40 |
| 31039V5T1 | Fiji | Oceania | 2006 | pharyngitis and/or tonsillitis | emm74.0 | 120 | 28 | 13 | 75 | 41 | 50 | 53 | 123 | 82 | 11 | 4 | 59 | 70 |
| 31176V1S1 | Fiji | Oceania | 2006 | skin/soft tissue infection, NOS | emm89.14 | 380 | 14 | 49 | 20 | 16 | 112 | 52 | 136 | 80 | 33 | 11 | 57 | 25 |
| 33014V1S1 | Fiji | Oceania | 2006 | skin/soft tissue infection, NOS | emm105.0 | 954 | 6 | 6 | 6 | 6 | 5 | 6 | 6 | 12 | 5 | 4 | 6 | 11 |
| 33017V1S1 | Fiji | Oceania | 2006 | skin/soft tissue infection, NOS | emm92.0 | 1026 | 5 | 65 | 34 | 27 | 39 | 39 | 22 | 152 | 13 | 2 | 20 | 22 |
| 33022V1S1 | Fiji | Oceania | 2006 | skin/soft tissue infection, NOS | emm82.1 | 320 | 56 | 70 | 4 | 1 | 20 | 2 | 89 | 98 | 38 | 2 | 65 | 3 |
| 33041V1S1 | Fiji | Oceania | 2006 | skin/soft tissue infection, NOS | emm33.0 | 3 | 11 | 11 | 7 | 10 | 35 | 36 | 85 | 79 | 1 | 2 | 6 | 20 |
| 33048V1S1 | Fiji | Oceania | 2006 | skin/soft tissue infection, NOS | emm25.0 | 1032 | 30 | 35 | 26 | 28 | 35 | 34 | 42 | 45 | 11 | 2 | 12 | 40 |
| 33050V1S1 | Fiji | Oceania | 2006 | skin/soft tissue infection, NOS | emm25.1 | 192 | 14 | 101 | 24 | 2 | 33 | 13 | 13 | 40 | 36 | 3 | 31 | 36 |
| 33052V1S1 | Fiji | Oceania | 2006 | skin/soft tissue infection, NOS | emm75.1 | 468 | 27 | 28 | 22 | 26 | 31 | 77 | 35 | 37 | 18 | 3 | 28 | 30 |
| 33061V1S1 | Fiji | Oceania | 2006 | skin/soft tissue infection, NOS | emm105.0 | 954 | 6 | 6 | 6 | 6 | 5 | 6 | 6 | 12 | 5 | 4 | 6 | 11 |
| 31097V2T1 | Fiji | Oceania | 2006 | pharyngitis and/or tonsillitis | emm82.1 | 320 | 56 | 70 | 4 | 1 | 20 | 2 | 89 | 98 | 38 | 2 | 65 | 3 |
| 33080V1S1 | Fiji | Oceania | 2006 | skin/soft tissue infection, NOS | emm92.0 | 1026 | 5 | 65 | 34 | 27 | 39 | 39 | 22 | 152 | 13 | 2 | 20 | 22 |
| 33082V1S1 | Fiji | Oceania | 2006 | skin/soft tissue infection, NOS | emm71.0 | 318 | 19 | 20 | 13 | 19 | 2 | 22 | 3 | 27 | 1 | 1 | 88 | 24 |
| 33089V1S1 | Fiji | Oceania | 2006 | skin/soft tissue infection, NOS | emm54.1 | 990 | 11 | 5 | 10 | 20 | 15 | 16 | 17 | 13 | 1 | 2 | 16 | 31 |
| 33100V1S1 | Fiji | Oceania | 2006 | skin/soft tissue infection, NOS | emm54.1 | 990 | 11 | 5 | 10 | 20 | 15 | 16 | 17 | 13 | 1 | 2 | 16 | 31 |
| 33107V1S1 | Fiji | Oceania | 2006 | skin/soft tissue infection, NOS | emm73.0 | 957 | 14 | 18 | 4 | 16 | 21 | 61 | 27 | 26 | 46 | 3 | 66 | 87 |
| 33117V1S1 | Fiji | Oceania | 2006 | skin/soft tissue infection, NOS | emm33.0 | 3 | 11 | 11 | 7 | 10 | 35 | 36 | 85 | 79 | 1 | 2 | 6 | 20 |
| 33122V1S1 | Fiji | Oceania | 2006 | skin/soft tissue infection, NOS | emm100.0 | 119 | 11 | 11 | 7 | 10 | 10 | 7 | 37 | 39 | 19 | 2 | 30 | 31 |
| 33122V1S2 | Fiji | Oceania | 2006 | skin/soft tissue infection, NOS | emm86.2 | 963 | 11 | 32 | 10 | 20 | 15 | 16 | 128 | 13 | 1 | 2 | 6 | 100 |
| 33124V1S1 | Fiji | Oceania | 2006 | skin/soft tissue infection, NOS | emm65.4 | 129 | 11 | 66 | 40 | 35 | 102 | 76 | 44 | 145 | 1 | 2 | 87 | 31 |
| 31001V1S1 | Fiji | Oceania | 2006 | skin/soft tissue infection, NOS | emm25.0 | 1032 | 30 | 35 | 26 | 28 | 35 | 34 | 42 | 45 | 11 | 2 | 12 | 40 |

| | | | | | | | | | | | | | | | | | | |
|---|---|---|---|---|---|---|---|---|---|---|---|---|---|---|---|---|---|---|
| 33134V1S1 | Fiji | Oceania | 2006 | skin/soft tissue infection, NOS | emm58.0 | 176 | 14 | 13 | 73 | 29 | 13 | 13 | 126 | 47 | 10 | 2 | 34 | 31 |
| 33153V1S1 | Fiji | Oceania | 2006 | skin/soft tissue infection, NOS | emm65.4 | 129 | 11 | 66 | 40 | 35 | 102 | 76 | 44 | 145 | 1 | 2 | 87 | 31 |
| 33156V1S1 | Fiji | Oceania | 2006 | skin/soft tissue infection, NOS | emm86.2 | 963 | 11 | 32 | 10 | 20 | 15 | 16 | 128 | 13 | 1 | 2 | 6 | 100 |
| 33156V1S2 | Fiji | Oceania | 2006 | skin/soft tissue infection, NOS | emm86.2 | 963 | 11 | 32 | 10 | 20 | 15 | 16 | 128 | 13 | 1 | 2 | 6 | 100 |
| 33187V1S1 | Fiji | Oceania | 2006 | skin/soft tissue infection, NOS | emm25.1 | 192 | 14 | 101 | 24 | 2 | 33 | 13 | 13 | 40 | 36 | 3 | 31 | 36 |
| 33192V1S1 | Fiji | Oceania | 2006 | skin/soft tissue infection, NOS | emm103.0 | 327 | 14 | 13 | 3 | 1 | 66 | 13 | 82 | 87 | 1 | 2 | 49 | 4 |
| 33193V1S1 | Fiji | Oceania | 2006 | skin/soft tissue infection, NOS | emm25.1 | 192 | 14 | 101 | 24 | 2 | 33 | 13 | 13 | 40 | 36 | 3 | 31 | 36 |
| 33194V1S1 | Fiji | Oceania | 2006 | skin/soft tissue infection, NOS | emm100.0 | 119 | 11 | 11 | 7 | 10 | 10 | 7 | 37 | 39 | 19 | 2 | 30 | 31 |
| 33201V1S1 | Fiji | Oceania | 2006 | skin/soft tissue infection, NOS | emm92.0 | 1026 | 5 | 65 | 34 | 27 | 39 | 39 | 22 | 152 | 13 | 2 | 20 | 22 |
| 33052V2S1 | Fiji | Oceania | 2006 | skin/soft tissue infection, NOS | emm25.0 | 1032 | 30 | 35 | 26 | 28 | 35 | 34 | 42 | 45 | 11 | 2 | 12 | 40 |
| 31010V1S1 | Fiji | Oceania | 2006 | skin/soft tissue infection, NOS | emm25.0 | 1032 | 30 | 35 | 26 | 28 | 35 | 34 | 42 | 45 | 11 | 2 | 12 | 40 |
| 33094V2S1 | Fiji | Oceania | 2006 | skin/soft tissue infection, NOS | emm25.0 | 1032 | 30 | 35 | 26 | 28 | 35 | 34 | 42 | 45 | 11 | 2 | 12 | 40 |
| 33120V2S1 | Fiji | Oceania | 2006 | skin/soft tissue infection, NOS | emm33.0 | 3 | 11 | 11 | 7 | 10 | 35 | 36 | 85 | 79 | 1 | 2 | 6 | 20 |
| 33123V2S1 | Fiji | Oceania | 2006 | skin/soft tissue infection, NOS | emm71.0 | 318 | 19 | 20 | 13 | 19 | 2 | 22 | 3 | 27 | 1 | 1 | 4 | 24 |
| 33139V2S1 | Fiji | Oceania | 2006 | skin/soft tissue infection, NOS | emm85.0 | 109 | 10 | 10 | 4 | 9 | 9 | 11 | 72 | 11 | 7 | 3 | 10 | 10 |
| 33158V2S1 | Fiji | Oceania | 2006 | skin/soft tissue infection, NOS | emm19.4 | 616 | 50 | 32 | 10 | 1 | 6 | 55 | 78 | 13 | 1 | 2 | 6 | 106 |
| 33169V2S1 | Fiji | Oceania | 2006 | skin/soft tissue infection, NOS | emm19.4 | 616 | 50 | 32 | 10 | 1 | 6 | 55 | 78 | 13 | 1 | 2 | 6 | 106 |
| 33169V2S2 | Fiji | Oceania | 2006 | skin/soft tissue infection, NOS | emm19.4 | 616 | 50 | 32 | 10 | 1 | 6 | 55 | 78 | 13 | 1 | 2 | 6 | 106 |
| 33188V2S2 | Fiji | Oceania | 2006 | skin/soft tissue infection, NOS | emm73.0 | 957 | 14 | 18 | 4 | 16 | 21 | 61 | 27 | 26 | 46 | 3 | 66 | 87 |
| 33189V2S2 | Fiji | Oceania | 2006 | skin/soft tissue infection, NOS | emm100.0 | 119 | 11 | 11 | 7 | 10 | 10 | 7 | 37 | 39 | 19 | 2 | 30 | 31 |
| 33190V2S1 | Fiji | Oceania | 2006 | skin/soft tissue infection, NOS | emm33.0 | 3 | 11 | 11 | 7 | 10 | 35 | 36 | 85 | 79 | 1 | 2 | 6 | 20 |
| 31012V1S1 | Fiji | Oceania | 2006 | skin/soft tissue infection, NOS | emm73.0 | 957 | 14 | 18 | 4 | 16 | 21 | 61 | 27 | 26 | 46 | 3 | 66 | 87 |
| 33191V2S1 | Fiji | Oceania | 2006 | skin/soft tissue infection, NOS | emm52.1 | 180 | 15 | 17 | 10 | 17 | 18 | 18 | 131 | 23 | 12 | 8 | 19 | 21 |
| 33192V2S1 | Fiji | Oceania | 2006 | skin/soft tissue infection, NOS | emm25.0 | 1032 | 30 | 35 | 26 | 28 | 35 | 34 | 42 | 45 | 11 | 2 | 12 | 40 |
| 33194V2S1 | Fiji | Oceania | 2006 | skin/soft tissue infection, NOS | emm101.0 | 182 | 15 | 31 | 10 | 20 | 15 | 31 | 38 | 41 | 1 | 3 | 32 | 37 |
| 33011V3S1 | Fiji | Oceania | 2006 | skin/soft tissue infection, NOS | emm44.0 | 178 | 5 | 7 | 7 | 3 | 2 | 3 | 24 | 8 | 14 | 3 | 7 | 7 |
| 33016V3S1 | Fiji | Oceania | 2006 | skin/soft tissue infection, NOS | emm53.0 | 1033 | 46 | 32 | 7 | 1 | 6 | 7 | 73 | 79 | 1 | 16 | 56 | 31 |
| 33045V3S1 | Fiji | Oceania | 2006 | skin/soft tissue infection, NOS | emm105.0 | 954 | 6 | 6 | 6 | 6 | 5 | 6 | 6 | 12 | 5 | 4 | 6 | 11 |
| 33080V3S1 | Fiji | Oceania | 2006 | skin/soft tissue infection, NOS | emm82.1 | 320 | 56 | 70 | 4 | 1 | 20 | 2 | 89 | 98 | 38 | 2 | 65 | 3 |
| 33093V3S1 | Fiji | Oceania | 2006 | skin/soft tissue infection, NOS | emm71.0 | 318 | 19 | 20 | 13 | 19 | 2 | 22 | 3 | 27 | 1 | 1 | 88 | 24 |
| 33101V3S1 | Fiji | Oceania | 2006 | skin/soft tissue infection, NOS | emm65.4 | 129 | 11 | 66 | 40 | 35 | 102 | 76 | 44 | 145 | 1 | 2 | 87 | 31 |
| 33107V3S1 | Fiji | Oceania | 2006 | skin/soft tissue infection, NOS | emm89.14 | 380 | 14 | 49 | 20 | 16 | 112 | 52 | 136 | 80 | 33 | 11 | 57 | 25 |
| 31032V1S2 | Fiji | Oceania | 2006 | skin/soft tissue infection, NOS | emm25.0 | 1032 | 30 | 35 | 26 | 28 | 35 | 34 | 42 | 45 | 11 | 2 | 12 | 40 |
| 33115V3S1 | Fiji | Oceania | 2006 | skin/soft tissue infection, NOS | emm93.4 | 814 | 28 | 104 | 10 | 1 | 108 | 32 | 61 | 42 | 50 | 2 | 93 | 69 |
| 33119V3S1 | Fiji | Oceania | 2006 | skin/soft tissue infection, NOS | emm123.0 | 325 | 6 | 6 | 6 | 6 | 5 | 6 | 6 | 7 | 5 | 4 | 6 | 6 |
| 33126V3S1 | Fiji | Oceania | 2006 | skin/soft tissue infection, NOS | emm73.0 | 957 | 14 | 18 | 4 | 16 | 21 | 61 | 27 | 26 | 46 | 3 | 66 | 87 |
| 33132V3S1 | Fiji | Oceania | 2006 | skin/soft tissue infection, NOS | emm101.0 | 182 | 15 | 31 | 10 | 20 | 15 | 31 | 38 | 41 | 53 | 3 | 32 | 37 |
| 33139V3S1 | Fiji | Oceania | 2006 | skin/soft tissue infection, NOS | emm60.7 | 193 | 14 | 34 | 25 | 27 | 2 | 33 | 41 | 44 | 1 | 2 | 52 | 39 |
| 33139V3S2 | Fiji | Oceania | 2006 | skin/soft tissue infection, NOS | emm60.7 | 193 | 14 | 34 | 25 | 27 | 2 | 33 | 41 | 44 | 1 | 2 | 52 | 39 |
| 31158V1S1_01 | Fiji | Oceania | 2006 | skin/soft tissue infection, NOS | emm25.0 | 1032 | 30 | 35 | 26 | 28 | 35 | 34 | 42 | 45 | 11 | 2 | 12 | 40 |
| 31158V1S2 | Fiji | Oceania | 2006 | skin/soft tissue infection, NOS | emm25.0 | 1032 | 30 | 35 | 26 | 28 | 35 | 34 | 42 | 45 | 11 | 2 | 12 | 40 |
| 31158V3S1 | Fiji | Oceania | 2006 | skin/soft tissue infection, NOS | emm93.4 | 814 | 28 | 104 | 10 | 1 | 108 | 32 | 61 | 42 | 50 | 2 | 93 | 69 |
| 31158V3S2_01 | Fiji | Oceania | 2006 | skin/soft tissue infection, NOS | emm89.14 | 380 | 14 | 49 | 20 | 16 | 112 | 52 | 136 | 80 | 33 | 11 | 57 | 25 |
| 31158V4S1 | Fiji | Oceania | 2006 | skin/soft tissue infection, NOS | emm89.14 | 380 | 14 | 49 | 20 | 16 | 112 | 52 | 136 | 80 | 33 | 11 | 57 | 25 |
| 31158V5S1_01 | Fiji | Oceania | 2006 | skin/soft tissue infection, NOS | emm49.0 | 433 | 4 | 8 | 4 | 11 | 14 | 14 | 16 | 18 | 9 | 2 | 12 | 13 |
| 33181V1T1_01 | Fiji | Oceania | 2006 | pharyngitis and/or tonsillitis | emm137.0 | 268 | 1 | 60 | 49 | 45 | 61 | 19 | 13 | 83 | 34 | 3 | 61 | 29 |
| 33181V3S1 | Fiji | Oceania | 2006 | skin/soft tissue infection, NOS | emm218.1 | 292 | 38 | 45 | 3 | 16 | 44 | 44 | 53 | 61 | 54 | 2 | 12 | 3 |
| 33181V4T1 | Fiji | Oceania | 2006 | pharyngitis and/or tonsillitis | emm205.0 | 182 | 15 | 31 | 10 | 20 | 15 | 31 | 38 | 41 | 1 | 3 | 32 | 37 |

| | | | | | | | | | | | | | | | | | |
|---|---|---|---|---|---|---|---|---|---|---|---|---|---|---|---|---|---|
| 31086V2S2 | Fiji | Oceania | 2006 | skin/soft tissue infection, NOS | emm70.0 | 10 | 16 | 16 | 3 | 16 | 17 | 7 | 20 | 22 | 1 | 2 | 18 | 3 |
| 31142V2S1 | Fiji | Oceania | 2006 | skin/soft tissue infection, NOS | emm70.0 | 10 | 16 | 16 | 3 | 16 | 17 | 7 | 20 | 22 | 1 | 2 | 18 | 3 |
| 31144V2S1 | Fiji | Oceania | 2006 | skin/soft tissue infection, NOS | emm82.1 | 320 | 56 | 70 | 4 | 1 | 20 | 2 | 89 | 98 | 38 | 2 | 65 | 3 |
| 31152V2S1 | Fiji | Oceania | 2006 | skin/soft tissue infection, NOS | emm93.4 | 814 | 28 | 104 | 10 | 1 | 108 | 32 | 61 | 42 | 50 | 2 | 93 | 69 |
| 31154V2S1 | Fiji | Oceania | 2006 | skin/soft tissue infection, NOS | emm25.0 | 1032 | 30 | 35 | 26 | 28 | 35 | 34 | 42 | 45 | 11 | 2 | 12 | 40 |
| 31165V2S1 | Fiji | Oceania | 2006 | skin/soft tissue infection, NOS | emm93.4 | 814 | 28 | 104 | 10 | 1 | 108 | 32 | 61 | 42 | 50 | 2 | 93 | 69 |
| 31166V2S1 | Fiji | Oceania | 2006 | skin/soft tissue infection, NOS | emm82.1 | 320 | 56 | 70 | 4 | 1 | 20 | 2 | 89 | 98 | 38 | 2 | 65 | 3 |
| 31168V2S1 | Fiji | Oceania | 2006 | skin/soft tissue infection, NOS | emm70.0 | 10 | 16 | 16 | 3 | 16 | 17 | 7 | 20 | 22 | 1 | 2 | 18 | 3 |
| 31170V2S1 | Fiji | Oceania | 2006 | skin/soft tissue infection, NOS | emm82.1 | 320 | 56 | 70 | 4 | 1 | 20 | 2 | 89 | 98 | 38 | 2 | 65 | 3 |
| 31175V2S1 | Fiji | Oceania | 2006 | skin/soft tissue infection, NOS | emm70.0 | 10 | 16 | 16 | 3 | 16 | 17 | 7 | 20 | 22 | 1 | 2 | 18 | 3 |
| 31176V2S2 | Fiji | Oceania | 2006 | skin/soft tissue infection, NOS | emm103.0 | 327 | 14 | 13 | 3 | 1 | 66 | 13 | 82 | 87 | 1 | 2 | 49 | 4 |
| 31010V3S1 | Fiji | Oceania | 2006 | skin/soft tissue infection, NOS | emm123.0 | 325 | 6 | 6 | 6 | 6 | 5 | 6 | 6 | 7 | 5 | 4 | 6 | 6 |
| 31013V3S1 | Fiji | Oceania | 2006 | skin/soft tissue infection, NOS | emm82.1 | 320 | 56 | 70 | 4 | 1 | 20 | 2 | 89 | 98 | 38 | 2 | 65 | 3 |
| 31029V3S1 | Fiji | Oceania | 2006 | skin/soft tissue infection, NOS | emm57.0 | 1025 | 80 | 9 | 8 | 1 | 32 | 79 | 129 | 149 | 8 | 2 | 23 | 103 |
| 31048V3S1 | Fiji | Oceania | 2006 | skin/soft tissue infection, NOS | emm25.0 | 1032 | 30 | 35 | 26 | 28 | 35 | 34 | 42 | 45 | 11 | 2 | 12 | 40 |
| 31050V3S1 | Fiji | Oceania | 2006 | skin/soft tissue infection, NOS | emm103.0 | 327 | 14 | 13 | 3 | 1 | 66 | 13 | 82 | 87 | 1 | 2 | 49 | 4 |
| 31069V3S1 | Fiji | Oceania | 2006 | skin/soft tissue infection, NOS | emm4.5 | 1027 | 14 | 51 | 42 | 39 | 47 | 33 | 132 | 79 | 8 | 3 | 91 | 107 |
| 31089V2S1 | Fiji | Oceania | 2006 | skin/soft tissue infection, NOS | emm4.5 | 1027 | 14 | 51 | 42 | 39 | 47 | 33 | 132 | 79 | 8 | 3 | 91 | 107 |
| 31086V3S1 | Fiji | Oceania | 2006 | skin/soft tissue infection, NOS | emm104.0 | 789 | 14 | 13 | 10 | 8 | 13 | 13 | 15 | 17 | 10 | 2 | 15 | 16 |
| 31089V3S1 | Fiji | Oceania | 2006 | skin/soft tissue infection, NOS | emm70.0 | 10 | 16 | 16 | 3 | 16 | 17 | 7 | 20 | 22 | 1 | 2 | 18 | 3 |
| 31094V3S1 | Fiji | Oceania | 2006 | skin/soft tissue infection, NOS | emm82.1 | 320 | 56 | 70 | 4 | 1 | 20 | 2 | 89 | 98 | 38 | 2 | 65 | 3 |
| 31099V3S1 | Fiji | Oceania | 2006 | skin/soft tissue infection, NOS | emm123.0 | 325 | 6 | 6 | 6 | 6 | 5 | 6 | 6 | 7 | 5 | 4 | 6 | 6 |
| 31118V3S1 | Fiji | Oceania | 2006 | skin/soft tissue infection, NOS | emm82.1 | 320 | 56 | 70 | 4 | 1 | 20 | 2 | 89 | 98 | 38 | 2 | 65 | 3 |
| 31118V3S2 | Fiji | Oceania | 2006 | skin/soft tissue infection, NOS | emm74.0 | 120 | 28 | 13 | 75 | 41 | 50 | 53 | 123 | 82 | 11 | 4 | 59 | 70 |
| 31128V3S1 | Fiji | Oceania | 2006 | skin/soft tissue infection, NOS | emm42.0 | 80 | 37 | 32 | 37 | 35 | 43 | 43 | 44 | 60 | 2 | 2 | 39 | 52 |
| 31132V3S1 | Fiji | Oceania | 2006 | skin/soft tissue infection, NOS | emm65.4 | 129 | 11 | 66 | 40 | 35 | 68 | 76 | 44 | 145 | 1 | 2 | 87 | 31 |
| 31143V3S1 | Fiji | Oceania | 2006 | skin/soft tissue infection, NOS | emm89.14 | 380 | 14 | 49 | 20 | 16 | 112 | 52 | 136 | 80 | 33 | 11 | 57 | 25 |
| 31104V2S1 | Fiji | Oceania | 2006 | skin/soft tissue infection, NOS | emm4.5 | 1027 | 14 | 51 | 42 | 39 | 47 | 33 | 132 | 79 | 8 | 3 | 91 | 107 |
| 31147V3S1 | Fiji | Oceania | 2006 | skin/soft tissue infection, NOS | emm56.0 | 115 | 14 | 44 | 24 | 2 | 12 | 41 | 51 | 153 | 23 | 1 | 6 | 49 |
| 31158V3S1a | Fiji | Oceania | 2006 | skin/soft tissue infection, NOS | emm93.4 | 814 | 28 | 104 | 10 | 1 | 108 | 32 | 61 | 42 | 50 | 2 | 93 | 69 |
| 31158V3S1b | Fiji | Oceania | 2006 | skin/soft tissue infection, NOS | emm89.14 | 380 | 14 | 49 | 20 | 16 | 112 | 52 | 136 | 80 | 33 | 11 | 57 | 25 |
| 31163V3S1 | Fiji | Oceania | 2006 | skin/soft tissue infection, NOS | emm97.1 | 216 | 21 | 21 | 15 | 8 | 46 | 24 | 26 | 30 | 8 | 10 | 12 | 26 |
| 31170V3S1 | Fiji | Oceania | 2006 | skin/soft tissue infection, NOS | emm73.0 | 957 | 14 | 18 | 4 | 16 | 21 | 61 | 27 | 26 | 46 | 3 | 66 | 87 |
| 31171V3S1 | Fiji | Oceania | 2006 | skin/soft tissue infection, NOS | emm93.4 | 814 | 28 | 104 | 10 | 1 | 108 | 32 | 61 | 42 | 50 | 2 | 93 | 69 |
| 31177V3S1 | Fiji | Oceania | 2006 | skin/soft tissue infection, NOS | emm123.0 | 325 | 6 | 6 | 6 | 6 | 5 | 6 | 6 | 7 | 5 | 4 | 6 | 6 |
| 31004V4S1 | Fiji | Oceania | 2006 | skin/soft tissue infection, NOS | emm93.4 | 814 | 28 | 104 | 10 | 1 | 108 | 32 | 61 | 42 | 50 | 2 | 93 | 69 |
| 31009V4S1 | Fiji | Oceania | 2006 | skin/soft tissue infection, NOS | emm93.4 | 814 | 28 | 104 | 10 | 1 | 108 | 32 | 61 | 42 | 50 | 2 | 93 | 69 |
| 31032V4S1 | Fiji | Oceania | 2006 | skin/soft tissue infection, NOS | emm104.0 | 789 | 14 | 13 | 10 | 8 | 13 | 13 | 15 | 17 | 10 | 2 | 15 | 16 |
| 31034V4S1 | Fiji | Oceania | 2006 | skin/soft tissue infection, NOS | emm57.0 | 1025 | 80 | 9 | 8 | 1 | 32 | 79 | 129 | 149 | 8 | 2 | 23 | 103 |
| 31037V4S1 | Fiji | Oceania | 2006 | skin/soft tissue infection, NOS | emm70.0 | 10 | 16 | 16 | 3 | 16 | 17 | 7 | 20 | 22 | 1 | 2 | 18 | 3 |
| 31044V4S1 | Fiji | Oceania | 2006 | skin/soft tissue infection, NOS | emm39.4 | 268 | 1 | 60 | 49 | 45 | 61 | 19 | 13 | 83 | 34 | 3 | 61 | 29 |
| 31053V4S1 | Fiji | Oceania | 2006 | skin/soft tissue infection, NOS | emm93.4 | 814 | 28 | 104 | 10 | 1 | 108 | 32 | 61 | 42 | 50 | 2 | 93 | 69 |
| 31058V4S1 | Fiji | Oceania | 2006 | skin/soft tissue infection, NOS | emm103.0 | 327 | 14 | 13 | 3 | 1 | 66 | 13 | 82 | 87 | 1 | 2 | 49 | 4 |
| 31059V4S1 | Fiji | Oceania | 2006 | skin/soft tissue infection, NOS | emm93.4 | 814 | 28 | 104 | 10 | 1 | 108 | 32 | 61 | 42 | 50 | 2 | 93 | 69 |
| 31063V4S1 | Fiji | Oceania | 2006 | skin/soft tissue infection, NOS | emm70.0 | 10 | 16 | 16 | 3 | 16 | 17 | 7 | 20 | 22 | 1 | 2 | 18 | 3 |
| 31075V4S1 | Fiji | Oceania | 2006 | skin/soft tissue infection, NOS | emm11.0 | ND | 1 | 6 | 76 | 1 | 20 | 20 | 13 | 25 | 8 | 3 | 21 | 23 |
| 31077V4S1 | Fiji | Oceania | 2006 | skin/soft tissue infection, NOS | emm65.4 | 129 | 11 | 66 | 40 | 35 | 68 | 76 | 44 | 145 | 1 | 2 | 87 | 31 |
| 31107V2S1 | Fiji | Oceania | 2006 | skin/soft tissue infection, NOS | emm105.0 | 954 | 6 | 6 | 6 | 6 | 5 | 6 | 6 | 12 | 5 | 4 | 6 | 11 |

| | | | | | | | | | | | | | | | | | | |
|---|---|---|---|---|---|---|---|---|---|---|---|---|---|---|---|---|---|---|
| 31081V4S1 | Fiji | Oceania | 2006 | skin/soft tissue infection, NOS | emm65.4 | 129 | 11 | 66 | 40 | 35 | 68 | 76 | 44 | 145 | 1 | 2 | 87 | 31 |
| 31088V4S1 | Fiji | Oceania | 2006 | skin/soft tissue infection, NOS | emm39.4 | 268 | 1 | 60 | 49 | 45 | 61 | 19 | 13 | 83 | 34 | 3 | 61 | 29 |
| 31091V4S1 | Fiji | Oceania | 2006 | skin/soft tissue infection, NOS | emm97.1 | 216 | 21 | 21 | 15 | 8 | 46 | 24 | 26 | 30 | 8 | 10 | 12 | 26 |
| 31101V4S1 | Fiji | Oceania | 2006 | skin/soft tissue infection, NOS | emm89.14 | 380 | 14 | 49 | 20 | 16 | 112 | 52 | 136 | 80 | 33 | 11 | 57 | 25 |
| 31109V4S1 | Fiji | Oceania | 2006 | skin/soft tissue infection, NOS | emm93.4 | 814 | 28 | 104 | 10 | 1 | 108 | 32 | 61 | 42 | 50 | 2 | 93 | 69 |
| 31114V4S1 | Fiji | Oceania | 2006 | skin/soft tissue infection, NOS | emm93.4 | 814 | 28 | 104 | 10 | 1 | 108 | 32 | 61 | 42 | 50 | 2 | 93 | 69 |
| 31117V4S1 | Fiji | Oceania | 2006 | skin/soft tissue infection, NOS | emm89.14 | 380 | 14 | 49 | 20 | 16 | 112 | 52 | 136 | 80 | 33 | 11 | 57 | 25 |
| 31119V4S1 | Fiji | Oceania | 2006 | skin/soft tissue infection, NOS | emm89.14 | 380 | 14 | 49 | 20 | 16 | 112 | 52 | 136 | 80 | 33 | 11 | 57 | 25 |
| 31118V2S1 | Fiji | Oceania | 2006 | skin/soft tissue infection, NOS | emm65.4 | 129 | 11 | 66 | 40 | 35 | 68 | 76 | 44 | 145 | 1 | 2 | 87 | 31 |
| 31121V4S1 | Fiji | Oceania | 2006 | skin/soft tissue infection, NOS | emm39.4 | 268 | 1 | 60 | 49 | 45 | 61 | 19 | 13 | 83 | 34 | 3 | 61 | 29 |
| 31124V4S1 | Fiji | Oceania | 2006 | skin/soft tissue infection, NOS | emm65.4 | 129 | 11 | 66 | 40 | 35 | 68 | 76 | 44 | 145 | 1 | 2 | 87 | 31 |
| 31125V4S1 | Fiji | Oceania | 2006 | skin/soft tissue infection, NOS | emm65.4 | 129 | 11 | 66 | 40 | 35 | 68 | 76 | 44 | 145 | 1 | 2 | 87 | 31 |
| 31128V4S1 | Fiji | Oceania | 2006 | skin/soft tissue infection, NOS | emm11.0 | ND | 1 | 6 | 76 | 1 | 20 | 20 | 13 | 25 | 8 | 3 | 21 | 23 |
| 31129V4S1 | Fiji | Oceania | 2006 | skin/soft tissue infection, NOS | emm39.4 | 268 | 1 | 60 | 49 | 45 | 61 | 19 | 13 | 83 | 34 | 3 | 61 | 29 |
| 31134V4S1 | Fiji | Oceania | 2006 | skin/soft tissue infection, NOS | emm39.4 | 268 | 1 | 60 | 49 | 45 | 61 | 19 | 13 | 83 | 34 | 3 | 61 | 29 |
| 31136V4S1 | Fiji | Oceania | 2006 | skin/soft tissue infection, NOS | emm49.0 | 433 | 4 | 8 | 4 | 11 | 14 | 14 | 16 | 18 | 9 | 2 | 12 | 13 |
| 31148V4S1 | Fiji | Oceania | 2006 | skin/soft tissue infection, NOS | emm70.0 | 10 | 16 | 16 | 3 | 16 | 17 | 7 | 20 | 22 | 1 | 2 | 18 | 3 |
| 31165V4S1 | Fiji | Oceania | 2006 | skin/soft tissue infection, NOS | emm56.0 | 115 | 14 | 44 | 24 | 2 | 12 | 41 | 51 | 153 | 23 | 1 | 6 | 49 |
| 31167V4S1 | Fiji | Oceania | 2006 | skin/soft tissue infection, NOS | emm11.0 | ND | 1 | 6 | 76 | 1 | 104 | 20 | 13 | 25 | 8 | 3 | 21 | 23 |
| 31176V4S1 | Fiji | Oceania | 2006 | skin/soft tissue infection, NOS | emm76.4 | 988 | 45 | 34 | 48 | 8 | 58 | 50 | 71 | 57 | 18 | 3 | 22 | 102 |
| 33160V2T1 | Fiji | Oceania | 2006 | pharyngitis and/or tonsillitis | emm92.0 | 1026 | 5 | 65 | 34 | 27 | 39 | 39 | 22 | 152 | 13 | 2 | 20 | 22 |
| 33157V2T1 | Fiji | Oceania | 2006 | pharyngitis and/or tonsillitis | emm101.0 | 182 | 15 | 31 | 10 | 20 | 15 | 31 | 38 | 41 | 1 | 3 | 32 | 37 |
| 33141V2T1 | Fiji | Oceania | 2006 | pharyngitis and/or tonsillitis | emm25.1 | 192 | 14 | 101 | 24 | 2 | 33 | 13 | 13 | 40 | 36 | 3 | 31 | 36 |
| 33098V2T1 | Fiji | Oceania | 2006 | pharyngitis and/or tonsillitis | emm44.0 | 178 | 5 | 7 | 7 | 3 | 2 | 3 | 24 | 8 | 14 | 3 | 7 | 7 |
| 33181V3T1 | Fiji | Oceania | 2006 | pharyngitis and/or tonsillitis | emm101.0 | 182 | 15 | 31 | 10 | 20 | 15 | 31 | 38 | 41 | 1 | 3 | 32 | 37 |
| 33133V3T1 | Fiji | Oceania | 2006 | pharyngitis and/or tonsillitis | emm238.2 | 867 | 1 | 1 | 1 | 1 | 1 | 15 | 14 | 93 | 5 | 6 | 14 | 15 |
| 33092V2T1 | Fiji | Oceania | 2006 | pharyngitis and/or tonsillitis | emm49.0 | 433 | 4 | 8 | 4 | 11 | 14 | 14 | 16 | 18 | 9 | 2 | 12 | 13 |
| 33141V4T1 | Fiji | Oceania | 2006 | pharyngitis and/or tonsillitis | emm238.2 | 867 | 1 | 1 | 1 | 1 | 1 | 15 | 14 | 93 | 5 | 6 | 14 | 15 |
| 33138V3T1 | Fiji | Oceania | 2006 | pharyngitis and/or tonsillitis | emm122.2 | 1029 | 15 | 15 | 10 | 1 | 110 | 7 | 134 | 22 | 23 | 1 | 6 | 108 |
| 31124V2S1 | Fiji | Oceania | 2006 | skin/soft tissue infection, NOS | emm71.0 | 318 | 19 | 20 | 13 | 19 | 2 | 22 | 3 | 27 | 1 | 1 | 4 | 24 |
| 33124V2T1 | Fiji | Oceania | 2006 | pharyngitis and/or tonsillitis | emm54.1 | 990 | 11 | 5 | 10 | 20 | 15 | 16 | 17 | 13 | 1 | 2 | 16 | 31 |
| 31027V4S1 | Fiji | Oceania | 2006 | skin/soft tissue infection, NOS | emm57.0 | 1025 | 80 | 9 | 8 | 1 | 32 | 79 | 129 | 149 | 8 | 2 | 23 | 103 |
| 33181V5T1 | Fiji | Oceania | 2006 | pharyngitis and/or tonsillitis | emm101.0 | 182 | 15 | 31 | 10 | 20 | 15 | 31 | 38 | 41 | 1 | 3 | 32 | 37 |
| 33181V6T1 | Fiji | Oceania | 2006 | pharyngitis and/or tonsillitis | emm92.0 | 1026 | 5 | 65 | 34 | 27 | 39 | 39 | 22 | 152 | 13 | 2 | 20 | 22 |
| M28PF1 | France | Europe | ND | puerperal fever | emm28.0 | 52 | 3 | 3 | 3 | 3 | 2 | 3 | 3 | 4 | 1 | 1 | 3 | 3 |
| STAB09014 | France | Europe | 2015 | skin/soft tissue infection, NOS | emm28.0 | 52 | 3 | 3 | 3 | 3 | 2 | 3 | 3 | 4 | 1 | 1 | 3 | 3 |
| STAB090229 | France | Europe | 2009 | sepsis | emm75.0 | 150 | 25 | 26 | 20 | 24 | 29 | 29 | 33 | 48 | 15 | 3 | 12 | 33 |
| STAB10015 | France | Europe | ND | skin/soft tissue infection, NOS | emm28.0 | 52 | 3 | 3 | 3 | 3 | 2 | 3 | 3 | 4 | 1 | 1 | 3 | 3 |
| 7F7 | France | Europe | 2011 | ND | emm83.1 | 5 | 11 | 5 | 10 | 8 | 15 | 65 | 97 | 115 | 11 | 2 | 69 | 83 |
| STAB13021 | France | Europe | 2013 | abscess | emm66.0 | 44 | 14 | 82 | 64 | 31 | 6 | 18 | 127 | 77 | 1 | 1 | 73 | 3 |
| STAB14018 | France | Europe | 2014 | sepsis | emm75.0 | 150 | 25 | 26 | 20 | 24 | 29 | 29 | 33 | 155 | 15 | 3 | 12 | 33 |
| STAB901 | France | Europe | 2009 | invasive, NOS | emm44.0 | 178 | 5 | 7 | 7 | 1 | 6 | 7 | 7 | [S] | [S] | 3 | 7 | 7 |
| STAB902 | France | Europe | ND | ND | emm3.2 | 406 | 15 | 15 | 10 | 1 | 10 | 32 | 111 | 129 | 1 | 2 | 6 | 31 |
| STAB1102 | France | Europe | ND | ND | emm83.1 | 5 | 11 | 5 | 10 | 8 | 15 | 65 | 97 | 115 | 11 | 2 | 69 | 83 |
| 1E1 | France | Europe | 2009 | ND | emm44.0 | 178 | 5 | 7 | 7 | 1 | 6 | 7 | 7 | 8 | 14 | 3 | 7 | 7 |
| HKU30 | China [Hong Kong] | Asia | 2011 | scarlet fever | emm12.0 | 36 | 14 | 3 | 3 | 3 | 2 | 3 | 52 | 59 | 24 | 12 | 38 | 3 |
| HKU22 | China [Hong Kong] | Asia | 2011 | ND | emm12.0 | 36 | 14 | 3 | 3 | 3 | 2 | 3 | 52 | 59 | 24 | 12 | 38 | 3 |
| HKU165 | China [Hong Kong] | Asia | 2005 | ND | emm12.0 | 36 | 14 | 3 | 3 | 3 | 2 | 3 | 52 | 59 | 24 | 12 | 94 | 3 |

| | | | | | | | | | | | | | | | | | | |
|---|---|---|---|---|---|---|---|---|---|---|---|---|---|---|---|---|---|---|
| HKU306 | China [Hong Kong] | Asia | 2011 | scarlet fever | emm12.0 | 36 | 14 | 3 | 3 | 3 | 2 | 3 | 52 | 59 | 24 | 12 | 38 | 3 |
| HKU360 | China [Hong Kong] | Asia | 2011 | scarlet fever | emm12.0 | 36 | 14 | 3 | 3 | 3 | 113 | 3 | 52 | 59 | 24 | 12 | 38 | 3 |
| HKU383 | China [Hong Kong] | Asia | 2011 | scarlet fever | emm12.0 | 36 | 14 | 3 | 3 | 3 | 2 | 3 | 52 | 59 | 24 | 12 | 38 | 3 |
| HKU388 | China [Hong Kong] | Asia | 2011 | scarlet fever | emm12.0 | 36 | 14 | 3 | 3 | 3 | 2 | 3 | 52 | 59 | 24 | 12 | 38 | 3 |
| HKU364 | China [Hong Kong] | Asia | 2008 | ND | emm12.0 | 36 | 14 | 3 | 3 | 3 | 2 | 3 | 52 | 59 | 24 | 12 | 38 | 3 |
| HKU397 | China [Hong Kong] | Asia | 2011 | scarlet fever | emm12.0 | 36 | 14 | 3 | 3 | 3 | 2 | 3 | 52 | 59 | 24 | 12 | 38 | 3 |
| HKU434 | China [Hong Kong] | Asia | 2011 | scarlet fever | emm1.0 | 28 | 1 | 1 | 1 | 1 | 1 | 1 | 1 | 156 | 1 | 1 | 1 | 1 |
| HKU464 | China [Hong Kong] | Asia | 2012 | pneumonia | emm1.0 | 28 | 1 | 1 | 1 | 1 | 1 | 1 | 1 | 1 | 1 | 1 | 1 | 1 |
| HKU474 | China [Hong Kong] | Asia | 2012 | scarlet fever | emm1.0 | 28 | 1 | 1 | 1 | 1 | 1 | 1 | 1 | 156 | 1 | 1 | 1 | 1 |
| HKU484 | China [Hong Kong] | Asia | 2012 | scarlet fever | emm1.0 | 28 | 1 | 1 | 1 | 1 | 114 | 1 | 1 | 1 | 1 | 1 | 1 | 1 |
| HKU486 | China [Hong Kong] | Asia | 2011 | ND | emm1.0 | 28 | 1 | 1 | 1 | 65 | 1 | 1 | 1 | 1 | 1 | 1 | 1 | 1 |
| HKU487 | China [Hong Kong] | Asia | 2011 | scarlet fever | emm1.0 | 28 | 1 | 1 | 1 | 1 | 1 | 1 | 1 | 1 | 1 | 1 | 1 | 1 |
| HKU488 | China [Hong Kong] | Asia | 2012 | scarlet fever | emm1.0 | 28 | 1 | 1 | 1 | 1 | 1 | 1 | 1 | 1 | 1 | 1 | 1 | 1 |
| A1268 | India | Asia | 2009 | invasive, NOS | emm1.0 | 28 | 1 | 1 | 1 | 1 | 1 | 1 | 1 | 1 | 1 | 1 | 1 | 1 |
| A1287 | India | Asia | 2009 | invasive, NOS | emm49.0 | 371 | 4 | 8 | 77 | 11 | 14 | 14 | 16 | 18 | 9 | 2 | 12 | 13 |
| A1267 | India | Asia | 2009 | invasive, NOS | emm43.3 | 1001 | 14 | 49 | 33 | 19 | 2 | 22 | 137 | 157 | 56 | 4 | 95 | 109 |
| A1152 | India | Asia | 2009 | invasive, NOS | emm12.0 | 36 | 14 | 3 | 3 | 3 | 2 | 3 | 52 | 59 | 24 | 12 | 38 | 3 |
| A1151 | India | Asia | 2009 | invasive, NOS | emm104.0 | 353 | 45 | 34 | 48 | 8 | 58 | 50 | 71 | 57 | 18 | 3 | 22 | 102 |
| A1144 | India | Asia | 2009 | pharyngitis and/or tonsillitis | emm12.40 | 36 | 14 | 3 | 3 | 3 | 2 | 3 | 52 | 59 | 24 | 12 | 38 | 3 |
| A1138 | India | Asia | 2009 | pharyngitis and/or tonsillitis | emm12.40 | 36 | 14 | 3 | 3 | 3 | 2 | 3 | 52 | 59 | 24 | 12 | 38 | 3 |
| A1013 | India | Asia | 2009 | ND | emm82.1 | 320 | 56 | 70 | 4 | 1 | 20 | 2 | 89 | 158 | 38 | 2 | 65 | 3 |
| A998 | India | Asia | 2009 | ND | emm80.0 | 701 | 16 | 16 | 3 | 16 | 17 | 7 | 20 | 22 | 1 | 2 | 18 | 110 |
| A997 | India | Asia | 2009 | ND | emm75.3 | 230 | 10 | 106 | 45 | 6 | 55 | 2 | 138 | 74 | 1 | 2 | 30 | 31 |
| A996 | India | Asia | 2009 | ND | emm44.0 | 987 | 5 | 7 | 7 | 1 | 6 | 7 | 7 | 8 | 14 | 3 | 7 | 7 |
| A995 | India | Asia | 2009 | ND | emm22.8 | 360 | 14 | 51 | 42 | 39 | 115 | 81 | 139 | 159 | 57 | 2 | 96 | 111 |
| A993 | India | Asia | 2009 | ND | emm80.0 | 715 | 16 | 16 | 3 | 16 | 17 | 82 | 20 | 22 | 1 | 2 | 18 | 3 |
| A982 | India | Asia | 2009 | ND | emm80.0 | 701 | 16 | 16 | 3 | 16 | 116 | 7 | 20 | 22 | 1 | 2 | 18 | 110 |
| A981 | India | Asia | 2009 | ND | emm102.2 | 349 | 14 | 37 | 31 | 3 | 55 | 38 | 47 | 81 | 1 | 2 | 58 | 25 |
| A980 | India | Asia | 2009 | ND | emm82.1 | 320 | 56 | 70 | 4 | 1 | 20 | 2 | 89 | 158 | 38 | 2 | 65 | 3 |
| A969 | India | Asia | 2009 | ND | emm100.6 | 119 | 11 | 11 | 7 | 10 | 10 | 7 | 37 | 39 | 19 | 2 | 30 | 31 |
| A967 | India | Asia | 2009 | ND | emm44.0 | 178 | 5 | 7 | 7 | 1 | 6 | 7 | 7 | 8 | 14 | 3 | 7 | 7 |
| A960 | India | Asia | 2009 | ND | emm168.1 | 358 | 24 | 19 | 78 | 23 | 117 | 83 | 45 | 160 | 58 | 3 | 97 | 112 |
| A920 | India | Asia | 2009 | ND | emm112.2 | 497 | 18 | 107 | 4 | 66 | 118 | 61 | 27 | 161 | 8 | 3 | 21 | 113 |
| A915 | India | Asia | 2009 | ND | emm49.4 | 534 | 14 | 108 | 79 | 22 | 27 | 13 | 26 | 29 | 8 | 3 | 25 | 114 |
| A914 | India | Asia | 2009 | ND | emm74.0 | 120 | 28 | 13 | 3 | 41 | 50 | 53 | 123 | 82 | 11 | 4 | 59 | 70 |
| A906 | India | Asia | 2009 | ND | emm104.0 | 353 | 45 | 34 | 48 | 8 | 58 | 50 | 71 | 57 | 18 | 3 | 22 | 102 |
| A905 | India | Asia | 2009 | ND | emm82.1 | 320 | 56 | 70 | 4 | 1 | 20 | 2 | 89 | 158 | 38 | 2 | 65 | 3 |
| A904 | India | Asia | 2009 | ND | emm113.0 | 499 | 18 | 18 | 4 | 4 | 119 | 7 | 25 | 57 | 18 | 3 | 35 | 115 |
| A900 | India | Asia | 2009 | ND | emm110.0 | 495 | 14 | 33 | 18 | 6 | 25 | 27 | 40 | 43 | 1 | 2 | 18 | 28 |
| A896 | India | Asia | 2009 | ND | emm18.12 | 535 | 83 | 57 | 2 | 2 | 2 | 2 | 67 | 75 | 8 | 3 | 53 | 2 |
| A895 | India | Asia | 2009 | ND | emm100.6 | 119 | 11 | 11 | 7 | 10 | 10 | 7 | 37 | 39 | 19 | 2 | 30 | 31 |
| A879 | India | Asia | 2009 | ND | emm12.21 | 991 | 14 | 3 | 3 | 3 | 2 | 3 | 52 | 59 | 24 | 12 | 38 | 3 |
| A876 | India | Asia | 2009 | ND | emm18.12 | 535 | 83 | 57 | 2 | 2 | 2 | 2 | 67 | 75 | 8 | 3 | 53 | 2 |
| A873 | India | Asia | 2009 | ND | emm3.23 | 15 | 15 | 15 | 10 | 1 | 10 | 32 | 111 | 129 | 1 | 2 | 6 | 31 |
| A870 | India | Asia | 2009 | ND | emm168.1 | 358 | 24 | 19 | 78 | 23 | 117 | 83 | 45 | 160 | 58 | 3 | 97 | 112 |
| A868 | India | Asia | 2009 | ND | emm108.0 | 304 | 11 | 109 | 10 | 20 | 15 | 16 | 17 | 13 | 59 | 2 | 16 | 31 |
| A856 | India | Asia | 2009 | ND | emm3.23 | 15 | 15 | 15 | 10 | 1 | 10 | 32 | 111 | 129 | 1 | 2 | 6 | 31 |
| A855 | India | Asia | 2009 | ND | emm108.0 | 304 | 11 | 109 | 10 | 20 | 15 | 16 | 17 | 13 | 59 | 2 | 16 | 31 |

| | | | | | | | | | | | | | | | | | | |
|---|---|---|---|---|---|---|---|---|---|---|---|---|---|---|---|---|---|---|
| A848 | India | Asia | 2009 | pharyngitis and/or tonsillitis | emm63.3 | 297 | 24 | 19 | 3 | 23 | 120 | 7 | 32 | 34 | 17 | 11 | 26 | 32 |
| A846 | India | Asia | 2009 | ND | emm28.5 | 473 | 38 | 45 | 3 | 16 | 44 | 19 | 140 | 162 | 60 | 3 | 12 | 30 |
| A843 | India | Asia | 2009 | pharyngitis and/or tonsillitis | emm3.22 | 1002 | 9 | 9 | 80 | 8 | 8 | 10 | 9 | 163 | 5 | 5 | 98 | 9 |
| A842 | India | Asia | 2009 | ND | emm3.22 | 1002 | 9 | 9 | 80 | 8 | 8 | 10 | 9 | 163 | 5 | 5 | 98 | 9 |
| A841 | India | Asia | 2009 | ND | emm77.0 | 482 | 14 | 110 | 3 | 22 | 27 | 13 | 26 | 29 | 8 | 3 | 25 | 30 |
| A840 | India | Asia | 2009 | ND | emm77.0 | 482 | 14 | 110 | 3 | 22 | 27 | 13 | 26 | 29 | 8 | 3 | 25 | 30 |
| A839 | India | Asia | 2009 | pharyngitis and/or tonsillitis | emm75.1 | 357 | 14 | 34 | 55 | 50 | 71 | 45 | 35 | 37 | 18 | 3 | 28 | 30 |
| A838 | India | Asia | 2009 | ND | emm49.4 | 534 | 14 | 25 | 79 | 22 | 27 | 13 | 26 | 29 | 8 | 3 | 25 | 114 |
| A837 | India | Asia | 2009 | ND | emm82.1 | 320 | 56 | 70 | 4 | 1 | 20 | 2 | 89 | 158 | 38 | 2 | 65 | 3 |
| A829 | India | Asia | 2009 | pharyngitis and/or tonsillitis | emm28.5 | 473 | 38 | 45 | 3 | 16 | 44 | 19 | 140 | 162 | 60 | 3 | 12 | 30 |
| JS12 | Israel | Asia | 1997 | ND | emm179.0 | 619 | 21 | 111 | 15 | 8 | 46 | 24 | 26 | 30 | 8 | 10 | 12 | 26 |
| MGAS23530 | Italy | Europe | ND | pharyngitis and/or tonsillitis | emm89.0 | 101 | 2 | 2 | 2 | 2 | 2 | 2 | 2 | 2 | 2 | 2 | 2 | 2 |
| JMUB1235 | Japan | Asia | 2016 | ND | emm89.0 | 101 | 2 | 2 | 2 | 2 | 121 | 2 | 2 | 2 | 2 | 2 | 2 | 2 |
| M1_476 | Japan | Asia | 1994 | invasive, NOS | emm1.0 | 28 | 1 | 1 | 1 | 1 | 1 | 1 | 1 | 1 | 1 | 1 | [S] | 1 |
| MTB313 | Japan | Asia | ND | meningitis | emm1.0 | 28 | 1 | 1 | 1 | 1 | 1 | 1 | 141 | [S] | [S] | 1 | 1 | 1 |
| MTB314 | Japan | Asia | ND | meningitis | emm1.0 | 28 | 1 | 1 | 1 | 1 | 1 | 1 | 141 | 1 | 1 | 1 | 1 | [S] |
| SSI-1 | Japan | Asia | 1994 | invasive, NOS | emm3.1 | 15 | 15 | 15 | 10 | 1 | 10 | 32 | 111 | 129 | 1 | 2 | 6 | 31 |
| K3534 | Kenya | Africa | 1998 | skin/soft tissue infection, NOS | emm65.0 | 716 | 14 | 34 | 81 | 27 | 122 | 13 | 142 | 164 | 15 | 3 | 100 | 117 |
| K5499 | Kenya | Africa | 2000 | skin/soft tissue infection, NOS | emm65.5 | 215 | 1 | 112 | 24 | 1 | 2 | 13 | 143 | 165 | 8 | 3 | 101 | 118 |
| K5721 | Kenya | Africa | 2000 | skin/soft tissue infection, NOS | emm26.3 | 745 | 14 | 6 | 82 | 67 | 123 | 84 | 144 | 166 | 8 | 3 | 102 | 119 |
| K5911 | Kenya | Africa | 2000 | invasive, NOS | emm217.0 | 728 | 14 | 37 | 20 | 24 | 64 | 57 | 145 | 167 | 15 | 3 | 18 | 89 |
| K6239 | Kenya | Africa | 2000 | pneumonia | emm9.0 | 740 | 45 | 34 | 40 | 8 | 124 | 19 | 45 | 168 | 62 | 3 | 12 | 120 |
| K6429 | Kenya | Africa | 2000 | skin/soft tissue infection, NOS | emm38.1 | 757 | 14 | 37 | 20 | 24 | 125 | 13 | 146 | 23 | 5 | 25 | 103 | 117 |
| K6821 | Kenya | Africa | 2001 | skin/soft tissue infection, NOS | emm177.0 | 758 | 84 | 113 | 21 | 23 | 126 | 13 | 147 | 169 | 1 | 2 | 104 | 121 |
| K7367 | Kenya | Africa | 2001 | skin/soft tissue infection, NOS | emm55.0 | 248 | 11 | 11 | 83 | 10 | 10 | 13 | 148 | 13 | 8 | 3 | 11 | 12 |
| K9340 | Kenya | Africa | 2002 | skin/soft tissue infection, NOS | emm18.21 | 221 | 11 | 114 | 84 | 16 | 82 | 13 | 20 | 170 | 63 | 3 | 13 | 30 |
| K9404 | Kenya | Africa | 2002 | invasive, NOS | emm90.5 | 708 | 14 | 115 | 85 | 2 | 127 | 13 | 149 | 171 | 8 | 3 | 12 | 122 |
| K9429 | Kenya | Africa | 2002 | pneumonia | emm95.0 | 712 | 85 | 6 | 6 | 6 | 5 | 6 | 6 | 172 | 5 | 4 | 6 | 11 |
| K9466 | Kenya | Africa | 2002 | pneumonia | emm78.5 | 255 | 86 | 34 | 40 | 8 | 128 | 19 | 45 | 173 | 62 | 3 | 12 | 120 |
| K12363 | Kenya | Africa | 2004 | skin/soft tissue infection, NOS | emm79.5 | 714 | 14 | 25 | 3 | 22 | 27 | 13 | 26 | 29 | 8 | 3 | 25 | 30 |
| K12452 | Kenya | Africa | 2004 | skin/soft tissue infection, NOS | emm183.2 | 761 | 14 | 25 | 3 | 22 | 27 | 13 | 26 | 174 | 8 | 3 | 25 | 30 |
| K13569 | Kenya | Africa | 2004 | invasive, NOS | emm63.5 | 764 | 14 | 116 | 86 | 27 | 129 | 9 | 150 | 175 | 64 | 2 | 12 | 123 |
| K16587 | Kenya | Africa | 2005 | skin/soft tissue infection, NOS | ND | 450 | 87 | 117 | 3 | 1 | 26 | 13 | 20 | 176 | 65 | 3 | 105 | 124 |
| K16849 | Kenya | Africa | 2005 | invasive, NOS | emm83.12 | 393 | 88 | 45 | 3 | 16 | 130 | 85 | 151 | 177 | 60 | 3 | 12 | 30 |
| K19464 | Kenya | Africa | 2006 | skin/soft tissue infection, NOS | ND | 254 | 6 | 118 | 71 | 1 | 131 | 13 | 152 | 178 | 8 | 3 | 106 | 125 |
| K24357 | Kenya | Africa | 2007 | skin/soft tissue infection, NOS | emm209.0 | 260 | 14 | 37 | 3 | 27 | 132 | 13 | 153 | 179 | 34 | 2 | 107 | 126 |
| K39244 | Kenya | Africa | 2009 | pneumonia | emm80.0 | 715 | 16 | 16 | 3 | 16 | 133 | 7 | 20 | 22 | 1 | 2 | 18 | 3 |
| K47483 | Kenya | Africa | 2011 | skin/soft tissue infection, NOS | emm225.0 | 262 | 14 | 44 | 24 | 2 | 12 | 41 | 154 | 180 | 8 | 3 | 101 | 127 |
| K51725 | Kenya | Africa | 2011 | skin/soft tissue infection, NOS | emm18.21 | 402 | 11 | 13 | 3 | 41 | 50 | 13 | 112 | 42 | 31 | 1 | 51 | 4 |
| K9374 | Kenya | Africa | 2002 | skin/soft tissue infection, NOS | emm18.21 | 221 | 11 | 114 | 84 | 16 | 82 | 13 | 20 | 170 | 63 | 3 | 13 | 30 |
| K9400 | Kenya | Africa | 2002 | skin/soft tissue infection, NOS | emm90.5 | 708 | 14 | 115 | 85 | 2 | 127 | 13 | 149 | 171 | 8 | 3 | 12 | 122 |
| K9887 | Kenya | Africa | 2003 | skin/soft tissue infection, NOS | emm119.2 | 239 | 38 | 45 | 3 | 16 | 44 | 44 | 53 | 61 | 54 | 2 | 12 | 3 |
| K9927 | Kenya | Africa | 2003 | skin/soft tissue infection, NOS | emm223.0 | 613 | 85 | 6 | 6 | 6 | 5 | 6 | 6 | 12 | 5 | 4 | 6 | 11 |
| K10234 | Kenya | Africa | 2003 | skin/soft tissue infection, NOS | emm65.5 | 215 | 1 | 112 | 24 | 1 | 2 | 13 | 143 | 165 | 8 | 3 | 101 | 118 |
| K10246 | Kenya | Africa | 2003 | pneumonia | emm162.1 | 412 | 14 | 119 | 24 | 1 | 134 | 1 | 121 | 140 | 5 | 4 | 95 | 128 |
| K13254 | Kenya | Africa | 2004 | skin/soft tissue infection, NOS | emm63.5 | 764 | 14 | 116 | 86 | 27 | 129 | 9 | 150 | 175 | 64 | 2 | 12 | 123 |
| K17011 | Kenya | Africa | 2005 | skin/soft tissue infection, NOS | emm79.5 | 714 | 14 | 25 | 3 | 22 | 27 | 13 | 26 | 29 | 8 | 3 | 25 | 30 |
| K17074 | Kenya | Africa | 2005 | skin/soft tissue infection, NOS | emm218.1 | 292 | 38 | 45 | 3 | 16 | 44 | 44 | 53 | 61 | 54 | 2 | 12 | 3 |

| | | | | | | | | | | | | | | | | | | |
|---|---|---|---|---|---|---|---|---|---|---|---|---|---|---|---|---|---|---|
| K19188 | Kenya | Africa | 2006 | skin/soft tissue infection, NOS | emm8.3 | 505 | 89 | 120 | 3 | 1 | 135 | 45 | 103 | 181 | 66 | 3 | 12 | 30 |
| K21771 | Kenya | Africa | 2007 | invasive, NOS | emm60.7 | 700 | 90 | 12 | 87 | 68 | 132 | 13 | 153 | 179 | 34 | 3 | 108 | 129 |
| K36563 | Kenya | Africa | 2009 | skin/soft tissue infection, NOS | emm229.0 | 703 | 91 | 9 | 8 | 8 | 136 | 13 | 129 | 182 | 8 | 3 | 106 | 130 |
| K38470 | Kenya | Africa | 2009 | skin/soft tissue infection, NOS | emm18.21 | 402 | 11 | 13 | 3 | 41 | 50 | 13 | 112 | 42 | 31 | 1 | 51 | 4 |
| K42952 | Kenya | Africa | 2010 | skin/soft tissue infection, NOS | emm111.2 | 737 | 14 | 37 | 88 | 16 | 12 | 13 | 26 | 29 | 67 | 3 | 109 | 89 |
| K43037 | Kenya | Africa | 2010 | pneumonia | emm44.0 | 178 | 5 | 7 | 7 | 1 | 6 | 7 | 7 | 8 | 14 | 3 | 7 | 7 |
| K50977 | Kenya | Africa | 2011 | invasive, NOS | emm179.0 | 619 | 21 | 111 | 15 | 8 | 46 | 24 | 26 | 30 | 8 | 10 | 12 | 26 |
| K6847 | Kenya | Africa | 2001 | skin/soft tissue infection, NOS | emm65.5 | 215 | 1 | 112 | 24 | 1 | 2 | 13 | 143 | 165 | 8 | 3 | 101 | 118 |
| K6932 | Kenya | Africa | 2001 | skin/soft tissue infection, NOS | emm217.0 | 728 | 14 | 37 | 20 | 24 | 64 | 57 | 145 | 167 | 15 | 3 | 18 | 89 |
| K7275 | Kenya | Africa | 2001 | skin/soft tissue infection, NOS | ND | 450 | 14 | 121 | 3 | 1 | 26 | 13 | 20 | 176 | 65 | 3 | 105 | 124 |
| K7478 | Kenya | Africa | 2001 | skin/soft tissue infection, NOS | emm65.5 | 215 | 1 | 112 | 24 | 1 | 2 | 13 | 143 | 165 | 8 | 3 | 101 | 118 |
| K7928 | Kenya | Africa | 2001 | pneumonia | emm4.5 | 771 | 5 | 65 | 34 | 50 | 26 | 13 | 155 | 183 | 18 | 3 | 50 | 129 |
| K37990 | Kenya | Africa | 2009 | meningitis | emm64.3 | 223 | 14 | 122 | 89 | 24 | 64 | 57 | 156 | 128 | 15 | 3 | 110 | 131 |
| K42771 | Kenya | Africa | 2010 | pneumonia | emm124.2 | 231 | 18 | 123 | 84 | 16 | 82 | 13 | 157 | 184 | 8 | 3 | 111 | 132 |
| K48186 | Kenya | Africa | 2011 | skin/soft tissue infection, NOS | ND | 731 | 92 | 16 | 3 | 16 | 137 | 24 | 26 | 185 | 60 | 3 | 112 | 133 |
| K48650 | Kenya | Africa | 2011 | skin/soft tissue infection, NOS | emm44.0 | 178 | 5 | 7 | 7 | 1 | 6 | 7 | 7 | 8 | 14 | 3 | 7 | 7 |
| K47581 | Kenya | Africa | 2011 | skin/soft tissue infection, NOS | emm80.0 | 729 | 16 | 16 | 3 | 16 | 17 | 7 | 20 | 22 | 1 | 2 | 18 | 110 |
| K16727 | Kenya | Africa | 2005 | skin/soft tissue infection, NOS | emm65.0 | 778 | 5 | 65 | 34 | 27 | 39 | 13 | 158 | 186 | 15 | 3 | 35 | 2 |
| K48083 | Kenya | Africa | 2011 | skin/soft tissue infection, NOS | emm179.0 | 619 | 21 | 111 | 15 | 8 | 46 | 24 | 26 | 187 | 8 | 10 | 12 | 26 |
| K47118 | Kenya | Africa | 2011 | skin/soft tissue infection, NOS | emm217.0 | 728 | 14 | 37 | 20 | 69 | 64 | 57 | 159 | 167 | 15 | 3 | 18 | 89 |
| K46187 | Kenya | Africa | 2010 | skin/soft tissue infection, NOS | emm68.8 | 726 | 1 | 124 | 4 | 70 | 138 | 41 | 160 | 188 | 68 | 3 | 12 | 134 |
| K47020 | Kenya | Africa | 2011 | ND | emm80.0 | 701 | 16 | 16 | 3 | 16 | 17 | 7 | 20 | 22 | 1 | 2 | 18 | 110 |
| K37698 | Kenya | Africa | 2009 | meningitis | emm85.1 | 709 | 14 | 125 | 90 | 71 | 139 | 9 | 161 | 189 | 11 | 3 | 113 | 25 |
| K30465 | Kenya | Africa | 2008 | skin/soft tissue infection, NOS | emm8.3 | 505 | 89 | 120 | 3 | 1 | 135 | 45 | 103 | 181 | 66 | 3 | 12 | 30 |
| K23653 | Kenya | Africa | 2007 | skin/soft tissue infection, NOS | emm8.3 | 241 | 89 | 120 | 3 | 1 | 135 | 45 | 103 | 181 | 66 | 3 | 12 | 30 |
| K36535 | Kenya | Africa | 2009 | skin/soft tissue infection, NOS | emm100.2 | 773 | 93 | 25 | 3 | 26 | 71 | 45 | 162 | 190 | 60 | 26 | 114 | 135 |
| K37741 | Kenya | Africa | 2009 | invasive, NOS | emm239.1 | 776 | 63 | 79 | 91 | 1 | 26 | 13 | 20 | 176 | 65 | 3 | 105 | 124 |
| K6102 | Kenya | Africa | 2000 | skin/soft tissue infection, NOS | emm109.1 | 718 | 94 | 126 | 3 | 27 | 19 | 37 | 163 | 191 | 15 | 3 | 35 | 34 |
| K45527 | Kenya | Africa | 2010 | invasive, NOS | emm192.0 | 724 | 14 | 44 | 24 | 2 | 12 | 41 | 154 | [S] | 8 | 3 | 112 | 3 |
| K45900 | Kenya | Africa | 2010 | skin/soft tissue infection, NOS | emm118.2 | 725 | 1 | 124 | 4 | 72 | 138 | 41 | 164 | 188 | 68 | 3 | 12 | 134 |
| K44869 | Kenya | Africa | 2010 | skin/soft tissue infection, NOS | ND | 450 | 87 | 117 | 3 | 1 | 26 | 13 | 20 | 176 | 65 | 3 | 83 | 136 |
| K21345 | Kenya | Africa | 2007 | skin/soft tissue infection, NOS | ND | 265 | 87 | 117 | 3 | 1 | 26 | 13 | 20 | 176 | 65 | 3 | 105 | 124 |
| K44896 | Kenya | Africa | 2010 | invasive, NOS | emm99.5 | 781 | 14 | 41 | 92 | 1 | 20 | 40 | 26 | 56 | 5 | 4 | 20 | 46 |
| K44098 | Kenya | Africa | 2010 | invasive, NOS | emm18.21 | 402 | 11 | 13 | 3 | 41 | 140 | 13 | 165 | 193 | 31 | 1 | 51 | 4 |
| K44582 | Kenya | Africa | 2010 | skin/soft tissue infection, NOS | emm57.0 | 723 | 21 | 54 | 44 | 1 | 35 | 24 | 166 | 194 | 8 | 10 | 12 | 137 |
| K43101 | Kenya | Africa | 2010 | ND | emm55.0 | 248 | 11 | 11 | 83 | 10 | 10 | 13 | 148 | 13 | 8 | 3 | 11 | 12 |
| K43304 | Kenya | Africa | 2010 | ND | emm18.21 | 402 | 11 | 13 | 3 | 41 | 50 | 13 | 165 | 42 | 31 | 1 | 51 | 4 |
| K48877 | Kenya | Africa | 2011 | ND | emm74.0 | 732 | 11 | 13 | 3 | 41 | 50 | 13 | 112 | 195 | 5 | 4 | 95 | 138 |
| K49285 | Kenya | Africa | 2011 | skin/soft tissue infection, NOS | emm103.0 | 733 | 63 | 79 | 91 | 1 | 26 | 24 | 26 | 196 | 65 | 3 | 105 | 124 |
| K48807 | Kenya | Africa | 2011 | invasive, NOS | emm75.1 | 704 | 14 | 33 | 18 | 6 | 25 | 27 | 40 | 37 | 18 | 3 | 28 | 30 |
| K48817 | Kenya | Africa | 2011 | invasive, NOS | emm74.0 | 120 | 11 | 13 | 3 | 41 | 50 | 13 | 112 | 195 | 5 | 4 | 95 | 138 |
| K12434 | Kenya | Africa | 2004 | skin/soft tissue infection, NOS | emm44.0 | 178 | 5 | 7 | 7 | 1 | 6 | 7 | 7 | 8 | 14 | 3 | 7 | 7 |
| K25147 | Kenya | Africa | 2007 | pneumonia | emm90.5 | 708 | 14 | 115 | 85 | 2 | 127 | 13 | 149 | 171 | 8 | 3 | 12 | 122 |
| K11898 | Kenya | Africa | 2004 | skin/soft tissue infection, NOS | emm111.2 | 737 | 14 | 37 | 88 | 16 | 12 | 13 | 26 | 29 | 69 | 3 | 109 | 89 |
| K16781 | Kenya | Africa | 2005 | skin/soft tissue infection, NOS | ND | 450 | 87 | 117 | 3 | 1 | 26 | 13 | 20 | 176 | 65 | 3 | 105 | 124 |
| K16837 | Kenya | Africa | 2005 | skin/soft tissue infection, NOS | emm83.12 | 393 | 88 | 45 | 3 | 16 | 130 | 85 | 151 | 177 | 60 | 3 | 12 | 30 |
| K13994 | Kenya | Africa | 2005 | invasive, NOS | emm77.0 | 747 | 95 | 127 | 93 | 73 | 2 | 86 | 167 | 197 | 8 | 3 | 20 | 139 |
| K16544 | Kenya | Africa | 2005 | skin/soft tissue infection, NOS | emm11.0 | 250 | 92 | 128 | 3 | 26 | 141 | 24 | 168 | 198 | 70 | 3 | 83 | 28 |

| | | | | | | | | | | | | | | | | | | |
|---|---|---|---|---|---|---|---|---|---|---|---|---|---|---|---|---|---|---|
| K22338 | Kenya | Africa | 2006 | skin/soft tissue infection, NOS | ND | 450 | 87 | 117 | 3 | 1 | 26 | 13 | 20 | 176 | 65 | 3 | 105 | 124 |
| K22633 | Kenya | Africa | 2007 | skin/soft tissue infection, NOS | emm89.8 | 772 | 14 | 49 | 33 | 8 | 138 | 50 | 71 | 199 | 18 | 3 | 83 | 126 |
| K22757 | Kenya | Africa | 2007 | skin/soft tissue infection, NOS | emm83.12 | 393 | 88 | 45 | 3 | 16 | 130 | 85 | 151 | 177 | 60 | 3 | 12 | 30 |
| K23799 | Kenya | Africa | 2007 | skin/soft tissue infection, NOS | emm124.2 | 280 | 18 | 129 | 94 | 16 | 64 | 50 | 71 | 184 | 8 | 3 | 111 | 132 |
| K22813 | Kenya | Africa | 2007 | skin/soft tissue infection, NOS | emm121.0 | 262 | 14 | 44 | 24 | 2 | 12 | 41 | 154 | 200 | 8 | 3 | 101 | 118 |
| K23180 | Kenya | Africa | 2007 | skin/soft tissue infection, NOS | emm44.0 | 178 | 5 | 7 | 7 | 1 | 6 | 7 | 7 | 8 | 14 | 3 | 7 | 7 |
| K23323 | Kenya | Africa | 2007 | skin/soft tissue infection, NOS | emm84.1 | 259 | 14 | 130 | 95 | 1 | 142 | 13 | 169 | 201 | 60 | 3 | 115 | 30 |
| K21710 | Kenya | Africa | 2007 | skin/soft tissue infection, NOS | emm109.1 | 718 | 94 | 126 | 3 | 27 | 19 | 37 | 45 | 191 | 15 | 3 | 35 | 34 |
| K19083 | Kenya | Africa | 2006 | skin/soft tissue infection, NOS | emm56.0 | 115 | 14 | 44 | 24 | 2 | 12 | 41 | 154 | 180 | 8 | 3 | 112 | 3 |
| K49882 | Kenya | Africa | 2011 | skin/soft tissue infection, NOS | emm111.2 | 737 | 14 | 37 | 88 | 16 | 12 | 13 | 26 | 29 | 67 | 3 | 109 | 89 |
| K5698 | Kenya | Africa | 2000 | skin/soft tissue infection, NOS | emm109.1 | 718 | 94 | 126 | 3 | 27 | 19 | 37 | 45 | 191 | 15 | 3 | 35 | 34 |
| K49294 | Kenya | Africa | 2011 | skin/soft tissue infection, NOS | emm90.5 | 734 | 14 | 115 | 85 | 2 | 127 | 13 | 149 | 171 | 8 | 3 | 12 | 122 |
| K49551 | Kenya | Africa | 2011 | pneumonia | emm44.0 | 178 | 5 | 7 | 7 | 1 | 6 | 7 | 7 | 8 | 14 | 3 | 7 | 7 |
| K38181 | Kenya | Africa | 2009 | skin/soft tissue infection, NOS | emm56.0 | 115 | 14 | 44 | 24 | 2 | 12 | 41 | 154 | 180 | 8 | 3 | 112 | 3 |
| K25325 | Kenya | Africa | 2007 | pneumonia | emm90.5 | 708 | 14 | 115 | 85 | 2 | 127 | 13 | 67 | 171 | 8 | 3 | 12 | 122 |
| K40810 | Kenya | Africa | 2010 | skin/soft tissue infection, NOS | emm93.0 | 10 | 16 | 16 | 96 | 16 | 17 | 7 | 20 | 22 | 1 | 2 | 18 | 3 |
| K41947 | Kenya | Africa | 2010 | skin/soft tissue infection, NOS | emm109.1 | 718 | 94 | 126 | 3 | 27 | 19 | 37 | 45 | 191 | 15 | 3 | 35 | 34 |
| K23617 | Kenya | Africa | 2007 | skin/soft tissue infection, NOS | emm48.0 | 278 | 24 | 50 | 97 | 50 | 71 | 45 | 35 | 183 | 18 | 3 | 28 | 30 |
| K6108 | Kenya | Africa | 2000 | skin/soft tissue infection, NOS | emm25.1 | 323 | 14 | 7 | 98 | 2 | 2 | 13 | 26 | 202 | 8 | 3 | 116 | 140 |
| K3637 | Kenya | Africa | 1998 | skin/soft tissue infection, NOS | emm179.0 | 619 | 21 | 111 | 15 | 8 | 46 | 24 | 26 | 30 | 8 | 10 | 12 | 26 |
| K4254 | Kenya | Africa | 1999 | invasive, NOS | emm85.1 | 774 | 14 | 125 | 90 | 71 | 139 | 9 | 161 | 189 | 11 | 3 | 113 | 25 |
| K40818 | Kenya | Africa | 2010 | invasive, NOS | emm112.5 | 777 | 96 | 131 | 3 | 71 | 143 | 13 | 11 | 13 | 8 | 3 | 11 | 12 |
| K6635 | Kenya | Africa | 2000 | skin/soft tissue infection, NOS | emm177.0 | 758 | 84 | 113 | 21 | 23 | 126 | 13 | 147 | 169 | 1 | 2 | 104 | 121 |
| K12473 | Kenya | Africa | 2004 | invasive, NOS | emm55.0 | 232 | 11 | 11 | 83 | 10 | 10 | 13 | 148 | 13 | 8 | 3 | 11 | 12 |
| K5248 | Kenya | Africa | 1999 | skin/soft tissue infection, NOS | emm43.1 | 770 | 16 | 16 | 3 | 16 | 17 | 7 | 170 | 22 | 1 | 2 | 18 | 3 |
| K16898 | Kenya | Africa | 2005 | invasive, NOS | emm112.5 | 246 | 89 | 131 | 3 | 71 | 143 | 13 | 11 | 13 | 8 | 3 | 11 | 12 |
| K7024 | Kenya | Africa | 2001 | skin/soft tissue infection, NOS | emm18.21 | 402 | 11 | 13 | 3 | 41 | 50 | 13 | 112 | 42 | 31 | 1 | 51 | 4 |
| K7087 | Kenya | Africa | 2001 | skin/soft tissue infection, NOS | emm11.0 | 742 | 92 | 128 | 3 | 26 | 141 | 24 | 168 | 198 | 70 | 3 | 83 | 28 |
| K7113 | Kenya | Africa | 2001 | skin/soft tissue infection, NOS | emm11.0 | 742 | 92 | 128 | 3 | 26 | 141 | 24 | 168 | 198 | 70 | 3 | 83 | 28 |
| K38591 | Kenya | Africa | 2009 | skin/soft tissue infection, NOS | emm124.2 | 231 | 18 | 123 | 84 | 16 | 82 | 13 | 157 | 184 | 8 | 3 | 111 | 132 |
| K41948 | Kenya | Africa | 2010 | ND | emm208.0 | 719 | 6 | 132 | 33 | 1 | 144 | 19 | 171 | 203 | 1 | 3 | 18 | 141 |
| K42600 | Kenya | Africa | 2010 | skin/soft tissue infection, NOS | emm42.3 | 721 | 63 | 45 | 3 | 16 | 82 | 13 | 172 | 204 | 1 | 2 | 11 | 142 |
| K37164 | Kenya | Africa | 2009 | skin/soft tissue infection, NOS | emm224.1 | 707 | 14 | 49 | 33 | 1 | 42 | 13 | 173 | 205 | 1 | 3 | 18 | 3 |
| K37287 | Kenya | Africa | 2009 | skin/soft tissue infection, NOS | emm55.0 | 248 | 11 | 11 | 83 | 10 | 10 | 13 | 148 | 13 | 8 | 3 | 11 | 12 |
| K36294 | Kenya | Africa | 2009 | skin/soft tissue infection, NOS | emm49.9 | 705 | 14 | 133 | 3 | 1 | 1 | 13 | 174 | 206 | 71 | 3 | 21 | 143 |
| K36347 | Kenya | Africa | 2009 | skin/soft tissue infection, NOS | emm65.0 | 716 | 14 | 34 | 81 | 27 | 122 | 13 | 142 | 164 | 15 | 3 | 100 | 117 |
| K11464 | Kenya | Africa | 2003 | skin/soft tissue infection, NOS | emm99.5 | 781 | 14 | 41 | 92 | 1 | 20 | 40 | 26 | 56 | 5 | 4 | 20 | 46 |
| K11814 | Kenya | Africa | 2004 | skin/soft tissue infection, NOS | emm169.1 | 226 | 5 | 37 | 99 | 1 | 145 | 87 | 175 | 207 | 72 | 2 | 117 | 144 |
| K17716 | Kenya | Africa | 2005 | skin/soft tissue infection, NOS | emm19.10 | 769 | 12 | 134 | 100 | 12 | 146 | 13 | 13 | 208 | 8 | 3 | 13 | 14 |
| K17300 | Kenya | Africa | 2005 | skin/soft tissue infection, NOS | ND | 450 | 87 | 117 | 3 | 1 | 26 | 13 | 20 | 176 | 65 | 3 | 105 | 124 |
| K19219 | Kenya | Africa | 2006 | skin/soft tissue infection, NOS | emm124.2 | 231 | 18 | 123 | 84 | 16 | 82 | 13 | 157 | 184 | 8 | 3 | 111 | 145 |
| K50658 | Kenya | Africa | 2011 | ND | emm9.0 | 740 | 45 | 34 | 40 | 8 | 124 | 19 | 45 | 168 | 62 | 3 | 12 | 120 |
| K24601 | Kenya | Africa | 2007 | skin/soft tissue infection, NOS | emm89.8 | 772 | 14 | 49 | 33 | 8 | 138 | 50 | 71 | 199 | 18 | 3 | 83 | 126 |
| K24635 | Kenya | Africa | 2007 | skin/soft tissue infection, NOS | emm44.0 | 178 | 5 | 7 | 7 | 1 | 6 | 7 | 7 | 8 | 14 | 3 | 7 | 7 |
| K25713 | Kenya | Africa | 2007 | skin/soft tissue infection, NOS | emm124.2 | 280 | 18 | 129 | 101 | 8 | 64 | 50 | 71 | 184 | 8 | 3 | 111 | 132 |
| K26504 | Kenya | Africa | 2007 | skin/soft tissue infection, NOS | emm44.0 | 178 | 5 | 7 | 7 | 74 | 6 | 7 | 7 | 8 | 14 | 3 | 7 | 7 |
| K28162 | Kenya | Africa | 2007 | skin/soft tissue infection, NOS | emm83.12 | 393 | 88 | 45 | 3 | 16 | 130 | 85 | 151 | 177 | 60 | 3 | 12 | 30 |
| K29166 | Kenya | Africa | 2008 | invasive, NOS | emm162.1 | 412 | 14 | 119 | 102 | 1 | 134 | 1 | 121 | 140 | 5 | 4 | 95 | 128 |

| | | | | | | | | | | | | | | | | | |
|---|---|---|---|---|---|---|---|---|---|---|---|---|---|---|---|---|---|
| K29527 | Kenya | Africa | 2008 | skin/soft tissue infection, NOS | emm4.5 | 771 | 5 | 65 | 34 | 27 | 39 | 13 | 176 | 192 | 15 | 3 | 35 | 2 |
| K29655 | Kenya | Africa | 2008 | skin/soft tissue infection, NOS | emm53.4 | 460 | 14 | 135 | 24 | 1 | 147 | 13 | 177 | 23 | 8 | 4 | 95 | 146 |
| K29743 | Kenya | Africa | 2008 | pneumonia | emm114.5 | 220 | 18 | 136 | 103 | 43 | 148 | 40 | 26 | 56 | 5 | 27 | 83 | 147 |
| K30067 | Kenya | Africa | 2008 | skin/soft tissue infection, NOS | emm73.0 | 469 | 92 | 137 | 104 | 27 | 149 | 50 | 71 | 209 | 18 | 3 | 83 | 25 |
| K31063 | Kenya | Africa | 2008 | invasive, NOS | emm209.0 | 260 | 14 | 37 | 3 | 27 | 132 | 13 | 153 | 179 | 34 | 2 | 107 | 126 |
| K33937 | Kenya | Africa | 2008 | skin/soft tissue infection, NOS | emm60.7 | 700 | 90 | 12 | 87 | 68 | 132 | 13 | 153 | 179 | 34 | 3 | 108 | 129 |
| K31539 | Kenya | Africa | 2008 | invasive, NOS | emm238.1 | 713 | 6 | 117 | 3 | 43 | 150 | 57 | 178 | 210 | 8 | 10 | 12 | 28 |
| K31611 | Kenya | Africa | 2008 | skin/soft tissue infection, NOS | emm93.6 | 613 | 85 | 6 | 6 | 6 | 5 | 6 | 6 | 12 | 5 | 4 | 6 | 11 |
| K32502 | Kenya | Africa | 2008 | skin/soft tissue infection, NOS | emm165.0 | 768 | 58 | 7 | 56 | 1 | 76 | 88 | 27 | 102 | 8 | 3 | 61 | 29 |
| K33560 | Kenya | Africa | 2008 | meningitis | emm179.0 | 619 | 21 | 111 | 15 | 8 | 46 | 24 | 26 | 30 | 8 | 10 | 12 | 26 |
| K20641 | Kenya | Africa | 2006 | skin/soft tissue infection, NOS | emm80.0 | 701 | 16 | 16 | 3 | 16 | 17 | 89 | 20 | 22 | 1 | 2 | 18 | 110 |
| K20653 | Kenya | Africa | 2007 | skin/soft tissue infection, NOS | emm209.0 | 260 | 14 | 37 | 3 | 27 | 132 | 13 | 153 | 179 | 34 | 2 | 107 | 126 |
| K20746 | Kenya | Africa | 2007 | skin/soft tissue infection, NOS | emm97.1 | 283 | 38 | 45 | 3 | 16 | 44 | 44 | 53 | 61 | 54 | 2 | 12 | 3 |
| K20747 | Kenya | Africa | 2007 | skin/soft tissue infection, NOS | emm84.1 | 259 | 14 | 130 | 105 | 1 | 142 | 13 | 169 | 201 | 60 | 3 | 115 | 30 |
| K33951 | Kenya | Africa | 2008 | skin/soft tissue infection, NOS | emm80.0 | 701 | 16 | 16 | 3 | 16 | 17 | 7 | 20 | 22 | 1 | 2 | 18 | 110 |
| K20882 | Kenya | Africa | 2007 | skin/soft tissue infection, NOS | emm209.0 | 260 | 14 | 37 | 3 | 27 | 132 | 13 | 153 | 179 | 34 | 2 | 107 | 126 |
| K20910 | Kenya | Africa | 2007 | skin/soft tissue infection, NOS | emm9.0 | 447 | 96 | 65 | 34 | 27 | 39 | 13 | 179 | 183 | 18 | 3 | 50 | 148 |
| K21246 | Kenya | Africa | 2007 | skin/soft tissue infection, NOS | emm77.0 | 747 | 95 | 127 | 93 | 73 | 2 | 86 | 167 | 197 | 8 | 3 | 20 | 139 |
| K10332 | Kenya | Africa | 2003 | skin/soft tissue infection, NOS | emm183.2 | 761 | 14 | 25 | 3 | 22 | 27 | 13 | 26 | 174 | 8 | 3 | 25 | 30 |
| K10474 | Kenya | Africa | 2003 | skin/soft tissue infection, NOS | emm118.2 | 752 | 1 | 124 | 4 | 72 | 138 | 41 | 164 | 188 | 68 | 3 | 12 | 134 |
| K10586 | Kenya | Africa | 2003 | skin/soft tissue infection, NOS | emm50.3 | 217 | 111 | 138 | 33 | 1 | 151 | 13 | 32 | 211 | 42 | 2 | 118 | 149 |
| K10722 | Kenya | Africa | 2003 | skin/soft tissue infection, NOS | emm22.5 | 213 | 112 | 37 | 20 | 24 | 64 | 57 | 110 | 128 | 15 | 3 | 18 | 150 |
| K11116 | Kenya | Africa | 2003 | skin/soft tissue infection, NOS | emm18.21 | 402 | 11 | 13 | 3 | 41 | 50 | 13 | 112 | 42 | 31 | 1 | 51 | 4 |
| K11239 | Kenya | Africa | 2003 | skin/soft tissue infection, NOS | emm169.1 | 238 | 5 | 37 | 99 | 1 | 152 | 87 | 175 | 207 | 72 | 2 | 117 | 144 |
| K11243 | Kenya | Africa | 2003 | skin/soft tissue infection, NOS | emm114.5 | 220 | 18 | 136 | 103 | 43 | 148 | 40 | 26 | 56 | 5 | 27 | 83 | 147 |
| K33983 | Kenya | Africa | 2008 | skin/soft tissue infection, NOS | emm30.15 | 537 | 21 | 21 | 15 | 8 | 46 | 24 | 26 | 30 | 8 | 10 | 12 | 26 |
| K11319 | Kenya | Africa | 2003 | skin/soft tissue infection, NOS | emm64.3 | 223 | 14 | 122 | 20 | 24 | 64 | 57 | 156 | 128 | 15 | 28 | 110 | 131 |
| K13065 | Kenya | Africa | 2004 | skin/soft tissue infection, NOS | emm75.1 | 578 | 59 | 74 | 97 | 50 | 71 | 45 | 35 | 183 | 18 | 3 | 18 | 131 |
| K13107 | Kenya | Africa | 2004 | skin/soft tissue infection, NOS | emm183.2 | 219 | 14 | 25 | 3 | 22 | 27 | 13 | 26 | 29 | 8 | 3 | 25 | 30 |
| K13190 | Kenya | Africa | 2004 | skin/soft tissue infection, NOS | emm39.4 | 236 | 14 | 139 | 4 | 12 | 12 | 13 | 13 | 212 | 8 | 3 | 13 | 14 |
| K13372 | Kenya | Africa | 2004 | skin/soft tissue infection, NOS | emm81.2 | 766 | 113 | 140 | 106 | 1 | 26 | 13 | 180 | 213 | 66 | 3 | 12 | 30 |
| K17276 | Kenya | Africa | 2005 | skin/soft tissue infection, NOS | emm92.0 | 674 | 114 | 65 | 34 | 27 | 39 | 13 | 22 | 152 | 13 | 2 | 20 | 22 |
| K23685 | Kenya | Africa | 2007 | skin/soft tissue infection, NOS | emm79.5 | 714 | 14 | 25 | 3 | 22 | 27 | 13 | 26 | 29 | 8 | 3 | 25 | 30 |
| K23745 | Kenya | Africa | 2007 | skin/soft tissue infection, NOS | emm60.7 | 279 | 59 | 74 | 107 | 50 | 71 | 57 | 181 | 214 | 15 | 3 | 18 | 89 |
| K23866 | Kenya | Africa | 2007 | skin/soft tissue infection, NOS | emm82.5 | 257 | 14 | 141 | 7 | 8 | 2 | 1 | 121 | 23 | 15 | 3 | 18 | 151 |
| K35870 | Kenya | Africa | 2008 | skin/soft tissue infection, NOS | emm229.0 | 703 | 91 | 9 | 8 | 8 | 136 | 13 | 129 | 182 | 8 | 3 | 106 | 130 |
| K23890 | Kenya | Africa | 2007 | skin/soft tissue infection, NOS | emm97.1 | 283 | 38 | 45 | 3 | 16 | 44 | 44 | 53 | 61 | 54 | 2 | 12 | 3 |
| K24146 | Kenya | Africa | 2007 | skin/soft tissue infection, NOS | emm165.0 | 768 | 58 | 7 | 56 | 1 | 76 | 88 | 27 | 102 | 8 | 3 | 61 | 29 |
| K24190 | Kenya | Africa | 2007 | skin/soft tissue infection, NOS | emm28.0 | 763 | 96 | 65 | 34 | 27 | 39 | 13 | 182 | 215 | 13 | 2 | 20 | 22 |
| K24525 | Kenya | Africa | 2007 | skin/soft tissue infection, NOS | emm11.0 | ND | 24 | 19 | 78 | 23 | 117 | 83 | 183 | 216 | 8 | 3 | 18 | 4 |
| K19347 | Kenya | Africa | 2006 | skin/soft tissue infection, NOS | ND | 258 | 92 | 16 | 3 | 16 | 17 | 24 | 26 | 185 | 60 | 3 | 112 | 133 |
| K19376 | Kenya | Africa | 2006 | skin/soft tissue infection, NOS | emm75.1 | 578 | 59 | 74 | 97 | 50 | 71 | 45 | 35 | 183 | 18 | 3 | 18 | 131 |
| K19669 | Kenya | Africa | 2006 | skin/soft tissue infection, NOS | ND | 258 | 92 | 16 | 3 | 16 | 17 | 24 | 26 | 217 | 73 | 3 | 112 | 133 |
| K35909 | Kenya | Africa | 2009 | skin/soft tissue infection, NOS | emm75.1 | 704 | 14 | 33 | 18 | 6 | 25 | 27 | 40 | 37 | 18 | 3 | 28 | 30 |
| K50593 | Kenya | Africa | 2011 | pneumonia | emm9.0 | 740 | 45 | 34 | 40 | 8 | 124 | 19 | 45 | 168 | 62 | 3 | 12 | 120 |
| K19875 | Kenya | Africa | 2006 | skin/soft tissue infection, NOS | emm111.2 | 256 | 14 | 37 | 88 | 16 | 12 | 90 | 26 | 29 | 69 | 3 | 109 | 89 |
| K19912 | Kenya | Africa | 2006 | skin/soft tissue infection, NOS | emm92.0 | 674 | 114 | 65 | 34 | 27 | 39 | 13 | 22 | 152 | 13 | 2 | 20 | 22 |
| K19952 | Kenya | Africa | 2006 | skin/soft tissue infection, NOS | emm92.0 | 674 | 114 | 65 | 34 | 27 | 39 | 13 | 22 | 152 | 13 | 2 | 20 | 22 |

| | | | | | | | | | | | | | | | | | |
|---|---|---|---|---|---|---|---|---|---|---|---|---|---|---|---|---|---|
| K20201 | Kenya | Africa | 2006 | skin/soft tissue infection, NOS | emm44.0 | 178 | 5 | 7 | 7 | 1 | 6 | 7 | 7 | 8 | 14 | 3 | 7 | 7 |
| K36395 | Kenya | Africa | 2009 | skin/soft tissue infection, NOS | emm81.2 | 766 | 113 | 140 | 106 | 1 | 26 | 13 | 180 | 213 | 66 | 3 | 12 | 30 |
| K35215 | Kenya | Africa | 2008 | skin/soft tissue infection, NOS | ND | 450 | 87 | 117 | 3 | 1 | 26 | 13 | 20 | 176 | 65 | 3 | 105 | 124 |
| K36067 | Kenya | Africa | 2009 | ND | emm85.1 | 774 | 14 | 125 | 90 | 71 | 139 | 9 | 161 | 189 | 11 | 3 | 113 | 25 |
| K50105 | Kenya | Africa | 2011 | skin/soft tissue infection, NOS | emm22.5 | 213 | 112 | 37 | 20 | 24 | 64 | 57 | 110 | 128 | 15 | 3 | 18 | 150 |
| K31028 | Kenya | Africa | 2008 | skin/soft tissue infection, NOS | emm93.6 | 613 | 85 | 6 | 6 | 6 | 5 | 6 | 6 | 12 | 5 | 4 | 6 | 11 |
| K35129 | Kenya | Africa | 2008 | skin/soft tissue infection, NOS | ND | 258 | 92 | 16 | 3 | 16 | 17 | 24 | 26 | 218 | 60 | 3 | 112 | 133 |
| K27345 | Kenya | Africa | 2007 | skin/soft tissue infection, NOS | ND | 450 | 87 | 117 | 3 | 1 | 26 | 13 | 20 | 176 | 65 | 3 | 105 | 124 |
| K28044 | Kenya | Africa | 2007 | pneumonia | emm25.1 | 323 | 14 | 7 | 98 | 2 | 2 | 13 | 26 | 202 | 8 | 3 | 116 | 140 |
| K19417 | Kenya | Africa | 2006 | skin/soft tissue infection, NOS | emm122.0 | 200 | 11 | 13 | 3 | 41 | 50 | 13 | 112 | 42 | 31 | 1 | 51 | 4 |
| K50316 | Kenya | Africa | 2011 | skin/soft tissue infection, NOS | emm64.3 | 223 | 14 | 122 | 20 | 24 | 64 | 57 | 156 | 128 | 15 | 3 | 110 | 131 |
| K7541 | Kenya | Africa | 2001 | skin/soft tissue infection, NOS | emm81.2 | 766 | 113 | 140 | 106 | 1 | 26 | 13 | 180 | 213 | 66 | 3 | 12 | 30 |
| K7114 | Kenya | Africa | 2001 | skin/soft tissue infection, NOS | ND | 450 | 87 | 117 | 3 | 1 | 26 | 13 | 20 | 176 | 65 | 3 | 105 | 124 |
| K7151 | Kenya | Africa | 2001 | skin/soft tissue infection, NOS | emm183.2 | 761 | 14 | 25 | 3 | 22 | 27 | 13 | 26 | 174 | 8 | 3 | 25 | 30 |
| K8955 | Kenya | Africa | 2002 | skin/soft tissue infection, NOS | emm65.0 | 778 | 5 | 65 | 34 | 27 | 39 | [S] | 158 | 186 | 15 | 3 | 35 | 2 |
| K9333 | Kenya | Africa | 2002 | skin/soft tissue infection, NOS | emm90.5 | 708 | 14 | 115 | 85 | 2 | 127 | 13 | 149 | 171 | 8 | 3 | 12 | 122 |
| K10016 | Kenya | Africa | 2003 | skin/soft tissue infection, NOS | emm103.0 | 233 | 115 | 79 | 91 | 1 | 26 | 24 | 26 | 196 | 65 | 3 | 105 | 124 |
| K10021 | Kenya | Africa | 2003 | skin/soft tissue infection, NOS | emm103.0 | 233 | 63 | 79 | 108 | 1 | 26 | 24 | 26 | 219 | 65 | 3 | 105 | 124 |
| K9408 | Kenya | Africa | 2002 | skin/soft tissue infection, NOS | emm65.0 | 778 | 5 | 65 | 34 | 27 | 39 | 13 | 158 | 186 | 15 | 3 | 35 | 2 |
| K10040 | Kenya | Africa | 2003 | skin/soft tissue infection, NOS | emm22.5 | 213 | 112 | 37 | 20 | 24 | 64 | 57 | 110 | 128 | 15 | 3 | 18 | 150 |
| K7829 | Kenya | Africa | 2001 | skin/soft tissue infection, NOS | emm186.1 | 262 | 14 | 44 | 24 | 2 | 12 | 41 | 154 | 180 | 74 | 3 | 101 | 118 |
| K10105 | Kenya | Africa | 2003 | skin/soft tissue infection, NOS | emm22.5 | 213 | 112 | 37 | 20 | 24 | 64 | 57 | 110 | 128 | 15 | 3 | 18 | 150 |
| K10712 | Kenya | Africa | 2003 | invasive, NOS | emm183.2 | 219 | 14 | 25 | 3 | 22 | 27 | 13 | 26 | 29 | 8 | 3 | 25 | 30 |
| K5086 | Kenya | Africa | 1999 | invasive, NOS | emm238.1 | 713 | 6 | 117 | 3 | 43 | 150 | 57 | 178 | 210 | 8 | 10 | 12 | 28 |
| K10987 | Kenya | Africa | 2003 | pneumonia | emm230.1 | 755 | 14 | 142 | 24 | 1 | 153 | 13 | 177 | 23 | 5 | 4 | 95 | 146 |
| K10238 | Kenya | Africa | 2003 | APSGN | emm218.1 | 292 | 38 | 45 | 3 | 16 | 44 | 44 | 53 | 61 | 54 | 2 | 12 | 3 |
| K9440 | Kenya | Africa | 2002 | invasive, NOS | emm99.5 | 779 | 14 | 41 | 92 | 1 | 20 | 40 | 26 | 56 | 5 | 4 | 20 | 46 |
| K19639 | Kenya | Africa | 2006 | invasive, NOS | emm111.1 | 496 | 11 | 11 | 109 | 10 | 10 | 7 | 77 | 39 | 19 | 2 | 30 | 152 |
| K17494 | Kenya | Africa | 2005 | invasive, NOS | emm55.0 | 248 | 11 | 11 | 83 | 10 | 10 | 13 | 148 | 13 | 8 | 3 | 11 | 12 |
| K7908 | Kenya | Africa | 2001 | skin/soft tissue infection, NOS | emm65.4 | 1031 | 11 | 66 | 40 | 35 | 68 | 76 | 44 | 145 | 1 | 2 | 87 | 31 |
| K7175 | Kenya | Africa | 2001 | invasive, NOS | emm165.0 | 762 | 59 | 74 | 97 | 50 | 71 | 45 | 35 | 183 | 5 | 4 | 20 | 46 |
| K18724 | Kenya | Africa | 2006 | pneumonia | emm11.0 | 250 | 92 | 128 | 3 | 26 | 141 | 24 | 168 | 198 | 70 | 3 | 83 | 28 |
| K7554 | Kenya | Africa | 2001 | invasive, NOS | emm90.5 | 708 | 14 | 143 | 85 | 2 | 127 | 13 | 67 | 171 | 8 | 3 | 12 | 122 |
| K10697 | Kenya | Africa | 2003 | invasive, NOS | emm18.21 | 221 | 11 | 114 | 84 | 16 | 82 | 13 | 20 | 170 | 63 | 3 | 13 | 30 |
| K5192 | Kenya | Africa | 1999 | invasive, NOS | emm147.0 | 741 | 88 | 45 | 3 | 16 | 130 | 85 | 151 | 177 | 60 | 3 | 12 | 30 |
| K4761 | Kenya | Africa | 1999 | pneumonia | emm8.3 | 730 | 89 | 120 | 3 | 1 | 135 | 45 | 103 | 181 | 66 | 3 | 12 | 30 |
| K16772 | Kenya | Africa | 2005 | invasive, NOS | emm89.8 | 245 | 14 | 49 | 33 | 8 | 138 | 50 | 71 | 199 | 18 | 3 | 83 | 126 |
| K3573 | Kenya | Africa | 1998 | invasive, NOS | emm116.2 | 702 | 14 | 119 | 24 | 1 | 153 | 13 | 177 | 23 | 75 | 4 | 95 | 146 |
| K16738 | Kenya | Africa | 2005 | invasive, NOS | emm64.3 | 223 | 14 | 122 | 20 | 24 | 64 | 57 | 156 | 128 | 76 | 28 | 110 | 131 |
| K7559 | Kenya | Africa | 2001 | invasive, NOS | emm110.0 | 767 | 84 | 144 | 110 | 27 | 154 | 33 | 144 | 220 | 1 | 3 | 119 | 28 |
| K3525 | Kenya | Africa | 1998 | invasive, NOS | emm55.0 | 100 | 11 | 11 | 83 | 10 | 10 | 13 | [S] | 13 | 8 | 3 | 11 | 12 |
| K3879 | Kenya | Africa | 1998 | invasive, NOS | emm183.2 | 714 | 14 | 25 | 3 | 22 | 27 | 13 | 26 | 29 | 8 | 3 | 25 | 30 |
| K19873 | Kenya | Africa | 2006 | meningitis | emm78.5 | 255 | 86 | 34 | 40 | 8 | 128 | 19 | 45 | 173 | 62 | 3 | 12 | 120 |
| K4973 | Kenya | Africa | 1999 | invasive, NOS | emm25.4 | 735 | 116 | 145 | 111 | 23 | 155 | 92 | 185 | 221 | 77 | 2 | 120 | 3 |
| K8861 | Kenya | Africa | 2002 | invasive, NOS | emm44.0 | 178 | 5 | 7 | 7 | 1 | 6 | 7 | 7 | 8 | 14 | 3 | 7 | 7 |
| K13045 | Kenya | Africa | 2004 | pneumonia | emm192.0 | 261 | 14 | 44 | 24 | 2 | 12 | 41 | 154 | 180 | 8 | 3 | 112 | 3 |
| K12183 | Kenya | Africa | 2004 | invasive, NOS | emm124.2 | 231 | 18 | 123 | 84 | 16 | 82 | 13 | 157 | 184 | 8 | 3 | 111 | 132 |
| K37914 | Kenya | Africa | 2009 | ND | emm119.2 | 239 | 38 | 45 | 3 | 16 | 44 | 44 | 53 | 61 | 54 | 2 | 12 | 3 |

| | | | | | | | | | | | | | | | | | |
|---|---|---|---|---|---|---|---|---|---|---|---|---|---|---|---|---|---|
| K4728 | Kenya | Africa | 1999 | pneumonia | emm44.0 | 178 | 5 | 7 | 7 | 1 | 6 | 7 | 7 | 8 | 14 | 3 | 7 | 7 |
| K9679 | Kenya | Africa | 2002 | pneumonia | emm112.5 | 777 | 96 | 131 | 3 | 71 | 143 | 13 | 11 | 13 | 8 | 3 | 11 | 153 |
| K17786 | Kenya | Africa | 2006 | meningitis | emm11.0 | 251 | 92 | 128 | 3 | 26 | 141 | 24 | 168 | 198 | [S] | 3 | 83 | 28 |
| K3730 | Kenya | Africa | 1998 | invasive, NOS | emm90.5 | 708 | 14 | 115 | 85 | 2 | 127 | 13 | 67 | 171 | 8 | 3 | 12 | 122 |
| K10378 | Kenya | Africa | 2003 | invasive, NOS | emm50.3 | 217 | 111 | 138 | 33 | 1 | 151 | 13 | 32 | 211 | 42 | 2 | 118 | 149 |
| K8460 | Kenya | Africa | 2002 | invasive, NOS | emm82.5 | 257 | 14 | 141 | 7 | 8 | 2 | 1 | 121 | 23 | 15 | 3 | 18 | 151 |
| K12669 | Kenya | Africa | 2004 | pneumonia | emm183.2 | 234 | 19 | 25 | 3 | 22 | 27 | 13 | 26 | 29 | 8 | 3 | 25 | 30 |
| K5851 | Kenya | Africa | 2000 | invasive, NOS | emm230.1 | 755 | 14 | 142 | 24 | 1 | 156 | 13 | 177 | 23 | [S] | 4 | 95 | 146 |
| K7792 | Kenya | Africa | 2001 | skin/soft tissue infection, NOS | emm43.1 | 770 | 16 | 16 | 3 | 16 | 17 | 7 | 20 | 222 | 1 | 2 | 18 | 3 |
| K9454 | Kenya | Africa | 2002 | invasive, NOS | emm50.3 | 217 | 111 | 138 | 33 | 1 | 151 | 13 | 32 | 211 | 42 | 2 | 118 | 149 |
| K9612 | Kenya | Africa | 2002 | pneumonia | emm99.5 | 781 | 14 | 41 | 92 | 1 | 20 | 40 | 26 | 56 | 5 | 4 | 20 | 46 |
| K21633 | Kenya | Africa | 2007 | invasive, NOS | emm15.1 | 266 | 117 | 146 | 20 | 24 | 64 | 2 | 67 | 223 | 66 | 3 | 12 | 174 |
| K7219 | Kenya | Africa | 2001 | invasive, NOS | emm28.0 | 763 | 96 | 65 | 34 | 27 | 39 | 13 | 182 | 215 | 13 | 2 | 20 | 22 |
| K10676 | Kenya | Africa | 2003 | invasive, NOS | emm103.0 | 233 | 63 | 79 | 91 | 1 | 26 | 24 | 26 | 196 | 65 | 3 | 105 | 124 |
| K10167 | Kenya | Africa | 2003 | pneumonia | emm98.3 | 136 | 118 | 44 | 24 | 75 | 12 | 41 | 154 | 180 | 8 | 3 | 101 | 118 |
| K8015 | Kenya | Africa | 2001 | skin/soft tissue infection, NOS | emm99.5 | 781 | 14 | 41 | 92 | 1 | 20 | 40 | 26 | 56 | 5 | 4 | 20 | 175 |
| K9521 | Kenya | Africa | 2002 | pneumonia | emm84.1 | 780 | 14 | 130 | 112 | 1 | 142 | 13 | 186 | 75 | 80 | 2 | 121 | 2 |
| K5679 | Kenya | Africa | 2000 | pneumonia | emm177.0 | 743 | 84 | 113 | 21 | 23 | 138 | 13 | 147 | 169 | 1 | 2 | 104 | 121 |
| K7393 | Kenya | Africa | 2001 | invasive, NOS | emm63.5 | 764 | 14 | 116 | 86 | 27 | 129 | 9 | 150 | 175 | 64 | 2 | 12 | 123 |
| K6222 | Kenya | Africa | 2000 | invasive, NOS | emm70.0 | 754 | 14 | 147 | 3 | 26 | 157 | 50 | 187 | 224 | 60 | 3 | 115 | 30 |
| K17097 | Kenya | Africa | 2005 | pneumonia | emm223.0 | 536 | 85 | 6 | 6 | 6 | 5 | 6 | 6 | 12 | 5 | 4 | 6 | 11 |
| K9215 | Kenya | Africa | 2002 | meningitis | emm112.5 | 777 | 96 | 131 | 3 | 71 | 143 | 13 | 11 | 13 | 8 | 3 | 11 | 153 |
| K12554 | Kenya | Africa | 2004 | invasive, NOS | emm44.0 | 178 | 5 | 7 | 7 | 1 | 6 | 7 | 7 | 8 | 14 | 3 | 7 | 7 |
| K13484 | Kenya | Africa | 2004 | pneumonia | emm119.2 | 239 | 38 | 45 | 3 | 16 | 44 | 44 | 53 | 61 | 54 | 2 | 12 | 3 |
| K3944 | Kenya | Africa | 1998 | invasive, NOS | emm122.0 | 200 | 11 | 13 | 3 | 41 | 50 | 13 | 112 | 42 | 31 | 1 | 51 | 4 |
| K10311 | Kenya | Africa | 2003 | invasive, NOS | emm50.3 | 217 | 111 | 138 | 33 | 1 | 151 | 13 | 32 | 211 | 42 | 2 | 118 | 149 |
| K10812 | Kenya | Africa | 2003 | pneumonia | emm82.5 | 257 | 14 | 141 | 7 | 8 | 2 | 1 | 121 | 23 | 15 | 3 | 18 | 151 |
| K7846 | Kenya | Africa | 2001 | invasive, NOS | emm74.0 | 120 | 11 | 13 | 3 | 41 | 50 | 13 | 112 | 195 | 5 | 4 | 95 | 138 |
| K3589 | Kenya | Africa | 1998 | meningitis | emm8.3 | 241 | 89 | 120 | 3 | 1 | 135 | 45 | 103 | 181 | 66 | 3 | 12 | 30 |
| K8543 | Kenya | Africa | 2002 | invasive, NOS | emm85.1 | 774 | 14 | 125 | 90 | 71 | 139 | 9 | 161 | 189 | 11 | 3 | 113 | 25 |
| K8057 | Kenya | Africa | 2001 | pneumonia | emm179.0 | 619 | 21 | 111 | 15 | 8 | 158 | 24 | 26 | 225 | 8 | 10 | 12 | 26 |
| K11254 | Kenya | Africa | 2003 | invasive, NOS | emm18.21 | 221 | 11 | 114 | 84 | 16 | 82 | 13 | 20 | 170 | 60 | 3 | 13 | 30 |
| K11271 | Kenya | Africa | 2003 | pneumonia | emm25.4 | 222 | 116 | 145 | 111 | 23 | 155 | 92 | 185 | 221 | 77 | 2 | 120 | 3 |
| K9037 | Kenya | Africa | 2002 | invasive, NOS | emm239.1 | 776 | 63 | 79 | 91 | 1 | 26 | 13 | 20 | 176 | 65 | 3 | 105 | 124 |
| K4350 | Kenya | Africa | 1999 | invasive, NOS | emm22.5 | 213 | 112 | 37 | 20 | 24 | 64 | 57 | 110 | 128 | 15 | 3 | 18 | 150 |
| K7736 | Kenya | Africa | 2001 | APSGN | emm19.10 | 769 | 12 | 134 | 100 | 12 | 146 | 13 | 13 | 208 | 8 | 3 | 13 | 14 |
| K7731 | Kenya | Africa | 2001 | invasive, NOS | emm100.2 | 773 | 93 | 25 | 3 | 26 | 71 | 45 | 162 | 190 | 60 | 26 | 114 | 135 |
| K14810 | Kenya | Africa | 2005 | invasive, NOS | emm18.21 | 402 | 11 | 13 | 3 | 41 | 50 | 13 | 112 | 42 | 31 | 1 | 51 | 4 |
| K10213 | Kenya | Africa | 2003 | skin/soft tissue infection, NOS | emm169.1 | 238 | 5 | 37 | 99 | 1 | 145 | 87 | 175 | 207 | 72 | 2 | 117 | 144 |
| K3808 | Kenya | Africa | 1998 | skin/soft tissue infection, NOS | emm95.0 | 712 | 85 | 6 | 6 | 6 | 5 | 6 | 6 | 226 | 5 | 4 | 6 | 176 |
| K20001 | Kenya | Africa | 2006 | invasive, NOS | emm77.0 | 747 | 95 | 127 | 93 | 73 | 2 | 86 | 167 | 197 | 8 | 3 | 20 | 139 |
| K3828 | Kenya | Africa | 1998 | skin/soft tissue infection, NOS | emm238.1 | 713 | 6 | 117 | 3 | 43 | 150 | 57 | 178 | 210 | 8 | 10 | 12 | 28 |
| K3964 | Kenya | Africa | 1999 | skin/soft tissue infection, NOS | emm65.0 | 716 | 14 | 34 | 81 | 27 | 122 | 13 | 142 | 164 | 15 | 3 | 100 | 117 |
| K3997 | Kenya | Africa | 1999 | skin/soft tissue infection, NOS | emm65.0 | 716 | 14 | 34 | 81 | 27 | 122 | 13 | 142 | 164 | 15 | 3 | 100 | 117 |
| K4134 | Kenya | Africa | 1999 | invasive, NOS | emm209.1 | 717 | 1 | 148 | 115 | 1 | 159 | 13 | 188 | 227 | 15 | 2 | 122 | 177 |
| K4240 | Kenya | Africa | 1999 | skin/soft tissue infection, NOS | emm74.0 | 720 | 11 | 13 | 3 | 41 | 50 | 13 | 112 | 195 | 5 | 4 | 95 | 138 |
| K4263 | Kenya | Africa | 1999 | skin/soft tissue infection, NOS | emm230.1 | 755 | 14 | 142 | 24 | 1 | 153 | 13 | 177 | 23 | 5 | 4 | 95 | 146 |
| K4656 | Kenya | Africa | 1999 | skin/soft tissue infection, NOS | emm92.1 | 727 | 5 | 65 | 34 | 27 | 150 | 57 | 189 | [S] | 18 | 3 | 18 | 178 |

| | | | | | | | | | | | | | | | | | | |
|---|---|---|---|---|---|---|---|---|---|---|---|---|---|---|---|---|---|---|
| K4819 | Kenya | Africa | 1999 | skin/soft tissue infection, NOS | emm230.1 | 755 | 14 | 142 | 24 | 1 | 153 | 13 | 177 | 23 | 5 | 4 | 95 | 146 |
| K4982 | Kenya | Africa | 1999 | skin/soft tissue infection, NOS | emm124.2 | 736 | 18 | 129 | 94 | 16 | 64 | 50 | 71 | 184 | 8 | 3 | 111 | 132 |
| K5460 | Kenya | Africa | 2000 | skin/soft tissue infection, NOS | emm11.0 | 742 | 92 | 128 | 3 | 26 | 141 | 24 | 168 | 198 | 70 | 3 | 83 | 28 |
| K8396 | Kenya | Africa | 2002 | invasive, NOS | emm89.8 | 772 | 14 | 49 | 33 | 8 | 138 | 50 | 71 | 199 | 18 | 3 | 83 | 126 |
| K5660 | Kenya | Africa | 2000 | skin/soft tissue infection, NOS | ND | 1085 | 87 | 117 | 3 | 1 | 26 | 13 | 20 | 176 | 65 | 3 | 105 | 124 |
| K5690 | Kenya | Africa | 2000 | skin/soft tissue infection, NOS | emm81.11 | 744 | 8 | 4 | 4 | 4 | 160 | 13 | 4 | 228 | 31 | 29 | 4 | 4 |
| K5727 | Kenya | Africa | 2000 | skin/soft tissue infection, NOS | emm25.1 | 746 | 14 | 7 | 117 | 76 | 55 | 13 | 13 | 40 | 36 | 3 | 83 | 179 |
| K8492 | Kenya | Africa | 2002 | skin/soft tissue infection, NOS | emm100.2 | 773 | 93 | 25 | 3 | 26 | 71 | 45 | 162 | 190 | 60 | 26 | 114 | 135 |
| K8744 | Kenya | Africa | 2002 | skin/soft tissue infection, NOS | emm8.3 | 241 | 89 | 120 | 3 | 1 | 135 | 45 | 190 | 181 | 66 | 3 | 12 | 30 |
| K5797 | Kenya | Africa | 2000 | skin/soft tissue infection, NOS | emm77.0 | 747 | 95 | 127 | 93 | 73 | 2 | 86 | 167 | 197 | 8 | 3 | 20 | 139 |
| K5898 | Kenya | Africa | 2000 | skin/soft tissue infection, NOS | emm209.1 | 749 | 1 | 148 | 115 | 1 | 159 | 13 | 188 | 227 | 15 | 2 | 122 | 177 |
| K5910 | Kenya | Africa | 2000 | skin/soft tissue infection, NOS | emm92.1 | 750 | 5 | 65 | 34 | 27 | 150 | 57 | 189 | 229 | 18 | 3 | 18 | 178 |
| K5965 | Kenya | Africa | 2000 | skin/soft tissue infection, NOS | emm147.0 | 753 | 88 | 45 | 3 | 16 | 130 | 85 | 151 | 177 | 60 | 3 | 12 | 30 |
| K6008 | Kenya | Africa | 2000 | skin/soft tissue infection, NOS | emm75.1 | 751 | 119 | 37 | 3 | 43 | 161 | 19 | 107 | 230 | 9 | 11 | 57 | 120 |
| K6038 | Kenya | Africa | 2000 | skin/soft tissue infection, NOS | emm118.2 | 752 | 1 | 124 | 4 | 72 | 138 | 41 | 164 | 188 | 68 | 3 | 12 | 134 |
| K6048 | Kenya | Africa | 2000 | skin/soft tissue infection, NOS | emm147.0 | 753 | 88 | 45 | 3 | 16 | 130 | 85 | 151 | 177 | 60 | 3 | 12 | 30 |
| K6085 | Kenya | Africa | 2000 | skin/soft tissue infection, NOS | emm8.3 | 241 | 89 | 120 | 3 | 1 | 135 | 45 | 191 | 181 | 66 | 3 | 12 | 30 |
| K6099 | Kenya | Africa | 2000 | skin/soft tissue infection, NOS | emm77.0 | 482 | 14 | 110 | 3 | 22 | 27 | 13 | 26 | 29 | 8 | 3 | 25 | 30 |
| K6363 | Kenya | Africa | 2000 | skin/soft tissue infection, NOS | emm230.1 | 755 | 14 | 142 | 24 | 77 | 153 | 13 | 177 | 23 | 5 | 4 | 95 | 146 |
| K6428 | Kenya | Africa | 2000 | skin/soft tissue infection, NOS | emm97.1 | 756 | 21 | 21 | 15 | 8 | 46 | 24 | 26 | 231 | 8 | 10 | 12 | 26 |
| K7498 | Kenya | Africa | 2001 | invasive, NOS | emm44.0 | 178 | 5 | 7 | 7 | 1 | 6 | 7 | 7 | 8 | 14 | 3 | 7 | 7 |
| K6917 | Kenya | Africa | 2001 | skin/soft tissue infection, NOS | emm65.5 | 215 | 1 | 112 | 24 | 1 | 2 | 13 | 143 | 165 | 8 | 3 | 101 | 118 |
| K4828 | Kenya | Africa | 1999 | invasive, NOS | emm179.0 | 619 | 21 | 111 | 15 | 8 | 162 | 24 | 26 | 30 | 8 | 10 | 12 | 26 |
| K8728 | Kenya | Africa | 2002 | invasive, NOS | emm64.3 | 223 | 14 | 122 | 20 | 24 | 64 | 57 | 156 | 128 | 15 | 3 | 110 | 131 |
| K13179 | Kenya | Africa | 2004 | invasive, NOS | ND | 235 | 6 | 118 | 71 | 1 | 163 | 13 | 152 | 232 | 8 | 3 | 106 | 180 |
| K12537 | Kenya | Africa | 2004 | meningitis | emm103.0 | 233 | 63 | 79 | 91 | 1 | 26 | 24 | 26 | 196 | 65 | 3 | 105 | 124 |
| K23182 | Kenya | Africa | 2007 | pneumonia | emm63.5 | 274 | 59 | 74 | 97 | 50 | 71 | 45 | 35 | 183 | 81 | 3 | 18 | 154 |
| K19961 | Kenya | Africa | 2006 | pneumonia | emm82.5 | 257 | 14 | 141 | 7 | 8 | 2 | 1 | 121 | 23 | 15 | 3 | 18 | 151 |
| K5017 | Kenya | Africa | 1999 | meningitis | emm44.0 | 178 | 5 | 7 | 7 | 1 | 6 | 7 | 7 | 8 | 14 | 3 | 7 | 7 |
| K10514 | Kenya | Africa | 2003 | invasive, NOS | emm77.0 | 218 | 14 | 110 | 3 | 22 | 27 | 13 | 26 | 29 | 8 | 3 | 25 | 30 |
| K12614 | Kenya | Africa | 2004 | invasive, NOS | emm179.0 | 619 | 21 | 111 | 15 | 8 | 46 | 24 | 26 | 30 | 8 | 10 | 12 | 26 |
| K9361 | Kenya | Africa | 2002 | pneumonia | emm44.0 | 178 | 5 | 7 | 7 | 1 | 6 | 7 | 7 | 8 | 14 | 3 | 7 | 7 |
| K13389 | Kenya | Africa | 2004 | invasive, NOS | emm169.1 | 238 | 5 | 37 | 99 | 1 | 145 | 87 | 175 | 207 | 72 | 2 | 117 | 144 |
| K9189 | Kenya | Africa | 2002 | pneumonia | emm182.1 | 229 | 14 | 41 | 120 | 1 | 164 | 40 | 26 | 56 | 5 | 4 | 123 | 46 |
| K6613 | Kenya | Africa | 2000 | invasive, NOS | emm218.1 | 292 | 38 | 45 | 3 | 16 | 44 | 44 | 53 | 61 | 54 | 2 | 12 | 3 |
| SP10LAU | Lebanon | Asia | 2011 | pharyngitis and/or tonsillitis | emm118.0 | 167 | 11 | 32 | 52 | 1 | 67 | 13 | 118 | 135 | 1 | 2 | 83 | 93 |
| SP1LAU | Lebanon | Asia | 2010 | pharyngitis and/or tonsillitis | emm12.0 | 36 | 14 | 3 | 3 | 3 | 2 | 3 | 52 | 59 | 24 | 12 | 38 | 3 |
| SP2LAU | Lebanon | Asia | 2010 | skin/soft tissue infection, NOS | emm108.0 | 304 | 11 | 109 | 10 | 20 | 15 | 16 | 17 | 13 | 59 | 2 | 16 | 31 |
| SP3LAU | Lebanon | Asia | 2010 | pharyngitis and/or tonsillitis | emm89.0 | 101 | 2 | 2 | 2 | 2 | 2 | 2 | 2 | 2 | 2 | 2 | 2 | 2 |
| SP4LAU | Lebanon | Asia | 2010 | pharyngitis and/or tonsillitis | emm28.0 | 52 | 3 | 3 | 3 | 3 | 2 | 3 | 3 | 4 | 1 | 1 | 3 | 3 |
| SP5LAU | Lebanon | Asia | 2011 | skin/soft tissue infection, NOS | emm1.0 | 28 | 1 | 1 | 1 | 1 | 1 | 1 | 1 | 1 | 1 | 1 | 1 | 1 |
| SP6LAU | Lebanon | Asia | 2011 | pharyngitis and/or tonsillitis | emm89.0 | 101 | 2 | 2 | 2 | 2 | 2 | 2 | 2 | 2 | 2 | 2 | 2 | 2 |
| SP7LAU | Lebanon | Asia | 2011 | pharyngitis and/or tonsillitis | emm22.0 | 46 | 7 | 149 | 5 | 5 | 4 | 5 | 5 | 233 | 4 | 3 | 5 | 5 |
| SP8LAU | Lebanon | Asia | 2011 | pharyngitis and/or tonsillitis | emm85.0 | 109 | 10 | 10 | 4 | 9 | 9 | 11 | 72 | 11 | 7 | 3 | 10 | 10 |
| HSC5 | Unknown | | ND | ND | emm14.3 | 84 | 11 | 5 | 10 | 8 | 15 | 16 | 184 | 13 | 1 | 2 | 6 | 20 |
| M23ND | Unknown | | 1965 | invasive, NOS | emm23.0 | 160 | 120 | 150 | 121 | 54 | 106 | 62 | 192 | 234 | 11 | 4 | 35 | 155 |
| GAS13507 | New Zealand | Oceania | 2013 | ARF | emm25.0 | 191 | 30 | 35 | 26 | 28 | 35 | 34 | 42 | 45 | 11 | 2 | 12 | 40 |
| 15GA0017 | New Zealand | Oceania | 2015 | ARF | emm41.2 | 579 | 11 | 5 | 10 | 51 | 15 | 16 | 77 | 99 | 1 | 2 | 6 | 20 |

| | | | | | | | | | | | | | | | | | | |
|---|---|---|---|---|---|---|---|---|---|---|---|---|---|---|---|---|---|---|
| 14GA0838 | New Zealand | Oceania | 2014 | skin/soft tissue infection, NOS | emm81.0 | 330 | 62 | 75 | 45 | 52 | 5 | 6 | 45 | 106 | 24 | 17 | 66 | 61 |
| GAS14243 | New Zealand | Oceania | 2014 | ARF | emm12.0 | 36 | 14 | 3 | 3 | 3 | 2 | 3 | 52 | 59 | 24 | 12 | 38 | 3 |
| GAS131474 | New Zealand | Oceania | 2013 | pharyngitis and/or tonsillitis | emm53.0 | 11 | 121 | 5 | 10 | 80 | 15 | 54 | 17 | 84 | 1 | 2 | 6 | 20 |
| GAS131267 | New Zealand | Oceania | 2013 | pharyngitis and/or tonsillitis | emm74.0 | 120 | 28 | 13 | 3 | 41 | 50 | 53 | 123 | 82 | 11 | 4 | 59 | 70 |
| GAS131159 | New Zealand | Oceania | 2013 | ARF | emm70.0 | 1003 | 16 | 16 | 3 | 16 | 17 | 7 | 193 | 22 | 1 | 2 | 18 | 3 |
| GAS13502 | New Zealand | Oceania | 2013 | invasive, NOS | emm116.1 | 227 | 28 | 105 | 32 | 16 | 111 | 63 | 87 | 235 | 51 | 4 | 11 | 77 |
| 15GA0011 | New Zealand | Oceania | 2015 | invasive, NOS | emm41.2 | 579 | 11 | 5 | 10 | 51 | 15 | 16 | 77 | 99 | 1 | 2 | 6 | 20 |
| 14GA0832 | New Zealand | Oceania | 2014 | invasive, NOS | emm81.0 | 330 | 62 | 75 | 45 | 52 | 5 | 6 | 45 | 106 | 24 | 17 | 66 | 61 |
| GAS14211 | New Zealand | Oceania | 2014 | invasive, NOS | emm116.1 | 227 | 28 | 105 | 32 | 16 | 111 | 63 | 87 | 154 | 51 | 4 | 11 | 77 |
| GAS131472 | New Zealand | Oceania | 2013 | pharyngitis and/or tonsillitis | emm100.0 | 119 | 11 | 11 | 7 | 10 | 10 | 7 | 37 | 39 | 19 | 2 | 30 | 31 |
| GAS131255 | New Zealand | Oceania | 2013 | pharyngitis and/or tonsillitis | emm58.2 | 176 | 14 | 13 | 10 | 8 | 13 | 13 | 194 | 47 | 10 | 2 | 34 | 31 |
| GAS131153 | New Zealand | Oceania | 2013 | pharyngitis and/or tonsillitis | emm92.0 | 82 | 17 | 4 | 3 | 18 | 19 | 19 | 22 | 24 | 13 | 2 | 20 | 22 |
| GAS13480 | New Zealand | Oceania | 2013 | ARF | emm116.1 | 227 | 28 | 105 | 32 | 16 | 111 | 63 | 87 | 154 | 51 | 4 | 11 | 77 |
| 14GA0830 | New Zealand | Oceania | 2014 | ND | emm81.0 | 330 | 62 | 75 | 45 | 52 | 5 | 6 | 45 | 106 | 24 | 17 | 66 | 61 |
| 14GA1137 | New Zealand | Oceania | 2014 | ARF | emm41.2 | 12 | 11 | 5 | 10 | 15 | 15 | 16 | 17 | 13 | 1 | 2 | 16 | 18 |
| GAS131465 | New Zealand | Oceania | 2013 | ARF | emm53.0 | 11 | 121 | 5 | 10 | 8 | 15 | 54 | 17 | 84 | 1 | 2 | 6 | 20 |
| GAS131252 | New Zealand | Oceania | 2013 | ARF | emm74.0 | 120 | 28 | 13 | 3 | 41 | 50 | 53 | 123 | 82 | 11 | 4 | 59 | 70 |
| GAS131142 | New Zealand | Oceania | 2013 | pharyngitis and/or tonsillitis | emm76.0 | 378 | 122 | 3 | 3 | 3 | 2 | 3 | 52 | 59 | 24 | 12 | 38 | 3 |
| GAS13476 | New Zealand | Oceania | 2013 | ARF | emm19.4 | 616 | 50 | 32 | 10 | 1 | 6 | 55 | 78 | 13 | 1 | 2 | 6 | 106 |
| 14GA0829 | New Zealand | Oceania | 2014 | ND | emm81.0 | 330 | 62 | 75 | 45 | 52 | 5 | 6 | 45 | 106 | 24 | 17 | 66 | 61 |
| 14GA1136 | New Zealand | Oceania | 2014 | ARF | emm41.2 | 579 | 11 | 5 | 10 | 51 | 15 | 16 | 77 | 99 | 1 | 2 | 6 | 20 |
| GAS131460 | New Zealand | Oceania | 2013 | ARF | emm76.0 | 378 | 122 | 3 | 3 | 3 | 2 | 3 | 52 | 59 | 24 | 12 | 38 | 3 |
| GAS131245 | New Zealand | Oceania | 2013 | pharyngitis and/or tonsillitis | emm89.0 | 101 | 123 | 2 | 2 | 58 | 2 | 2 | 2 | 2 | 2 | 2 | 2 | 2 |
| GAS131141 | New Zealand | Oceania | 2013 | pharyngitis and/or tonsillitis | emm25.0 | 191 | 30 | 35 | 26 | 28 | 35 | 34 | 42 | 45 | 11 | 2 | 12 | 40 |
| GAS13475 | New Zealand | Oceania | 2013 | ARF | emm197.0 | 998 | 11 | 151 | 10 | 8 | 165 | 16 | 17 | 236 | 1 | 2 | 6 | 20 |
| 14GA0828 | New Zealand | Oceania | 2014 | skin/soft tissue infection, NOS | emm81.0 | 330 | 62 | 75 | 45 | 52 | 5 | 6 | 45 | 106 | 24 | 17 | 66 | 61 |
| 14GA0305 | New Zealand | Oceania | 2014 | invasive, NOS | emm103.0 | 876 | 82 | 25 | 3 | 64 | 107 | 13 | 26 | 29 | 8 | 3 | 25 | 30 |
| 14GA1101 | New Zealand | Oceania | 2014 | invasive, NOS | emm118.0 | 997 | 11 | 32 | 52 | 1 | 67 | 13 | 118 | 237 | 1 | 2 | 83 | 93 |
| GAS131421 | New Zealand | Oceania | 2013 | invasive, NOS | emm74.0 | 120 | 28 | 13 | 75 | 41 | 50 | 53 | 123 | 82 | 11 | 4 | 59 | 70 |
| GAS131241 | New Zealand | Oceania | 2013 | ARF | emm58.2 | 176 | 14 | 13 | 10 | 8 | 13 | 13 | 194 | 47 | 10 | 2 | 34 | 31 |
| GAS131138 | New Zealand | Oceania | 2013 | invasive, NOS | emm74.0 | 120 | 28 | 13 | 3 | 41 | 50 | 53 | 123 | 82 | 11 | 4 | 59 | 70 |
| GAS13474 | New Zealand | Oceania | 2013 | ARF | emm95.0 | 604 | 11 | 32 | 30 | 31 | 37 | 7 | 46 | 51 | 1 | 2 | 6 | 44 |
| 14GA0826 | New Zealand | Oceania | 2014 | invasive, NOS | emm81.0 | 330 | 62 | 75 | 45 | 52 | 5 | 6 | 45 | 106 | 24 | 17 | 66 | 61 |
| 14GA0287 | New Zealand | Oceania | 2014 | ARF | emm74.0 | 120 | 28 | 13 | 3 | 41 | 50 | 53 | 123 | 238 | 11 | 4 | 59 | 70 |
| GAS1464 | New Zealand | Oceania | 2014 | pharyngitis and/or tonsillitis | emm58.7 | 1004 | 14 | 13 | 10 | 8 | 13 | 13 | 195 | 239 | 10 | 2 | 34 | 31 |
| GAS131413 | New Zealand | Oceania | 2013 | pharyngitis and/or tonsillitis | emm76.0 | 378 | 122 | 3 | 3 | 3 | 2 | 3 | 52 | 59 | 24 | 12 | 38 | 3 |
| GAS131223 | New Zealand | Oceania | 2013 | ARF | emm74.0 | 120 | 28 | 13 | 3 | 41 | 50 | 53 | 123 | 82 | 11 | 4 | 59 | 70 |
| GAS131111 | New Zealand | Oceania | 2013 | pharyngitis and/or tonsillitis | emm74.0 | 120 | 28 | 13 | 3 | 41 | 50 | 53 | 123 | 82 | 11 | 4 | 59 | 70 |
| GAS13472 | New Zealand | Oceania | 2013 | invasive, NOS | emm70.0 | 1003 | 16 | 16 | 3 | 16 | 17 | 7 | 193 | 22 | 1 | 2 | 18 | 3 |
| 14GA0809 | New Zealand | Oceania | 2014 | ARF | emm59.1 | 172 | 71 | 84 | 24 | 2 | 93 | 68 | 107 | 122 | 18 | 3 | 75 | 85 |
| 14GA0286 | New Zealand | Oceania | 2014 | ARF | emm103.0 | 876 | 82 | 25 | 3 | 64 | 107 | 13 | 26 | 29 | 8 | 3 | 25 | 30 |
| 14GA1044 | New Zealand | Oceania | 2014 | invasive, NOS | emm86.2 | 963 | 11 | 32 | 10 | 20 | 15 | 16 | 128 | 13 | 1 | 2 | 6 | 100 |
| GAS131348 | New Zealand | Oceania | 2013 | invasive, NOS | emm53.0 | 11 | 49 | 5 | 10 | 8 | 15 | 54 | 17 | 84 | 1 | 2 | 6 | 20 |
| GAS131222 | New Zealand | Oceania | 2013 | pharyngitis and/or tonsillitis | emm81.0 | 330 | 62 | 75 | 45 | 52 | 5 | 6 | 45 | 106 | 24 | 17 | 66 | 61 |
| GAS131103 | New Zealand | Oceania | 2013 | invasive, NOS | emm65.0 | 111 | 22 | 22 | 17 | 38 | 166 | 43 | 28 | 65 | 1 | 1 | 1 | 27 |
| GAS13450 | New Zealand | Oceania | 2013 | invasive, NOS | emm19.4 | 616 | 50 | 32 | 10 | 1 | 6 | 55 | 78 | 13 | 1 | 2 | 6 | 106 |
| 14GA0958 | New Zealand | Oceania | 2014 | invasive, NOS | emm90.5 | 184 | 20 | 20 | 14 | 19 | 2 | 23 | 25 | 28 | 15 | 3 | 22 | 25 |
| 14GA0794 | New Zealand | Oceania | 2014 | pharyngitis and/or tonsillitis | emm74.0 | 120 | 28 | 13 | 3 | 41 | 50 | 53 | 123 | 82 | 11 | 4 | 59 | 70 |

| | | | | | | | | | | | | | | | | | |
|---|---|---|---|---|---|---|---|---|---|---|---|---|---|---|---|---|---|
| GAS14274 | New Zealand | Oceania | 2014 | invasive, NOS | emm95.0 | 604 | 11 | 32 | 30 | 31 | 37 | 7 | 46 | 51 | 1 | 2 | 6 | 44 |
| GAS1441 | New Zealand | Oceania | 2014 | invasive, NOS | emm12.0 | 36 | 14 | 3 | 3 | 3 | 2 | 3 | 52 | 59 | 24 | 12 | 38 | 3 |
| GAS131322 | New Zealand | Oceania | 2013 | invasive, NOS | emm108.1 | 14 | 28 | 87 | 10 | 1 | 6 | 63 | 78 | 240 | 1 | 2 | 6 | 31 |
| GAS131204 | New Zealand | Oceania | 2013 | invasive, NOS | emm81.0 | 330 | 62 | 75 | 45 | 52 | 5 | 6 | 45 | 106 | 24 | 17 | 66 | 61 |
| GAS131015 | New Zealand | Oceania | 2013 | pharyngitis and/or tonsillitis | emm65.0 | 111 | 22 | 22 | 17 | 38 | 23 | 43 | 28 | 65 | 1 | 1 | 1 | 27 |
| GAS13432 | New Zealand | Oceania | 2013 | pharyngitis and/or tonsillitis | ND | 616 | 50 | 32 | 10 | 1 | 6 | 55 | 78 | 13 | 1 | 2 | 6 | 106 |
| GAS13426 | New Zealand | Oceania | 2013 | pharyngitis and/or tonsillitis | emm116.1 | 227 | 28 | 105 | 32 | 16 | 111 | 63 | 87 | 154 | 51 | 4 | 11 | 77 |
| GAS13118 | New Zealand | Oceania | 2013 | pharyngitis and/or tonsillitis | emm74.0 | 120 | 28 | 13 | 3 | 41 | 50 | 53 | 123 | 82 | 11 | 4 | 59 | 70 |
| GAS12394 | New Zealand | Oceania | 2012 | invasive, NOS | emm41.2 | 579 | 11 | 5 | 10 | 51 | 15 | 16 | 77 | 99 | 1 | 2 | 6 | 20 |
| GAS12235 | New Zealand | Oceania | 2012 | invasive, NOS | emm89.0 | 101 | 2 | 2 | 2 | 58 | 2 | 2 | 2 | 2 | 2 | 2 | 2 | 2 |
| GAS12118 | New Zealand | Oceania | 2012 | pharyngitis and/or tonsillitis | emm89.0 | 101 | 2 | 2 | 2 | 2 | 2 | 2 | 2 | 2 | 2 | 2 | 2 | 2 |
| GAS11549 | New Zealand | Oceania | 2011 | ARF | emm71.0 | 981 | 19 | 20 | 13 | 19 | 2 | 22 | 3 | 27 | 1 | 1 | 4 | 24 |
| GAS11291 | New Zealand | Oceania | 2011 | ARF | emm11.0 | ND | 1 | 6 | 76 | 1 | 104 | 20 | 13 | 25 | 8 | 3 | 21 | 23 |
| GAS11117 | New Zealand | Oceania | 2011 | ARF | emm68.0 | 993 | 81 | 103 | 33 | 63 | 105 | 7 | 130 | 150 | 34 | 3 | 90 | 104 |
| GAS13409 | New Zealand | Oceania | 2013 | ARF | emm19.4 | 616 | 50 | 32 | 10 | 1 | 6 | 55 | 78 | 13 | 1 | 2 | 6 | 106 |
| GAS1396 | New Zealand | Oceania | 2013 | pharyngitis and/or tonsillitis | emm116.1 | 227 | 28 | 105 | 32 | 16 | 111 | 63 | 87 | 241 | 51 | 4 | 11 | 77 |
| GAS12392 | New Zealand | Oceania | 2012 | pharyngitis and/or tonsillitis | emm59.1 | 172 | 71 | 84 | 24 | 2 | 93 | 68 | 107 | 122 | 18 | 3 | 75 | 85 |
| GAS12233 | New Zealand | Oceania | 2012 | invasive, NOS | emm1.0 | 28 | 1 | 1 | 1 | 1 | 1 | 1 | 1 | 3 | 1 | 1 | 1 | 1 |
| GAS12107 | New Zealand | Oceania | 2012 | ARF | emm218.1 | 292 | 38 | 45 | 3 | 16 | 44 | 93 | 53 | 61 | 54 | 2 | 12 | 3 |
| 14GA0862 | New Zealand | Oceania | 2014 | invasive, NOS | emm85.0 | 109 | 10 | 10 | 4 | 9 | 9 | 11 | 72 | 11 | 7 | 3 | 10 | 10 |
| 14GA0793 | New Zealand | Oceania | 2014 | pharyngitis and/or tonsillitis | emm59.1 | 172 | 71 | 84 | 24 | 2 | 93 | 68 | 107 | 122 | 18 | 3 | 75 | 85 |
| GAS14267 | New Zealand | Oceania | 2014 | invasive, NOS | emm54.1 | 990 | 11 | 5 | 10 | 20 | 15 | 16 | 17 | 13 | 1 | 2 | 16 | 31 |
| GAS1414 | New Zealand | Oceania | 2014 | pharyngitis and/or tonsillitis | emm90.5 | 184 | 20 | 20 | 14 | 19 | 2 | 23 | 25 | 28 | 15 | 3 | 22 | 25 |
| GAS131318 | New Zealand | Oceania | 2013 | ARF | emm108.1 | 14 | 28 | 87 | 10 | 1 | 6 | 63 | 78 | 240 | 1 | 2 | 6 | 31 |
| GAS131202 | New Zealand | Oceania | 2013 | ARF | emm232.0 | 902 | 14 | 37 | 31 | 3 | 2 | 38 | 47 | 242 | 1 | 30 | 58 | 25 |
| GAS13637 | New Zealand | Oceania | 2013 | ARF | emm65.4 | 129 | 11 | 66 | 40 | 35 | 102 | 76 | 44 | 145 | 1 | 2 | 87 | 31 |
| 14GA0857 | New Zealand | Oceania | 2014 | invasive, NOS | emm81.0 | 330 | 62 | 75 | 45 | 52 | 5 | 6 | 45 | 106 | 24 | 17 | 66 | 61 |
| 14GA0788 | New Zealand | Oceania | 2014 | ARF | emm85.0 | 109 | 10 | 10 | 4 | 9 | 9 | 11 | 72 | 11 | 7 | 3 | 10 | 10 |
| GAS14266 | New Zealand | Oceania | 2014 | pharyngitis and/or tonsillitis | emm118.0 | 997 | 11 | 32 | 52 | 1 | 67 | 13 | 118 | 237 | 1 | 2 | 83 | 93 |
| GAS146 | New Zealand | Oceania | 2014 | pharyngitis and/or tonsillitis | emm12.0 | 36 | 14 | 3 | 3 | 3 | 2 | 3 | 52 | 243 | 24 | 12 | 38 | 156 |
| GAS131286 | New Zealand | Oceania | 2013 | invasive, NOS | emm89.0 | 101 | 2 | 2 | 2 | 2 | 2 | 2 | 2 | 2 | 2 | 2 | 2 | 2 |
| GAS131175 | New Zealand | Oceania | 2013 | invasive, NOS | emm74.0 | 120 | 28 | 13 | 3 | 41 | 50 | 53 | 123 | 82 | 11 | 4 | 59 | 70 |
| GAS13599 | New Zealand | Oceania | 2013 | invasive, NOS | emm41.2 | 579 | 11 | 5 | 10 | 51 | 15 | 16 | 77 | 99 | 1 | 2 | 6 | 20 |
| 15GAS0097 | New Zealand | Oceania | 2015 | ND | emm95.0 | 604 | 11 | 32 | 30 | 31 | 37 | 7 | 46 | 51 | 1 | 2 | 6 | 44 |
| 14GA0844 | New Zealand | Oceania | 2014 | pharyngitis and/or tonsillitis | emm91.0 | 12 | 11 | 5 | 10 | 15 | 15 | 16 | 17 | 13 | 1 | 2 | 16 | 18 |
| GAS14265 | New Zealand | Oceania | 2014 | ARF | emm86.2 | 963 | 11 | 32 | 10 | 20 | 15 | 16 | 128 | 13 | 1 | 2 | 6 | 100 |
| 14GA1005 | New Zealand | Oceania | 2014 | pharyngitis and/or tonsillitis | emm1.0 | 28 | 1 | 1 | 1 | 1 | 1 | 1 | 1 | 3 | 1 | 1 | 1 | 1 |
| GAS131282 | New Zealand | Oceania | 2013 | ARF | emm89.0 | 101 | 2 | 2 | 2 | 2 | 2 | 2 | 2 | 2 | 2 | 2 | 2 | 2 |
| GAS131172 | New Zealand | Oceania | 2013 | ARF | emm74.0 | 120 | 28 | 13 | 3 | 41 | 50 | 53 | 123 | 82 | 11 | 4 | 59 | 70 |
| GAS13589 | New Zealand | Oceania | 2013 | ARF | emm41.2 | 579 | 11 | 5 | 10 | 51 | 15 | 16 | 77 | 99 | 1 | 2 | 6 | 20 |
| 15GAS0096 | New Zealand | Oceania | 2015 | ND | emm90.5 | 184 | 20 | 20 | 14 | 19 | 2 | 23 | 25 | 28 | 15 | 3 | 22 | 25 |
| 14GA0839 | New Zealand | Oceania | 2014 | ND | emm85.0 | 109 | 10 | 10 | 4 | 9 | 9 | 11 | 72 | 244 | 7 | 3 | 10 | 10 |
| GAS14246 | New Zealand | Oceania | 2014 | invasive, NOS | emm86.2 | 963 | 11 | 32 | 10 | 20 | 15 | 16 | 128 | 13 | 1 | 2 | 6 | 100 |
| GAS131487 | New Zealand | Oceania | 2013 | ARF | emm74.0 | 1011 | 11 | 13 | 122 | 81 | 167 | 13 | 196 | 42 | 31 | 1 | 51 | 4 |
| GAS131275 | New Zealand | Oceania | 2013 | pharyngitis and/or tonsillitis | emm77.0 | 63 | 23 | 24 | 19 | 1 | 26 | 28 | 30 | 33 | 8 | 3 | 24 | 29 |
| GAS131166 | New Zealand | Oceania | 2013 | invasive, NOS | emm58.2 | 176 | 14 | 13 | 10 | 8 | 13 | 13 | 194 | 47 | 10 | 2 | 34 | 31 |
| GAS11493 | New Zealand | Oceania | 2011 | invasive, NOS | emm19.4 | 616 | 50 | 32 | 10 | 1 | 6 | 55 | 78 | 13 | 1 | 2 | 6 | 106 |
| GAS11282 | New Zealand | Oceania | 2011 | invasive, NOS | emm100.5 | 119 | 11 | 11 | 7 | 10 | 10 | 7 | 37 | 39 | 19 | 2 | 30 | 31 |

| | | | | | | | | | | | | | | | | | | |
|---|---|---|---|---|---|---|---|---|---|---|---|---|---|---|---|---|---|---|
| GAS10411 | New Zealand | Oceania | 2010 | pharyngitis and/or tonsillitis | emm91.0 | 12 | 11 | 5 | 10 | 15 | 15 | 16 | 17 | 13 | 1 | 2 | 16 | 18 |
| GAS10213 | New Zealand | Oceania | 2010 | ARF | emm36.2 | 748 | 15 | 15 | 10 | 1 | 10 | 32 | 197 | 21 | 12 | 8 | 37 | 31 |
| GAS1073 | New Zealand | Oceania | 2010 | invasive, NOS | emm36.2 | 748 | 15 | 15 | 10 | 1 | 10 | 32 | 197 | 21 | 12 | 8 | 37 | 31 |
| GAS09398 | New Zealand | Oceania | 2009 | pharyngitis and/or tonsillitis | emm12.0 | 36 | 14 | 3 | 3 | 3 | 2 | 3 | 52 | 245 | 24 | 12 | 38 | 3 |
| GAS09190 | New Zealand | Oceania | 2009 | pharyngitis and/or tonsillitis | emm87.0 | 62 | 67 | 15 | 10 | 1 | 72 | 58 | 87 | 95 | 1 | 2 | 52 | 74 |
| GAS07167 | New Zealand | Oceania | 2007 | ARF | emm6.0 | 411 | 65 | 80 | 62 | 55 | 85 | 62 | 97 | 130 | 11 | 20 | 69 | 82 |
| GAS05161 | New Zealand | Oceania | 2005 | ARF | emm98.1 | 205 | 15 | 15 | 10 | 1 | 10 | 94 | 19 | 246 | 12 | 2 | 6 | 20 |
| GAS0387 | New Zealand | Oceania | 2003 | invasive, NOS | emm108.1 | 14 | 28 | 87 | 10 | 1 | 6 | 63 | 78 | 240 | 1 | 2 | 6 | 31 |
| GAS10367 | New Zealand | Oceania | 2010 | pharyngitis and/or tonsillitis | emm22.0 | 46 | 7 | 5 | 5 | 5 | 4 | 5 | 5 | 6 | 4 | 3 | 5 | 5 |
| GAS10210 | New Zealand | Oceania | 2010 | invasive, NOS | emm87.0 | 62 | 67 | 15 | 10 | 1 | 72 | 58 | 87 | 247 | 23 | 2 | 52 | 74 |
| GAS1186 | New Zealand | Oceania | 2011 | invasive, NOS | emm65.4 | 129 | 11 | 66 | 40 | 35 | 102 | 76 | 44 | 145 | 1 | 2 | 87 | 31 |
| GAS1050 | New Zealand | Oceania | 2010 | invasive, NOS | emm68.0 | 993 | 81 | 103 | 33 | 63 | 105 | 7 | 130 | 150 | 34 | 3 | 90 | 104 |
| GAS09379 | New Zealand | Oceania | 2009 | ARF | emm36.2 | 748 | 15 | 15 | 10 | 1 | 10 | 32 | 197 | 21 | 12 | 8 | 37 | 31 |
| GAS09155 | New Zealand | Oceania | 2009 | ARF | emm65.4 | 129 | 11 | 66 | 40 | 35 | 102 | 76 | 44 | 145 | 1 | 2 | 87 | 31 |
| GAS07147 | New Zealand | Oceania | 2007 | pharyngitis and/or tonsillitis | emm44.0 | 178 | 5 | 7 | 7 | 3 | 2 | 3 | 24 | 8 | 14 | 3 | 7 | 7 |
| GAS05154 | New Zealand | Oceania | 2005 | invasive, NOS | emm53.0 | 11 | 49 | 5 | 10 | 8 | 15 | 54 | 17 | 84 | 1 | 2 | 6 | 20 |
| GAS0371 | New Zealand | Oceania | 2003 | pharyngitis and/or tonsillitis | emm82.0 | 334 | 73 | 58 | 46 | 1 | 6 | 18 | 69 | 77 | 1 | 1 | 1 | 3 |
| GAS10358 | New Zealand | Oceania | 2010 | ARF | emm53.0 | 11 | 49 | 5 | 10 | 8 | 15 | 54 | 17 | 84 | 1 | 2 | 6 | 20 |
| GAS10197 | New Zealand | Oceania | 2010 | ARF | emm113.0 | 148 | 14 | 3 | 3 | 3 | 2 | 3 | 52 | 59 | 24 | 12 | 38 | 3 |
| GAS1047 | New Zealand | Oceania | 2010 | ARF | emm36.2 | 748 | 15 | 15 | 10 | 1 | 10 | 32 | 197 | 21 | 12 | 8 | 37 | 31 |
| GAS09344 | New Zealand | Oceania | 2009 | ARF | emm106.0 | 1012 | 14 | 152 | 21 | 25 | 106 | 19 | 34 | 36 | 15 | 3 | 124 | 34 |
| GAS13376 | New Zealand | Oceania | 2013 | invasive, NOS | emm49.0 | 433 | 4 | 8 | 4 | 11 | 14 | 14 | 16 | 18 | 9 | 2 | 12 | 13 |
| GAS0945 | New Zealand | Oceania | 2009 | pharyngitis and/or tonsillitis | emm6.0 | 382 | 65 | 80 | 62 | 55 | 85 | 62 | 97 | 130 | 11 | 20 | 69 | 82 |
| GAS06308 | New Zealand | Oceania | 2006 | ARF | emm54.1 | 990 | 11 | 5 | 10 | 20 | 15 | 16 | 17 | 13 | 1 | 2 | 16 | 31 |
| GAS05151 | New Zealand | Oceania | 2005 | ARF | emm89.0 | 142 | 4 | 8 | 3 | 11 | 14 | 14 | 16 | 18 | 9 | 2 | 12 | 13 |
| GAS0370 | New Zealand | Oceania | 2003 | invasive, NOS | emm82.0 | 334 | 73 | 58 | 46 | 1 | 6 | 18 | 69 | 77 | 1 | 1 | 1 | 3 |
| GAS10346 | New Zealand | Oceania | 2010 | invasive, NOS | emm71.0 | 318 | 19 | 20 | 13 | 19 | 2 | 22 | 3 | 27 | 1 | 1 | 4 | 24 |
| GAS10179 | New Zealand | Oceania | 2010 | ARF | emm91.0 | 12 | 11 | 5 | 10 | 15 | 15 | 16 | 17 | 13 | 1 | 2 | 16 | 18 |
| GAS1041 | New Zealand | Oceania | 2010 | invasive, NOS | emm91.0 | 12 | 11 | 5 | 10 | 15 | 15 | 16 | 17 | 13 | 1 | 2 | 16 | 18 |
| GAS09330 | New Zealand | Oceania | 2009 | ARF | emm49.0 | 433 | 4 | 8 | 4 | 11 | 14 | 14 | 16 | 18 | 9 | 2 | 12 | 13 |
| GAS0944 | New Zealand | Oceania | 2009 | ARF | ND | 382 | 65 | 80 | 62 | 55 | 85 | 62 | 97 | 130 | 11 | 20 | 69 | 82 |
| GAS06262 | New Zealand | Oceania | 2006 | invasive, NOS | emm54.1 | 990 | 11 | 5 | 10 | 20 | 15 | 16 | 17 | 13 | 1 | 2 | 16 | 31 |
| GAS1376 | New Zealand | Oceania | 2013 | pharyngitis and/or tonsillitis | emm70.0 | 1003 | 16 | 16 | 3 | 16 | 17 | 7 | 193 | 22 | 1 | 2 | 18 | 3 |
| GAS05150 | New Zealand | Oceania | 2005 | ARF | emm53.0 | 11 | 49 | 5 | 10 | 8 | 15 | 54 | 17 | 84 | 1 | 2 | 6 | 20 |
| GAS0361 | New Zealand | Oceania | 2003 | ARF | emm53.0 | 11 | 49 | 5 | 10 | 8 | 15 | 54 | 17 | 84 | 1 | 2 | 6 | 20 |
| GAS10339 | New Zealand | Oceania | 2010 | invasive, NOS | emm53.0 | 11 | 121 | 5 | 10 | 8 | 15 | 54 | 17 | 84 | 1 | 2 | 6 | 20 |
| GAS10157 | New Zealand | Oceania | 2010 | invasive, NOS | emm232.0 | 902 | 14 | 37 | 31 | 3 | 2 | 38 | 47 | 242 | 1 | 2 | 58 | 25 |
| GAS1023 | New Zealand | Oceania | 2010 | invasive, NOS | emm113.0 | 148 | 14 | 3 | 3 | 3 | 2 | 3 | 52 | 59 | 24 | 12 | 38 | 3 |
| GAS09267 | New Zealand | Oceania | 2009 | ARF | emm92.0 | 82 | 17 | 4 | 3 | 18 | 19 | 19 | 22 | 24 | 13 | 2 | 20 | 22 |
| GAS06220 | New Zealand | Oceania | 2006 | invasive, NOS | emm1.0 | 28 | 1 | 1 | 1 | 1 | 1 | 1 | 1 | 1 | 1 | 1 | 1 | 1 |
| GAS05134 | New Zealand | Oceania | 2005 | ARF | emm1.0 | 28 | 1 | 1 | 1 | 1 | 1 | 1 | 1 | 1 | 1 | 1 | 1 | 1 |
| GAS0359 | New Zealand | Oceania | 2003 | ARF | emm82.0 | 334 | 73 | 58 | 46 | 1 | 6 | 18 | 69 | 77 | 1 | 1 | 1 | 3 |
| GAS12332 | New Zealand | Oceania | 2012 | pharyngitis and/or tonsillitis | emm33.0 | 3 | 11 | 11 | 7 | 10 | 35 | 36 | 85 | 79 | 1 | 2 | 6 | 20 |
| GAS10325 | New Zealand | Oceania | 2010 | ARF | emm105.0 | 151 | 6 | 6 | 6 | 6 | 5 | 6 | 6 | 12 | 5 | 4 | 6 | 11 |
| GAS10144 | New Zealand | Oceania | 2010 | invasive, NOS | emm6.0 | 411 | 65 | 80 | 62 | 55 | 85 | 62 | 97 | 130 | 11 | 20 | 69 | 82 |
| GAS1019 | New Zealand | Oceania | 2010 | ARF | emm22.0 | 202 | 33 | 11 | 7 | 10 | 35 | 36 | 44 | 49 | 19 | 2 | 30 | 43 |
| GAS09257 | New Zealand | Oceania | 2009 | ARF | emm39.4 | 268 | 1 | 60 | 49 | 45 | 61 | 19 | 13 | 83 | 34 | 3 | 61 | 29 |
| GAS0925 | New Zealand | Oceania | 2009 | ARF | emm74.0 | 120 | 28 | 13 | 3 | 41 | 50 | 53 | 123 | 82 | 11 | 4 | 59 | 70 |

| | | | | | | | | | | | | | | | | | | |
|---|---|---|---|---|---|---|---|---|---|---|---|---|---|---|---|---|---|---|
| GAS06216 | New Zealand | Oceania | 2006 | pharyngitis and/or tonsillitis | emm1.0 | 28 | 1 | 1 | 1 | 1 | 1 | 1 | 1 | 1 | 1 | 1 | 1 | 1 |
| GAS0592 | New Zealand | Oceania | 2005 | invasive, NOS | emm44.0 | 178 | 5 | 7 | 7 | 3 | 2 | 3 | 24 | 8 | 14 | 3 | 7 | 7 |
| GAS0354 | New Zealand | Oceania | 2003 | invasive, NOS | emm101.0 | 182 | 15 | 31 | 10 | 20 | 15 | 31 | 38 | 41 | 1 | 3 | 32 | 37 |
| GAS10275 | New Zealand | Oceania | 2010 | pharyngitis and/or tonsillitis | emm74.0 | 120 | 28 | 13 | 3 | 41 | 50 | 53 | 123 | 82 | 11 | 4 | 59 | 70 |
| GAS10141 | New Zealand | Oceania | 2010 | ARF | emm232.0 | 967 | 14 | 37 | 31 | 3 | 2 | 38 | 47 | 242 | 1 | 2 | 58 | 25 |
| GAS12183 | New Zealand | Oceania | 2012 | ARF | emm89.14 | 380 | 14 | 49 | 20 | 16 | 112 | 52 | 136 | 248 | 33 | 11 | 57 | 25 |
| GAS09441 | New Zealand | Oceania | 2009 | pharyngitis and/or tonsillitis | emm113.0 | 148 | 14 | 3 | 3 | 3 | 2 | 3 | 52 | 59 | 24 | 12 | 38 | 3 |
| GAS09227 | New Zealand | Oceania | 2009 | invasive, NOS | emm92.0 | 82 | 17 | 4 | 3 | 18 | 19 | 19 | 22 | 24 | 13 | 2 | 20 | 22 |
| GAS0919 | New Zealand | Oceania | 2009 | invasive, NOS | ND | 382 | 65 | 80 | 62 | 55 | 85 | 62 | 97 | 130 | 11 | 20 | 69 | 82 |
| GAS06166 | New Zealand | Oceania | 2006 | pharyngitis and/or tonsillitis | emm12.0 | 36 | 14 | 3 | 3 | 3 | 2 | 3 | 52 | 59 | 24 | 12 | 38 | 3 |
| GAS03202 | New Zealand | Oceania | 2003 | pharyngitis and/or tonsillitis | emm53.0 | 11 | 49 | 5 | 10 | 8 | 15 | 54 | 17 | 84 | 1 | 2 | 6 | 20 |
| GAS0327 | New Zealand | Oceania | 2003 | invasive, NOS | emm3.1 | 15 | 15 | 15 | 10 | 1 | 10 | 32 | 111 | 129 | 1 | 2 | 6 | 31 |
| GAS10262 | New Zealand | Oceania | 2010 | invasive, NOS | emm22.0 | 992 | 11 | 11 | 7 | 10 | 35 | 95 | 198 | 49 | 28 | 1 | 30 | 43 |
| GAS10113 | New Zealand | Oceania | 2010 | pharyngitis and/or tonsillitis | emm74.0 | 120 | 28 | 13 | 75 | 41 | 50 | 53 | 123 | 82 | 11 | 4 | 59 | 70 |
| GAS09438 | New Zealand | Oceania | 2009 | invasive, NOS | emm36.2 | 748 | 15 | 15 | 10 | 1 | 10 | 32 | 197 | 21 | 12 | 8 | 37 | 31 |
| GAS09226 | New Zealand | Oceania | 2009 | ARF | emm59.1 | 172 | 71 | 84 | 24 | 2 | 93 | 68 | 107 | 122 | 18 | 3 | 75 | 85 |
| GAS1291 | New Zealand | Oceania | 2012 | ARF | emm77.0 | 572 | 15 | 99 | 3 | 39 | 56 | 13 | 68 | 146 | 32 | 3 | 54 | 157 |
| GAS0917 | New Zealand | Oceania | 2009 | pharyngitis and/or tonsillitis | emm6.0 | 382 | 65 | 80 | 62 | 55 | 85 | 62 | 97 | 130 | 11 | 20 | 69 | 82 |
| GAS0656 | New Zealand | Oceania | 2006 | invasive, NOS | emm65.0 | 111 | 22 | 22 | 17 | 38 | 43 | 43 | 28 | 65 | 1 | 1 | 1 | 27 |
| GAS03188 | New Zealand | Oceania | 2003 | pharyngitis and/or tonsillitis | emm89.0 | 142 | 4 | 8 | 3 | 11 | 14 | 14 | 16 | 18 | 9 | 2 | 12 | 13 |
| GAS0320 | New Zealand | Oceania | 2003 | ARF | emm3.1 | 15 | 15 | 15 | 10 | 1 | 10 | 32 | 111 | 129 | 1 | 2 | 6 | 31 |
| GAS10250 | New Zealand | Oceania | 2010 | ARF | emm87.0 | 541 | 55 | 68 | 10 | 1 | 72 | 58 | 87 | 95 | 1 | 2 | 52 | 74 |
| GAS10112 | New Zealand | Oceania | 2010 | ARF | emm71.0 | 318 | 19 | 20 | 13 | 19 | 2 | 22 | 3 | 27 | 1 | 1 | 4 | 24 |
| GAS09437 | New Zealand | Oceania | 2009 | ARF | emm12.0 | 36 | 14 | 3 | 3 | 3 | 2 | 3 | 52 | 59 | 24 | 12 | 38 | 3 |
| GAS09219 | New Zealand | Oceania | 2009 | invasive, NOS | emm59.1 | 172 | 71 | 84 | 24 | 2 | 93 | 68 | 107 | 122 | 18 | 3 | 75 | 85 |
| GAS08308 | New Zealand | Oceania | 2008 | pharyngitis and/or tonsillitis | emm6.0 | 382 | 65 | 80 | 62 | 55 | 85 | 62 | 97 | 130 | 11 | 20 | 69 | 82 |
| GAS0617 | New Zealand | Oceania | 2006 | invasive, NOS | emm12.0 | 36 | 14 | 3 | 3 | 3 | 62 | 3 | 52 | 59 | 24 | 12 | 38 | 3 |
| GAS11453 | New Zealand | Oceania | 2011 | invasive, NOS | emm36.2 | 748 | 15 | 15 | 10 | 1 | 10 | 32 | 197 | 21 | 12 | 8 | 37 | 31 |
| GAS0317 | New Zealand | Oceania | 2003 | pharyngitis and/or tonsillitis | emm3.1 | 15 | 15 | 15 | 10 | 1 | 10 | 32 | 111 | 129 | 1 | 2 | 6 | 31 |
| GAS10245 | New Zealand | Oceania | 2010 | invasive, NOS | emm90.5 | 184 | 20 | 20 | 14 | 19 | 2 | 23 | 25 | 28 | 15 | 3 | 22 | 25 |
| GAS10104 | New Zealand | Oceania | 2010 | pharyngitis and/or tonsillitis | emm90.5 | 184 | 20 | 20 | 14 | 19 | 2 | 23 | 25 | 28 | 15 | 3 | 22 | 25 |
| GAS09426 | New Zealand | Oceania | 2009 | ARF | emm101.0 | 182 | 15 | 31 | 10 | 20 | 15 | 31 | 38 | 41 | 1 | 3 | 32 | 37 |
| GAS09205 | New Zealand | Oceania | 2009 | ARF | emm108.1 | 14 | 28 | 87 | 10 | 1 | 6 | 63 | 78 | 240 | 1 | 2 | 6 | 31 |
| GAS08119 | New Zealand | Oceania | 2008 | pharyngitis and/or tonsillitis | emm53.0 | 11 | 49 | 5 | 10 | 8 | 15 | 54 | 17 | 84 | 1 | 2 | 6 | 20 |
| GAS0615 | New Zealand | Oceania | 2006 | pharyngitis and/or tonsillitis | emm53.0 | 11 | 121 | 5 | 10 | 8 | 15 | 54 | 17 | 84 | 1 | 2 | 6 | 20 |
| GAS03109 | New Zealand | Oceania | 2003 | ARF | emm81.3 | 995 | 8 | 4 | 4 | 4 | 3 | 13 | 4 | 42 | 21 | 1 | 4 | 4 |
| GAS02198 | New Zealand | Oceania | 2002 | ARF | emm78.3 | 1000 | 11 | 32 | 52 | 1 | 67 | 13 | 199 | 249 | 23 | 2 | 63 | 73 |
| GAS11262 | New Zealand | Oceania | 2011 | ARF | emm74.0 | 120 | 28 | 13 | 3 | 41 | 50 | 53 | 123 | 82 | 11 | 4 | 59 | 70 |
| GAS1161 | New Zealand | Oceania | 2011 | invasive, NOS | emm71.0 | 318 | 19 | 20 | 13 | 19 | 2 | 22 | 3 | 27 | 1 | 1 | 4 | 24 |
| GAS11290 | New Zealand | Oceania | 2011 | ARF | emm19.4 | 616 | 50 | 32 | 10 | 1 | 6 | 55 | 78 | 13 | 1 | 2 | 6 | 106 |
| GAS13362 | New Zealand | Oceania | 2013 | invasive, NOS | emm19.4 | 616 | 50 | 32 | 10 | 1 | 6 | 55 | 78 | 13 | 1 | 2 | 6 | 106 |
| GAS1341 | New Zealand | Oceania | 2013 | invasive, NOS | emm197.0 | 998 | 11 | 151 | 10 | 8 | 165 | 16 | 17 | 236 | 1 | 2 | 6 | 20 |
| GAS12316 | New Zealand | Oceania | 2012 | ARF | emm74.0 | 120 | 28 | 13 | 3 | 41 | 50 | 53 | 123 | 82 | 11 | 4 | 59 | 70 |
| GAS12180 | New Zealand | Oceania | 2012 | invasive, NOS | emm81.0 | 330 | 62 | 75 | 45 | 52 | 5 | 6 | 45 | 106 | 24 | 17 | 66 | 61 |
| GAS1282 | New Zealand | Oceania | 2012 | ARF | emm81.0 | 330 | 62 | 75 | 45 | 52 | 5 | 6 | 45 | 106 | 24 | 17 | 66 | 61 |
| GAS11441 | New Zealand | Oceania | 2011 | ARF | emm19.4 | 616 | 50 | 32 | 10 | 1 | 6 | 55 | 78 | 13 | 1 | 2 | 6 | 106 |
| GAS11237 | New Zealand | Oceania | 2011 | invasive, NOS | emm44.0 | 178 | 5 | 7 | 7 | 3 | 2 | 3 | 24 | 8 | 14 | 3 | 7 | 7 |
| GAS1156 | New Zealand | Oceania | 2011 | pharyngitis and/or tonsillitis | emm65.0 | 111 | 22 | 22 | 17 | 38 | 23 | 43 | 28 | 65 | 1 | 1 | 1 | 27 |

| | | | | | | | | | | | | | | | | | | |
|---|---|---|---|---|---|---|---|---|---|---|---|---|---|---|---|---|---|---|
| GAS13358 | New Zealand | Oceania | 2013 | invasive, NOS | emm197.0 | 998 | 11 | 151 | 10 | 8 | 165 | 16 | 17 | 236 | 1 | 2 | 6 | 20 |
| GAS1328 | New Zealand | Oceania | 2013 | invasive, NOS | emm74.0 | 120 | 28 | 13 | 3 | 41 | 50 | 53 | 123 | 82 | 11 | 4 | 59 | 70 |
| GAS1187 | New Zealand | Oceania | 2011 | ARF | emm71.0 | 318 | 19 | 20 | 13 | 19 | 2 | 22 | 3 | 27 | 1 | 1 | 4 | 24 |
| GAS12314 | New Zealand | Oceania | 2012 | pharyngitis and/or tonsillitis | emm53.0 | 11 | 49 | 5 | 10 | 8 | 15 | 54 | 17 | 84 | 1 | 2 | 6 | 20 |
| GAS12172 | New Zealand | Oceania | 2012 | pharyngitis and/or tonsillitis | emm3.1 | 15 | 15 | 15 | 10 | 1 | 10 | 32 | 111 | 129 | 1 | 2 | 6 | 31 |
| GAS1262 | New Zealand | Oceania | 2012 | invasive, NOS | emm104.0 | 789 | 14 | 13 | 10 | 8 | 13 | 13 | 15 | 17 | 10 | 2 | 15 | 16 |
| GAS11426 | New Zealand | Oceania | 2011 | pharyngitis and/or tonsillitis | emm53.0 | 11 | 49 | 5 | 10 | 8 | 15 | 54 | 17 | 84 | 1 | 2 | 6 | 20 |
| GAS11212 | New Zealand | Oceania | 2011 | invasive, NOS | emm74.0 | 120 | 28 | 13 | 3 | 41 | 50 | 53 | 123 | 82 | 11 | 4 | 59 | 70 |
| GAS1142 | New Zealand | Oceania | 2011 | ARF | emm65.4 | 129 | 11 | 66 | 40 | 35 | 102 | 76 | 44 | 145 | 1 | 2 | 87 | 31 |
| GAS13350 | New Zealand | Oceania | 2013 | pharyngitis and/or tonsillitis | emm19.4 | 616 | 50 | 32 | 10 | 1 | 6 | 55 | 78 | 13 | 1 | 2 | 6 | 106 |
| GAS1318 | New Zealand | Oceania | 2013 | invasive, NOS | emm232.0 | 902 | 14 | 37 | 31 | 3 | 2 | 38 | 47 | 242 | 1 | 2 | 58 | 25 |
| GAS12274 | New Zealand | Oceania | 2012 | pharyngitis and/or tonsillitis | emm218.1 | 292 | 38 | 45 | 3 | 16 | 44 | 44 | 53 | 61 | 54 | 2 | 12 | 3 |
| GAS12171 | New Zealand | Oceania | 2012 | ARF | emm3.33 | 15 | 15 | 15 | 10 | 1 | 10 | 32 | 111 | 129 | 1 | 2 | 6 | 31 |
| GAS13399 | New Zealand | Oceania | 2013 | ARF | emm100.5 | 119 | 11 | 11 | 7 | 10 | 10 | 7 | 37 | 39 | 19 | 31 | 30 | 31 |
| GAS1224 | New Zealand | Oceania | 2012 | ARF | emm238.2 | 868 | 1 | 20 | 123 | 1 | 1 | 15 | 14 | 16 | 5 | 6 | 14 | 15 |
| GAS11423 | New Zealand | Oceania | 2011 | ARF | emm53.0 | 11 | 49 | 5 | 10 | 8 | 15 | 54 | 17 | 84 | 1 | 2 | 6 | 20 |
| GAS11195 | New Zealand | Oceania | 2011 | invasive, NOS | emm59.1 | 172 | 71 | 84 | 24 | 2 | 93 | 68 | 107 | 122 | 18 | 3 | 75 | 85 |
| GAS1121 | New Zealand | Oceania | 2011 | ARF | emm41.2 | 579 | 11 | 5 | 10 | 51 | 15 | 16 | 77 | 99 | 1 | 2 | 6 | 20 |
| GAS13257 | New Zealand | Oceania | 2013 | pharyngitis and/or tonsillitis | emm95.0 | 604 | 11 | 32 | 30 | 31 | 37 | 7 | 46 | 51 | 1 | 2 | 6 | 44 |
| GAS1315 | New Zealand | Oceania | 2013 | invasive, NOS | emm76.0 | 378 | 122 | 3 | 3 | 3 | 2 | 3 | 52 | 59 | 24 | 12 | 38 | 3 |
| GAS12271 | New Zealand | Oceania | 2012 | invasive, NOS | emm1.0 | 28 | 1 | 1 | 1 | 1 | 1 | 1 | 1 | 1 | 1 | 1 | 1 | 1 |
| GAS12163 | New Zealand | Oceania | 2012 | ARF | emm108.1 | 14 | 28 | 87 | 10 | 1 | 6 | 63 | 78 | 240 | 1 | 2 | 6 | 31 |
| GAS1222 | New Zealand | Oceania | 2012 | pharyngitis and/or tonsillitis | emm71.0 | 981 | 19 | 20 | 13 | 19 | 2 | 22 | 3 | 27 | 1 | 1 | 4 | 24 |
| GAS11371 | New Zealand | Oceania | 2011 | pharyngitis and/or tonsillitis | emm11.0 | ND | 1 | 6 | 76 | 1 | 104 | 20 | 13 | 25 | 8 | 3 | 21 | 23 |
| GAS1377 | New Zealand | Oceania | 2013 | pharyngitis and/or tonsillitis | emm59.1 | 172 | 71 | 84 | 24 | 2 | 93 | 68 | 107 | 122 | 18 | 3 | 75 | 85 |
| GAS11186 | New Zealand | Oceania | 2011 | pharyngitis and/or tonsillitis | emm19.4 | 616 | 50 | 32 | 10 | 1 | 6 | 55 | 78 | 13 | 1 | 2 | 6 | 106 |
| GAS10495 | New Zealand | Oceania | 2010 | pharyngitis and/or tonsillitis | emm19.4 | 616 | 50 | 32 | 10 | 1 | 6 | 55 | 78 | 13 | 1 | 2 | 6 | 106 |
| GAS13208 | New Zealand | Oceania | 2013 | pharyngitis and/or tonsillitis | emm74.0 | 120 | 28 | 13 | 3 | 41 | 50 | 53 | 123 | 82 | 11 | 4 | 59 | 70 |
| GAS12484 | New Zealand | Oceania | 2012 | invasive, NOS | emm77.0 | 572 | 15 | 99 | 3 | 39 | 56 | 13 | 68 | 146 | 32 | 3 | 54 | 101 |
| GAS12260 | New Zealand | Oceania | 2012 | ARF | emm104.0 | 789 | 14 | 13 | 10 | 8 | 13 | 13 | 15 | 17 | 10 | 2 | 15 | 16 |
| GAS12150 | New Zealand | Oceania | 2012 | ARF | emm232.0 | 902 | 14 | 37 | 31 | 3 | 2 | 38 | 47 | 242 | 1 | 2 | 58 | 25 |
| GAS1219 | New Zealand | Oceania | 2012 | invasive, NOS | emm108.1 | 14 | 28 | 87 | 10 | 1 | 6 | 63 | 78 | 240 | 1 | 2 | 6 | 31 |
| GAS11364 | New Zealand | Oceania | 2011 | ARF | emm74.0 | 120 | 28 | 13 | 3 | 41 | 50 | 53 | 123 | 82 | 11 | 4 | 59 | 70 |
| GAS11154 | New Zealand | Oceania | 2011 | pharyngitis and/or tonsillitis | emm41.2 | 579 | 11 | 5 | 10 | 51 | 15 | 16 | 77 | 99 | 1 | 2 | 6 | 20 |
| GAS10481 | New Zealand | Oceania | 2010 | ARF | emm74.0 | 120 | 28 | 13 | 3 | 41 | 50 | 53 | 123 | 82 | 11 | 4 | 59 | 70 |
| GAS12357 | New Zealand | Oceania | 2012 | ARF | emm41.2 | 579 | 11 | 5 | 10 | 51 | 15 | 16 | 77 | 99 | 1 | 2 | 6 | 20 |
| GAS13169 | New Zealand | Oceania | 2013 | pharyngitis and/or tonsillitis | emm232.0 | 902 | 14 | 37 | 31 | 3 | 2 | 38 | 47 | 242 | 1 | 2 | 58 | 25 |
| GAS12463 | New Zealand | Oceania | 2012 | pharyngitis and/or tonsillitis | emm41.2 | 579 | 11 | 5 | 10 | 51 | 15 | 16 | 77 | 99 | 1 | 2 | 6 | 20 |
| GAS12244 | New Zealand | Oceania | 2012 | pharyngitis and/or tonsillitis | emm71.0 | 318 | 19 | 20 | 13 | 19 | 2 | 22 | 3 | 27 | 1 | 1 | 4 | 24 |
| GAS12144 | New Zealand | Oceania | 2012 | invasive, NOS | emm108.1 | 14 | 28 | 87 | 10 | 1 | 6 | 63 | 78 | 240 | 1 | 2 | 6 | 31 |
| GAS1218 | New Zealand | Oceania | 2012 | invasive, NOS | emm3.1 | 15 | 15 | 15 | 10 | 1 | 10 | 32 | 111 | 129 | 1 | 2 | 6 | 31 |
| GAS11353 | New Zealand | Oceania | 2011 | ARF | emm92.0 | 82 | 17 | 4 | 3 | 18 | 19 | 19 | 22 | 24 | 13 | 2 | 20 | 22 |
| GAS11146 | New Zealand | Oceania | 2011 | ARF | emm44.0 | 178 | 5 | 7 | 7 | 3 | 2 | 3 | 24 | 8 | 14 | 3 | 7 | 7 |
| GAS10476 | New Zealand | Oceania | 2010 | invasive, NOS | emm33.0 | 3 | 11 | 11 | 7 | 10 | 35 | 36 | 85 | 79 | 1 | 2 | 6 | 20 |
| GAS13137 | New Zealand | Oceania | 2013 | pharyngitis and/or tonsillitis | emm98.1 | 205 | 15 | 15 | 10 | 1 | 10 | 96 | 19 | 21 | 12 | 2 | 6 | 20 |
| GAS12446 | New Zealand | Oceania | 2012 | pharyngitis and/or tonsillitis | emm74.0 | 120 | 28 | 13 | 3 | 41 | 50 | 53 | 123 | 82 | 11 | 4 | 59 | 70 |
| GAS12229 | New Zealand | Oceania | 2012 | ARF | emm238.2 | 1074 | 1 | 1 | 1 | 1 | 1 | 15 | 14 | 93 | 5 | 6 | 14 | 15 |
| GAS12252 | New Zealand | Oceania | 2012 | invasive, NOS | emm1.0 | 28 | 1 | 1 | 1 | 1 | 1 | 1 | 1 | 3 | 1 | 1 | 1 | 1 |

| | | | | | | | | | | | | | | | | | | |
|---|---|---|---|---|---|---|---|---|---|---|---|---|---|---|---|---|---|---|
| GAS12140 | New Zealand | Oceania | 2012 | invasive, NOS | emm232.0 | 902 | 14 | 37 | 31 | 3 | 2 | 38 | 47 | 242 | 1 | 2 | 58 | 25 |
| GAS1213 | New Zealand | Oceania | 2012 | invasive, NOS | emm232.0 | 902 | 14 | 37 | 31 | 3 | 2 | 38 | 47 | 242 | 1 | 2 | 58 | 25 |
| GAS11340 | New Zealand | Oceania | 2011 | invasive, NOS | emm19.4 | 616 | 50 | 32 | 10 | 1 | 6 | 55 | 78 | 13 | 1 | 2 | 6 | 106 |
| GAS11141 | New Zealand | Oceania | 2011 | ARF | emm59.1 | 172 | 71 | 84 | 24 | 2 | 93 | 68 | 107 | 122 | 18 | 3 | 75 | 85 |
| GAS10470 | New Zealand | Oceania | 2010 | invasive, NOS | emm74.0 | 120 | 28 | 13 | 3 | 41 | 50 | 53 | 123 | 82 | 11 | 4 | 59 | 70 |
| GAS13132 | New Zealand | Oceania | 2013 | pharyngitis and/or tonsillitis | emm108.1 | 14 | 28 | 87 | 10 | 1 | 6 | 63 | 78 | 240 | 1 | 2 | 6 | 31 |
| GAS12411 | New Zealand | Oceania | 2012 | invasive, NOS | emm218.1 | 292 | 38 | 45 | 3 | 16 | 44 | 93 | 53 | 61 | 54 | 2 | 12 | 3 |
| GAS12243 | New Zealand | Oceania | 2012 | pharyngitis and/or tonsillitis | emm1.0 | 28 | 1 | 1 | 1 | 1 | 1 | 1 | 1 | 3 | 1 | 1 | 1 | 1 |
| GAS12139 | New Zealand | Oceania | 2012 | invasive, NOS | emm104.0 | 789 | 14 | 13 | 10 | 8 | 13 | 13 | 15 | 17 | 10 | 2 | 15 | 16 |
| GAS12106 | New Zealand | Oceania | 2012 | pharyngitis and/or tonsillitis | emm81.0 | 330 | 62 | 75 | 45 | 52 | 5 | 6 | 45 | 106 | 24 | 17 | 66 | 61 |
| GAS11551 | New Zealand | Oceania | 2011 | pharyngitis and/or tonsillitis | emm71.0 | 318 | 19 | 20 | 13 | 19 | 2 | 22 | 3 | 27 | 1 | 1 | 4 | 24 |
| GAS11334 | New Zealand | Oceania | 2011 | invasive, NOS | emm92.0 | 82 | 17 | 4 | 3 | 82 | 19 | 19 | 22 | 24 | 13 | 2 | 20 | 22 |
| GAS11129 | New Zealand | Oceania | 2011 | invasive, NOS | emm41.2 | 579 | 11 | 5 | 10 | 51 | 15 | 16 | 77 | 99 | 1 | 2 | 6 | 20 |
| GAS10463 | New Zealand | Oceania | 2010 | invasive, NOS | emm232.0 | 967 | 14 | 37 | 31 | 3 | 2 | 38 | 47 | 242 | 1 | 2 | 58 | 25 |
| GAS10458 | New Zealand | Oceania | 2010 | invasive, NOS | emm36.2 | 748 | 15 | 15 | 10 | 1 | 10 | 32 | 197 | 21 | 12 | 8 | 37 | 31 |
| GAS10243 | New Zealand | Oceania | 2010 | invasive, NOS | emm105.0 | 954 | 6 | 6 | 6 | 6 | 5 | 6 | 6 | 12 | 5 | 4 | 6 | 11 |
| GAS1084 | New Zealand | Oceania | 2010 | ARF | emm232.0 | 902 | 14 | 37 | 31 | 3 | 2 | 38 | 47 | 242 | 1 | 2 | 58 | 25 |
| GAS09422 | New Zealand | Oceania | 2009 | invasive, NOS | emm12.0 | 36 | 14 | 3 | 3 | 3 | 2 | 3 | 52 | 243 | 24 | 12 | 38 | 156 |
| GAS09204 | New Zealand | Oceania | 2009 | ARF | emm93.4 | 814 | 28 | 104 | 10 | 1 | 108 | 32 | 61 | 42 | 50 | 2 | 93 | 69 |
| GAS0830 | New Zealand | Oceania | 2008 | ARF | emm33.0 | 3 | 11 | 11 | 7 | 10 | 35 | 36 | 85 | 79 | 1 | 2 | 6 | 20 |
| GAS11456 | New Zealand | Oceania | 2011 | invasive, NOS | emm36.2 | 748 | 15 | 15 | 10 | 1 | 10 | 32 | 197 | 21 | 12 | 8 | 37 | 31 |
| GAS066 | New Zealand | Oceania | 2006 | ARF | emm65.0 | 111 | 22 | 22 | 17 | 38 | 43 | 43 | 28 | 65 | 1 | 1 | 1 | 27 |
| GAS03101 | New Zealand | Oceania | 2003 | ARF | emm101.0 | 182 | 15 | 31 | 10 | 20 | 15 | 31 | 38 | 41 | 1 | 3 | 32 | 37 |
| GAS10417 | New Zealand | Oceania | 2010 | invasive, NOS | emm232.0 | 902 | 14 | 37 | 124 | 3 | 2 | 38 | 47 | 242 | 1 | 2 | 58 | 25 |
| GAS10219 | New Zealand | Oceania | 2010 | invasive, NOS | emm232.0 | 902 | 14 | 37 | 31 | 3 | 2 | 38 | 47 | 242 | 1 | 2 | 58 | 25 |
| GAS1074 | New Zealand | Oceania | 2010 | invasive, NOS | emm71.0 | 318 | 19 | 20 | 13 | 19 | 2 | 22 | 3 | 27 | 1 | 1 | 4 | 24 |
| GAS09418 | New Zealand | Oceania | 2009 | pharyngitis and/or tonsillitis | emm105.0 | 954 | 6 | 6 | 6 | 6 | 5 | 6 | 6 | 12 | 5 | 4 | 6 | 11 |
| GAS09194 | New Zealand | Oceania | 2009 | invasive, NOS | emm65.0 | 111 | 22 | 22 | 17 | 38 | 23 | 43 | 28 | 65 | 1 | 1 | 1 | 27 |
| GAS07174 | New Zealand | Oceania | 2007 | ARF | emm232.1 | 1013 | 18 | 18 | 16 | 16 | 21 | 25 | 125 | 250 | 8 | 3 | 21 | 23 |
| GAS05207 | New Zealand | Oceania | 2005 | pharyngitis and/or tonsillitis | emm93.4 | 814 | 28 | 104 | 10 | 1 | 108 | 32 | 61 | 42 | 50 | 2 | 93 | 69 |
| GAS0391 | New Zealand | Oceania | 2003 | invasive, NOS | emm78.3 | 1000 | 11 | 32 | 52 | 1 | 67 | 13 | 199 | 249 | 23 | 2 | 63 | 73 |
| NZ131 | New Zealand | Oceania | 1991 | APSGN | emm49.1 | 1073 | 4 | 153 | 4 | 11 | 168 | 97 | 200 | 251 | 9 | 2 | 99 | 158 |
| GUR | Russia | Asia | 2017 | ND | emm111.2 | 1065 | 14 | 154 | 75 | 8 | 169 | 19 | 187 | 109 | 1 | 2 | 125 | 159 |
| GURSA1 | Russia | Asia | 2017 | ND | emm111.2 | 1065 | 14 | 154 | 75 | 8 | 169 | 19 | 187 | 109 | 1 | 2 | 125 | 159 |
| A20 | Taiwan | Asia | ND | invasive, NOS | emm1.0 | 28 | 1 | 1 | 1 | 1 | 1 | 1 | 1 | 1 | 1 | 1 | 1 | 160 |
| MGAS2096 | Trinidad and Tobago | North America | 1960 | APSGN | emm12.0 | 36 | 14 | 3 | 3 | 3 | 2 | 3 | 52 | 59 | 24 | 12 | 38 | 3 |
| PHE_12088 | UK | Europe | 2014 | scarlet fever | emm3.56 | 15 | 15 | 15 | 10 | 1 | 10 | 32 | 111 | 129 | 1 | 2 | 6 | 31 |
| PHE_12036 | UK | Europe | 2014 | invasive, NOS | emm131.0 | 785 | 1 | 1 | 1 | 1 | 1 | 1 | 1 | 1 | 1 | 1 | 1 | 1 |
| PHE_12783 | UK | Europe | 2014 | invasive, NOS | emm90.5 | 184 | 20 | 20 | 14 | 83 | 170 | 23 | 25 | 28 | 15 | 3 | 22 | 25 |
| PHE_33063 | UK | Europe | 2014 | scarlet fever | emm12.0 | 1068 | 14 | 3 | 3 | 3 | 2 | 3 | 52 | 59 | 24 | 12 | 38 | 3 |
| PHE_32280 | UK | Europe | 2014 | invasive, NOS | emm12.37 | 242 | 14 | 3 | 3 | 3 | 2 | 3 | 52 | 252 | 24 | 12 | 38 | 3 |
| PHE_24974 | UK | Europe | 2014 | invasive, NOS | emm73.0 | 331 | 14 | 96 | 3 | 41 | 50 | 13 | 3 | 137 | 48 | 2 | 49 | 60 |
| PHE_55087 | UK | Europe | 2014 | scarlet fever | emm2.0 | 55 | 32 | 26 | 28 | 24 | 29 | 13 | 43 | 48 | 15 | 3 | 12 | 42 |
| PHE_33054 | UK | Europe | 2014 | invasive, NOS | emm53.0 | 363 | 46 | 32 | 7 | 1 | 171 | 7 | 201 | 79 | 1 | 16 | 56 | 31 |
| PHE_107859 | UK | Europe | 2014 | scarlet fever | emm22.0 | 46 | 7 | 149 | 5 | 5 | 4 | 5 | 5 | 253 | 4 | 3 | 5 | 5 |
| PHE_12749 | UK | Europe | 2014 | scarlet fever | emm12.0 | 36 | 14 | 3 | 3 | 3 | 2 | 3 | 52 | 59 | 24 | 12 | 38 | 3 |
| PHE_39584 | UK | Europe | 2014 | scarlet fever | emm58.0 | 176 | 14 | 94 | 10 | 8 | 13 | 13 | 15 | 47 | 10 | 2 | 34 | 31 |
| PHE_27636 | UK | Europe | 2014 | scarlet fever | emm77.0 | 399 | 82 | 25 | 3 | 22 | 27 | 13 | 26 | 29 | 8 | 3 | 25 | 30 |

| | | | | | | | | | | | | | | | | | | |
|---|---|---|---|---|---|---|---|---|---|---|---|---|---|---|---|---|---|---|
| PHE_39586 | UK | Europe | 2014 | scarlet fever | emm28.0 | 458 | 3 | 3 | 3 | 3 | 2 | 3 | 202 | 4 | 1 | 1 | 3 | 3 |
| PHE_18921 | UK | Europe | 2014 | scarlet fever | emm131.0 | 382 | 65 | 80 | 62 | 55 | 85 | 62 | 97 | 130 | 11 | 20 | 69 | 82 |
| PHE_27701 | UK | Europe | 2014 | scarlet fever | emm3.1 | 406 | 15 | 155 | 10 | 1 | 10 | 32 | 111 | 129 | 1 | 2 | 6 | 31 |
| PHE_24945 | UK | Europe | 2014 | invasive, NOS | emm5.23 | 99 | 14 | 91 | 70 | 1 | 17 | 71 | 114 | 254 | 47 | 22 | 80 | 91 |
| PHE_12750 | UK | Europe | 2014 | scarlet fever | emm12.0 | 36 | 14 | 3 | 3 | 3 | 2 | 3 | 52 | 59 | 24 | 12 | 38 | 3 |
| PHE_12875 | UK | Europe | 2014 | scarlet fever | emm6.0 | 382 | 65 | 80 | 62 | 55 | 85 | 62 | 97 | 130 | 11 | 20 | 69 | 82 |
| PHE_25160 | UK | Europe | 2014 | invasive, NOS | emm76.0 | 50 | 14 | 3 | 3 | 3 | 2 | 3 | 52 | 59 | 24 | 12 | 38 | 3 |
| PHE_33034 | UK | Europe | 2014 | scarlet fever | emm4.0 | 1066 | 124 | 8 | 4 | 7 | 7 | 9 | 8 | 9 | 7 | 2 | 8 | 8 |
| PHE_12906 | UK | Europe | 2014 | scarlet fever | emm75.0 | 150 | 25 | 26 | 20 | 24 | 29 | 29 | 33 | 155 | 15 | 3 | 12 | 33 |
| PHE_26043 | UK | Europe | 2014 | scarlet fever | emm28.0 | 52 | 3 | 3 | 3 | 3 | 2 | 3 | 3 | 4 | 1 | 1 | 126 | 161 |
| PHE_26023 | UK | Europe | 2014 | scarlet fever | emm6.0 | 382 | 65 | 80 | 62 | 55 | 85 | 62 | 97 | 130 | 11 | 20 | 69 | 82 |
| PHE_12907 | UK | Europe | 2014 | scarlet fever | emm3.93 | 315 | 15 | 15 | 10 | 1 | 10 | 32 | 111 | 129 | 1 | 2 | 6 | 31 |
| PHE_30640 | UK | Europe | 2014 | invasive, NOS | emm219.0 | 25 | 125 | 58 | 125 | 1 | 6 | 18 | 69 | 255 | 1 | 1 | 127 | 162 |
| PHE_30600 | UK | Europe | 2014 | invasive, NOS | emm6.0 | 382 | 65 | 80 | 62 | 55 | 85 | 62 | 97 | 130 | 11 | 20 | 69 | 82 |
| PHE_33016 | UK | Europe | 2014 | invasive, NOS | emm81.0 | 624 | 5 | 53 | 43 | 42 | 172 | 47 | 64 | 119 | 30 | 14 | 30 | 61 |
| PHE_32292 | UK | Europe | 2014 | invasive, NOS | emm90.2 | 184 | 14 | 88 | 68 | 57 | 94 | 70 | 109 | 127 | 1 | 2 | 79 | 88 |
| PHE_12026 | UK | Europe | 2014 | invasive, NOS | emm3.56 | 15 | 15 | 15 | 10 | 1 | 10 | 32 | 111 | 129 | 1 | 2 | 6 | 31 |
| PHE_12861 | UK | Europe | 2014 | scarlet fever | emm3.1 | 315 | 15 | 15 | 10 | 1 | 10 | 32 | 111 | 129 | 1 | 2 | 6 | 31 |
| PHE_18945 | UK | Europe | 2014 | invasive, NOS | emm81.0 | 624 | 5 | 53 | 43 | 42 | 52 | 47 | 64 | 119 | 30 | 14 | 30 | 61 |
| PHE_25152 | UK | Europe | 2014 | scarlet fever | emm219.0 | 25 | 14 | 58 | 125 | 1 | 6 | 18 | 203 | 255 | 1 | 1 | 127 | 162 |
| PHE_45816 | UK | Europe | 2014 | scarlet fever | emm28.0 | 52 | 3 | 3 | 3 | 3 | 2 | 3 | 3 | 4 | 1 | 1 | 3 | 3 |
| PHE_12832 | UK | Europe | 2014 | scarlet fever | emm4.0 | 39 | 4 | 8 | 4 | 7 | 7 | 9 | 8 | 9 | 7 | 2 | 8 | 8 |
| PHE_12841 | UK | Europe | 2014 | scarlet fever | emm87.0 | 62 | 67 | 15 | 10 | 1 | 72 | 58 | 87 | 95 | 1 | 2 | 52 | 74 |
| PHE_12764 | UK | Europe | 2014 | scarlet fever | emm28.0 | 52 | 3 | 3 | 3 | 84 | 2 | 3 | 204 | 4 | 1 | 1 | 3 | 163 |
| PHE_12067 | UK | Europe | 2014 | invasive, NOS | emm81.0 | 624 | 5 | 53 | 43 | 42 | 52 | 47 | 64 | 119 | 30 | 14 | 30 | 61 |
| PHE_45811 | UK | Europe | 2014 | scarlet fever | emm75.0 | 150 | 25 | 26 | 20 | 24 | 29 | 29 | 33 | 155 | 15 | 3 | 12 | 33 |
| PHE_12021 | UK | Europe | 2014 | scarlet fever | emm12.60 | 36 | 14 | 3 | 3 | 3 | 2 | 3 | 52 | 59 | 24 | 12 | 38 | 3 |
| PHE_18939 | UK | Europe | 2014 | invasive, NOS | emm94.0 | 89 | 8 | 4 | 4 | 4 | 3 | 13 | 4 | 42 | 21 | 1 | 4 | 4 |
| PHE_25158 | UK | Europe | 2014 | invasive, NOS | emm82.0 | 314 | 14 | 58 | 46 | 85 | 6 | 18 | 69 | 77 | 1 | 1 | 1 | 3 |
| PHE_12046 | UK | Europe | 2014 | scarlet fever | emm82.0 | 36 | 14 | 3 | 3 | 3 | 2 | 3 | 52 | 59 | 24 | 12 | 38 | 3 |
| PHE_30630 | UK | Europe | 2014 | invasive, NOS | emm1.25 | 28 | 1 | 1 | 1 | 1 | 1 | 1 | 1 | 134 | 1 | 1 | 1 | 1 |
| PHE_24940 | UK | Europe | 2014 | invasive, NOS | emm90.2 | 184 | 14 | 88 | 68 | 57 | 94 | 70 | 109 | 127 | 1 | 2 | 79 | 88 |
| PHE_12912 | UK | Europe | 2014 | scarlet fever | emm3.93 | 315 | 15 | 15 | 10 | 1 | 10 | 32 | 111 | 129 | 1 | 2 | 6 | 31 |
| PHE_12855 | UK | Europe | 2014 | scarlet fever | emm89.0 | 101 | 2 | 2 | 2 | 2 | 2 | 2 | 2 | 2 | 2 | 2 | 2 | 2 |
| PHE_45825 | UK | Europe | 2014 | scarlet fever | emm12.37 | 242 | 14 | 3 | 3 | 3 | 2 | 3 | 52 | 252 | 24 | 12 | 38 | 3 |
| PHE_104230 | UK | Europe | 2014 | invasive, NOS | emm44.0 | 1086 | 125 | 58 | 125 | 1 | 6 | 18 | 69 | 255 | 1 | 1 | 127 | 162 |
| PHE_32243 | UK | Europe | 2014 | invasive, NOS | emm94.0 | 89 | 8 | 4 | 4 | 4 | 3 | 13 | 4 | 42 | 21 | 1 | 4 | 4 |
| PHE_30599 | UK | Europe | 2014 | invasive, NOS | emm58.8 | 985 | 20 | 20 | 14 | 19 | 2 | 23 | 25 | 57 | 18 | 3 | 22 | 25 |
| PHE_12810 | UK | Europe | 2014 | scarlet fever | emm6.0 | 382 | 65 | 80 | 62 | 55 | 85 | 62 | 205 | 130 | 11 | 20 | 69 | 82 |
| PHE_26033 | UK | Europe | 2014 | scarlet fever | emm4.0 | 38 | 4 | 8 | 4 | 11 | 51 | 9 | 8 | 9 | 7 | 2 | 8 | 8 |
| PHE_45819 | UK | Europe | 2014 | scarlet fever | emm4.0 | 39 | 4 | 8 | 4 | 7 | 51 | 9 | 8 | 9 | 7 | 2 | 8 | 8 |
| PHE_39598 | UK | Europe | 2014 | scarlet fever | emm6.0 | 382 | 65 | 80 | 126 | 55 | 85 | 62 | 97 | 130 | 11 | 20 | 69 | 82 |
| PHE_30647 | UK | Europe | 2014 | scarlet fever | emm4.0 | 38 | 4 | 8 | 4 | 11 | 173 | 9 | 8 | 9 | 7 | 2 | 8 | 8 |
| PHE_45826 | UK | Europe | 2014 | scarlet fever | emm3.103 | 406 | 15 | 15 | 10 | 1 | 10 | 32 | 111 | 129 | 1 | 2 | 6 | 31 |
| PHE_26068 | UK | Europe | 2014 | invasive, NOS | emm22.0 | 46 | 7 | 5 | 5 | 5 | 4 | 5 | 5 | 6 | 4 | 3 | 5 | 5 |
| PHE_12780 | UK | Europe | 2014 | invasive, NOS | emm90.2 | 184 | 11 | 32 | 52 | 1 | 67 | 13 | 118 | 256 | 1 | 2 | 83 | 93 |
| PHE_30624 | UK | Europe | 2014 | invasive, NOS | emm58.8 | 985 | 20 | 20 | 14 | 19 | 2 | 23 | 25 | 57 | 18 | 3 | 22 | 25 |
| PHE_104235 | UK | Europe | 2014 | invasive, NOS | emm73.0 | 331 | 14 | 96 | 3 | 41 | 50 | 13 | 3 | 137 | 48 | 2 | 49 | 60 |

| | | | | | | | | | | | | | | | | | | |
|---|---|---|---|---|---|---|---|---|---|---|---|---|---|---|---|---|---|---|
| PHE_26041 | UK | Europe | 2014 | invasive, NOS | emm3.5 | 15 | 15 | 15 | 10 | 1 | 10 | 32 | 111 | 129 | 1 | 2 | 6 | 31 |
| PHE_12853 | UK | Europe | 2014 | scarlet fever | emm4.0 | 39 | 4 | 8 | 4 | 7 | 7 | 9 | 8 | 9 | 7 | 2 | 8 | 8 |
| PHE_45820 | UK | Europe | 2014 | scarlet fever | emm87.0 | 62 | 67 | 15 | 10 | 1 | 72 | 58 | 87 | 95 | 1 | 2 | 52 | 74 |
| PHE_12908 | UK | Europe | 2014 | scarlet fever | emm3.1 | 315 | 15 | 15 | 10 | 1 | 10 | 32 | 111 | 129 | 1 | 2 | 6 | 31 |
| PHE_18896 | UK | Europe | 2014 | scarlet fever | emm131.0 | 382 | 65 | 80 | 62 | 55 | 85 | 62 | 97 | 130 | 11 | 20 | 69 | 82 |
| PHE_12833 | UK | Europe | 2014 | invasive, NOS | emm3.56 | 15 | 15 | 15 | 10 | 1 | 10 | 32 | 111 | 129 | 1 | 2 | 6 | 31 |
| PHE_12076 | UK | Europe | 2014 | invasive, NOS | emm94.0 | 89 | 8 | 4 | 4 | 4 | 3 | 13 | 4 | 42 | 21 | 1 | 4 | 4 |
| PHE_18919 | UK | Europe | 2014 | invasive, NOS | emm103.0 | 327 | 14 | 156 | 3 | 1 | 66 | 13 | 82 | 87 | 1 | 2 | 49 | 4 |
| PHE_55078 | UK | Europe | 2014 | scarlet fever | emm75.0 | 150 | 25 | 26 | 20 | 24 | 29 | 29 | 33 | 155 | 15 | 3 | 12 | 33 |
| PHE_104231 | UK | Europe | 2014 | invasive, NOS | emm18.0 | 42 | 75 | 92 | 61 | 5 | 98 | 72 | 115 | 132 | 47 | 2 | 69 | 5 |
| PHE_12894 | UK | Europe | 2014 | scarlet fever | emm3.1 | 315 | 15 | 15 | 10 | 1 | 10 | 32 | 111 | 129 | 1 | 2 | 6 | 31 |
| PHE_33069 | UK | Europe | 2014 | scarlet fever | emm12.0 | 1068 | 14 | 3 | 3 | 3 | 2 | 3 | 52 | 59 | 24 | 12 | 38 | 3 |
| PHE_33033 | UK | Europe | 2014 | invasive, NOS | emm1.0 | 785 | 1 | 1 | 1 | 1 | 1 | 1 | 1 | 1 | 1 | 1 | 1 | 1 |
| PHE_12792 | UK | Europe | 2014 | scarlet fever | emm3.1 | 15 | 15 | 15 | 10 | 1 | 10 | 32 | 111 | 129 | 1 | 2 | 6 | 31 |
| PHE_104228 | UK | Europe | 2014 | scarlet fever | emm2.0 | 55 | 32 | 26 | 28 | 24 | 29 | 13 | 43 | 257 | 15 | 3 | 12 | 42 |
| PHE_12903 | UK | Europe | 2014 | scarlet fever | emm131.0 | 382 | 65 | 80 | 62 | 55 | 85 | 62 | 97 | 130 | 11 | 20 | 69 | 82 |
| PHE_25109 | UK | Europe | 2014 | scarlet fever | emm12.0 | 36 | 14 | 3 | 3 | 3 | 2 | 3 | 52 | 59 | 24 | 12 | 38 | 3 |
| PHE_12757 | UK | Europe | 2014 | invasive, NOS | emm81.0 | 624 | 5 | 53 | 43 | 42 | 52 | 47 | 64 | 119 | 30 | 14 | 30 | 61 |
| PHE_24950 | UK | Europe | 2014 | invasive, NOS | emm76.0 | 50 | 14 | 3 | 3 | 3 | 2 | 3 | 52 | 59 | 24 | 12 | 38 | 3 |
| PHE_39585 | UK | Europe | 2014 | scarlet fever | emm3.1 | 406 | 15 | 155 | 10 | 1 | 10 | 32 | 111 | 129 | 1 | 2 | 6 | 31 |
| PHE_12854 | UK | Europe | 2014 | scarlet fever | emm89.0 | 101 | 2 | 2 | 2 | 2 | 2 | 2 | 2 | 2 | 55 | 2 | 2 | 2 |
| PHE_12066 | UK | Europe | 2014 | invasive, NOS | emm58.0 | 176 | 14 | 94 | 10 | 8 | 13 | 13 | 15 | 47 | 10 | 2 | 34 | 31 |
| PHE_39599 | UK | Europe | 2014 | scarlet fever | emm6.0 | 382 | 65 | 80 | 62 | 55 | 85 | 62 | 97 | 130 | 11 | 20 | 69 | 82 |
| PHE_12062 | UK | Europe | 2014 | scarlet fever | emm3.1 | 406 | 15 | 155 | 10 | 1 | 10 | 32 | 111 | 129 | 1 | 2 | 6 | 31 |
| PHE_104218 | UK | Europe | 2014 | scarlet fever | emm89.0 | 101 | 2 | 2 | 2 | 2 | 2 | 2 | 2 | 2 | 2 | 2 | 2 | 2 |
| PHE_45817 | UK | Europe | 2014 | scarlet fever | emm3.93 | 315 | 15 | 15 | 10 | 1 | 10 | 32 | 111 | 129 | 1 | 2 | 6 | 31 |
| PHE_18915 | UK | Europe | 2014 | scarlet fever | emm2.0 | 55 | 32 | 26 | 28 | 24 | 29 | 13 | 43 | 48 | 15 | 3 | 12 | 42 |
| PHE_32271 | UK | Europe | 2014 | invasive, NOS | emm22.0 | 46 | 7 | 149 | 5 | 5 | 4 | 5 | 5 | 6 | 4 | 3 | 5 | 5 |
| PHE_12031 | UK | Europe | 2014 | invasive, NOS | emm5.3 | 99 | 14 | 91 | 70 | 1 | 17 | 71 | 114 | 131 | 47 | 22 | 80 | 91 |
| PHE_18938 | UK | Europe | 2014 | invasive, NOS | emm171.1 | 1062 | 88 | 157 | 3 | 16 | 44 | 85 | 151 | 177 | 54 | 2 | 128 | 164 |
| PHE_30646 | UK | Europe | 2014 | scarlet fever | emm28.0 | 1067 | 3 | 3 | 3 | 3 | 2 | 3 | 3 | 4 | 1 | 1 | 3 | 3 |
| PHE_39608 | UK | Europe | 2014 | scarlet fever | emm9.0 | 75 | 34 | 77 | 60 | 32 | 2 | 1 | 95 | 113 | 5 | 3 | 11 | 45 |
| PHE_18958 | UK | Europe | 2014 | scarlet fever | emm3.23 | 15 | 15 | 15 | 10 | 1 | 10 | 32 | 111 | 129 | 1 | 2 | 6 | 31 |
| PHE_26066 | UK | Europe | 2014 | invasive, NOS | emm4.0 | 38 | 4 | 8 | 4 | 11 | 51 | 9 | 8 | 9 | 61 | 2 | 8 | 8 |
| PHE_12888 | UK | Europe | 2014 | scarlet fever | emm3.5 | 15 | 127 | 15 | 10 | 1 | 10 | 32 | 111 | 129 | 1 | 2 | 6 | 31 |
| PHE_12890 | UK | Europe | 2014 | scarlet fever | emm3.1 | 15 | 15 | 15 | 10 | 1 | 10 | 32 | 111 | 129 | 1 | 2 | 6 | 31 |
| PHE_27648 | UK | Europe | 2014 | invasive, NOS | emm9.0 | 75 | 34 | 77 | 60 | 32 | 2 | 1 | 95 | 113 | 5 | 3 | 11 | 45 |
| PHE_26062 | UK | Europe | 2014 | invasive, NOS | emm58.0 | 176 | 14 | 94 | 10 | 8 | 13 | 13 | 15 | 47 | 10 | 2 | 34 | 31 |
| PHE_33053 | UK | Europe | 2014 | invasive, NOS | emm9.0 | 75 | 34 | 77 | 60 | 32 | 2 | 1 | 95 | 113 | 5 | 3 | 11 | 45 |
| PHE_37685 | UK | Europe | 2014 | invasive, NOS | emm82.0 | 36 | 14 | 3 | 3 | 3 | 2 | 3 | 52 | 59 | 24 | 12 | 38 | 3 |
| PHE_12826 | UK | Europe | 2014 | scarlet fever | emm3.1 | 15 | 15 | 15 | 10 | 1 | 10 | 32 | 111 | 129 | 1 | 2 | 6 | 31 |
| PHE_12806 | UK | Europe | 2014 | scarlet fever | emm1.0 | 28 | 1 | 1 | 1 | 1 | 1 | 1 | 1 | 3 | 78 | 1 | 1 | 1 |
| PHE_32261 | UK | Europe | 2014 | invasive, NOS | emm3.122 | 406 | 15 | 15 | 10 | 1 | 10 | 32 | 111 | 129 | 1 | 2 | 6 | 31 |
| PHE_55095 | UK | Europe | 2014 | scarlet fever | emm6.0 | 382 | 65 | 80 | 62 | 55 | 85 | 62 | 97 | 130 | 11 | 20 | 69 | 82 |
| PHE_30587 | UK | Europe | 2014 | scarlet fever | emm3.111 | 315 | 15 | 15 | 10 | 1 | 10 | 32 | 111 | 129 | 1 | 2 | 6 | 31 |
| PHE_33064 | UK | Europe | 2014 | scarlet fever | emm12.0 | 1068 | 14 | 3 | 3 | 3 | 2 | 3 | 52 | 59 | 24 | 12 | 38 | 3 |
| PHE_12805 | UK | Europe | 2014 | scarlet fever | emm3.106 | 315 | 15 | 15 | 10 | 1 | 10 | 32 | 111 | 129 | 1 | 2 | 6 | 31 |
| PHE_12804 | UK | Europe | 2014 | invasive, NOS | emm77.0 | 63 | 23 | 24 | 57 | 1 | 26 | 28 | 30 | 33 | 8 | 3 | 24 | 29 |

| | | | | | | | | | | | | | | | | | | |
|---|---|---|---|---|---|---|---|---|---|---|---|---|---|---|---|---|---|---|
| PHE_27706 | UK | Europe | 2014 | invasive, NOS | emm77.0 | 63 | 23 | 24 | 57 | 1 | 26 | 28 | 30 | 33 | 8 | 3 | 24 | 29 |
| PHE_12864 | UK | Europe | 2014 | scarlet fever | emm87.0 | 62 | 67 | 15 | 10 | 1 | 72 | 58 | 87 | 95 | 1 | 2 | 52 | 74 |
| PHE_25153 | UK | Europe | 2014 | scarlet fever | emm28.0 | 1079 | 3 | 3 | 3 | 3 | 2 | 3 | 3 | 4 | 1 | 1 | 3 | 3 |
| GASEMM1195 | UK | Europe | 2015 | ND | emm11.0 | ND | 128 | 19 | 78 | 23 | 117 | 83 | 206 | 216 | 8 | 3 | 18 | 4 |
| GASEMM1871 | UK | Europe | 2015 | ND | emm100.2 | 773 | 93 | 25 | 3 | 26 | 71 | 45 | 162 | 190 | 60 | 26 | 114 | 135 |
| GASEMM2706 | UK | Europe | 2014 | ND | emm92.0 | 674 | 114 | 158 | 34 | 27 | 39 | 13 | 22 | 152 | 13 | 2 | 20 | 22 |
| GASEMM2087 | UK | Europe | 2015 | ND | emm18.38 | 42 | 75 | 92 | 61 | 5 | 98 | 72 | 115 | 132 | 47 | 2 | 69 | 165 |
| GASEMM2594 | UK | Europe | 2015 | ND | emm66.0 | 44 | 14 | 82 | 64 | 31 | 6 | 18 | 127 | 77 | 1 | 1 | 73 | 3 |
| GASEMM0115 | UK | Europe | 2015 | ND | emm168.1 | 819 | 14 | 25 | 3 | 22 | 27 | 13 | 26 | 258 | 8 | 3 | 83 | 28 |
| GASEMM1220 | UK | Europe | 2015 | ND | ND | 984 | 28 | 29 | 7 | 1 | 32 | 30 | 36 | 38 | 1 | 32 | 129 | 50 |
| GASEMM2296 | UK | Europe | 2015 | ND | emm44.0 | 367 | 14 | 58 | 46 | 1 | 6 | 18 | 69 | 77 | 1 | 1 | 1 | 3 |
| GASEMM2093 | UK | Europe | 2015 | ND | emm12.0 | 242 | 14 | 3 | 3 | 3 | 2 | 3 | 52 | 252 | 24 | 12 | 38 | 3 |
| GASEMM1829 | UK | Europe | 2015 | ND | emm9.0 | 204 | 34 | 39 | 33 | 32 | 174 | 1 | 49 | 54 | 5 | 3 | 11 | 45 |
| GASEMM1426 | UK | Europe | 2015 | ND | emm74.0 | 120 | 28 | 13 | 3 | 41 | 50 | 53 | 123 | 82 | 11 | 4 | 59 | 70 |
| GASEMM2806 | UK | Europe | 2015 | ND | emm19.4 | 616 | 50 | 32 | 10 | 1 | 6 | 55 | 78 | 13 | 1 | 2 | 6 | 106 |
| GASEMM2368 | UK | Europe | 2015 | ND | emm5.120 | 99 | 14 | 91 | 70 | 1 | 17 | 71 | 114 | 131 | 47 | 22 | 80 | 91 |
| GASEMM0105 | UK | Europe | 2015 | ND | emm168.1 | 819 | 14 | 25 | 3 | 22 | 27 | 13 | 26 | 258 | 8 | 3 | 83 | 28 |
| GASEMM0070 | UK | Europe | 2015 | ND | emm49.0 | 371 | 4 | 8 | 4 | 11 | 14 | 14 | 16 | 18 | 9 | 2 | 12 | 13 |
| GASEMM2885 | UK | Europe | 2015 | ND | emm113.0 | 677 | 14 | 3 | 3 | 3 | 2 | 3 | 207 | 59 | 24 | 12 | 38 | 3 |
| GASEMM0015 | UK | Europe | 2015 | ND | emm75.0 | 150 | 25 | 26 | 20 | 24 | 29 | 29 | 33 | 155 | 79 | 3 | 12 | 33 |
| GASEMM0145 | UK | Europe | 2015 | ND | emm88.4 | 919 | 14 | 89 | 69 | 1 | 20 | 2 | 67 | 74 | 82 | 2 | 6 | 31 |
| GASEMM2743 | UK | Europe | 2015 | ND | emm92.0 | 674 | 114 | 158 | 34 | 27 | 39 | 13 | 22 | 152 | 13 | 2 | 20 | 22 |
| GASEMM0052 | UK | Europe | 2015 | ND | emm75.0 | 788 | 25 | 26 | 20 | 24 | 29 | 29 | 33 | 155 | 15 | 3 | 12 | 33 |
| GASEMM0248 | UK | Europe | 2015 | ND | emm28.0 | 52 | 3 | 3 | 3 | 84 | 2 | 3 | 3 | 4 | 1 | 1 | 3 | 3 |
| GASEMM2187 | UK | Europe | 2015 | ND | emm76.0 | 875 | 14 | 13 | 10 | 8 | 13 | 13 | 15 | 17 | 10 | 2 | 15 | 16 |
| GASEMM1345 | UK | Europe | 2015 | ND | emm89.0 | 966 | 2 | 2 | 2 | 2 | 2 | 2 | 2 | 2 | 2 | 2 | 2 | 2 |
| GASEMM2863 | UK | Europe | 2015 | ND | emm5.23 | 99 | 14 | 91 | 70 | 1 | 17 | 71 | 114 | 131 | 47 | 22 | 80 | 91 |
| GASEMM2626 | UK | Europe | 2015 | ND | emm76.0 | 875 | 14 | 13 | 10 | 8 | 13 | 13 | 15 | 17 | 10 | 2 | 15 | 16 |
| GASEMM2825 | UK | Europe | 2015 | ND | emm6.4 | 37 | 65 | 80 | 62 | 55 | 85 | 62 | 97 | 115 | 11 | 20 | 69 | 82 |
| GASEMM0087 | UK | Europe | 2015 | ND | emm81.0 | 117 | 40 | 53 | 43 | 42 | 52 | 47 | 64 | 70 | 30 | 14 | 30 | 61 |
| GASEMM1403 | UK | Europe | 2015 | ND | emm3.108 | 406 | 15 | 155 | 10 | 1 | 10 | 32 | 111 | 129 | 1 | 2 | 6 | 31 |
| GASEMM1445 | UK | Europe | 2015 | ND | emm12.19 | 36 | 14 | 3 | 3 | 3 | 2 | 3 | 52 | 59 | 24 | 12 | 38 | 3 |
| GASEMM1730 | UK | Europe | 2015 | ND | emm99.3 | 229 | 14 | 41 | 120 | 1 | 20 | 13 | 208 | 56 | 5 | 4 | 20 | 98 |
| GASEMM0136 | UK | Europe | 2015 | ND | emm1.53 | 28 | 1 | 1 | 1 | 1 | 1 | 1 | 1 | 259 | 1 | 1 | 1 | 1 |
| GASEMM1699 | UK | Europe | 2015 | ND | emm11.0 | ND | 1 | 6 | 40 | 61 | 20 | 20 | 13 | 139 | 8 | 3 | 18 | 4 |
| GASEMM2100 | UK | Europe | 2015 | ND | emm12.0 | 242 | 14 | 3 | 3 | 3 | 2 | 3 | 52 | 252 | 24 | 12 | 38 | 3 |
| GASEMM1826 | UK | Europe | 2015 | ND | emm76.0 | 378 | 14 | 3 | 3 | 3 | 2 | 3 | 52 | 59 | 24 | 12 | 38 | 3 |
| GASEMM2311 | UK | Europe | 2015 | ND | emm12.84 | 36 | 14 | 3 | 3 | 3 | 2 | 3 | 52 | 59 | 24 | 12 | 38 | 3 |
| GASEMM1994 | UK | Europe | 2015 | ND | emm32.4 | 386 | 129 | 159 | 7 | 1 | 89 | 64 | 209 | 260 | 1 | 1 | 70 | 3 |
| GASEMM1462 | UK | Europe | 2015 | ND | emm57.0 | 348 | 29 | 32 | 7 | 1 | 34 | 32 | 39 | 42 | 1 | 2 | 23 | 3 |
| GASEMM1752 | UK | Europe | 2015 | ND | emm18.21 | 402 | 11 | 13 | 3 | 41 | 50 | 13 | 112 | 42 | 31 | 1 | 51 | 4 |
| GASEMM2438 | UK | Europe | 2015 | ND | emm3.7 | 15 | 15 | 15 | 10 | 1 | 10 | 32 | 111 | 129 | 1 | 2 | 6 | 31 |
| GASEMM1318 | UK | Europe | 2015 | ND | emm11.24 | ND | 1 | 97 | 40 | 61 | 20 | 20 | 120 | 139 | 8 | 3 | 85 | 95 |
| GASEMM1471 | UK | Europe | 2015 | ND | emm3.88 | 15 | 127 | 15 | 10 | 1 | 10 | 32 | 111 | 129 | 1 | 2 | 6 | 31 |
| GASEMM1536 | UK | Europe | 2015 | ND | emm232.1 | 1013 | 18 | 18 | 16 | 16 | 21 | 25 | 125 | 26 | 8 | 3 | 21 | 23 |
| GASEMM0963 | UK | Europe | 2015 | ND | emm3.86 | 406 | 15 | 15 | 10 | 1 | 10 | 32 | 111 | 129 | 1 | 2 | 6 | 31 |
| GASEMM0954 | UK | Europe | 2015 | ND | emm3.94 | 15 | 15 | 15 | 10 | 1 | 10 | 32 | 111 | 129 | 1 | 2 | 6 | 31 |
| GASEMM0568 | UK | Europe | 2015 | ND | emm12.37 | 242 | 14 | 3 | 3 | 3 | 2 | 3 | 52 | 252 | 24 | 12 | 38 | 3 |

| | | | | | | | | | | | | | | | | | | |
|---|---|---|---|---|---|---|---|---|---|---|---|---|---|---|---|---|---|---|
| GASEMM2733 | UK | Europe | 2015 | ND | emm5.119 | 99 | 14 | 91 | 70 | 1 | 17 | 71 | 114 | 131 | 47 | 22 | 80 | 91 |
| GASEMM2842 | UK | Europe | 2015 | ND | emm76.0 | 378 | 14 | 3 | 3 | 3 | 2 | 3 | 52 | 59 | 24 | 12 | 38 | 3 |
| GASEMM1778 | UK | Europe | 2015 | ND | emm131.0 | 382 | 65 | 80 | 62 | 55 | 85 | 62 | 97 | 261 | 11 | 20 | 69 | 82 |
| GASEMM1780 | UK | Europe | 2015 | ND | emm131.0 | 411 | 65 | 80 | 62 | 55 | 85 | 62 | 97 | 130 | 11 | 20 | 69 | 82 |
| GASEMM0053 | UK | Europe | 2015 | ND | emm75.0 | 788 | 25 | 26 | 20 | 24 | 29 | 29 | 33 | 155 | 15 | 3 | 12 | 33 |
| GASEMM0090 | UK | Europe | 2015 | ND | emm81.0 | 117 | 40 | 53 | 43 | 42 | 52 | 47 | 64 | 70 | 30 | 14 | 30 | 61 |
| GASEMM2575 | UK | Europe | 2015 | ND | emm218.1 | 224 | 38 | 45 | 3 | 86 | 44 | 44 | 53 | 61 | 54 | 2 | 12 | 3 |
| GASEMM1282 | UK | Europe | 2015 | ND | emm28.0 | 52 | 3 | 3 | 3 | 84 | 2 | 3 | 3 | 4 | 1 | 1 | 3 | 3 |
| GASEMM2682 | UK | Europe | 2015 | ND | emm3.41 | 15 | 15 | 15 | 10 | 1 | 10 | 32 | 111 | 129 | 1 | 2 | 6 | 31 |
| GASEMM0832 | UK | Europe | 2015 | ND | emm1.0 | 785 | 1 | 1 | 1 | 1 | 1 | 1 | 1 | 1 | 1 | 1 | 1 | 1 |
| GASEMM1031 | UK | Europe | 2015 | ND | emm218.1 | 292 | 38 | 45 | 3 | 16 | 44 | 44 | 53 | 61 | 8 | 3 | 18 | 4 |
| GASEMM1603 | UK | Europe | 2015 | ND | emm217.0 | 594 | 42 | 55 | 3 | 6 | 5 | 48 | 211 | 73 | 1 | 2 | 52 | 11 |
| GASEMM2962 | UK | Europe | 2015 | ND | emm49.4 | 228 | 14 | 25 | 79 | 22 | 27 | 13 | 26 | 29 | 8 | 3 | 25 | 114 |
| GASEMM2653 | UK | Europe | 2015 | ND | emm126.1 | 101 | 2 | 2 | 2 | 2 | 2 | 2 | 2 | 2 | 2 | 2 | 2 | 116 |
| GASEMM1338 | UK | Europe | 2015 | ND | emm123.0 | 123 | 11 | 13 | 3 | 41 | 50 | 99 | 4 | 42 | 3 | 1 | 51 | 4 |
| GASEMM0065 | UK | Europe | 2015 | ND | emm49.0 | 371 | 4 | 8 | 4 | 11 | 14 | 14 | 16 | 18 | 9 | 2 | 12 | 13 |
| GASEMM2308 | UK | Europe | 2015 | ND | emm169.3 | 53 | 70 | 4 | 3 | 87 | 19 | 19 | 101 | 118 | 42 | 2 | 72 | 3 |
| GASEMM0141 | UK | Europe | 2015 | ND | emm88.4 | 919 | 14 | 89 | 69 | 1 | 20 | 2 | 67 | 74 | 82 | 2 | 6 | 31 |
| GASEMM2656 | UK | Europe | 2015 | ND | emm80.0 | 701 | 16 | 16 | 3 | 16 | 116 | 7 | 20 | 22 | 1 | 2 | 18 | 110 |
| GASEMM1755 | UK | Europe | 2015 | ND | emm102.2 | 60 | 130 | 18 | 4 | 4 | 167 | 7 | 25 | 262 | 1 | 3 | 130 | 3 |
| GASEMM2260 | UK | Europe | 2015 | ND | emm12.84 | 36 | 14 | 3 | 3 | 3 | 2 | 3 | 52 | 59 | 24 | 12 | 38 | 3 |
| GASEMM2454 | UK | Europe | 2015 | ND | emm5.121 | 99 | 14 | 91 | 70 | 1 | 17 | 71 | 114 | 263 | 47 | 22 | 80 | 91 |
| GASEMM1319 | UK | Europe | 2015 | ND | emm11.12 | ND | 1 | 97 | 40 | 61 | 20 | 20 | 120 | 139 | 8 | 3 | 85 | 95 |
| GASEMM0274 | UK | Europe | 2015 | ND | emm12.37 | 242 | 14 | 3 | 3 | 3 | 2 | 3 | 52 | 252 | 24 | 12 | 38 | 3 |
| GASEMM1880 | UK | Europe | 2015 | ND | emm169.3 | 53 | 70 | 4 | 3 | 87 | 19 | 19 | 101 | 118 | 42 | 2 | 72 | 3 |
| GASEMM1189 | UK | Europe | 2015 | ND | emm3.116 | 15 | 15 | 15 | 10 | 1 | 10 | 32 | 111 | 129 | 1 | 2 | 6 | 31 |
| GASEMM2898 | UK | Europe | 2015 | ND | emm12.85 | 36 | 14 | 3 | 3 | 3 | 2 | 3 | 52 | 59 | 24 | 12 | 38 | 3 |
| GASEMM2106 | UK | Europe | 2015 | ND | emm3.19 | 15 | 15 | 15 | 10 | 1 | 10 | 32 | 111 | 129 | 1 | 2 | 6 | 31 |
| GASEMM2778 | UK | Europe | 2015 | ND | emm11.23 | ND | 1 | 97 | 40 | 61 | 20 | 20 | 120 | 139 | 8 | 3 | 85 | 95 |
| GASEMM2115 | UK | Europe | 2015 | ND | emm12.40 | 36 | 14 | 3 | 3 | 3 | 2 | 3 | 212 | 59 | 24 | 12 | 38 | 3 |
| GASEMM1161 | UK | Europe | 2015 | ND | emm78.3 | 253 | 2 | 160 | 7 | 88 | 176 | 43 | 213 | 264 | 2 | 34 | 2 | 166 |
| GASEMM2204 | UK | Europe | 2015 | ND | emm95.0 | 240 | 85 | 6 | 6 | 6 | 5 | 6 | 214 | 172 | 5 | 4 | 6 | 11 |
| GASEMM2512 | UK | Europe | 2015 | ND | emm5.16 | 99 | 14 | 91 | 70 | 1 | 17 | 71 | 114 | 254 | 47 | 22 | 80 | 91 |
| GASEMM2268 | UK | Europe | 2015 | ND | emm66.0 | 44 | 14 | 82 | 64 | 31 | 6 | 18 | 127 | 77 | 1 | 1 | 73 | 3 |
| GASEMM2718 | UK | Europe | 2015 | ND | emm6.49 | 382 | 65 | 80 | 62 | 55 | 85 | 62 | 97 | 130 | 11 | 20 | 69 | 82 |
| GASEMM1801 | UK | Europe | 2015 | ND | emm23.0 | 160 | 120 | 150 | 121 | 54 | 106 | 62 | 192 | 234 | 11 | 4 | 35 | 155 |
| GASEMM0851 | UK | Europe | 2015 | ND | emm6.0 | 839 | 65 | 80 | 62 | 55 | 85 | 62 | 97 | 130 | 11 | 20 | 69 | 82 |
| GASEMM0022 | UK | Europe | 2015 | ND | emm75.0 | 150 | 25 | 26 | 20 | 24 | 29 | 29 | 33 | 155 | 79 | 3 | 12 | 33 |
| GASEMM2280 | UK | Europe | 2015 | ND | emm109.1 | 718 | 94 | 126 | 3 | 27 | 177 | 37 | 45 | 191 | 15 | 3 | 35 | 34 |
| GASEMM1932 | UK | Europe | 2015 | ND | emm5.113 | 99 | 14 | 91 | 70 | 1 | 17 | 71 | 114 | 263 | 47 | 22 | 80 | 91 |
| GASEMM0669 | UK | Europe | 2015 | ND | emm32.2 | 386 | 129 | 159 | 7 | 1 | 89 | 64 | 209 | 260 | 1 | 1 | 70 | 3 |
| GASEMM0619 | UK | Europe | 2015 | ND | emm12.1 | 36 | 14 | 3 | 3 | 3 | 2 | 3 | 52 | 59 | 24 | 12 | 38 | 3 |
| GASEMM1383 | UK | Europe | 2015 | ND | emm5.86 | 99 | 14 | 91 | 70 | 1 | 17 | 71 | 114 | 131 | 47 | 22 | 80 | 91 |
| GASEMM3004 | UK | Europe | 2015 | ND | emm89.0 | 791 | 2 | 2 | 2 | 2 | 2 | 2 | 2 | 2 | 2 | 2 | 2 | 2 |
| GASEMM1714 | UK | Europe | 2015 | ND | emm165.0 | 277 | 131 | 161 | 113 | 89 | 106 | 100 | 215 | 265 | 77 | 2 | 131 | 25 |
| GASEMM0726 | UK | Europe | 2015 | ND | emm12.82 | 36 | 14 | 162 | 3 | 3 | 62 | 3 | 52 | 59 | 24 | 12 | 38 | 3 |
| GASEMM1431 | UK | Europe | 2015 | ND | emm58.7 | 176 | 14 | 13 | 10 | 8 | 13 | 13 | 195 | 47 | 10 | 2 | 34 | 31 |
| GASEMM1947 | UK | Europe | 2015 | ND | emm103.0 | 311 | 14 | 163 | 114 | 90 | 178 | 13 | 216 | 266 | 8 | 3 | 12 | 167 |

| | | | | | | | | | | | | | | | | | | |
|---|---|---|---|---|---|---|---|---|---|---|---|---|---|---|---|---|---|---|
| GASEMM1878 | UK | Europe | 2015 | ND | emm1.19 | 28 | 1 | 1 | 1 | 1 | 1 | 1 | 1 | 3 | 1 | 1 | 1 | 1 |
| GASEMM2496 | UK | Europe | 2015 | ND | emm112.2 | 497 | 132 | 107 | 4 | 66 | 21 | 61 | 27 | 161 | 8 | 3 | 21 | 113 |
| GASEMM1434 | UK | Europe | 2015 | ND | emm12.37 | 36 | 14 | 3 | 3 | 3 | 2 | 3 | 52 | 59 | 24 | 12 | 38 | 3 |
| GASEMM2044 | UK | Europe | 2015 | ND | emm5.120 | 99 | 14 | 91 | 70 | 1 | 17 | 71 | 114 | 131 | 47 | 22 | 80 | 91 |
| GASEMM0500 | UK | Europe | 2015 | ND | emm131.0 | 785 | 1 | 1 | 1 | 1 | 1 | 1 | 1 | 1 | 1 | 1 | 1 | 1 |
| GASEMM1541 | UK | Europe | 2015 | ND | emm90.2 | 184 | 14 | 88 | 68 | 57 | 94 | 70 | 109 | 127 | 1 | 2 | 79 | 88 |
| GASEMM0295 | UK | Europe | 2015 | ND | emm5.86 | 99 | 14 | 91 | 70 | 1 | 17 | 71 | 114 | 131 | 47 | 22 | 80 | 91 |
| GASEMM0584 | UK | Europe | 2015 | ND | emm3.91 | 15 | 127 | 15 | 10 | 1 | 10 | 32 | 111 | 129 | 1 | 2 | 6 | 31 |
| GASEMM1933 | UK | Europe | 2015 | ND | emm58.8 | 549 | 51 | 61 | 50 | 27 | 63 | 56 | 217 | 267 | 35 | 3 | 35 | 72 |
| GASEMM0956 | UK | Europe | 2015 | ND | emm131.0 | 99 | 14 | 91 | 70 | 1 | 17 | 71 | 114 | 263 | 47 | 22 | 80 | 91 |
| GASEMM2201 | UK | Europe | 2015 | ND | emm100.0 | 119 | 11 | 11 | 116 | 10 | 10 | 7 | 37 | 39 | 83 | 2 | 132 | 31 |
| H293 | UK | Europe | ND | invasive, NOS | emm89.0 | 101 | 2 | 2 | 2 | 2 | 2 | 2 | 2 | 2 | 2 | 2 | 2 | 2 |
| Alab49 | USA | North America | 1986 | ND | emm53.0 | 11 | 11 | 164 | 10 | 8 | 15 | 54 | 17 | 84 | 1 | 2 | 6 | 20 |
| D471 | USA | North America | ND | ARF | emm6.52 | 37 | 65 | 80 | 62 | 55 | 85 | 62 | 97 | 268 | 11 | 20 | 69 | 82 |
| FDAARGOS_149 | USA | North America | 2014 | invasive, NOS | emm1.0 | 28 | 1 | 1 | 1 | 1 | 1 | 1 | 1 | 134 | 1 | 1 | 1 | 1 |
| HarveyGAS | USA | North America | 2017 | necrotizing fasciitis | emm28.0 | 52 | 3 | 3 | 3 | 3 | 2 | 3 | 3 | 4 | 1 | 1 | 3 | 3 |
| Manfredo | USA | North America | 1952 | ARF | emm5.0 | 99 | [S] | 91 | 70 | 1 | 17 | 71 | 114 | 131 | 47 | 22 | 80 | 91 |
| MEW123 | USA | North America | ND | pharyngitis and/or tonsillitis | emm28.0 | 52 | 3 | 3 | 3 | 3 | 2 | 3 | 117 | 4 | 1 | 1 | 3 | 3 |
| MEW427 | USA | North America | ND | ND | emm4.0 | 39 | 4 | 8 | [S] | 7 | 51 | 9 | 8 | 9 | 7 | 2 | 8 | 8 |
| MGAS10270 | USA | North America | 2002 | pharyngitis and/or tonsillitis | emm2.0 | 55 | 32 | 26 | 28 | 24 | 29 | 13 | 43 | 48 | 15 | 3 | 12 | 42 |
| MGAS10394 | USA | North America | 1998 | pharyngitis and/or tonsillitis | emm6.4 | 382 | 65 | 80 | 62 | 55 | 85 | 62 | 218 | 130 | 11 | 20 | 69 | 82 |
| MGAS10750 | USA | North America | 2001 | pharyngitis and/or tonsillitis | emm4.0 | 1072 | 4 | 8 | 4 | 7 | 7 | 9 | 8 | 9 | 7 | 2 | 8 | 8 |
| MGAS11027 | USA | North America | ND | pharyngitis and/or tonsillitis | emm89.0 | 407 | 2 | 2 | 2 | 2 | 2 | 2 | 2 | 269 | 2 | 2 | 2 | 2 |
| MGAS1882 | USA | North America | ND | ND | emm59.0 | 172 | 71 | 84 | 24 | 2 | 93 | 68 | 107 | 122 | 18 | 3 | 75 | 85 |
| MGAS27061 | USA | North America | ND | invasive, NOS | emm89.0 | 101 | 2 | 2 | 119 | 58 | 2 | 2 | 2 | 2 | 2 | 2 | 2 | 2 |
| MGAS315 | USA | North America | ND | invasive, NOS | emm3.1 | 15 | 15 | 15 | 10 | 1 | 10 | 32 | 111 | 129 | 1 | 2 | 6 | 31 |
| MGAS5005 | USA | North America | 1996 | invasive, NOS | emm1.0 | 28 | 1 | 1 | 1 | 1 | 1 | 1 | 1 | 1 | 1 | 1 | 1 | 1 |
| MGAS6180 | USA | North America | 1998 | invasive, NOS | emm28.4 | 52 | 3 | 3 | 3 | 3 | 2 | 3 | 3 | 4 | 1 | 1 | 3 | 3 |
| MGAS8232 | USA | North America | 1987 | ARF | emm18.19 | 42 | 75 | 92 | 61 | 5 | 98 | 72 | 115 | 132 | 47 | 2 | 69 | 5 |
| MGAS9429 | USA | North America | 2001 | pharyngitis and/or tonsillitis | emm12.0 | 36 | 14 | 3 | 3 | 3 | 2 | 3 | 52 | 59 | 24 | 12 | 38 | 3 |
| SF370 | USA | North America | 1985 | ND | emm1.6 | 28 | 1 | 1 | 1 | 1 | 1 | 1 | 1 | 270 | 1 | 1 | 1 | 1 |
| 20161543 | USA | North America | 2015 | invasive, NOS | emm106.0 | 900 | 1 | 124 | 4 | 70 | 138 | 41 | 154 | 188 | 68 | 3 | 12 | 134 |
| 20163526 | USA | North America | 2015 | invasive, NOS | emm22.0 | 46 | 7 | 5 | 127 | 5 | 4 | 5 | 5 | 6 | 4 | 3 | 5 | 5 |
| 20162416 | USA | North America | 2015 | invasive, NOS | emm101.0 | 182 | 15 | 31 | 10 | 20 | 15 | 31 | 38 | 41 | 1 | 21 | 32 | 37 |
| 20162417 | USA | North America | 2015 | invasive, NOS | emm41.2 | 579 | 11 | 5 | 10 | 51 | 15 | 16 | 77 | 99 | 1 | 2 | 6 | 20 |
| 20165275 | USA | North America | 2015 | invasive, NOS | emm151.1 | 433 | 4 | 8 | 4 | 11 | 14 | 14 | 16 | 18 | 84 | 2 | 12 | 13 |
| 20162136 | USA | North America | 2015 | invasive, NOS | emm15.1 | 872 | 14 | 48 | 41 | 2 | 2 | 2 | 56 | 13 | 8 | 3 | 43 | 105 |
| 20162134 | USA | North America | 2015 | invasive, NOS | emm151.1 | 433 | 4 | 8 | 4 | 11 | 14 | 14 | 16 | 18 | 84 | 2 | 12 | 13 |
| 20162139 | USA | North America | 2015 | invasive, NOS | emm60.7 | 53 | 70 | 4 | 3 | 18 | 19 | 19 | 101 | 118 | 42 | 2 | 72 | 3 |
| 20160937 | USA | North America | 2015 | invasive, NOS | emm22.0 | 46 | 134 | 5 | 5 | 5 | 4 | 5 | 5 | 6 | 4 | 3 | 5 | 5 |
| 20170001 | USA | North America | 2015 | invasive, NOS | emm33.0 | 3 | 11 | 165 | 128 | 8 | 15 | 16 | 219 | 79 | 1 | 2 | 6 | 20 |
| 20162669 | USA | North America | 2015 | invasive, NOS | emm90.5 | 184 | 20 | 20 | 14 | 19 | 2 | 23 | 25 | 28 | 15 | 3 | 22 | 25 |
| 20161833 | USA | North America | 2015 | invasive, NOS | emm90.2 | 184 | 20 | 20 | 14 | 19 | 2 | 23 | 25 | 28 | 15 | 3 | 22 | 25 |
| 20162642 | USA | North America | 2015 | invasive, NOS | emm1.99 | 28 | 1 | 1 | 1 | 1 | 1 | 1 | 1 | 134 | 1 | 1 | 1 | 1 |
| 20162638 | USA | North America | 2015 | invasive, NOS | emm87.3 | 62 | 67 | 15 | 10 | 1 | 72 | 58 | 87 | 95 | 1 | 2 | 52 | 74 |
| 20162634 | USA | North America | 2015 | invasive, NOS | emm76.4 | 631 | 130 | 36 | 27 | 1 | 36 | 35 | 4 | 46 | 21 | 1 | 4 | 4 |
| 20162633 | USA | North America | 2015 | invasive, NOS | emm11.10 | ND | 1 | 97 | 40 | 61 | 20 | 20 | 13 | 139 | 8 | 3 | 18 | 4 |
| 20160980 | USA | North America | 2015 | invasive, NOS | emm49.1 | 29 | 4 | 8 | 4 | 11 | 11 | 14 | 12 | 271 | 9 | 2 | 12 | 13 |

| | | | | | | | | | | | | | | | | | | |
|---|---|---|---|---|---|---|---|---|---|---|---|---|---|---|---|---|---|---|
| 20160989 | USA | North America | 2015 | invasive, NOS | emm73.0 | 331 | 14 | 96 | 3 | 41 | 50 | 13 | 3 | 137 | 48 | 2 | 49 | 94 |
| 20161725 | USA | North America | 2015 | invasive, NOS | emm68.1 | 247 | 25 | 26 | 129 | 24 | 29 | 13 | 124 | 48 | 15 | 3 | 12 | 42 |
| 20161207 | USA | North America | 2015 | invasive, NOS | emm25.1 | 192 | 14 | 7 | 24 | 2 | 33 | 13 | 13 | 40 | 36 | 3 | 31 | 36 |
| 20161237 | USA | North America | 2015 | invasive, NOS | emm82.0 | 896 | 14 | 147 | 3 | 16 | 6 | 18 | 220 | 77 | 1 | 1 | 1 | 3 |
| 20162629 | USA | North America | 2015 | invasive, NOS | emm1.0 | 858 | 1 | 1 | 1 | 1 | 1 | 1 | 1 | 134 | 1 | 1 | 1 | 1 |
| 20162594 | USA | North America | 2015 | invasive, NOS | emm80.1 | 8 | 16 | 16 | 3 | 16 | 17 | 7 | 20 | 22 | 1 | 2 | 18 | 3 |
| 20160981 | USA | North America | 2015 | invasive, NOS | emm12.75 | 36 | 14 | 3 | 3 | 3 | 2 | 3 | 52 | 59 | 24 | 12 | 38 | 3 |
| 20162407 | USA | North America | 2015 | invasive, NOS | emm9.0 | 891 | 34 | 77 | 60 | 32 | 2 | 1 | 95 | 113 | 5 | 3 | 84 | 45 |
| 20162410 | USA | North America | 2015 | invasive, NOS | emm87.0 | 890 | 67 | 15 | 10 | 1 | 72 | 58 | 87 | 95 | 1 | 2 | 52 | 74 |
| 20161074 | USA | North America | 2015 | invasive, NOS | emm81.2 | 901 | 96 | 65 | 130 | 27 | 39 | 19 | 215 | 152 | 85 | 2 | 20 | 22 |
| 20160749 | USA | North America | 2015 | invasive, NOS | emm89.0 | 407 | 2 | 2 | 2 | 2 | 2 | 2 | 2 | 89 | 2 | 2 | 2 | 2 |
| 20160756 | USA | North America | 2015 | invasive, NOS | emm29.14 | 12 | 11 | 5 | 10 | 20 | 15 | 16 | 17 | 138 | 1 | 2 | 16 | 31 |
| 20161447 | USA | North America | 2015 | invasive, NOS | emm1.3 | 28 | 1 | 1 | 1 | 1 | 1 | 1 | 1 | 134 | 1 | 1 | 1 | 1 |
| 20161448 | USA | North America | 2015 | invasive, NOS | emm41.2 | 579 | 11 | 5 | 10 | 51 | 15 | 16 | 77 | 99 | 1 | 2 | 6 | 20 |
| 20161432 | USA | North America | 2015 | invasive, NOS | emm11.17 | ND | 1 | 97 | 40 | 61 | 20 | 20 | 120 | 139 | 8 | 3 | 85 | 95 |
| 20161430 | USA | North America | 2015 | invasive, NOS | emm68.4 | 894 | 25 | 26 | 20 | 24 | 29 | 13 | 124 | 48 | 49 | 3 | 12 | 42 |
| 20160762 | USA | North America | 2015 | invasive, NOS | emm219.0 | 25 | 14 | 58 | 125 | 1 | 6 | 18 | 69 | 255 | 1 | 1 | 127 | 162 |
| 20164926 | USA | North America | 2015 | invasive, NOS | emm103.0 | 327 | 14 | 13 | 3 | 1 | 66 | 13 | 82 | 87 | 1 | 2 | 49 | 4 |
| 20160970 | USA | North America | 2015 | invasive, NOS | emm9.2 | 891 | 34 | 77 | 60 | 32 | 2 | 1 | 95 | 113 | 5 | 3 | 84 | 45 |
| 20161736 | USA | North America | 2015 | invasive, NOS | emm11.16 | ND | 1 | 97 | 40 | 61 | 20 | 20 | 120 | 139 | 8 | 3 | 85 | 95 |
| 20161731 | USA | North America | 2015 | invasive, NOS | emm63.3 | 385 | 24 | 19 | 7 | 1 | 15 | 101 | 32 | 272 | 17 | 11 | 133 | 3 |
| 20161729 | USA | North America | 2015 | invasive, NOS | emm8.0 | 59 | 14 | 13 | 3 | 41 | 50 | 13 | 3 | 273 | 7 | 2 | 49 | 4 |
| 20161730 | USA | North America | 2015 | invasive, NOS | emm63.3 | 385 | 24 | 19 | 7 | 1 | 15 | 101 | 32 | 272 | 17 | 11 | 133 | 3 |
| 20161737 | USA | North America | 2015 | invasive, NOS | emm63.3 | 385 | 24 | 19 | 7 | 1 | 15 | 101 | 32 | 272 | 17 | 11 | 133 | 3 |
| 20161863 | USA | North America | 2015 | invasive, NOS | emm77.0 | 133 | 135 | 37 | 71 | 1 | 179 | 102 | 221 | 274 | 8 | 3 | 134 | 168 |
| 20162158 | USA | North America | 2015 | invasive, NOS | emm151.1 | 433 | 4 | 8 | 4 | 11 | 14 | 14 | 16 | 18 | 84 | 2 | 12 | 13 |
| 20161065 | USA | North America | 2015 | invasive, NOS | emm1.0 | 858 | 1 | 1 | 1 | 1 | 1 | 1 | 1 | 134 | 1 | 1 | 1 | 1 |
| 20161097 | USA | North America | 2015 | invasive, NOS | emm42.0 | 80 | 37 | 32 | 37 | 35 | 43 | 43 | 44 | 60 | 2 | 2 | 39 | 52 |
| 20161201 | USA | North America | 2015 | invasive, NOS | emm83.1 | 853 | 11 | 5 | 10 | 8 | 15 | 73 | 97 | 115 | 11 | 2 | 82 | 83 |
| 20161200 | USA | North America | 2015 | invasive, NOS | emm91.0 | 12 | 11 | 5 | 10 | 15 | 15 | 16 | 17 | 13 | 1 | 2 | 16 | 18 |
| 20161199 | USA | North America | 2015 | invasive, NOS | emm81.0 | 624 | 5 | 53 | 43 | 42 | 52 | 47 | 64 | 119 | 30 | 14 | 30 | 61 |
| 20161203 | USA | North America | 2015 | invasive, NOS | emm82.0 | 866 | 73 | 58 | 46 | 1 | 6 | 18 | 69 | 77 | 1 | 1 | 1 | 3 |
| 20160332 | USA | North America | 2015 | invasive, NOS | emm68.4 | 894 | 25 | 26 | 20 | 24 | 29 | 13 | 124 | 48 | 49 | 3 | 12 | 42 |
| 20160333 | USA | North America | 2015 | invasive, NOS | emm111.2 | 214 | 11 | 11 | 109 | 10 | 180 | 7 | 77 | 275 | 19 | 2 | 30 | 31 |
| 20162430 | USA | North America | 2015 | invasive, NOS | emm2.0 | 55 | 32 | 26 | 28 | 24 | 29 | 13 | 43 | 48 | 15 | 3 | 12 | 42 |
| 20162423 | USA | North America | 2015 | invasive, NOS | emm81.0 | 909 | 5 | 53 | 43 | 42 | 52 | 47 | 64 | 276 | 24 | 17 | 66 | 61 |
| 20162105 | USA | North America | 2015 | invasive, NOS | emm60.7 | 53 | 70 | 4 | 3 | 18 | 19 | 19 | 101 | 118 | 42 | 2 | 72 | 3 |
| 20162060 | USA | North America | 2015 | invasive, NOS | emm73.0 | 331 | 14 | 96 | 3 | 41 | 50 | 13 | 3 | 137 | 48 | 2 | 49 | 60 |
| 20161425 | USA | North America | 2015 | invasive, NOS | emm90.2 | 184 | 14 | 88 | 68 | 57 | 94 | 70 | 109 | 127 | 1 | 2 | 79 | 88 |
| 20160944 | USA | North America | 2015 | invasive, NOS | emm89.0 | 892 | 2 | 2 | 2 | 2 | 2 | 8 | 2 | 2 | 2 | 2 | 2 | 2 |
| 20160946 | USA | North America | 2015 | invasive, NOS | emm24.8 | 70 | 11 | 32 | 32 | 16 | 38 | 18 | 77 | 277 | 11 | 7 | 60 | 71 |
| 20160945 | USA | North America | 2015 | invasive, NOS | emm12.8 | 36 | 74 | 3 | 3 | 3 | 2 | 3 | 52 | 59 | 24 | 12 | 38 | 3 |
| 20161741 | USA | North America | 2015 | invasive, NOS | emm9.0 | 75 | 34 | 77 | 60 | 32 | 2 | 1 | 95 | 113 | 5 | 3 | 11 | 45 |
| 20161832 | USA | North America | 2015 | invasive, NOS | emm60.7 | 53 | 70 | 4 | 3 | 18 | 19 | 19 | 101 | 118 | 42 | 2 | 72 | 3 |
| 20162146 | USA | North America | 2015 | invasive, NOS | emm89.0 | 101 | 2 | 2 | 2 | 2 | 2 | 2 | 2 | [S] | 2 | 2 | 2 | 2 |
| 20164918 | USA | North America | 2015 | invasive, NOS | emm4.0 | 911 | 4 | 8 | 4 | 7 | 51 | 9 | 8 | 9 | 7 | 2 | 8 | 8 |
| 20164915 | USA | North America | 2015 | invasive, NOS | emm82.7 | 36 | 14 | 3 | 3 | 3 | 2 | 3 | 52 | 59 | 24 | 12 | 38 | 3 |
| 20161091 | USA | North America | 2015 | invasive, NOS | emm9.0 | 75 | 34 | 77 | 60 | 32 | 2 | 1 | 95 | 113 | 5 | 3 | 11 | 45 |

| | | | | | | | | | | | | | | | | | | |
|---|---|---|---|---|---|---|---|---|---|---|---|---|---|---|---|---|---|---|
| 20162124 | USA | North America | 2015 | invasive, NOS | emm28.0 | 456 | 3 | 3 | 3 | 3 | 2 | 3 | 3 | 4 | 1 | 1 | 3 | 3 |
| 20160491 | USA | North America | 2015 | invasive, NOS | emm108.1 | 14 | 28 | 87 | 10 | 1 | 6 | 63 | 78 | 240 | 1 | 2 | 6 | 31 |
| 20162118 | USA | North America | 2015 | invasive, NOS | emm81.3 | 903 | 8 | 166 | 4 | 4 | 181 | 13 | 222 | 42 | 31 | 1 | 135 | 4 |
| 20162110 | USA | North America | 2015 | invasive, NOS | emm90.5 | 184 | 20 | 20 | 14 | 19 | 2 | 23 | 25 | 28 | 15 | 3 | 22 | 25 |
| 20162109 | USA | North America | 2015 | invasive, NOS | emm227.1 | 28 | 1 | 1 | 1 | 1 | 1 | 1 | 1 | 134 | 1 | 1 | 1 | 1 |
| 20162115 | USA | North America | 2015 | invasive, NOS | emm28.0 | 456 | 3 | 3 | 3 | 3 | 2 | 3 | 3 | 4 | 1 | 1 | 3 | 3 |
| 20161333 | USA | North America | 2015 | invasive, NOS | emm2.0 | 55 | 32 | 26 | 28 | 24 | 29 | 13 | 43 | 48 | 15 | 3 | 12 | 42 |
| 20163031 | USA | North America | 2015 | invasive, NOS | emm27.0 | 308 | 10 | 85 | 65 | 56 | 1 | 69 | 104 | 18 | 9 | 2 | 12 | 13 |
| 20160948 | USA | North America | 2015 | invasive, NOS | emm89.0 | 101 | 2 | 2 | 2 | 2 | 2 | 2 | 2 | 2 | 2 | 2 | 2 | 2 |
| 20160953 | USA | North America | 2015 | invasive, NOS | emm77.0 | 904 | 14 | 110 | 3 | 22 | 27 | 13 | 26 | 29 | 8 | 3 | 25 | 30 |
| 20160951 | USA | North America | 2015 | invasive, NOS | emm73.0 | 331 | 14 | 96 | 3 | 41 | 50 | 13 | 3 | 137 | 48 | 2 | 49 | 60 |
| 20160955 | USA | North America | 2015 | invasive, NOS | emm91.0 | 12 | 11 | 5 | 10 | 15 | 15 | 16 | 17 | 13 | 1 | 2 | 16 | 18 |
| 20155415 | USA | North America | 2015 | invasive, NOS | emm11.1 | ND | 1 | 97 | 40 | 61 | 20 | 20 | 13 | 139 | 8 | 3 | 18 | 4 |
| 20160028 | USA | North America | 2015 | invasive, NOS | emm63.0 | 210 | 5 | 167 | 34 | 27 | 39 | 13 | 84 | 279 | 13 | 2 | 20 | 22 |
| 20156364 | USA | North America | 2015 | invasive, NOS | emm77.0 | 399 | 82 | 25 | 3 | 22 | 27 | 13 | 26 | 29 | 8 | 3 | 25 | 30 |
| 20155408 | USA | North America | 2015 | invasive, NOS | emm165.0 | 768 | 58 | 7 | 131 | 1 | 76 | 103 | 27 | 102 | 8 | 3 | 61 | 29 |
| 20156321 | USA | North America | 2015 | invasive, NOS | emm75.0 | 150 | 25 | 26 | 20 | 24 | 29 | 29 | 33 | 48 | 15 | 3 | 12 | 33 |
| 20156324 | USA | North America | 2015 | invasive, NOS | emm77.0 | 399 | 82 | 25 | 3 | 22 | 27 | 13 | 26 | 29 | 8 | 3 | 25 | 30 |
| 20156359 | USA | North America | 2015 | invasive, NOS | emm58.2 | 176 | 14 | 13 | 10 | 8 | 13 | 13 | 194 | 47 | 10 | 2 | 34 | 31 |
| 20156362 | USA | North America | 2015 | invasive, NOS | emm22.0 | 46 | 7 | 5 | 127 | 5 | 4 | 5 | 5 | 6 | 4 | 3 | 5 | 5 |
| 20155617 | USA | North America | 2015 | invasive, NOS | emm75.0 | 49 | 25 | 26 | 20 | 24 | 29 | 74 | 33 | 136 | 15 | 3 | 12 | 33 |
| 20156430 | USA | North America | 2015 | invasive, NOS | emm81.0 | 837 | 5 | 53 | 43 | 42 | 52 | 47 | 64 | 119 | 30 | 14 | 30 | 61 |
| 20156432 | USA | North America | 2015 | invasive, NOS | emm238.2 | 867 | 1 | 1 | 1 | 1 | 1 | 15 | 14 | 93 | 5 | 6 | 14 | 15 |
| 20156319 | USA | North America | 2015 | invasive, NOS | emm68.3 | 247 | 25 | 26 | 4 | 24 | 29 | 13 | 124 | 48 | 15 | 3 | 12 | 42 |
| 20155197 | USA | North America | 2015 | invasive, NOS | emm81.0 | 624 | 5 | 53 | 43 | 42 | 52 | 47 | 64 | 119 | 86 | 14 | 30 | 61 |
| 20155204 | USA | North America | 2015 | invasive, NOS | emm9.2 | 891 | 34 | 77 | 60 | 32 | 2 | 1 | 95 | 113 | 5 | 3 | 84 | 45 |
| 20155202 | USA | North America | 2015 | invasive, NOS | emm87.0 | 62 | 67 | 15 | 10 | 1 | 72 | 58 | 87 | 95 | 23 | 2 | 52 | 74 |
| 20155579 | USA | North America | 2015 | invasive, NOS | emm77.0 | 133 | 135 | 37 | 71 | 1 | 179 | 102 | 221 | 274 | 8 | 3 | 134 | 168 |
| 20155417 | USA | North America | 2015 | invasive, NOS | emm44.0 | 641 | 5 | 7 | 7 | 1 | 81 | 13 | 26 | 280 | 8 | 18 | 25 | 30 |
| 20155580 | USA | North America | 2015 | invasive, NOS | emm77.0 | 133 | 135 | 37 | 71 | 1 | 179 | 102 | 221 | 274 | 8 | 3 | 134 | 168 |
| 20155581 | USA | North America | 2015 | invasive, NOS | emm89.0 | 407 | 2 | 2 | 2 | 2 | 2 | 2 | 2 | 281 | 2 | 2 | 2 | 2 |
| 20155208 | USA | North America | 2015 | invasive, NOS | emm4.0 | 39 | 4 | 8 | 4 | 7 | 51 | 9 | 8 | 9 | 7 | 2 | 8 | 8 |
| 20155209 | USA | North America | 2015 | invasive, NOS | emm76.0 | 50 | 136 | 3 | 3 | 3 | 2 | 3 | 52 | 59 | 24 | 12 | 38 | 3 |
| 20155267 | USA | North America | 2015 | invasive, NOS | emm49.0 | 433 | 4 | 8 | 4 | 11 | 14 | 14 | 16 | 18 | 84 | 2 | 12 | 13 |
| 20155269 | USA | North America | 2015 | invasive, NOS | emm87.0 | 62 | 67 | 15 | 10 | 1 | 72 | 58 | 87 | 95 | 1 | 2 | 52 | 74 |
| 20160177 | USA | North America | 2015 | invasive, NOS | emm28.0 | 458 | 3 | 3 | 3 | 3 | 2 | 3 | 3 | 4 | 1 | 1 | 3 | 3 |
| 20156635 | USA | North America | 2015 | invasive, NOS | emm28.0 | 859 | 3 | 3 | 3 | 3 | 2 | 3 | 117 | 4 | 1 | 1 | 3 | 3 |
| 20156154 | USA | North America | 2015 | invasive, NOS | emm81.0 | 837 | 5 | 53 | 43 | 42 | 52 | 47 | 64 | 119 | 30 | 14 | 30 | 61 |
| 20160329 | USA | North America | 2015 | invasive, NOS | emm101.0 | 182 | 15 | 31 | 10 | 20 | 15 | 31 | 38 | 41 | 1 | 21 | 32 | 169 |
| 20156401 | USA | North America | 2015 | invasive, NOS | emm83.1 | 5 | 11 | 5 | 10 | 8 | 15 | 65 | 97 | 282 | 11 | 2 | 69 | 83 |
| 20156400 | USA | North America | 2015 | invasive, NOS | emm104.0 | 137 | 8 | 168 | 71 | 1 | 1 | 64 | 70 | 1 | 1 | 1 | 1 | 3 |
| 20160016 | USA | North America | 2015 | invasive, NOS | emm28.0 | 458 | 3 | 3 | 3 | 3 | 2 | 3 | 3 | 4 | 1 | 1 | 3 | 3 |
| 20156270 | USA | North America | 2015 | invasive, NOS | emm82.0 | 866 | 73 | 58 | 46 | 1 | 6 | 18 | 69 | 77 | 1 | 1 | 1 | 3 |
| 20156271 | USA | North America | 2015 | invasive, NOS | emm75.0 | 150 | 25 | 26 | 20 | 24 | 29 | 29 | 33 | 155 | 15 | 3 | 12 | 33 |
| 20155380 | USA | North America | 2015 | invasive, NOS | emm58.0 | 176 | 14 | 94 | 10 | 8 | 13 | 13 | 15 | 47 | 10 | 2 | 34 | 31 |
| 20155381 | USA | North America | 2015 | invasive, NOS | emm81.0 | 837 | 5 | 53 | 43 | 42 | 52 | 47 | 64 | 119 | 30 | 14 | 30 | 61 |
| 20160140 | USA | North America | 2015 | invasive, NOS | emm12.91 | 36 | 14 | 3 | 3 | 3 | 2 | 3 | 52 | 59 | 24 | 12 | 38 | 3 |
| 20156447 | USA | North America | 2015 | invasive, NOS | emm12.32 | 36 | 14 | 3 | 3 | 3 | 2 | 3 | 52 | 59 | 24 | 12 | 38 | 3 |

| | | | | | | | | | | | | | | | | | | |
|---|---|---|---|---|---|---|---|---|---|---|---|---|---|---|---|---|---|---|
| 20156102 | USA | North America | 2015 | invasive, NOS | emm118.0 | 863 | 11 | 32 | 52 | 1 | 67 | 13 | 118 | 135 | 1 | 23 | 83 | 93 |
| 20156448 | USA | North America | 2015 | invasive, NOS | emm104.0 | 137 | 8 | 168 | 71 | 1 | 1 | 64 | 70 | 1 | 1 | 1 | 1 | 3 |
| 20156019 | USA | North America | 2015 | invasive, NOS | emm6.80 | 382 | 65 | 80 | 62 | 55 | 85 | 62 | 97 | 130 | 11 | 20 | 69 | 82 |
| 20155585 | USA | North America | 2015 | invasive, NOS | emm77.0 | 133 | 135 | 37 | 71 | 1 | 179 | 102 | 221 | 274 | 8 | 3 | 134 | 168 |
| 20155589 | USA | North America | 2015 | invasive, NOS | emm75.0 | 861 | 25 | 26 | 20 | 24 | 29 | 74 | 33 | 283 | 15 | 3 | 12 | 33 |
| 20155588 | USA | North America | 2015 | invasive, NOS | emm75.0 | 150 | 25 | 26 | 20 | 24 | 29 | 29 | 33 | 48 | 15 | 3 | 12 | 33 |
| 20160010 | USA | North America | 2015 | invasive, NOS | emm89.0 | 101 | 2 | 2 | 2 | 58 | 2 | 2 | 2 | 2 | 2 | 2 | 2 | 2 |
| 20156107 | USA | North America | 2015 | invasive, NOS | emm76.0 | 50 | 14 | 3 | 3 | 3 | 2 | 3 | 52 | 59 | 24 | 12 | 38 | 3 |
| 20156105 | USA | North America | 2015 | invasive, NOS | emm59.0 | 172 | 72 | 84 | 24 | 2 | 93 | 68 | 107 | 122 | 18 | 3 | 75 | 85 |
| 20156103 | USA | North America | 2015 | invasive, NOS | emm58.0 | 176 | 14 | 94 | 10 | 8 | 13 | 13 | 15 | 47 | 10 | 2 | 34 | 31 |
| 20155639 | USA | North America | 2015 | invasive, NOS | emm76.0 | 50 | 14 | 3 | 3 | 3 | 2 | 3 | 52 | 59 | 24 | 12 | 38 | 3 |
| 20160313 | USA | North America | 2015 | invasive, NOS | emm11.18 | ND | 1 | 97 | 40 | 61 | 20 | 20 | 13 | 139 | 8 | 3 | 18 | 4 |
| 20156730 | USA | North America | 2015 | invasive, NOS | emm68.3 | 247 | 25 | 26 | 4 | 24 | 29 | 13 | 124 | 48 | 15 | 3 | 12 | 42 |
| 20156725 | USA | North America | 2015 | invasive, NOS | emm102.2 | 895 | 14 | 93 | 71 | 60 | 99 | 13 | 116 | 133 | 13 | 2 | 81 | 92 |
| 20156726 | USA | North America | 2015 | invasive, NOS | emm58.0 | 176 | 14 | 94 | 10 | 8 | 13 | 13 | 15 | 47 | 10 | 2 | 34 | 31 |
| 20156185 | USA | North America | 2015 | invasive, NOS | emm114.6 | 220 | 18 | 136 | 103 | 43 | 148 | 40 | 26 | 56 | 5 | 27 | 83 | 147 |
| 20156188 | USA | North America | 2015 | invasive, NOS | emm25.1 | 192 | 14 | 7 | 24 | 2 | 33 | 13 | 13 | 40 | 36 | 3 | 31 | 36 |
| 20155373 | USA | North America | 2015 | invasive, NOS | emm9.0 | 75 | 34 | 77 | 60 | 32 | 2 | 1 | 95 | 113 | 5 | 3 | 11 | 45 |
| 20156592 | USA | North America | 2015 | invasive, NOS | emm83.1 | 853 | 11 | 5 | 10 | 8 | 15 | 73 | 97 | 115 | 11 | 2 | 82 | 83 |
| 20160031 | USA | North America | 2015 | invasive, NOS | emm232.0 | 902 | 14 | 37 | 31 | 3 | 2 | 38 | 47 | 242 | 1 | 2 | 58 | 25 |
| 20156457 | USA | North America | 2015 | invasive, NOS | emm92.2 | 82 | 17 | 4 | 3 | 18 | 19 | 19 | 22 | 24 | 13 | 2 | 20 | 22 |
| 20155603 | USA | North America | 2015 | invasive, NOS | emm9.2 | 891 | 34 | 77 | 60 | 32 | 2 | 1 | 95 | 113 | 5 | 3 | 84 | 45 |
| 20156711 | USA | North America | 2015 | invasive, NOS | emm18.7 | ND | 75 | 92 | 61 | 5 | 98 | 72 | 115 | 132 | 47 | 2 | 69 | 5 |
| 20155386 | USA | North America | 2015 | invasive, NOS | emm83.1 | 853 | 11 | 5 | 10 | 8 | 15 | 73 | 97 | 115 | 11 | 2 | 82 | 83 |
| 20155657 | USA | North America | 2015 | invasive, NOS | emm49.0 | 862 | 4 | 8 | 4 | 11 | 14 | 14 | 16 | 18 | 9 | 2 | 12 | 13 |
| 20155393 | USA | North America | 2015 | invasive, NOS | emm87.0 | 62 | 67 | 15 | 10 | 1 | 72 | 58 | 87 | 95 | 1 | 2 | 52 | 74 |
| 20156790 | USA | North America | 2015 | invasive, NOS | emm28.0 | 458 | 3 | 3 | 3 | 3 | 2 | 3 | 3 | 4 | 1 | 1 | 3 | 3 |
| 20156794 | USA | North America | 2015 | invasive, NOS | emm28.0 | 458 | 3 | 3 | 3 | 3 | 2 | 3 | 3 | 4 | 1 | 1 | 3 | 3 |
| 20156713 | USA | North America | 2015 | invasive, NOS | emm9.0 | 75 | 34 | 77 | 60 | 32 | 2 | 1 | 95 | 113 | 5 | 3 | 11 | 45 |
| 20156717 | USA | North America | 2015 | invasive, NOS | emm11.16 | ND | 1 | 97 | 40 | 61 | 20 | 20 | 120 | 139 | 8 | 3 | 85 | 95 |
| 20156781 | USA | North America | 2015 | invasive, NOS | emm8.0 | 59 | 14 | 13 | 3 | 41 | 50 | 13 | 3 | 273 | 7 | 2 | 49 | 4 |
| 20155615 | USA | North America | 2015 | invasive, NOS | emm27.0 | 308 | 10 | 85 | 65 | 56 | 1 | 69 | 104 | [S] | 9 | 2 | 12 | 13 |
| 20155609 | USA | North America | 2015 | invasive, NOS | emm75.0 | 49 | 25 | 26 | 20 | 24 | 29 | 74 | 33 | 136 | 15 | 3 | 12 | 33 |
| 20155360 | USA | North America | 2015 | invasive, NOS | emm89.0 | 407 | 2 | 2 | 2 | 2 | 2 | 2 | 2 | 89 | 2 | 2 | 2 | 170 |
| 20155356 | USA | North America | 2015 | invasive, NOS | emm81.0 | 837 | 5 | 53 | 43 | 42 | 52 | 47 | 64 | 119 | 30 | 14 | 30 | 61 |
| 20156595 | USA | North America | 2015 | invasive, NOS | emm8.0 | 869 | 18 | 169 | 4 | 8 | 64 | 50 | 223 | 285 | 18 | 3 | 83 | 171 |
| 20160246 | USA | North America | 2015 | invasive, NOS | emm82.0 | 866 | 73 | 58 | 46 | 1 | 6 | 18 | 69 | 77 | 1 | 1 | 1 | 3 |
| 20156593 | USA | North America | 2015 | invasive, NOS | emm238.1 | 868 | 1 | 20 | 123 | 1 | 1 | 15 | 14 | 16 | 5 | 6 | 14 | 15 |
| 20156622 | USA | North America | 2015 | invasive, NOS | emm73.0 | 331 | 14 | 96 | 3 | 41 | 50 | 13 | 3 | 286 | 48 | 2 | 49 | 60 |
| 20156597 | USA | North America | 2015 | invasive, NOS | emm91.0 | 12 | 11 | 5 | 10 | 15 | 15 | 16 | 17 | 13 | 1 | 2 | 16 | 18 |
| 20156169 | USA | North America | 2015 | invasive, NOS | emm63.3 | 385 | 24 | 19 | 7 | 1 | 15 | 101 | 32 | 272 | 17 | 11 | 133 | 3 |
| 20156012 | USA | North America | 2015 | invasive, NOS | emm77.0 | 399 | 82 | 25 | 3 | 22 | 27 | 13 | 26 | 29 | 8 | 3 | 25 | 30 |
| 20156165 | USA | North America | 2015 | invasive, NOS | emm101.0 | 182 | 15 | 31 | 10 | 20 | 15 | 31 | 38 | 41 | 1 | 21 | 32 | 37 |
| 20155387 | USA | North America | 2015 | invasive, NOS | emm89.0 | 860 | 2 | 2 | 2 | 58 | 2 | 2 | 2 | 2 | 2 | 2 | 2 | 2 |
| 20155047 | USA | North America | 2015 | invasive, NOS | emm5.128 | 99 | 14 | 91 | 70 | 1 | 17 | 71 | 114 | 131 | 47 | 22 | 80 | 91 |
| 20155046 | USA | North America | 2015 | invasive, NOS | emm5.18 | 99 | 14 | 91 | 70 | 1 | 17 | 71 | 114 | 131 | 47 | 22 | 80 | 91 |
| 20155042 | USA | North America | 2015 | invasive, NOS | emm101.0 | 182 | 15 | 31 | 10 | 20 | 15 | 31 | 38 | 41 | 1 | 21 | 32 | 37 |
| 20156410 | USA | North America | 2015 | invasive, NOS | emm12.75 | 36 | 14 | 170 | 3 | 3 | 2 | 3 | 52 | 59 | 24 | 12 | 38 | 3 |

| | | | | | | | | | | | | | | | | | | |
|---|---|---|---|---|---|---|---|---|---|---|---|---|---|---|---|---|---|---|
| 20156414 | USA | North America | 2015 | invasive, NOS | emm1.0 | 897 | 1 | 1 | 1 | 1 | 1 | 1 | 224 | 134 | 1 | 1 | 1 | 1 |
| 20156413 | USA | North America | 2015 | invasive, NOS | emm49.0 | 433 | 4 | 8 | 4 | 11 | 14 | 14 | 16 | 18 | 84 | 2 | 12 | 13 |
| 20154297 | USA | North America | 2015 | invasive, NOS | emm59.2 | 864 | 72 | 84 | 24 | 2 | 93 | 68 | 107 | 122 | 18 | 3 | 75 | 85 |
| 20154375 | USA | North America | 2015 | invasive, NOS | emm11.0 | ND | 1 | 6 | 40 | 78 | 20 | 20 | 13 | 139 | 8 | 3 | 18 | 4 |
| 20154055 | USA | North America | 2015 | invasive, NOS | emm92.0 | 855 | 17 | 4 | 3 | 18 | 19 | 19 | 22 | 24 | 13 | 2 | 20 | 22 |
| 20154137 | USA | North America | 2015 | invasive, NOS | emm12.7 | 36 | 14 | 3 | 3 | 3 | 2 | 3 | 52 | 59 | 24 | 12 | 38 | 3 |
| 20154144 | USA | North America | 2015 | invasive, NOS | emm41.2 | 579 | 11 | 5 | 10 | 51 | 15 | 16 | 77 | 99 | 1 | 2 | 6 | 20 |
| 20154142 | USA | North America | 2015 | invasive, NOS | emm118.0 | 167 | 11 | 32 | 52 | 1 | 67 | 13 | 118 | 135 | 1 | 23 | 83 | 93 |
| 20154591 | USA | North America | 2015 | invasive, NOS | emm89.0 | 801 | 137 | 2 | 2 | 58 | 2 | 2 | 2 | 2 | 2 | 2 | 2 | 2 |
| 20154590 | USA | North America | 2015 | invasive, NOS | emm219.0 | 25 | 14 | 58 | 125 | 1 | 6 | 18 | 69 | 255 | 1 | 1 | 127 | 162 |
| 20155018 | USA | North America | 2015 | invasive, NOS | emm77.0 | 63 | 23 | 24 | 19 | 1 | 26 | 28 | 30 | 33 | 8 | 3 | 24 | 172 |
| 20155015 | USA | North America | 2015 | invasive, NOS | emm83.1 | 5 | 11 | 5 | 10 | 8 | 15 | 65 | 97 | 282 | 11 | 2 | 69 | 83 |
| 20154949 | USA | North America | 2015 | invasive, NOS | emm83.1 | 853 | 11 | 5 | 10 | 8 | 15 | 73 | 97 | 115 | 11 | 2 | 82 | 83 |
| 20154031 | USA | North America | 2015 | invasive, NOS | emm12.0 | 36 | 14 | 3 | 3 | 3 | 2 | 3 | 52 | 287 | 24 | 12 | 38 | 3 |
| 20154030 | USA | North America | 2015 | invasive, NOS | emm76.0 | 50 | 14 | 3 | 3 | 3 | 2 | 3 | 52 | 59 | 24 | 12 | 38 | 3 |
| 20154029 | USA | North America | 2015 | invasive, NOS | emm82.0 | 334 | 138 | 58 | 46 | 1 | 6 | 18 | 69 | 288 | 1 | 1 | 1 | 3 |
| 20154028 | USA | North America | 2015 | invasive, NOS | emm60.7 | 53 | 70 | 4 | 3 | 18 | 19 | 19 | 101 | 118 | 42 | 2 | 72 | 3 |
| 20154026 | USA | North America | 2015 | invasive, NOS | emm89.0 | 101 | 2 | 2 | 2 | 58 | 2 | 2 | 2 | 2 | 2 | 2 | 2 | 2 |
| 20154024 | USA | North America | 2015 | invasive, NOS | emm28.0 | 52 | 3 | 3 | 3 | 3 | 2 | 3 | 3 | 4 | 1 | 1 | 3 | 3 |
| 20154025 | USA | North America | 2015 | invasive, NOS | emm11.0 | ND | 1 | 97 | 40 | 61 | 20 | 20 | 120 | 139 | 8 | 3 | 85 | 95 |
| 20154805 | USA | North America | 2015 | invasive, NOS | emm81.0 | 624 | 5 | 53 | 43 | 42 | 52 | 47 | 64 | 119 | 30 | 14 | 30 | 61 |
| 20154799 | USA | North America | 2015 | invasive, NOS | emm3.1 | 15 | 15 | 15 | 10 | 1 | 10 | 32 | 111 | 129 | 1 | 2 | 6 | 31 |
| 20154797 | USA | North America | 2015 | invasive, NOS | emm6.101 | 382 | 65 | 80 | 62 | 55 | 182 | 62 | 97 | 130 | 11 | 20 | 69 | 82 |
| 20154803 | USA | North America | 2015 | invasive, NOS | emm1.0 | 899 | 1 | 1 | 1 | 1 | 1 | 1 | 1 | 1 | 1 | 1 | 1 | 1 |
| 20154293 | USA | North America | 2015 | invasive, NOS | emm59.0 | 864 | 72 | 84 | 24 | 2 | 93 | 68 | 107 | 122 | 18 | 3 | 75 | 85 |
| 20154292 | USA | North America | 2015 | invasive, NOS | emm87.0 | 62 | 67 | 15 | 10 | 1 | 72 | 58 | 87 | 95 | 1 | 2 | 52 | 74 |
| 20154289 | USA | North America | 2015 | invasive, NOS | emm59.0 | 864 | 72 | 84 | 24 | 2 | 93 | 68 | 107 | 122 | 18 | 3 | 75 | 85 |
| 20154288 | USA | North America | 2015 | invasive, NOS | emm59.2 | 864 | 72 | 84 | 24 | 2 | 93 | 68 | 107 | 122 | 18 | 3 | 75 | 85 |
| 20154168 | USA | North America | 2015 | invasive, NOS | emm12.7 | 36 | 14 | 3 | 3 | 3 | 2 | 3 | 52 | 59 | 24 | 12 | 38 | 3 |
| 20154167 | USA | North America | 2015 | invasive, NOS | emm4.0 | 39 | 4 | 8 | 4 | 7 | 51 | 9 | 8 | 9 | 7 | 2 | 8 | 8 |
| 20154813 | USA | North America | 2015 | invasive, NOS | emm3.4 | 15 | 15 | 15 | 10 | 1 | 10 | 32 | 111 | 129 | 1 | 2 | 6 | 31 |
| 20154042 | USA | North America | 2015 | invasive, NOS | emm108.2 | 14 | 28 | 87 | 10 | 1 | 183 | 91 | 78 | 240 | 1 | 2 | 6 | 31 |
| 20154038 | USA | North America | 2015 | invasive, NOS | emm92.0 | 82 | 17 | 4 | 3 | 18 | 19 | 19 | 22 | 24 | 13 | 2 | 20 | 22 |
| 20154035 | USA | North America | 2015 | invasive, NOS | emm12.0 | 36 | 14 | 3 | 3 | 3 | 2 | 3 | 52 | 252 | 24 | 12 | 38 | 3 |
| 20154043 | USA | North America | 2015 | invasive, NOS | emm83.1 | 5 | 11 | 5 | 10 | 8 | 15 | 65 | 97 | 282 | 11 | 2 | 69 | 83 |
| 20154036 | USA | North America | 2015 | invasive, NOS | emm92.0 | 82 | 17 | 4 | 3 | 18 | 19 | 19 | 22 | 24 | 13 | 2 | 20 | 22 |
| 20154044 | USA | North America | 2015 | invasive, NOS | emm4.0 | 39 | 4 | 8 | 4 | 7 | 51 | 9 | 8 | 9 | 7 | 2 | 8 | 8 |
| 20154565 | USA | North America | 2015 | invasive, NOS | emm89.0 | 910 | 2 | 2 | 2 | 2 | 2 | 2 | 2 | 89 | 2 | 2 | 2 | 2 |
| 20154558 | USA | North America | 2015 | invasive, NOS | emm77.0 | 63 | 23 | 24 | 57 | 1 | 184 | 28 | 30 | 33 | 8 | 3 | 24 | 29 |
| 20154557 | USA | North America | 2015 | invasive, NOS | emm1.0 | 856 | 1 | 1 | 1 | 1 | 1 | 1 | 224 | 134 | 1 | 1 | 1 | 1 |
| 20154555 | USA | North America | 2015 | invasive, NOS | emm28.0 | 52 | 3 | 3 | 3 | 3 | 2 | 3 | 117 | 4 | 1 | 1 | 3 | 3 |
| 20154563 | USA | North America | 2015 | invasive, NOS | emm106.0 | 338 | 14 | 27 | 132 | 25 | 106 | 19 | 225 | 36 | 15 | 3 | 27 | 34 |
| 20154623 | USA | North America | 2015 | invasive, NOS | emm1.49 | 28 | 1 | 1 | 1 | 1 | 1 | 1 | 1 | 134 | 1 | 1 | 1 | 1 |
| 20154628 | USA | North America | 2015 | invasive, NOS | emm75.0 | 150 | 25 | 26 | 20 | 24 | 29 | 29 | 33 | 48 | 15 | 3 | 12 | 33 |
| 20154166 | USA | North America | 2015 | invasive, NOS | emm3.1 | 15 | 15 | 15 | 10 | 1 | 10 | 32 | 111 | 129 | 1 | 2 | 6 | 31 |
| 20154161 | USA | North America | 2015 | invasive, NOS | emm59.0 | 172 | 72 | 84 | 24 | 79 | 93 | 68 | 107 | 122 | 18 | 3 | 75 | 85 |
| 20154158 | USA | North America | 2015 | invasive, NOS | emm11.0 | ND | 1 | 97 | 40 | 61 | 20 | 20 | 120 | 139 | 8 | 3 | 85 | 95 |
| 20154946 | USA | North America | 2015 | invasive, NOS | emm12.7 | 36 | 14 | 3 | 3 | 3 | 2 | 3 | 52 | 59 | 24 | 12 | 38 | 3 |

| | | | | | | | | | | | | | | | | | | |
|---|---|---|---|---|---|---|---|---|---|---|---|---|---|---|---|---|---|---|
| 20154547 | USA | North America | 2015 | invasive, NOS | emm89.0 | 407 | 2 | 2 | 2 | 2 | 2 | 2 | 2 | 89 | 2 | 2 | 2 | 170 |
| 20154548 | USA | North America | 2015 | invasive, NOS | emm89.0 | 101 | 2 | 2 | 2 | 2 | 2 | 2 | 2 | 2 | 2 | 2 | 2 | 2 |
| 20154550 | USA | North America | 2015 | invasive, NOS | emm6.0 | 382 | 65 | 80 | 133 | 55 | 85 | 62 | 97 | 130 | 11 | 20 | 69 | 82 |
| 20154944 | USA | North America | 2015 | invasive, NOS | emm28.0 | 850 | 3 | 3 | 3 | 3 | 2 | 3 | 3 | 4 | 1 | 1 | 3 | 3 |
| 20154821 | USA | North America | 2015 | invasive, NOS | emm1.0 | 858 | 1 | 1 | 1 | 1 | 1 | 1 | 1 | 134 | 1 | 1 | 1 | 1 |
| 20155031 | USA | North America | 2015 | invasive, NOS | emm3.1 | 15 | 15 | 15 | 10 | 1 | 10 | 32 | 111 | 129 | 1 | 2 | 6 | 31 |
| 20155025 | USA | North America | 2015 | invasive, NOS | emm54.1 | 302 | 11 | 5 | 10 | 20 | 15 | 16 | 17 | 13 | 1 | 2 | 16 | 31 |
| 20154049 | USA | North America | 2015 | invasive, NOS | emm12.0 | 36 | 14 | 3 | 3 | 3 | 2 | 3 | 52 | 287 | 24 | 12 | 38 | 3 |
| 20154051 | USA | North America | 2015 | invasive, NOS | emm82.0 | 36 | 14 | 3 | 3 | 3 | 2 | 3 | 52 | 59 | 24 | 12 | 38 | 3 |
| 20154046 | USA | North America | 2015 | invasive, NOS | emm18.12 | 535 | 14 | 57 | 2 | 2 | 2 | 78 | 67 | 75 | 8 | 3 | 53 | 2 |
| 20154045 | USA | North America | 2015 | invasive, NOS | emm77.0 | 63 | 23 | 24 | 57 | 1 | 26 | 28 | 30 | 33 | 8 | 3 | 24 | 29 |
| 20154048 | USA | North America | 2015 | invasive, NOS | emm49.0 | 433 | 4 | 8 | 4 | 11 | 14 | 14 | 16 | 18 | 84 | 2 | 12 | 13 |
| 20154047 | USA | North America | 2015 | invasive, NOS | emm49.0 | 433 | 4 | 8 | 4 | 11 | 14 | 14 | 16 | 18 | 84 | 2 | 12 | 13 |
| 20154054 | USA | North America | 2015 | invasive, NOS | emm92.0 | 82 | 17 | 4 | 3 | 18 | 19 | 19 | 22 | 24 | 13 | 2 | 20 | 22 |
| 20154053 | USA | North America | 2015 | invasive, NOS | emm92.0 | 82 | 17 | 4 | 3 | 18 | 19 | 19 | 22 | 24 | 13 | 2 | 20 | 22 |
| 20154615 | USA | North America | 2015 | invasive, NOS | emm75.0 | 49 | 25 | 26 | 20 | 24 | 29 | 74 | 33 | 283 | 15 | 3 | 12 | 33 |
| 20154616 | USA | North America | 2015 | invasive, NOS | emm6.64 | 382 | 65 | 80 | 62 | 55 | 185 | 62 | 97 | 130 | 11 | 20 | 69 | 82 |
| 20154612 | USA | North America | 2015 | invasive, NOS | emm77.0 | 399 | 82 | 25 | 3 | 22 | 27 | 13 | 26 | 29 | 8 | 3 | 25 | 30 |
| 20154146 | USA | North America | 2015 | invasive, NOS | emm28.0 | 52 | 3 | 3 | 3 | 3 | 2 | 3 | 117 | 4 | 1 | 1 | 3 | 3 |
| 20154630 | USA | North America | 2015 | invasive, NOS | emm118.0 | 167 | 11 | 32 | 52 | 1 | 67 | 13 | 118 | 135 | 1 | 23 | 83 | 93 |
| 20154145 | USA | North America | 2015 | invasive, NOS | emm1.29 | 28 | 1 | 1 | 1 | 1 | 1 | 1 | 1 | 134 | 1 | 1 | 1 | 1 |
| 20154679 | USA | North America | 2015 | invasive, NOS | emm81.0 | 624 | 5 | 53 | 43 | 42 | 52 | 47 | 64 | 119 | 30 | 14 | 30 | 61 |
| 20154138 | USA | North America | 2015 | invasive, NOS | emm4.0 | 39 | 4 | 8 | 4 | 7 | 51 | 9 | 8 | 9 | 7 | 2 | 8 | 8 |
| 20154143 | USA | North America | 2015 | invasive, NOS | emm6.0 | 382 | 65 | 80 | 134 | 55 | 85 | 62 | 97 | 130 | 11 | 20 | 69 | 82 |
| 20154156 | USA | North America | 2015 | invasive, NOS | emm6.0 | 382 | 65 | 80 | 62 | 55 | 85 | 62 | 97 | 130 | 11 | 20 | 69 | 82 |
| 20154155 | USA | North America | 2015 | invasive, NOS | emm2.0 | 55 | 32 | 26 | 28 | 24 | 29 | 13 | 226 | 48 | 15 | 3 | 12 | 42 |
| 20154151 | USA | North America | 2015 | invasive, NOS | emm77.0 | 63 | 23 | 24 | 19 | 1 | 26 | 28 | 30 | 33 | 8 | 3 | 24 | 29 |
| 20154148 | USA | North America | 2015 | invasive, NOS | emm12.7 | 36 | 14 | 3 | 3 | 3 | 2 | 3 | 52 | 59 | 24 | 12 | 38 | 3 |
| 20154153 | USA | North America | 2015 | invasive, NOS | emm83.1 | 5 | 11 | 5 | 10 | 8 | 15 | 65 | 97 | 282 | 11 | 2 | 69 | 83 |
| 20154154 | USA | North America | 2015 | invasive, NOS | emm118.0 | 167 | 11 | 32 | 52 | 1 | 67 | 13 | 118 | 135 | 1 | 23 | 83 | 93 |
| 20154147 | USA | North America | 2015 | invasive, NOS | emm28.0 | 52 | 3 | 171 | 3 | 3 | 2 | 3 | 117 | 4 | 1 | 1 | 3 | 3 |
| 20154150 | USA | North America | 2015 | invasive, NOS | emm2.0 | 55 | 32 | 172 | 118 | 24 | 29 | 13 | 43 | 48 | 15 | 3 | 12 | 42 |
| 20154149 | USA | North America | 2015 | invasive, NOS | emm2.0 | 55 | 32 | 172 | 28 | 24 | 29 | 13 | 43 | 48 | 15 | 3 | 12 | 42 |
| 20154952 | USA | North America | 2015 | invasive, NOS | emm59.0 | 172 | 72 | 84 | 24 | 2 | 93 | 68 | 107 | 122 | 18 | 3 | 75 | 85 |
| 20154135 | USA | North America | 2015 | invasive, NOS | emm4.0 | 39 | 4 | 8 | 4 | 7 | 51 | 9 | 8 | 9 | 7 | 2 | 8 | 8 |
| 20154131 | USA | North America | 2015 | invasive, NOS | emm2.0 | 55 | 32 | 172 | 118 | 24 | 29 | 13 | 43 | 48 | 15 | 3 | 12 | 42 |
| 20154133 | USA | North America | 2015 | invasive, NOS | emm3.1 | 15 | 15 | 15 | 10 | 1 | 10 | 32 | 111 | 129 | 1 | 2 | 6 | 31 |
| 20154127 | USA | North America | 2015 | invasive, NOS | emm1.0 | 28 | 1 | 1 | 1 | 1 | 1 | 1 | 224 | 134 | 1 | 1 | 1 | 1 |
| 20154129 | USA | North America | 2015 | invasive, NOS | emm12.0 | 36 | 14 | 173 | 3 | 3 | 2 | 3 | 52 | 59 | 24 | 12 | 38 | 3 |
| 20154384 | USA | North America | 2015 | invasive, NOS | emm75.0 | 49 | 25 | 26 | 20 | 24 | 29 | 74 | 33 | 136 | 87 | 3 | 12 | 33 |
| 20154381 | USA | North America | 2015 | invasive, NOS | emm76.4 | 631 | 130 | 36 | 27 | 1 | 36 | 35 | 4 | 46 | 21 | 1 | 4 | 4 |
| 20154386 | USA | North America | 2015 | invasive, NOS | emm11.0 | ND | 1 | 6 | 40 | 61 | 20 | 20 | 13 | 139 | 8 | 3 | 18 | 4 |
| 20154796 | USA | North America | 2015 | invasive, NOS | emm28.0 | 859 | 3 | 3 | 3 | 3 | 2 | 3 | 117 | 4 | 1 | 1 | 3 | 3 |
| 20154787 | USA | North America | 2015 | invasive, NOS | emm1.0 | 858 | 1 | 1 | 1 | 1 | 1 | 1 | 1 | 134 | 1 | 1 | 1 | 1 |
| 20154017 | USA | North America | 2015 | invasive, NOS | emm89.0 | 101 | 2 | 2 | 2 | 58 | 2 | 2 | 2 | 2 | 2 | 2 | 2 | 2 |
| 20154018 | USA | North America | 2015 | invasive, NOS | emm89.0 | 101 | 2 | 2 | 2 | 2 | 2 | 2 | 2 | 2 | 2 | 2 | 2 | 2 |
| 20154019 | USA | North America | 2015 | invasive, NOS | emm82.0 | 334 | 138 | 58 | 46 | 1 | 6 | 18 | 69 | 288 | 1 | 1 | 1 | 3 |
| 20154020 | USA | North America | 2015 | invasive, NOS | emm89.0 | 101 | 2 | 2 | 2 | 2 | 2 | 2 | 2 | 2 | 2 | 2 | 2 | 2 |

| | | | | | | | | | | | | | | | | | |
|---|---|---|---|---|---|---|---|---|---|---|---|---|---|---|---|---|---|
| 20154021 | USA | North America | 2015 | invasive, NOS | emm81.0 | 624 | 5 | 53 | 43 | 42 | 52 | 47 | 64 | 119 | 30 | 14 | 30 | 173 |
| 20154023 | USA | North America | 2015 | invasive, NOS | emm82.0 | 334 | 138 | 58 | 46 | 1 | 6 | 18 | 69 | 288 | 1 | 1 | 1 | 3 |
| 20154022 | USA | North America | 2015 | invasive, NOS | emm82.0 | 334 | 138 | 58 | 46 | 1 | 6 | 18 | 69 | 288 | 1 | 1 | 1 | 3 |
| 20154605 | USA | North America | 2015 | invasive, NOS | emm114.8 | 188 | 14 | 18 | 67 | 16 | 21 | 61 | 108 | 26 | 46 | 3 | 78 | 87 |
| 20154599 | USA | North America | 2015 | invasive, NOS | emm59.0 | 172 | 72 | 84 | 24 | 2 | 93 | 68 | 107 | 122 | 18 | 3 | 75 | 85 |
| 20154600 | USA | North America | 2015 | invasive, NOS | emm118.0 | 167 | 11 | 32 | 52 | 1 | 67 | 13 | 118 | 135 | 1 | 23 | 83 | 93 |
| 20154792 | USA | North America | 2015 | invasive, NOS | emm77.0 | 63 | 23 | 24 | 19 | 1 | 26 | 28 | 30 | 33 | 8 | 3 | 24 | 29 |
| 20154607 | USA | North America | 2015 | invasive, NOS | emm6.0 | 382 | 65 | 80 | 62 | 55 | 85 | 62 | 97 | 130 | 11 | 20 | 69 | 82 |

Footnotes
ND = not determinded
^ primary disease: APSGN = acute post-streptococcal glomerul nephritis; ARF = acute rheumatic fever; NOS = not otherwise specified
### number indicates allele number of *gac* gene; allele sequences of *gac* genes can be downloaded from PubMLST; [S] indicates gene with premature stopcodon

**Supplementary Table S5 Overview of SNPs in gac gene cluster including intergenic regions in 2,021 S. pyogenes isolates**

| position | REF | ALT | annotation 1 | S/NS | gene | locus_tag | nucl_sub | AA_sub | freq (n=2021) | annotation 2 | S/NS 2 | gene | nucl_sub 2 | AA_sub 2 | freq 2 (n=2021) |
|---|---|---|---|---|---|---|---|---|---|---|---|---|---|---|---|
| 7 | G | A | A | upstream_gene_variant | intergenic | | | | 4 | | | | | | 0 |
| 12 | G | A | A | upstream_gene_variant | intergenic | | | | 7 | | | | | | 0 |
| 13 | G | A | A | upstream_gene_variant | intergenic | | | | 6 | | | | | | 0 |
| 20 | A | G | G | upstream_gene_variant | intergenic | | | | 1 | | | | | | 0 |
| 21 | T | A | A | upstream_gene_variant | intergenic | | | | 1 | | | | | | 0 |
| 23 | T | C | C | upstream_gene_variant | intergenic | | | | 1 | | | | | | 0 |
| 24 | T | C | C | upstream_gene_variant | intergenic | | | | 5 | | | | | | 0 |
| 35 | A | G | G | upstream_gene_variant | intergenic | | | | 1 | | | | | | 0 |
| 45 | A | G | G | upstream_gene_variant | intergenic | | | | 11 | | | | | | 0 |
| 47 | A | G | G | upstream_gene_variant | intergenic | | | | 2 | | | | | | 0 |
| 48 | G | A | A | upstream_gene_variant | intergenic | | | | 2 | | | | | | 0 |
| 51 | A | G | G | upstream_gene_variant | intergenic | | | | 55 | | | | | | 0 |
| 59 | T | C | C | upstream_gene_variant | intergenic | | | | 3 | | | | | | 0 |
| 69 | T | C | C | upstream_gene_variant | intergenic | | | | 5 | | | | | | 0 |
| 70 | G | A | A | upstream_gene_variant | intergenic | | | | 85 | | | | | | 0 |
| 73 | T | G | G | upstream_gene_variant | intergenic | | | | 302 | | | | | | 0 |
| 92 | G | A | A | upstream_gene_variant | intergenic | | | | 12 | | | | | | 0 |
| 93 | A | T | T | upstream_gene_variant | intergenic | | | | 4 | | | | | | 0 |
| 101 | G | A | A | upstream_gene_variant | intergenic | | | | 151 | | | | | | 0 |
| 105 | A | G | G | upstream_gene_variant | intergenic | | | | 7 | | | | | | 0 |
| 108 | G | A | A | upstream_gene_variant | intergenic | | | | 1 | | | | | | 0 |
| 111 | C | A | A | upstream_gene_variant | intergenic | | | | 14 | | | | | | 0 |
| 115 | C | T | T | upstream_gene_variant | intergenic | | | | 75 | | | | | | 0 |
| 119 | A | G | G | upstream_gene_variant | intergenic | | | | 1 | | | | | | 0 |
| 121 | G | A | A | upstream_gene_variant | intergenic | | | | 6 | | | | | | 0 |
| 122 | G | A,T | A | upstream_gene_variant | intergenic | | | | 3 | T | upstream_gene_variant | intergenic | c.-3759C>A | | 6 |
| 126 | A | G | G | upstream_gene_variant | intergenic | | | | 3 | | | | | | 0 |
| 147 | A | G | G | missense_variant | gacA | M5005_Spy0602 | c.10A>G | p.Ile4Val | 4 | | | | | | 0 |
| 158 | C | T | T | synonymous_variant | gacA | M5005_Spy0602 | c.21C>T | p.Ser7Ser | 11 | | | | | | 0 |
| 173 | G | A | A | synonymous_variant | gacA | M5005_Spy0602 | c.36G>A | p.Gly12Gly | 34 | | | | | | 0 |
| 176 | A | G | G | synonymous_variant | gacA | M5005_Spy0602 | c.39A>G | p.Thr13Thr | 20 | | | | | | 0 |
| 188 | C | T | T | synonymous_variant | gacA | M5005_Spy0602 | c.51C>T | p.Tyr17Tyr | 8 | | | | | | 0 |
| 204 | G | A | A | missense_variant | gacA | M5005_Spy0602 | c.67G>A | p.Gly23Ser | 22 | | | | | | 0 |
| 205 | G | A | A | missense_variant | gacA | M5005_Spy0602 | c.68G>A | p.Gly23Asp | 1 | | | | | | 0 |
| 211 | A | G | G | missense_variant | gacA | M5005_Spy0602 | c.74A>G | p.Asp25Gly | 19 | | | | | | 0 |
| 224 | T | G | G | synonymous_variant | gacA | M5005_Spy0602 | c.87T>G | p.Val29Val | 1 | | | | | | 0 |
| 233 | A | G | G | synonymous_variant | gacA | M5005_Spy0602 | c.96A>G | p.Ala32Ala | 8 | | | | | | 0 |
| 255 | G | A | A | missense_variant | gacA | M5005_Spy0602 | c.118G>A | p.Asp40Asn | 1 | | | | | | 0 |
| 257 | C | A,T | A | missense_variant | gacA | M5005_Spy0602 | c.120C>A | p.Asp40Glu | 4 | T | synonymous_variant | gacA | c.120C>T | p.Asp40Asp | 1 |
| 258 | A | G | G | missense_variant | gacA | M5005_Spy0602 | c.121A>G | p.Lys41Glu | 1 | | | | | | 0 |
| 267 | G | A | A | missense_variant | gacA | M5005_Spy0602 | c.130G>A | p.Ala44Thr | 11 | | | | | | 0 |
| 270 | G | A | A | missense_variant | gacA | M5005_Spy0602 | c.133G>A | p.Val45Ile | 8 | | | | | | 0 |
| 277 | C | T | T | missense_variant | gacA | M5005_Spy0602 | c.140C>T | p.Ala47Val | 5 | | | | | | 0 |
| 284 | C | T | T | synonymous_variant | gacA | M5005_Spy0602 | c.147C>T | p.Val49Val | 7 | | | | | | 0 |
| 287 | G | A | A | synonymous_variant | gacA | M5005_Spy0602 | c.150G>A | p.Lys50Lys | 3 | | | | | | 0 |
| 292 | C | G | G | missense_variant | gacA | M5005_Spy0602 | c.155C>G | p.Thr52Arg | 1 | | | | | | 0 |
| 298 | T | C | C | missense_variant | gacA | M5005_Spy0602 | c.161T>C | p.Val54Ala | 2 | | | | | | 0 |
| 310 | C | T | T | missense_variant | gacA | M5005_Spy0602 | c.173C>T | p.Ala58Val | 1 | | | | | | 0 |
| 321 | G | A | A | missense_variant | gacA | M5005_Spy0602 | c.184G>A | p.Ala62Thr | 15 | | | | | | 0 |
| 331 | C | T | T | missense_variant | gacA | M5005_Spy0602 | c.194C>T | p.Ala65Val | 12 | | | | | | 0 |
| 340 | A | G | G | missense_variant | gacA | M5005_Spy0602 | c.203A>G | p.Asp68Gly | 2 | | | | | | 0 |
| 341 | C | T | T | synonymous_variant | gacA | M5005_Spy0602 | c.204C>T | p.Asp68Asp | 15 | | | | | | 0 |
| 342 | G | A | A | missense_variant | gacA | M5005_Spy0602 | c.205G>A | p.Glu69Lys | 2 | | | | | | 0 |
| 351 | G | A | A | missense_variant | gacA | M5005_Spy0602 | c.214G>A | p.Ala72Thr | 38 | | | | | | 0 |
| 352 | C | T | T | missense_variant | gacA | M5005_Spy0602 | c.215C>T | p.Ala72Val | 13 | | | | | | 0 |
| 353 | C | T | T | synonymous_variant | gacA | M5005_Spy0602 | c.216C>T | p.Ala72Ala | 131 | | | | | | 0 |
| 364 | C | A | A | missense_variant | gacA | M5005_Spy0602 | c.227C>A | p.Ala76Asp | 4 | | | | | | 0 |
| 375 | A | G | G | missense_variant | gacA | M5005_Spy0602 | c.238A>G | p.Thr80Ala | 24 | | | | | | 0 |
| 380 | C | A | A | synonymous_variant | gacA | M5005_Spy0602 | c.243C>A | p.Gly81Gly | 45 | | | | | | 0 |
| 386 | A | G | G | synonymous_variant | gacA | M5005_Spy0602 | c.249A>G | p.Glu83Glu | 1 | | | | | | 0 |

| | | | | | | | | | | | | | | | |
|---|---|---|---|---|---|---|---|---|---|---|---|---|---|---|---|
| 399 | G | A | A | missense_variant | gacA | M5005_Spy0602 | c.262G>A | p.Ala88Thr | 4 | | | | | | 0 |
| 400 | C | T | T | missense_variant | gacA | M5005_Spy0602 | c.263C>T | p.Ala88Val | 4 | | | | | | 0 |
| 412 | A | G | G | missense_variant | gacA | M5005_Spy0602 | c.275A>G | p.Tyr92Cys | 3 | | | | | | 0 |
| 416 | G | A | A | synonymous_variant | gacA | M5005_Spy0602 | c.279G>A | p.Gly93Gly | 14 | | | | | | 0 |
| 417 | G | A | A | missense_variant | gacA | M5005_Spy0602 | c.280G>A | p.Ala94Thr | 4 | | | | | | 0 |
| 420 | A | G | G | missense_variant | gacA | M5005_Spy0602 | c.283A>G | p.Thr95Ala | 13 | | | | | | 0 |
| 422 | C | T | T | synonymous_variant | gacA | M5005_Spy0602 | c.285C>T | p.Thr95Thr | 21 | | | | | | 0 |
| 431 | C | T | T | synonymous_variant | gacA | M5005_Spy0602 | c.294C>T | p.Tyr98Tyr | 6 | | | | | | 0 |
| 432 | A | G | G | missense_variant | gacA | M5005_Spy0602 | c.295A>G | p.Ile99Val | 56 | | | | | | 0 |
| 455 | T | C | C | synonymous_variant | gacA | M5005_Spy0602 | c.318T>C | p.Asp106Asp | 17 | | | | | | 0 |
| 458 | C | T | T | synonymous_variant | gacA | M5005_Spy0602 | c.321C>T | p.Gly107Gly | 2 | | | | | | 0 |
| 468 | G | A | A | missense_variant | gacA | M5005_Spy0602 | c.331G>A | p.Val111Ile | 1 | | | | | | 0 |
| 484 | T | C | C | missense_variant | gacA | M5005_Spy0602 | c.347T>C | p.Val116Ala | 2 | | | | | | 0 |
| 488 | A | G | G | synonymous_variant | gacA | M5005_Spy0602 | c.351A>G | p.Glu117Glu | 6 | | | | | | 0 |
| 490 | C | T | T | missense_variant | gacA | M5005_Spy0602 | c.353C>T | p.Thr118Ile | 2 | | | | | | 0 |
| 495 | C | T | T | missense_variant | gacA | M5005_Spy0602 | c.358C>T | p.His120Tyr | 131 | | | | | | 0 |
| 496 | A | G | G | missense_variant | gacA | M5005_Spy0602 | c.359A>G | p.His120Arg | 1838 | | | | | | 0 |
| 503 | C | T | T | synonymous_variant | gacA | M5005_Spy0602 | c.366C>T | p.Asp122Asp | 3 | | | | | | 0 |
| 507 | A | G | G | missense_variant | gacA | M5005_Spy0602 | c.370A>G | p.Lys124Glu | 1 | | | | | | 0 |
| 508 | A | C | C | missense_variant | gacA | M5005_Spy0602 | c.371A>C | p.Lys124Thr | 1 | | | | | | 0 |
| 521 | C | T | T | synonymous_variant | gacA | M5005_Spy0602 | c.384C>T | p.Gly128Gly | 1 | | | | | | 0 |
| 524 | T | C | C | synonymous_variant | gacA | M5005_Spy0602 | c.387T>C | p.Arg129Arg | 3 | | | | | | 0 |
| 532 | G | A | A | missense_variant | gacA | M5005_Spy0602 | c.395G>A | p.Arg132His | 1 | | | | | | 0 |
| 542 | A | G | G | synonymous_variant | gacA | M5005_Spy0602 | c.405A>G | p.Glu135Glu | 40 | | | | | | 0 |
| 543 | C | T | T | synonymous_variant | gacA | M5005_Spy0602 | c.406C>T | p.Leu136Leu | 3 | | | | | | 0 |
| 555 | C | G | G | missense_variant | gacA | M5005_Spy0602 | c.418C>G | p.Arg140Gly | 6 | | | | | | 0 |
| 560 | C | T | T | synonymous_variant | gacA | M5005_Spy0602 | c.423C>T | p.Tyr141Tyr | 64 | | | | | | 0 |
| 566 | G | A | A | synonymous_variant | gacA | M5005_Spy0602 | c.429G>A | p.Glu143Glu | 4 | | | | | | 0 |
| 567 | C | T | T | missense_variant | gacA | M5005_Spy0602 | c.430C>T | p.His144Tyr | 30 | | | | | | 0 |
| 577 | T | G | G | missense_variant | gacA | M5005_Spy0602 | c.440T>G | p.Ile147Ser | 5 | | | | | | 0 |
| 581 | C | T | T | synonymous_variant | gacA | M5005_Spy0602 | c.444C>T | p.Ile148Ile | 1 | | | | | | 0 |
| 602 | A | G | G | synonymous_variant | gacA | M5005_Spy0602 | c.465A>G | p.Gly155Gly | 7 | | | | | | 0 |
| 617 | C | T | T | synonymous_variant | gacA | M5005_Spy0602 | c.480C>T | p.Asn160Asn | 1 | | | | | | 0 |
| 627 | A | G | G | missense_variant | gacA | M5005_Spy0602 | c.490A>G | p.Thr164Ala | 34 | | | | | | 0 |
| 629 | C | T | T | synonymous_variant | gacA | M5005_Spy0602 | c.492C>T | p.Thr164Thr | 18 | | | | | | 0 |
| 630 | A | C | C | missense_variant | gacA | M5005_Spy0602 | c.493A>C | p.Met165Leu | 2 | | | | | | 0 |
| 632 | G | A | A | missense_variant | gacA | M5005_Spy0602 | c.495G>A | p.Met165Ile | 43 | | | | | | 0 |
| 635 | G | A | A | synonymous_variant | gacA | M5005_Spy0602 | c.498G>A | p.Glu166Glu | 610 | | | | | | 0 |
| 637 | A | G | G | missense_variant | gacA | M5005_Spy0602 | c.500A>G | p.Gln167Arg | 6 | | | | | | 0 |
| 638 | G | A | A | synonymous_variant | gacA | M5005_Spy0602 | c.501G>A | p.Gln167Gln | 1 | | | | | | 0 |
| 644 | A | G | G | synonymous_variant | gacA | M5005_Spy0602 | c.507A>G | p.Ala169Ala | 1757 | | | | | | 0 |
| 650 | T | C | C | synonymous_variant | gacA | M5005_Spy0602 | c.513T>C | p.Asn171Asn | 2 | | | | | | 0 |
| 653 | C | T | T | synonymous_variant | gacA | M5005_Spy0602 | c.516C>T | p.His172His | 3 | | | | | | 0 |
| 657 | C | T | T | missense_variant | gacA | M5005_Spy0602 | c.520C>T | p.Arg174Cys | 21 | | | | | | 0 |
| 666 | G | A | A | missense_variant | gacA | M5005_Spy0602 | c.529G>A | p.Val177Ile | 1 | | | | | | 0 |
| 674 | T | A | A | missense_variant | gacA | M5005_Spy0602 | c.537T>A | p.Asn179Lys | 1 | | | | | | 0 |
| 680 | A | G | G | synonymous_variant | gacA | M5005_Spy0602 | c.543A>G | p.Gln181Gln | 1 | | | | | | 0 |
| 681 | C | T | T | missense_variant | gacA | M5005_Spy0602 | c.544C>T | p.His182Tyr | 4 | | | | | | 0 |
| 683 | C | T | T | synonymous_variant | gacA | M5005_Spy0602 | c.546C>T | p.His182His | 6 | | | | | | 0 |
| 704 | G | A | A | synonymous_variant | gacA | M5005_Spy0602 | c.567G>A | p.Arg189Arg | 1813 | | | | | | 0 |
| 719 | C | T | T | synonymous_variant | gacA | M5005_Spy0602 | c.582C>T | p.Phe194Phe | 1 | | | | | | 0 |
| 722 | G | T | T | missense_variant | gacA | M5005_Spy0602 | c.585G>T | p.Met195Ile | 1 | | | | | | 0 |
| 723 | T | A | A | missense_variant | gacA | M5005_Spy0602 | c.586T>A | p.Cys196Ser | 2 | | | | | | 0 |
| 724 | G | A | A | missense_variant | gacA | M5005_Spy0602 | c.587G>A | p.Cys196Tyr | 6 | | | | | | 0 |
| 726 | T | C | C | missense_variant | gacA | M5005_Spy0602 | c.589T>C | p.Tyr197His | 55 | | | | | | 0 |
| 752 | T | C | C | synonymous_variant | gacA | M5005_Spy0602 | c.615T>C | p.Phe205Phe | 1 | | | | | | 0 |
| 757 | A | G | G | missense_variant | gacA | M5005_Spy0602 | c.620A>G | p.Tyr207Cys | 5 | | | | | | 0 |
| 758 | C | T | T | synonymous_variant | gacA | M5005_Spy0602 | c.621C>T | p.Tyr207Tyr | 17 | | | | | | 0 |
| 782 | G | T | T | missense_variant | gacA | M5005_Spy0602 | c.645G>T | p.Lys215Asn | 4 | | | | | | 0 |
| 791 | C | T | T | synonymous_variant | gacA | M5005_Spy0602 | c.654C>T | p.Thr218Thr | 131 | | | | | | 0 |
| 809 | C | T | T | synonymous_variant | gacA | M5005_Spy0602 | c.672C>T | p.Ala224Ala | 12 | | | | | | 0 |
| 832 | C | T | T | missense_variant | gacA | M5005_Spy0602 | c.695C>T | p.Ala232Val | 10 | | | | | | 0 |

| | | | | | | | | | | | | | | | |
|---|---|---|---|---|---|---|---|---|---|---|---|---|---|---|---|
| 834 | G | A | A | missense_variant | gacA | M5005_Spy0602 | c.697G>A | p.Val233Ile | 3 | | | | | | 0 |
| 842 | G | C | C | synonymous_variant | gacA | M5005_Spy0602 | c.705G>C | p.Val235Val | 207 | | | | | | 0 |
| 857 | A | G | G | synonymous_variant | gacA | M5005_Spy0602 | c.720A>G | p.Ser240Ser | 4 | | | | | | 0 |
| 866 | C | T | T | synonymous_variant | gacA | M5005_Spy0602 | c.729C>T | p.Phe243Phe | 28 | | | | | | 0 |
| 868 | C | T | T | missense_variant | gacA | M5005_Spy0602 | c.731C>T | p.Pro244Leu | 25 | | | | | | 0 |
| 870 | G | A | A | missense_variant | gacA | M5005_Spy0602 | c.733G>A | p.Ala245Thr | 2 | | | | | | 0 |
| 871 | C | T | T | missense_variant | gacA | M5005_Spy0602 | c.734C>T | p.Ala245Val | 1 | | | | | | 0 |
| 876 | G | A | A | missense_variant | gacA | M5005_Spy0602 | c.739G>A | p.Ala247Thr | 1 | | | | | | 0 |
| 878 | C | T | T | synonymous_variant | gacA | M5005_Spy0602 | c.741C>T | p.Ala247Ala | 4 | | | | | | 0 |
| 881 | A | T | T | missense_variant | gacA | M5005_Spy0602 | c.744A>T | p.Lys248Asn | 15 | | | | | | 0 |
| 905 | T | A | A | missense_variant | gacA | M5005_Spy0602 | c.768T>A | p.Asn256Lys | 1 | | | | | | 0 |
| 909 | G | T | T | missense_variant | gacA | M5005_Spy0602 | c.772G>T | p.Asp258Tyr | 1 | | | | | | 0 |
| 923 | A | G | G | synonymous_variant | gacA | M5005_Spy0602 | c.786A>G | p.Ala262Ala | 39 | | | | | | 0 |
| 926 | A | G | G | synonymous_variant | gacA | M5005_Spy0602 | c.789A>G | p.Thr263Thr | 4 | | | | | | 0 |
| 940 | C | T | T | missense_variant | gacA | M5005_Spy0602 | c.803C>T | p.Pro268Leu | 2 | | | | | | 0 |
| 948 | C | A | A | missense_variant | gacA | M5005_Spy0602 | c.811C>A | p.Gln271Lys | 468 | | | | | | 0 |
| 953 | A | G | G | synonymous_variant | gacA | M5005_Spy0602 | c.816A>G | p.Glu272Glu | 34 | | | | | | 0 |
| 959 | G | A | A | synonymous_variant | gacA | M5005_Spy0602 | c.822G>A | p.Leu274Leu | 68 | | | | | | 0 |
| 962 | G | A | A | synonymous_variant | gacA | M5005_Spy0602 | c.825G>A | p.Lys275Lys | 1 | | | | | | 0 |
| 963 | G | T | T | missense_variant | gacA | M5005_Spy0602 | c.826G>T | p.Ala276Ser | 14 | | | | | | 0 |
| 971 | C | T | T | synonymous_variant | gacA | M5005_Spy0602 | c.834C>T | p.Tyr278Tyr | 1 | | | | | | 0 |
| 989 | G | A | A | synonymous_variant | gacA | M5005_Spy0602 | c.852G>A | p.Lys284Lys | 4 | | | | | | 0 |
| 1002 | A | G | G | upstream_gene_variant | intergenic | | | | 1 | | | | | | 0 |
| 1013 | T | C | C | upstream_gene_variant | intergenic | | | | 84 | | | | | | 0 |
| 1016 | C | T,A | A | upstream_gene_variant | intergenic | | | | 2 | T | upstream_gene_variant | intergenic | c.-4653G>A | | 46 |
| 1017 | T | G | G | upstream_gene_variant | intergenic | | | | 46 | | | | | | 0 |
| 1018 | A | G | G | upstream_gene_variant | intergenic | | | | 2 | | | | | | 0 |
| 1020 | G | A,T | A | upstream_gene_variant | intergenic | | | | 41 | T | upstream_gene_variant | intergenic | c.-4657C>A | | 1 |
| 1021 | G | C,A | A | upstream_gene_variant | intergenic | | | | 1 | C | upstream_gene_variant | intergenic | c.-4658C>G | | 2 |
| 1022 | C | T,G | G | upstream_gene_variant | intergenic | | | | 56 | T | upstream_gene_variant | intergenic | c.-4659G>A | | 4 |
| 1023 | C | T | T | upstream_gene_variant | intergenic | | | | 4 | | | | | | 0 |
| 1025 | A | C | C | upstream_gene_variant | intergenic | | | | 1 | | | | | | 0 |
| 1037 | G | A | A | upstream_gene_variant | intergenic | | | | 16 | | | | | | 0 |
| 1039 | G | A | A | upstream_gene_variant | intergenic | | | | 1 | | | | | | 0 |
| 1044 | A | G | G | upstream_gene_variant | intergenic | | | | 2 | | | | | | 0 |
| 1054 | A | G | G | upstream_gene_variant | intergenic | | | | 76 | | | | | | 0 |
| 1069 | G | A | A | upstream_gene_variant | intergenic | | | | 370 | | | | | | 0 |
| 1076 | A | G | G | upstream_gene_variant | intergenic | | | | 27 | | | | | | 0 |
| 1078 | C | T | T | upstream_gene_variant | intergenic | | | | 5 | | | | | | 0 |
| 1080 | C | T | T | upstream_gene_variant | intergenic | | | | 1 | | | | | | 0 |
| 1082 | G | A | A | upstream_gene_variant | intergenic | | | | 25 | | | | | | 0 |
| 1087 | T | C | C | upstream_gene_variant | intergenic | | | | 1 | | | | | | 0 |
| 1094 | G | A | A | upstream_gene_variant | intergenic | | | | 13 | | | | | | 0 |
| 1097 | T | C | C | upstream_gene_variant | intergenic | | | | 5 | | | | | | 0 |
| 1106 | A | G | G | upstream_gene_variant | intergenic | | | | 2 | | | | | | 0 |
| 1117 | G | A | A | missense_variant | gacB | M5005_Spy0603 | c.7G>A | p.Asp3Asn | 4 | | | | | | 0 |
| 1118 | A | C | C | missense_variant | gacB | M5005_Spy0603 | c.8A>C | p.Asp3Ala | 8 | | | | | | 0 |
| 1144 | T | G | G | missense_variant | gacB | M5005_Spy0603 | c.34T>G | p.Leu12Val | 1 | | | | | | 0 |
| 1146 | A | G | G | synonymous_variant | gacB | M5005_Spy0603 | c.36A>G | p.Leu12Leu | 34 | | | | | | 0 |
| 1148 | C | T | T | missense_variant | gacB | M5005_Spy0603 | c.38C>T | p.Pro13Leu | 2 | | | | | | 0 |
| 1158 | C | T | T | synonymous_variant | gacB | M5005_Spy0603 | c.48C>T | p.Tyr16Tyr | 15 | | | | | | 0 |
| 1161 | T | C | C | synonymous_variant | gacB | M5005_Spy0603 | c.51T>C | p.Gly17Gly | 13 | | | | | | 0 |
| 1164 | C | T | T | synonymous_variant | gacB | M5005_Spy0603 | c.54C>T | p.Gly18Gly | 2 | | | | | | 0 |
| 1173 | C | T | T | synonymous_variant | gacB | M5005_Spy0603 | c.63C>T | p.Thr21Thr | 386 | | | | | | 0 |
| 1177 | G | A | A | missense_variant | gacB | M5005_Spy0603 | c.67G>A | p.Val23Ile | 8 | | | | | | 0 |
| 1179 | C | T | T | synonymous_variant | gacB | M5005_Spy0603 | c.69C>T | p.Val23Val | 27 | | | | | | 0 |
| 1186 | T | C | C | synonymous_variant | gacB | M5005_Spy0603 | c.76T>C | p.Leu26Leu | 7 | | | | | | 0 |
| 1191 | T | C | C | synonymous_variant | gacB | M5005_Spy0603 | c.81T>C | p.Ile27Ile | 1 | | | | | | 0 |
| 1204 | T | C | C | missense_variant | gacB | M5005_Spy0603 | c.94T>C | p.Ser32Pro | 19 | | | | | | 0 |
| 1217 | G | A | A | missense_variant | gacB | M5005_Spy0603 | c.107G>A | p.Arg36His | 21 | | | | | | 0 |
| 1224 | T | C | C | synonymous_variant | gacB | M5005_Spy0603 | c.114T>C | p.His38His | 1593 | | | | | | 0 |
| 1266 | T | C | C | synonymous_variant | gacB | M5005_Spy0603 | c.156T>C | p.Asp52Asp | 9 | | | | | | 0 |

| | | | | | | | | | | | | | | | |
|---|---|---|---|---|---|---|---|---|---|---|---|---|---|---|---|
| 1279 | G | T | T | missense_variant | gacB | M5005_Spy0603 | c.169G>T | p.Asp57Tyr | 5 | | | | | | 0 |
| 1281 | T | C | C | synonymous_variant | gacB | M5005_Spy0603 | c.171T>C | p.Asp57Asp | 4 | | | | | | 0 |
| 1284 | C | T | T | synonymous_variant | gacB | M5005_Spy0603 | c.174C>T | p.Cys58Cys | 436 | | | | | | 0 |
| 1293 | T | C | C | synonymous_variant | gacB | M5005_Spy0603 | c.183T>C | p.Leu61Leu | 4 | | | | | | 0 |
| 1299 | C | T | T | synonymous_variant | gacB | M5005_Spy0603 | c.189C>T | p.Pro63Pro | 1 | | | | | | 0 |
| 1300 | C | T | T | missense_variant | gacB | M5005_Spy0603 | c.190C>T | p.Pro64Ser | 1 | | | | | | 0 |
| 1301 | C | A | A | missense_variant | gacB | M5005_Spy0603 | c.191C>A | p.Pro64Gln | 3 | | | | | | 0 |
| 1303 | A | C | C | missense_variant | gacB | M5005_Spy0603 | c.193A>C | p.Lys65Gln | 4 | | | | | | 0 |
| 1311 | T | A | A | synonymous_variant | gacB | M5005_Spy0603 | c.201T>A | p.Gly67Gly | 433 | | | | | | 0 |
| 1312 | C | T | T | missense_variant | gacB | M5005_Spy0603 | c.202C>T | p.Pro68Ser | 6 | | | | | | 0 |
| 1327 | G | A | A | missense_variant | gacB | M5005_Spy0603 | c.217G>A | p.Ala73Thr | 3 | | | | | | 0 |
| 1328 | C | T | T | missense_variant | gacB | M5005_Spy0603 | c.218C>T | p.Ala73Val | 2 | | | | | | 0 |
| 1345 | A | G | G | missense_variant | gacB | M5005_Spy0603 | c.235A>G | p.Ile79Val | 1 | | | | | | 0 |
| 1350 | C | T | T | synonymous_variant | gacB | M5005_Spy0603 | c.240C>T | p.Thr80Thr | 13 | | | | | | 0 |
| 1354 | G | A | A | missense_variant | gacB | M5005_Spy0603 | c.244G>A | p.Ala82Thr | 1 | | | | | | 0 |
| 1356 | C | T | T | synonymous_variant | gacB | M5005_Spy0603 | c.246C>T | p.Ala82Ala | 13 | | | | | | 0 |
| 1359 | G | A | A | synonymous_variant | gacB | M5005_Spy0603 | c.249G>A | p.Leu83Leu | 2 | | | | | | 0 |
| 1367 | G | A | A | missense_variant | gacB | M5005_Spy0603 | c.257G>A | p.Ser86Asn | 3 | | | | | | 0 |
| 1373 | A | G | G | missense_variant | gacB | M5005_Spy0603 | c.263A>G | p.Gln88Arg | 4 | | | | | | 0 |
| 1375 | C | A,T | A | missense_variant | gacB | M5005_Spy0603 | c.265C>A | p.His89Asn | 83 | T | missense_variant | gacB | c.265C>T | p.His89Tyr | 2 |
| 1376 | A | G | G | missense_variant | gacB | M5005_Spy0603 | c.266A>G | p.His89Arg | 20 | | | | | | 0 |
| 1388 | A | C | C | missense_variant | gacB | M5005_Spy0603 | c.278A>C | p.Asn93Thr | 3 | | | | | | 0 |
| 1389 | C | T | T | synonymous_variant | gacB | M5005_Spy0603 | c.279C>T | p.Asn93Asn | 3 | | | | | | 0 |
| 1404 | G | A | A | synonymous_variant | gacB | M5005_Spy0603 | c.294G>A | p.Val98Val | 6 | | | | | | 0 |
| 1407 | G | A | A | synonymous_variant | gacB | M5005_Spy0603 | c.297G>A | p.Leu99Leu | 68 | | | | | | 0 |
| 1410 | C | T | T | synonymous_variant | gacB | M5005_Spy0603 | c.300C>T | p.Gly100Gly | 1 | | | | | | 0 |
| 1422 | C | A,T | A | synonymous_variant | gacB | M5005_Spy0603 | c.312C>A | p.Gly104Gly | 13 | T | synonymous_variant | gacB | c.312C>T | p.Gly104Gly | 44 |
| 1428 | T | C | C | synonymous_variant | gacB | M5005_Spy0603 | c.318T>C | p.Phe106Phe | 8 | | | | | | 0 |
| 1429 | A | G | G | missense_variant | gacB | M5005_Spy0603 | c.319A>G | p.Ile107Val | 6 | | | | | | 0 |
| 1437 | A | G | G | synonymous_variant | gacB | M5005_Spy0603 | c.327A>G | p.Pro109Pro | 45 | | | | | | 0 |
| 1443 | G | A | A | synonymous_variant | gacB | M5005_Spy0603 | c.333G>A | p.Val111Val | 3 | | | | | | 0 |
| 1447 | C | A | A | missense_variant | gacB | M5005_Spy0603 | c.337C>A | p.Gln113Lys | 18 | | | | | | 0 |
| 1450 | A | G | G | missense_variant | gacB | M5005_Spy0603 | c.340A>G | p.Ile114Val | 14 | | | | | | 0 |
| 1451 | T | C | C | missense_variant | gacB | M5005_Spy0603 | c.341T>C | p.Ile114Thr | 4 | | | | | | 0 |
| 1461 | A | G | G | synonymous_variant | gacB | M5005_Spy0603 | c.351A>G | p.Arg117Arg | 5 | | | | | | 0 |
| 1468 | C | T | T | missense_variant | gacB | M5005_Spy0603 | c.358C>T | p.Arg120Cys | 2 | | | | | | 0 |
| 1479 | C | T | T | synonymous_variant | gacB | M5005_Spy0603 | c.369C>T | p.Ile123Ile | 2 | | | | | | 0 |
| 1488 | C | T | T | synonymous_variant | gacB | M5005_Spy0603 | c.378C>T | p.Asp126Asp | 3 | | | | | | 0 |
| 1491 | G | T | T | synonymous_variant | gacB | M5005_Spy0603 | c.381G>T | p.Gly127Gly | 11 | | | | | | 0 |
| 1492 | T | C | C | synonymous_variant | gacB | M5005_Spy0603 | c.382T>C | p.Leu128Leu | 45 | | | | | | 0 |
| 1520 | G | A | A | missense_variant | gacB | M5005_Spy0603 | c.410G>A | p.Arg137Lys | 1393 | | | | | | 0 |
| 1531 | G | A | A | missense_variant | gacB | M5005_Spy0603 | c.421G>A | p.Ala141Thr | 1 | | | | | | 0 |
| 1537 | C | T | T | synonymous_variant | gacB | M5005_Spy0603 | c.427C>T | p.Leu143Leu | 1 | | | | | | 0 |
| 1539 | G | A | A | synonymous_variant | gacB | M5005_Spy0603 | c.429G>A | p.Leu143Leu | 1 | | | | | | 0 |
| 1548 | C | T | T | synonymous_variant | gacB | M5005_Spy0603 | c.438C>T | p.Ser146Ser | 8 | | | | | | 0 |
| 1566 | A | G | G | synonymous_variant | gacB | M5005_Spy0603 | c.456A>G | p.Arg152Arg | 12 | | | | | | 0 |
| 1572 | T | A,C | A | synonymous_variant | gacB | M5005_Spy0603 | c.462T>A | p.Ala154Ala | 1 | C | synonymous_variant | gacB | c.462T>C | p.Ala154Ala | 1 |
| 1575 | T | C | C | synonymous_variant | gacB | M5005_Spy0603 | c.465T>C | p.Asp155Asp | 224 | | | | | | 0 |
| 1598 | G | C | C | missense_variant | gacB | M5005_Spy0603 | c.488G>C | p.Gly163Ala | 2 | | | | | | 0 |
| 1599 | C | T | T | synonymous_variant | gacB | M5005_Spy0603 | c.489C>T | p.Gly163Gly | 3 | | | | | | 0 |
| 1617 | A | G | G | synonymous_variant | gacB | M5005_Spy0603 | c.507A>G | p.Lys169Lys | 6 | | | | | | 0 |
| 1620 | A | G | G | synonymous_variant | gacB | M5005_Spy0603 | c.510A>G | p.Gln170Gln | 1694 | | | | | | 0 |
| 1623 | T | C | C | synonymous_variant | gacB | M5005_Spy0603 | c.513T>C | p.Val171Val | 2 | | | | | | 0 |
| 1636 | A | C | C | missense_variant | gacB | M5005_Spy0603 | c.526A>C | p.Lys176Gln | 40 | | | | | | 0 |
| 1643 | G | A | A | missense_variant | gacB | M5005_Spy0603 | c.533G>A | p.Cys178Tyr | 105 | | | | | | 0 |
| 1647 | T | C | C | synonymous_variant | gacB | M5005_Spy0603 | c.537T>C | p.Phe179Phe | 97 | | | | | | 0 |
| 1663 | C | A | A | missense_variant | gacB | M5005_Spy0603 | c.553C>A | p.Gln185Lys | 1 | | | | | | 0 |
| 1665 | G | A | A | synonymous_variant | gacB | M5005_Spy0603 | c.555G>A | p.Gln185Gln | 2 | | | | | | 0 |
| 1667 | C | T | T | missense_variant | gacB | M5005_Spy0603 | c.557C>T | p.Thr186Ile | 1 | | | | | | 0 |
| 1671 | A | T | T | missense_variant | gacB | M5005_Spy0603 | c.561A>T | p.Gln187His | 2 | | | | | | 0 |
| 1677 | A | C | C | synonymous_variant | gacB | M5005_Spy0603 | c.567A>C | p.Ser189Ser | 1 | | | | | | 0 |
| 1681 | T | C | C | synonymous_variant | gacB | M5005_Spy0603 | c.571T>C | p.Leu191Leu | 232 | | | | | | 0 |

| | | | | | | | | | | | | | | | |
|---|---|---|---|---|---|---|---|---|---|---|---|---|---|---|---|
| 1683 | G | A | A | synonymous_variant | gacB | M5005_Spy0603 | c.573G>A | p.Leu191Leu | 18 | | | | | | 0 |
| 1684 | G | A | A | missense_variant | gacB | M5005_Spy0603 | c.574G>A | p.Ala192Thr | 1853 | | | | | | 0 |
| 1685 | C | T | T | missense_variant | gacB | M5005_Spy0603 | c.575C>T | p.Ala192Val | 4 | | | | | | 0 |
| 1688 | C | T | T | missense_variant | gacB | M5005_Spy0603 | c.578C>T | p.Thr193Ile | 1 | | | | | | 0 |
| 1698 | C | T | T | synonymous_variant | gacB | M5005_Spy0603 | c.588C>T | p.Ser196Ser | 1 | | | | | | 0 |
| 1701 | G | A | A | synonymous_variant | gacB | M5005_Spy0603 | c.591G>A | p.Lys197Lys | 14 | | | | | | 0 |
| 1706 | G | T | T | missense_variant | gacB | M5005_Spy0603 | c.596G>T | p.Arg199Ile | 26 | | | | | | 0 |
| 1708 | G | A | A | missense_variant | gacB | M5005_Spy0603 | c.598G>A | p.Ala200Thr | 41 | | | | | | 0 |
| 1721 | C | T | T | missense_variant | gacB | M5005_Spy0603 | c.611C>T | p.Thr204Ile | 1 | | | | | | 0 |
| 1722 | C | T | T | synonymous_variant | gacB | M5005_Spy0603 | c.612C>T | p.Thr204Thr | 39 | | | | | | 0 |
| 1734 | A | C | C | missense_variant | gacB | M5005_Spy0603 | c.624A>C | p.Arg208Ser | 753 | | | | | | 0 |
| 1741 | G | A | A | missense_variant | gacB | M5005_Spy0603 | c.631G>A | p.Asp211Asn | 3 | | | | | | 0 |
| 1746 | C | T | T | synonymous_variant | gacB | M5005_Spy0603 | c.636C>T | p.Tyr212Tyr | 5 | | | | | | 0 |
| 1779 | T | C | C | synonymous_variant | gacB | M5005_Spy0603 | c.669T>C | p.Asn223Asn | 1 | | | | | | 0 |
| 1789 | A | G | G | missense_variant | gacB | M5005_Spy0603 | c.679A>G | p.Thr227Ala | 43 | | | | | | 0 |
| 1791 | C | T | T | synonymous_variant | gacB | M5005_Spy0603 | c.681C>T | p.Thr227Thr | 1097 | | | | | | 0 |
| 1792 | G | A | A | missense_variant | gacB | M5005_Spy0603 | c.682G>A | p.Ala228Thr | 1 | | | | | | 0 |
| 1793 | C | T | T | missense_variant | gacB | M5005_Spy0603 | c.683C>T | p.Ala228Val | 4 | | | | | | 0 |
| 1797 | T | C | C | synonymous_variant | gacB | M5005_Spy0603 | c.687T>C | p.Ile229Ile | 4 | | | | | | 0 |
| 1809 | G | A | A | missense_variant | gacB | M5005_Spy0603 | c.699G>A | p.Met233Ile | 6 | | | | | | 0 |
| 1810 | G | T | T | missense_variant | gacB | M5005_Spy0603 | c.700G>T | p.Ala234Ser | 16 | | | | | | 0 |
| 1817 | C | T | T | missense_variant | gacB | M5005_Spy0603 | c.707C>T | p.Ser236Leu | 3 | | | | | | 0 |
| 1823 | A | G | G | missense_variant | gacB | M5005_Spy0603 | c.713A>G | p.Lys238Arg | 3 | | | | | | 0 |
| 1824 | G | A | A | synonymous_variant | gacB | M5005_Spy0603 | c.714G>A | p.Lys238Lys | 3 | | | | | | 0 |
| 1826 | G | A | A | missense_variant | gacB | M5005_Spy0603 | c.716G>A | p.Arg239His | 96 | | | | | | 0 |
| 1842 | T | C,A | A | synonymous_variant | gacB | M5005_Spy0603 | c.732T>A | p.Ile244Ile | 38 | C | synonymous_variant | gacB | c.732T>C | p.Ile244Ile | 3 |
| 1851 | T | C | C | synonymous_variant | gacB | M5005_Spy0603 | c.741T>C | p.His247His | 114 | | | | | | 0 |
| 1854 | A | G | G | synonymous_variant | gacB | M5005_Spy0603 | c.744A>G | p.Glu248Glu | 49 | | | | | | 0 |
| 1856 | G | A | A | missense_variant | gacB | M5005_Spy0603 | c.746G>A | p.Gly249Asp | 140 | | | | | | 0 |
| 1860 | T | C | C | synonymous_variant | gacB | M5005_Spy0603 | c.750T>C | p.Asn250Asn | 1 | | | | | | 0 |
| 1862 | C | T | T | missense_variant | gacB | M5005_Spy0603 | c.752C>T | p.Ala251Val | 7 | | | | | | 0 |
| 1884 | T | G | G | synonymous_variant | gacB | M5005_Spy0603 | c.774T>G | p.Ala258Ala | 12 | | | | | | 0 |
| 1889 | C | T | T | missense_variant | gacB | M5005_Spy0603 | c.779C>T | p.Thr260Ile | 1 | | | | | | 0 |
| 1895 | G | A | A | missense_variant | gacB | M5005_Spy0603 | c.785G>A | p.Cys262Tyr | 17 | | | | | | 0 |
| 1898 | A | T | T | missense_variant | gacB | M5005_Spy0603 | c.788A>T | p.Asp263Val | 15 | | | | | | 0 |
| 1905 | C | T | T | synonymous_variant | gacB | M5005_Spy0603 | c.795C>T | p.Asp265Asp | 3 | | | | | | 0 |
| 1906 | C | T | T | missense_variant | gacB | M5005_Spy0603 | c.796C>T | p.Pro266Ser | 86 | | | | | | 0 |
| 1928 | C | T | T | missense_variant | gacB | M5005_Spy0603 | c.818C>T | p.Thr273Ile | 1 | | | | | | 0 |
| 1938 | C | T | T | synonymous_variant | gacB | M5005_Spy0603 | c.828C>T | p.Asp276Asp | 168 | | | | | | 0 |
| 1945 | C | T | T | synonymous_variant | gacB | M5005_Spy0603 | c.835C>T | p.Leu279Leu | 153 | | | | | | 0 |
| 1968 | G | A | A | synonymous_variant | gacB | M5005_Spy0603 | c.858G>A | p.Gln286Gln | 47 | | | | | | 0 |
| 1971 | T | C | C | synonymous_variant | gacB | M5005_Spy0603 | c.861T>C | p.Ala287Ala | 11 | | | | | | 0 |
| 1977 | C | T | T | synonymous_variant | gacB | M5005_Spy0603 | c.867C>T | p.Ala289Ala | 2 | | | | | | 0 |
| 1986 | T | C | C | synonymous_variant | gacB | M5005_Spy0603 | c.876T>C | p.His292His | 1 | | | | | | 0 |
| 2020 | C | T | T | synonymous_variant | gacB | M5005_Spy0603 | c.910C>T | p.Leu304Leu | 9 | | | | | | 0 |
| 2054 | T | C | C | missense_variant | gacB | M5005_Spy0603 | c.944T>C | p.Val315Ala | 4 | | | | | | 0 |
| 2060 | G | A | A | missense_variant | gacB | M5005_Spy0603 | c.950G>A | p.Gly317Glu | 27 | | | | | | 0 |
| 2065 | G | A | A | missense_variant | gacB | M5005_Spy0603 | c.955G>A | p.Asp319Asn | 1 | | | | | | 0 |
| 2078 | C | G | G | missense_variant | gacB | M5005_Spy0603 | c.968C>G | p.Ser323Trp | 8 | | | | | | 0 |
| 2089 | T | G | G | missense_variant | gacB | M5005_Spy0603 | c.979T>G | p.Ser327Ala | 1 | | | | | | 0 |
| 2092 | G | A | A | missense_variant | gacB | M5005_Spy0603 | c.982G>A | p.Ala328Thr | 5 | | | | | | 0 |
| 2093 | C | T | T | missense_variant | gacB | M5005_Spy0603 | c.983C>T | p.Ala328Val | 1 | | | | | | 0 |
| 2100 | T | G | G | synonymous_variant | gacB | M5005_Spy0603 | c.990T>G | p.Leu330Leu | 1685 | | | | | | 0 |
| 2107 | A | T,C | C | missense_variant | gacB | M5005_Spy0603 | c.997A>C | p.Thr333Pro | 496 | T | missense_variant | gacB | c.997A>T | p.Thr333Ser | 8 |
| 2115 | A | G | G | synonymous_variant | gacB | M5005_Spy0603 | c.1005A>G | p.Gln335Gln | 1 | | | | | | 0 |
| 2118 | A | T | T | missense_variant | gacB | M5005_Spy0603 | c.1008A>T | p.Lys336Asn | 13 | | | | | | 0 |
| 2128 | G | A | A | missense_variant | gacB | M5005_Spy0603 | c.1018G>A | p.Ala340Thr | 3 | | | | | | 0 |
| 2129 | C | T | T | missense_variant | gacB | M5005_Spy0603 | c.1019C>T | p.Ala340Val | 39 | | | | | | 0 |
| 2137 | A | G | G | missense_variant | gacB | M5005_Spy0603 | c.1027A>G | p.Ile343Val | 1 | | | | | | 0 |
| 2140 | A | G | G | missense_variant | gacB | M5005_Spy0603 | c.1030A>G | p.Asn344Asp | 32 | | | | | | 0 |
| 2142 | C | T | T | synonymous_variant | gacB | M5005_Spy0603 | c.1032C>T | p.Asn344Asn | 8 | | | | | | 0 |
| 2152 | G | A | A | missense_variant | gacB | M5005_Spy0603 | c.1042G>A | p.Ala348Thr | 7 | | | | | | 0 |

| | | | | | | | | | | | | | | | |
|---|---|---|---|---|---|---|---|---|---|---|---|---|---|---|---|
| 2156 | G | T | T | missense_variant | gacB | M5005_Spy0603 | c.1046G>T | p.Gly349Val | 9 | | | | | | 0 |
| 2167 | G | T | T | missense_variant | gacB | M5005_Spy0603 | c.1057G>T | p.Asp353Tyr | 34 | | | | | | 0 |
| 2171 | A | G | G | missense_variant | gacB | M5005_Spy0603 | c.1061A>G | p.His354Arg | 1574 | | | | | | 0 |
| 2192 | C | T | T | missense_variant | gacB | M5005_Spy0603 | c.1082C>T | p.Ala361Val | 1 | | | | | | 0 |
| 2201 | A | G | G | missense_variant | gacB | M5005_Spy0603 | c.1091A>G | p.Gln364Arg | 2 | | | | | | 0 |
| 2203 | G | C | C | missense_variant | gacB | M5005_Spy0603 | c.1093G>C | p.Glu365Gln | 9 | | | | | | 0 |
| 2211 | C | T | T | synonymous_variant | gacB | M5005_Spy0603 | c.1101C>T | p.Tyr367Tyr | 34 | | | | | | 0 |
| 2214 | T | C | C | synonymous_variant | gacB | M5005_Spy0603 | c.1104T>C | p.Thr368Thr | 1 | | | | | | 0 |
| 2218 | G | A | A | missense_variant | gacB | M5005_Spy0603 | c.1108G>A | p.Glu370Lys | 13 | | | | | | 0 |
| 2219 | A | G | G | missense_variant | gacB | M5005_Spy0603 | c.1109A>G | p.Glu370Gly | 3 | | | | | | 0 |
| 2226 | T | C | C | synonymous_variant | gacB | M5005_Spy0603 | c.1116T>C | p.Ile372Ile | 13 | | | | | | 0 |
| 2233 | G | A | A | missense_variant | gacB | M5005_Spy0603 | c.1123G>A | p.Glu375Lys | 14 | | | | | | 0 |
| 2238 | C | T | T | synonymous_variant | gacB | M5005_Spy0603 | c.1128C>T | p.Tyr376Tyr | 4 | | | | | | 0 |
| 2243 | C | T | T | missense_variant | gacB | M5005_Spy0603 | c.1133C>T | p.Ala378Val | 4 | | | | | | 0 |
| 2260 | C | T | T | missense_variant | gacB_overlap | M5005_Spy0603 | c.1150C>T | p.His384Tyr | 18 | | synonymous_variant | gacC_overlap | | p.Asn2Asn | 0 |
| 2278 | C | T | T | synonymous_variant | gacC | M5005_Spy0604 | c.24C>T | p.Ser8Ser | 1 | | | | | | 0 |
| 2284 | C | T | T | synonymous_variant | gacC | M5005_Spy0604 | c.30C>T | p.Tyr10Tyr | 7 | | | | | | 0 |
| 2290 | A | G | G | synonymous_variant | gacC | M5005_Spy0604 | c.36A>G | p.Gly12Gly | 89 | | | | | | 0 |
| 2293 | A | C | C | missense_variant | gacC | M5005_Spy0604 | c.39A>C | p.Glu13Asp | 4 | | | | | | 0 |
| 2294 | C | T | T | missense_variant | gacC | M5005_Spy0604 | c.40C>T | p.Arg14Cys | 1 | | | | | | 0 |
| 2296 | C | T | T | synonymous_variant | gacC | M5005_Spy0604 | c.42C>T | p.Arg14Arg | 10 | | | | | | 0 |
| 2299 | C | T | T | synonymous_variant | gacC | M5005_Spy0604 | c.45C>T | p.Phe15Phe | 18 | | | | | | 0 |
| 2304 | C | T | T | missense_variant | gacC | M5005_Spy0604 | c.50C>T | p.Ala17Val | 14 | | | | | | 0 |
| 2312 | A | G | G | missense_variant | gacC | M5005_Spy0604 | c.58A>G | p.Ile20Val | 1 | | | | | | 0 |
| 2320 | T | C | C | synonymous_variant | gacC | M5005_Spy0604 | c.66T>C | p.Ser22Ser | 1 | | | | | | 0 |
| 2328 | G | T | T | missense_variant | gacC | M5005_Spy0604 | c.74G>T | p.Arg25Met | 50 | | | | | | 0 |
| 2329 | G | A | A | synonymous_variant | gacC | M5005_Spy0604 | c.75G>A | p.Arg25Arg | 357 | | | | | | 0 |
| 2332 | A | G | G | synonymous_variant | gacC | M5005_Spy0604 | c.78A>G | p.Gln26Gln | 41 | | | | | | 0 |
| 2335 | A | G | G | synonymous_variant | gacC | M5005_Spy0604 | c.81A>G | p.Thr27Thr | 41 | | | | | | 0 |
| 2341 | C | A | A | missense_variant | gacC | M5005_Spy0604 | c.87C>A | p.Asn29Lys | 14 | | | | | | 0 |
| 2342 | G | A,C | A | missense_variant | gacC | M5005_Spy0604 | c.88G>A | p.Asp30Asn | 9 | C | missense_variant | gacC | c.88G>C | p.Asp30His | 1 |
| 2344 | C | T | T | synonymous_variant | gacC | M5005_Spy0604 | c.90C>T | p.Asp30Asp | 22 | | | | | | 0 |
| 2350 | C | T | T | synonymous_variant | gacC | M5005_Spy0604 | c.96C>T | p.Thr32Thr | 1 | | | | | | 0 |
| 2356 | G | A | A | synonymous_variant | gacC | M5005_Spy0604 | c.102G>A | p.Leu34Leu | 4 | | | | | | 0 |
| 2359 | T | C | C | synonymous_variant | gacC | M5005_Spy0604 | c.105T>C | p.Ile35Ile | 2 | | | | | | 0 |
| 2371 | T | A,G | A | synonymous_variant | gacC | M5005_Spy0604 | c.117T>A | p.Gly39Gly | 1411 | G | synonymous_variant | gacC | c.117T>G | p.Gly39Gly | 2 |
| 2376 | C | T | T | missense_variant | gacC | M5005_Spy0604 | c.122C>T | p.Thr41Ile | 1 | | | | | | 0 |
| 2383 | C | T | T | synonymous_variant | gacC | M5005_Spy0604 | c.129C>T | p.Gly43Gly | 3 | | | | | | 0 |
| 2386 | C | T | T | synonymous_variant | gacC | M5005_Spy0604 | c.132C>T | p.Thr44Thr | 1 | | | | | | 0 |
| 2389 | G | T | T | missense_variant | gacC | M5005_Spy0604 | c.135G>T | p.Gln45His | 1 | | | | | | 0 |
| 2400 | G | A | A | missense_variant | gacC | M5005_Spy0604 | c.146G>A | p.Arg49His | 1 | | | | | | 0 |
| 2401 | C | A | A | synonymous_variant | gacC | M5005_Spy0604 | c.147C>A | p.Arg49Arg | 6 | | | | | | 0 |
| 2408 | G | A | A | missense_variant | gacC | M5005_Spy0604 | c.154G>A | p.Val52Ile | 19 | | | | | | 0 |
| 2410 | C | T | T | synonymous_variant | gacC | M5005_Spy0604 | c.156C>T | p.Val52Val | 2 | | | | | | 0 |
| 2435 | A | G | G | missense_variant | gacC | M5005_Spy0604 | c.181A>G | p.Ile61Val | 3 | | | | | | 0 |
| 2440 | T | C | C | synonymous_variant | gacC | M5005_Spy0604 | c.186T>C | p.Asn62Asn | 1 | | | | | | 0 |
| 2452 | A | G | G | synonymous_variant | gacC | M5005_Spy0604 | c.198A>G | p.Thr66Thr | 1 | | | | | | 0 |
| 2469 | T | C | C | missense_variant | gacC | M5005_Spy0604 | c.215T>C | p.Ile72Thr | 1 | | | | | | 0 |
| 2488 | C | G | G | synonymous_variant | gacC | M5005_Spy0604 | c.234C>G | p.Leu78Leu | 25 | | | | | | 0 |
| 2493 | A | G | G | missense_variant | gacC | M5005_Spy0604 | c.239A>G | p.Lys80Arg | 39 | | | | | | 0 |
| 2497 | C | T | T | synonymous_variant | gacC | M5005_Spy0604 | c.243C>T | p.His81His | 11 | | | | | | 0 |
| 2510 | G | A | A | missense_variant | gacC | M5005_Spy0604 | c.256G>A | p.Val86Ile | 46 | | | | | | 0 |
| 2527 | C | T | T | synonymous_variant | gacC | M5005_Spy0604 | c.273C>T | p.Asp91Asp | 1 | | | | | | 0 |
| 2533 | T | C | C | synonymous_variant | gacC | M5005_Spy0604 | c.279T>C | p.Asp93Asp | 1 | | | | | | 0 |
| 2537 | A | G | G | missense_variant | gacC | M5005_Spy0604 | c.283A>G | p.Ile95Val | 167 | | | | | | 0 |
| 2569 | A | G | G | synonymous_variant | gacC | M5005_Spy0604 | c.315A>G | p.Leu105Leu | 1 | | | | | | 0 |
| 2575 | G | A | A | synonymous_variant | gacC | M5005_Spy0604 | c.321G>A | p.Glu107Glu | 16 | | | | | | 0 |
| 2576 | G | A | A | missense_variant | gacC | M5005_Spy0604 | c.322G>A | p.Ala108Thr | 28 | | | | | | 0 |
| 2583 | A | G | G | missense_variant | gacC | M5005_Spy0604 | c.329A>G | p.Lys110Arg | 1 | | | | | | 0 |
| 2594 | A | G | G | missense_variant | gacC | M5005_Spy0604 | c.340A>G | p.Thr114Ala | 45 | | | | | | 0 |
| 2595 | C | A | A | missense_variant | gacC | M5005_Spy0604 | c.341C>A | p.Thr114Lys | 128 | | | | | | 0 |
| 2608 | G | A | A | synonymous_variant | gacC | M5005_Spy0604 | c.354G>A | p.Leu118Leu | 42 | | | | | | 0 |

| | | | | | | | | | | | | | | | |
|---|---|---|---|---|---|---|---|---|---|---|---|---|---|---|---|
| 2620 | C | T | T | synonymous_variant | gacC | M5005_Spy0604 | c.366C>T | p.Asp122Asp | 9 | | | | | | 0 |
| 2626 | A | G | G | synonymous_variant | gacC | M5005_Spy0604 | c.372A>G | p.Lys124Lys | 2 | | | | | | 0 |
| 2641 | T | C | C | synonymous_variant | gacC | M5005_Spy0604 | c.387T>C | p.His129His | 3 | | | | | | 0 |
| 2645 | G | A | A | missense_variant | gacC | M5005_Spy0604 | c.391G>A | p.Ala131Thr | 1 | | | | | | 0 |
| 2648 | G | A | A | missense_variant | gacC | M5005_Spy0604 | c.394G>A | p.Val132Ile | 1886 | | | | | | 0 |
| 2652 | G | A | A | missense_variant | gacC | M5005_Spy0604 | c.398G>A | p.Cys133Tyr | 5 | | | | | | 0 |
| 2654 | C | T | T | missense_variant | gacC | M5005_Spy0604 | c.400C>T | p.His134Tyr | 9 | | | | | | 0 |
| 2657 | G | A | A | missense_variant | gacC | M5005_Spy0604 | c.403G>A | p.Asp135Asn | 519 | | | | | | 0 |
| 2659 | C | T | T | synonymous_variant | gacC | M5005_Spy0604 | c.405C>T | p.Asp135Asp | 16 | | | | | | 0 |
| 2668 | T | A | A | synonymous_variant | gacC | M5005_Spy0604 | c.414T>A | p.Ile138Ile | 4 | | | | | | 0 |
| 2671 | G | A | A | synonymous_variant | gacC | M5005_Spy0604 | c.417G>A | p.Lys139Lys | 2 | | | | | | 0 |
| 2674 | C | T | T | synonymous_variant | gacC | M5005_Spy0604 | c.420C>T | p.Thr140Thr | 28 | | | | | | 0 |
| 2683 | T | C | C | synonymous_variant | gacC | M5005_Spy0604 | c.429T>C | p.Gly143Gly | 21 | | | | | | 0 |
| 2692 | C | T | T | synonymous_variant | gacC | M5005_Spy0604 | c.438C>T | p.Asn146Asn | 2 | | | | | | 0 |
| 2697 | G | C | C | missense_variant | gacC | M5005_Spy0604 | c.443G>C | p.Ser148Thr | 1 | | | | | | 0 |
| 2725 | C | T | T | synonymous_variant | gacC | M5005_Spy0604 | c.471C>T | p.Thr157Thr | 2 | | | | | | 0 |
| 2741 | A | G | G | missense_variant | gacC | M5005_Spy0604 | c.487A>G | p.Met163Val | 1 | | | | | | 0 |
| 2759 | C | T | T | synonymous_variant | gacC | M5005_Spy0604 | c.505C>T | p.Leu169Leu | 2 | | | | | | 0 |
| 2761 | G | A | A | synonymous_variant | gacC | M5005_Spy0604 | c.507G>A | p.Leu169Leu | 5 | | | | | | 0 |
| 2764 | T | C | C | synonymous_variant | gacC | M5005_Spy0604 | c.510T>C | p.Ala170Ala | 2 | | | | | | 0 |
| 2767 | G | T | T | missense_variant | gacC | M5005_Spy0604 | c.513G>T | p.Glu171Asp | 2 | | | | | | 0 |
| 2769 | A | G | G | missense_variant | gacC | M5005_Spy0604 | c.515A>G | p.Glu172Gly | 4 | | | | | | 0 |
| 2785 | C | T | T | synonymous_variant | gacC | M5005_Spy0604 | c.531C>T | p.Asp177Asp | 1 | | | | | | 0 |
| 2787 | G | A | A | missense_variant | gacC | M5005_Spy0604 | c.533G>A | p.Gly178Asp | 8 | | | | | | 0 |
| 2828 | G | A | A | missense_variant | gacC | M5005_Spy0604 | c.574G>A | p.Ala192Thr | 1 | | | | | | 0 |
| 2832 | T | C | C | missense_variant | gacC | M5005_Spy0604 | c.578T>C | p.Ile193Thr | 1938 | | | | | | 0 |
| 2833 | T | C | C | synonymous_variant | gacC | M5005_Spy0604 | c.579T>C | p.Ile193Ile | 4 | | | | | | 0 |
| 2839 | A | G | G | synonymous_variant | gacC | M5005_Spy0604 | c.585A>G | p.Lys195Lys | 2 | | | | | | 0 |
| 2858 | C | T | T | missense_variant | gacC | M5005_Spy0604 | c.604C>T | p.Pro202Ser | 1 | | | | | | 0 |
| 2860 | A | G | G | synonymous_variant | gacC | M5005_Spy0604 | c.606A>G | p.Pro202Pro | 5 | | | | | | 0 |
| 2872 | C | T | T | synonymous_variant | gacC | M5005_Spy0604 | c.618C>T | p.Tyr206Tyr | 8 | | | | | | 0 |
| 2875 | T | C | C | synonymous_variant | gacC | M5005_Spy0604 | c.621T>C | p.Arg207Arg | 3 | | | | | | 0 |
| 2878 | G | T | T | missense_variant | gacC | M5005_Spy0604 | c.624G>T | p.Gln208His | 1 | | | | | | 0 |
| 2885 | G | A | A | missense_variant | gacC | M5005_Spy0604 | c.631G>A | p.Ala211Thr | 5 | | | | | | 0 |
| 2896 | G | A | A | synonymous_variant | gacC | M5005_Spy0604 | c.642G>A | p.Leu214Leu | 36 | | | | | | 0 |
| 2923 | G | A | A | missense_variant | gacC | M5005_Spy0604 | c.669G>A | p.Met223Ile | 1 | | | | | | 0 |
| 2928 | A | G | G | missense_variant | gacC | M5005_Spy0604 | c.674A>G | p.Asn225Ser | 3 | | | | | | 0 |
| 2936 | A | C | C | missense_variant | gacC | M5005_Spy0604 | c.682A>C | p.Thr228Pro | 37 | | | | | | 0 |
| 2937 | C | A,* | A | missense_variant | gacC | M5005_Spy0604 | c.683C>A | p.Thr228Asn | 537 | | | | | | 1 |
| 2938 | C | A | A | synonymous_variant | gacC | M5005_Spy0604 | c.684C>A | p.Thr228Thr | 1 | | | | | | 0 |
| 2941 | C | T | T | synonymous_variant | gacC | M5005_Spy0604 | c.687C>T | p.Pro229Pro | 3 | | | | | | 0 |
| 2947 | T | C | C | synonymous_variant | gacC | M5005_Spy0604 | c.693T>C | p.His231His | 11 | | | | | | 0 |
| 2953 | C | A | A | synonymous_variant | gacC | M5005_Spy0604 | c.699C>A | p.Val233Val | 1 | | | | | | 0 |
| 2956 | C | A,T | A | missense_variant | gacC | M5005_Spy0604 | c.702C>A | p.Asn234Lys | 1 | T | synonymous_variant | gacC | c.702C>T | p.Asn234Asn | 1 |
| 2968 | G | T | T | missense_variant | gacC | M5005_Spy0604 | c.714G>T | p.Trp238Cys | 3 | | | | | | 0 |
| 2980 | A | G | G | synonymous_variant | gacC | M5005_Spy0604 | c.726A>G | p.Ser242Ser | 18 | | | | | | 0 |
| 3010 | T | C | C | synonymous_variant | gacC | M5005_Spy0604 | c.756T>C | p.Asp252Asp | 8 | | | | | | 0 |
| 3020 | A | G | G | missense_variant | gacC | M5005_Spy0604 | c.766A>G | p.Lys256Glu | 17 | | | | | | 0 |
| 3023 | C | G | G | missense_variant | gacC | M5005_Spy0604 | c.769C>G | p.Pro257Ala | 1 | | | | | | 0 |
| 3024 | C | T | T | missense_variant | gacC | M5005_Spy0604 | c.770C>T | p.Pro257Leu | 1 | | | | | | 0 |
| 3031 | C | T | T | synonymous_variant | gacC | M5005_Spy0604 | c.777C>T | p.Asp259Asp | 39 | | | | | | 0 |
| 3032 | C | T | T | missense_variant | gacC | M5005_Spy0604 | c.778C>T | p.His260Tyr | 1 | | | | | | 0 |
| 3040 | G | A | A | synonymous_variant | gacC | M5005_Spy0604 | c.786G>A | p.Leu262Leu | 2 | | | | | | 0 |
| 3046 | G | A | A | synonymous_variant | gacC | M5005_Spy0604 | c.792G>A | p.Thr264Thr | 16 | | | | | | 0 |
| 3047 | G | A | A | missense_variant | gacC | M5005_Spy0604 | c.793G>A | p.Ala265Thr | 1 | | | | | | 0 |
| 3048 | C | A | A | missense_variant | gacC | M5005_Spy0604 | c.794C>A | p.Ala265Asp | 4 | | | | | | 0 |
| 3057 | C | T | T | missense_variant | gacC | M5005_Spy0604 | c.803C>T | p.Ser268Phe | 9 | | | | | | 0 |
| 3071 | C | T | T | missense_variant | gacC | M5005_Spy0604 | c.817C>T | p.Pro273Ser | 9 | | | | | | 0 |
| 3073 | T | C | C | synonymous_variant | gacC | M5005_Spy0604 | c.819T>C | p.Pro273Pro | 5 | | | | | | 0 |
| 3079 | C | T | T | synonymous_variant | gacC | M5005_Spy0604 | c.825C>T | p.Thr275Thr | 28 | | | | | | 0 |
| 3081 | A | G | G | missense_variant | gacC | M5005_Spy0604 | c.827A>G | p.Lys276Arg | 8 | | | | | | 0 |
| 3090 | C | T | T | missense_variant | gacC | M5005_Spy0604 | c.836C>T | p.Ala279Val | 1 | | | | | | 0 |

| | | | | | | | | | | | | | | | |
|---|---|---|---|---|---|---|---|---|---|---|---|---|---|---|---|
| 3096 | T | C | C | missense_variant | gacC | M5005_Spy0604 | c.842T>C | p.Leu281Ser | 14 | | | | | | 0 |
| 3103 | G | A | A | synonymous_variant | gacC | M5005_Spy0604 | c.849G>A | p.Arg283Arg | 11 | | | | | | 0 |
| 3107 | G | A | A | missense_variant | gacC | M5005_Spy0604 | c.853G>A | p.Gly285Ser | 6 | | | | | | 0 |
| 3114 | G | A | A | missense_variant | gacC | M5005_Spy0604 | c.860G>A | p.Arg287Lys | 2 | | | | | | 0 |
| 3115 | A | G,C | C | missense_variant | gacC | M5005_Spy0604 | c.861A>C | p.Arg287Ser | 13 | G | synonymous_variant | gacC | c.861A>G | p.Arg287Arg | 1 |
| 3123 | G | A | A | missense_variant | gacC | M5005_Spy0604 | c.869G>A | p.Arg290His | 4 | | | | | | 0 |
| 3127 | C | T | T | synonymous_variant | gacC | M5005_Spy0604 | c.873C>T | p.Ile291Ile | 29 | | | | | | 0 |
| 3133 | C | T | T | synonymous_variant | gacC | M5005_Spy0604 | c.879C>T | p.His293His | 64 | | | | | | 0 |
| 3136 | C | A | A | synonymous_variant | gacC | M5005_Spy0604 | c.882C>A | p.Thr294Thr | 3 | | | | | | 0 |
| 3156 | T | C | C | missense_variant | gacC | M5005_Spy0604 | c.902T>C | p.Val301Ala | 1 | | | | | | 0 |
| 3164 | T | C | C | synonymous_variant | gacC | M5005_Spy0604 | c.910T>C | p.Leu304Leu | 3 | | | | | | 0 |
| 3170 | G | T | T | missense_variant | gacC | M5005_Spy0604 | c.916G>T | p.Gly306Cys | 1 | | | | | | 0 |
| 3187 | A | G | G | splice_region_variant&stop_ret | gacC | M5005_Spy0604 | c.933A>G | p.Ter311Ter | 1 | | | | | | 0 |
| 3217 | A | G | G | missense_variant | gacD | M5005_Spy0605 | c.28A>G | p.Ile10Val | 16 | | | | | | 0 |
| 3223 | C | T | T | synonymous_variant | gacD | M5005_Spy0605 | c.34C>T | p.Leu12Leu | 1 | | | | | | 0 |
| 3255 | A | G | G | synonymous_variant | gacD | M5005_Spy0605 | c.66A>G | p.Leu22Leu | 63 | | | | | | 0 |
| 3267 | G | A | A | synonymous_variant | gacD | M5005_Spy0605 | c.78G>A | p.Gly26Gly | 354 | | | | | | 0 |
| 3270 | C | T | T | synonymous_variant | gacD | M5005_Spy0605 | c.81C>T | p.Ser27Ser | 9 | | | | | | 0 |
| 3310 | A | C | C | missense_variant | gacD | M5005_Spy0605 | c.121A>C | p.Met41Leu | 1 | | | | | | 0 |
| 3315 | T | C | C | synonymous_variant | gacD | M5005_Spy0605 | c.126T>C | p.Phe42Phe | 3 | | | | | | 0 |
| 3318 | G | A | A | synonymous_variant | gacD | M5005_Spy0605 | c.129G>A | p.Thr43Thr | 4 | | | | | | 0 |
| 3330 | G | A | A | synonymous_variant | gacD | M5005_Spy0605 | c.141G>A | p.Leu47Leu | 3 | | | | | | 0 |
| 3333 | A | G | G | synonymous_variant | gacD | M5005_Spy0605 | c.144A>G | p.Val48Val | 57 | | | | | | 0 |
| 3337 | A | C | C | missense_variant | gacD | M5005_Spy0605 | c.148A>C | p.Ile50Leu | 11 | | | | | | 0 |
| 3351 | C | T | T | synonymous_variant | gacD | M5005_Spy0605 | c.162C>T | p.Arg54Arg | 3 | | | | | | 0 |
| 3352 | C | T | T | synonymous_variant | gacD | M5005_Spy0605 | c.163C>T | p.Leu55Leu | 5 | | | | | | 0 |
| 3354 | G | A | A | synonymous_variant | gacD | M5005_Spy0605 | c.165G>A | p.Leu55Leu | 1 | | | | | | 0 |
| 3359 | G | T | T | missense_variant | gacD | M5005_Spy0605 | c.170G>T | p.Gly57Val | 9 | | | | | | 0 |
| 3360 | A | G | G | synonymous_variant | gacD | M5005_Spy0605 | c.171A>G | p.Gly57Gly | 1 | | | | | | 0 |
| 3361 | A | G | G | missense_variant | gacD | M5005_Spy0605 | c.172A>G | p.Asn58Asp | 182 | | | | | | 0 |
| 3364 | G | A | A | missense_variant | gacD | M5005_Spy0605 | c.175G>A | p.Val59Ile | 23 | | | | | | 0 |
| 3367 | C | T | T | missense_variant | gacD | M5005_Spy0605 | c.178C>T | p.Pro60Ser | 1 | | | | | | 0 |
| 3378 | A | G | G | synonymous_variant | gacD | M5005_Spy0605 | c.189A>G | p.Pro63Pro | 1 | | | | | | 0 |
| 3384 | G | A | A | synonymous_variant | gacD | M5005_Spy0605 | c.195G>A | p.Ala65Ala | 1 | | | | | | 0 |
| 3392 | T | C | C | missense_variant | gacD | M5005_Spy0605 | c.203T>C | p.Leu68Ser | 1 | | | | | | 0 |
| 3393 | G | A | A | synonymous_variant | gacD | M5005_Spy0605 | c.204G>A | p.Leu68Leu | 25 | | | | | | 0 |
| 3409 | T | C | C | missense_variant | gacD | M5005_Spy0605 | c.220T>C | p.Ser74Pro | 1 | | | | | | 0 |
| 3425 | C | T | T | missense_variant | gacD | M5005_Spy0605 | c.236C>T | p.Ala79Val | 1 | | | | | | 0 |
| 3426 | A | C | C | synonymous_variant | gacD | M5005_Spy0605 | c.237A>C | p.Ala79Ala | 2 | | | | | | 0 |
| 3442 | G | A | A | missense_variant | gacD | M5005_Spy0605 | c.253G>A | p.Val85Ile | 1 | | | | | | 0 |
| 3452 | T | C | C | missense_variant | gacD | M5005_Spy0605 | c.263T>C | p.Val88Ala | 3 | | | | | | 0 |
| 3456 | T | C | C | synonymous_variant | gacD | M5005_Spy0605 | c.267T>C | p.Ser89Ser | 2 | | | | | | 0 |
| 3458 | G | A | A | missense_variant | gacD | M5005_Spy0605 | c.269G>A | p.Arg90Gln | 1 | | | | | | 0 |
| 3465 | C | T | T | synonymous_variant | gacD | M5005_Spy0605 | c.276C>T | p.Asp92Asp | 73 | | | | | | 0 |
| 3466 | T | C | C | synonymous_variant | gacD | M5005_Spy0605 | c.277T>C | p.Leu93Leu | 39 | | | | | | 0 |
| 3468 | G | A | A | synonymous_variant | gacD | M5005_Spy0605 | c.279G>A | p.Leu93Leu | 5 | | | | | | 0 |
| 3472 | C | A | A | synonymous_variant | gacD | M5005_Spy0605 | c.283C>A | p.Arg95Arg | 1 | | | | | | 0 |
| 3495 | C | T | T | synonymous_variant | gacD | M5005_Spy0605 | c.306C>T | p.His102His | 6 | | | | | | 0 |
| 3522 | A | T | T | synonymous_variant | gacD | M5005_Spy0605 | c.333A>T | p.Gly111Gly | 1 | | | | | | 0 |
| 3561 | A | G | G | synonymous_variant | gacD | M5005_Spy0605 | c.372A>G | p.Leu124Leu | 14 | | | | | | 0 |
| 3568 | G | T | T | missense_variant | gacD | M5005_Spy0605 | c.379G>T | p.Ala127Ser | 1 | | | | | | 0 |
| 3574 | A | G | G | missense_variant | gacD | M5005_Spy0605 | c.385A>G | p.Ile129Val | 13 | | | | | | 0 |
| 3594 | A | G | G | synonymous_variant | gacD | M5005_Spy0605 | c.405A>G | p.Ser135Ser | 1 | | | | | | 0 |
| 3597 | G | A | A | synonymous_variant | gacD | M5005_Spy0605 | c.408G>A | p.Gly136Gly | 134 | | | | | | 0 |
| 3599 | A | G | G | missense_variant | gacD | M5005_Spy0605 | c.410A>G | p.Tyr137Cys | 32 | | | | | | 0 |
| 3601 | G | T,A | A | missense_variant | gacD | M5005_Spy0605 | c.412G>A | p.Ala138Thr | 41 | T | missense_variant | gacD | c.412G>T | p.Ala138Ser | 1 |
| 3603 | T | C | C | synonymous_variant | gacD | M5005_Spy0605 | c.414T>C | p.Ala138Ala | 1 | | | | | | 0 |
| 3605 | A | C | C | missense_variant | gacD | M5005_Spy0605 | c.416A>C | p.Tyr139Ser | 62 | | | | | | 0 |
| 3634 | G | A | A | missense_variant | gacD | M5005_Spy0605 | c.445G>A | p.Val149Ile | 16 | | | | | | 0 |
| 3637 | G | A | A | missense_variant | gacD | M5005_Spy0605 | c.448G>A | p.Val150Ile | 24 | | | | | | 0 |
| 3656 | C | T | T | missense_variant | gacD | M5005_Spy0605 | c.467C>T | p.Ala156Val | 3 | | | | | | 0 |
| 3714 | A | G | G | synonymous_variant | gacD | M5005_Spy0605 | c.525A>G | p.Val175Val | 61 | | | | | | 0 |

| | | | | | | | | | | | | | | | |
|---|---|---|---|---|---|---|---|---|---|---|---|---|---|---|---|
| 3715 | C | T | T | synonymous_variant | gacD | M5005_Spy0605 | c.526C>T | p.Leu176Leu | 150 | | | | | | 0 |
| 3730 | A | T | T | missense_variant | gacD | M5005_Spy0605 | c.541A>T | p.Met181Leu | 18 | | | | | | 0 |
| 3731 | T | C | C | missense_variant | gacD | M5005_Spy0605 | c.542T>C | p.Met181Thr | 65 | | | | | | 0 |
| 3774 | T | C | C | synonymous_variant | gacD | M5005_Spy0605 | c.585T>C | p.Asp195Asp | 14 | | | | | | 0 |
| 3777 | C | T | T | synonymous_variant | gacD | M5005_Spy0605 | c.588C>T | p.Ser196Ser | 9 | | | | | | 0 |
| 3778 | C | T | T | missense_variant | gacD | M5005_Spy0605 | c.589C>T | p.His197Tyr | 1 | | | | | | 0 |
| 3792 | A | G,C | C | synonymous_variant | gacD | M5005_Spy0605 | c.603A>C | p.Ala201Ala | 5 | G | synonymous_variant | gacD | c.603A>G | p.Ala201Ala | 1 |
| 3795 | G | A | A | synonymous_variant | gacD | M5005_Spy0605 | c.606G>A | p.Lys202Lys | 34 | | | | | | 0 |
| 3858 | G | A | A | synonymous_variant | gacD | M5005_Spy0605 | c.669G>A | p.Arg223Arg | 972 | | | | | | 0 |
| 3870 | G | A | A | synonymous_variant | gacD | M5005_Spy0605 | c.681G>A | p.Thr227Thr | 23 | | | | | | 0 |
| 3879 | G | A | A | synonymous_variant | gacD | M5005_Spy0605 | c.690G>A | p.Gln230Gln | 3 | | | | | | 0 |
| 3882 | G | A | A | missense_variant | gacD | M5005_Spy0605 | c.693G>A | p.Met231Ile | 8 | | | | | | 0 |
| 3888 | C | T | T | synonymous_variant | gacD | M5005_Spy0605 | c.699C>T | p.Thr233Thr | 215 | | | | | | 0 |
| 3900 | C | T | T | synonymous_variant | gacD | M5005_Spy0605 | c.711C>T | p.Tyr237Tyr | 3 | | | | | | 0 |
| 3901 | A | G | G | missense_variant | gacD | M5005_Spy0605 | c.712A>G | p.Ile238Val | 155 | | | | | | 0 |
| 3918 | A | G | G | synonymous_variant | gacD | M5005_Spy0605 | c.729A>G | p.Leu243Leu | 55 | | | | | | 0 |
| 3920 | T | C | C | missense_variant | gacD | M5005_Spy0605 | c.731T>C | p.Val244Ala | 31 | | | | | | 0 |
| 3921 | A | G | G | synonymous_variant | gacD | M5005_Spy0605 | c.732A>G | p.Val244Val | 1 | | | | | | 0 |
| 3943 | G | A | A | missense_variant | gacD | M5005_Spy0605 | c.754G>A | p.Gly252Ser | 17 | | | | | | 0 |
| 3945 | C | T | T | synonymous_variant | gacD | M5005_Spy0605 | c.756C>T | p.Gly252Gly | 2 | | | | | | 0 |
| 3952 | G | T | T | missense_variant | gacD | M5005_Spy0605 | c.763G>T | p.Val255Phe | 38 | | | | | | 0 |
| 3954 | C | T | T | synonymous_variant | gacD | M5005_Spy0605 | c.765C>T | p.Val255Val | 9 | | | | | | 0 |
| 3958 | A | G | G | missense_variant | gacD | M5005_Spy0605 | c.769A>G | p.Lys257Glu | 1 | | | | | | 0 |
| 3959 | A | G | G | missense_variant | gacD | M5005_Spy0605 | c.770A>G | p.Lys257Arg | 2 | | | | | | 0 |
| 3962 | A | G | G | missense_variant | gacD | M5005_Spy0605 | c.773A>G | p.Lys258Arg | 5 | | | | | | 0 |
| 3966 | T | C | C | synonymous_variant | gacD | M5005_Spy0605 | c.777T>C | p.Asn259Asn | 3 | | | | | | 0 |
| 3990 | T | C | C | synonymous_variant | gacD | M5005_Spy0605 | c.801T>C | p.Ile267Ile | 3 | | | | | | 0 |
| 3996 | A | G | G | missense_variant | gacE | M5005_Spy0606 | c.4A>G | p.Ile2Val | 1 | | | | | | 0 |
| 3997 | T | C | C | missense_variant | gacE | M5005_Spy0606 | c.5T>C | p.Ile2Thr | 41 | | | | | | 0 |
| 4012 | C | T | T | missense_variant | gacE | M5005_Spy0606 | c.20C>T | p.Ala7Val | 3 | | | | | | 0 |
| 4022 | A | C | C | synonymous_variant | gacE | M5005_Spy0606 | c.30A>C | p.Val10Val | 40 | | | | | | 0 |
| 4025 | G | T | T | missense_variant | gacE | M5005_Spy0606 | c.33G>T | p.Gln11His | 5 | | | | | | 0 |
| 4026 | C | T | T | missense_variant | gacE | M5005_Spy0606 | c.34C>T | p.His12Tyr | 1 | | | | | | 0 |
| 4039 | C | T | T | missense_variant | gacE | M5005_Spy0606 | c.47C>T | p.Thr16Met | 1 | | | | | | 0 |
| 4040 | G | A | A | synonymous_variant | gacE | M5005_Spy0606 | c.48G>A | p.Thr16Thr | 1603 | | | | | | 0 |
| 4049 | G | T | T | synonymous_variant | gacE | M5005_Spy0606 | c.57G>T | p.Leu19Leu | 1 | | | | | | 0 |
| 4060 | C | T | T | missense_variant | gacE | M5005_Spy0606 | c.68C>T | p.Ala23Val | 18 | | | | | | 0 |
| 4076 | C | T | T | synonymous_variant | gacE | M5005_Spy0606 | c.84C>T | p.Arg28Arg | 11 | | | | | | 0 |
| 4078 | C | T | T | missense_variant | gacE | M5005_Spy0606 | c.86C>T | p.Thr29Met | 1 | | | | | | 0 |
| 4092 | C | T | T | missense_variant | gacE | M5005_Spy0606 | c.100C>T | p.Arg34Cys | 1 | | | | | | 0 |
| 4094 | C | T | T | synonymous_variant | gacE | M5005_Spy0606 | c.102C>T | p.Arg34Arg | 12 | | | | | | 0 |
| 4097 | A | G | G | synonymous_variant | gacE | M5005_Spy0606 | c.105A>G | p.Leu35Leu | 45 | | | | | | 0 |
| 4106 | T | C | C | synonymous_variant | gacE | M5005_Spy0606 | c.114T>C | p.Ile38Ile | 1 | | | | | | 0 |
| 4112 | A | C | C | synonymous_variant | gacE | M5005_Spy0606 | c.120A>C | p.Gly40Gly | 1 | | | | | | 0 |
| 4131 | C | T | T | synonymous_variant | gacE | M5005_Spy0606 | c.139C>T | p.Leu47Leu | 9 | | | | | | 0 |
| 4156 | A | G | G | missense_variant | gacE | M5005_Spy0606 | c.164A>G | p.Tyr55Cys | 1 | | | | | | 0 |
| 4160 | A | G | G | synonymous_variant | gacE | M5005_Spy0606 | c.168A>G | p.Lys56Lys | 19 | | | | | | 0 |
| 4163 | C | T | T | synonymous_variant | gacE | M5005_Spy0606 | c.171C>T | p.Gly57Gly | 26 | | | | | | 0 |
| 4175 | A | G | G | synonymous_variant | gacE | M5005_Spy0606 | c.183A>G | p.Gly61Gly | 495 | | | | | | 0 |
| 4176 | A | G | G | missense_variant | gacE | M5005_Spy0606 | c.184A>G | p.Ile62Val | 1 | | | | | | 0 |
| 4190 | C | T | T | synonymous_variant | gacE | M5005_Spy0606 | c.198C>T | p.Asn66Asn | 77 | | | | | | 0 |
| 4196 | A | G | G | synonymous_variant | gacE | M5005_Spy0606 | c.204A>G | p.Ser68Ser | 1481 | | | | | | 0 |
| 4199 | A | G | G | synonymous_variant | gacE | M5005_Spy0606 | c.207A>G | p.Gly69Gly | 2 | | | | | | 0 |
| 4202 | A | T | T | missense_variant | gacE | M5005_Spy0606 | c.210A>T | p.Lys70Asn | 1 | | | | | | 0 |
| 4203 | T | A,C | A | missense_variant | gacE | M5005_Spy0606 | c.211T>A | p.Ser71Thr | 1 | C | missense_variant | gacE | c.211T>C | p.Ser71Pro | 1 |
| 4204 | C | A | A | stop_gained | gacE | M5005_Spy0606 | c.212C>A | p.Ser71* | 1 | | | | | | 0 |
| 4205 | A | G | G | synonymous_variant | gacE | M5005_Spy0606 | c.213A>G | p.Ser71Ser | 4 | | | | | | 0 |
| 4207 | C | A | A | missense_variant | gacE | M5005_Spy0606 | c.215C>A | p.Thr72Lys | 2 | | | | | | 0 |
| 4208 | G | A | A | synonymous_variant | gacE | M5005_Spy0606 | c.216G>A | p.Thr72Thr | 39 | | | | | | 0 |
| 4210 | T | A | A | missense_variant | gacE | M5005_Spy0606 | c.218T>A | p.Leu73His | 1 | | | | | | 0 |
| 4226 | C | T | T | synonymous_variant | gacE | M5005_Spy0606 | c.234C>T | p.Ser78Ser | 57 | | | | | | 0 |
| 4228 | A | C | C | missense_variant | gacE | M5005_Spy0606 | c.236A>C | p.Gln79Pro | 1 | | | | | | 0 |

| | | | | | | | | | | | | | | | |
|---|---|---|---|---|---|---|---|---|---|---|---|---|---|---|---|
| 4229 | G | A | A | synonymous_variant | gacE | M5005_Spy0606 | c.237G>A | p.Gln79Gln | 11 | | | | | | 0 |
| 4236 | G | A | A | missense_variant | gacE | M5005_Spy0606 | c.244G>A | p.Val82Ile | 114 | | | | | | 0 |
| 4250 | G | A,T | A | synonymous_variant | gacE | M5005_Spy0606 | c.258G>A | p.Gly86Gly | 945 | T | synonymous_variant | gacE | c.258G>T | p.Gly86Gly | 2 |
| 4258 | C | T | T | missense_variant | gacE | M5005_Spy0606 | c.266C>T | p.Thr89Ile | 19 | | | | | | 0 |
| 4268 | G | A | A | synonymous_variant | gacE | M5005_Spy0606 | c.276G>A | p.Gly92Gly | 93 | | | | | | 0 |
| 4274 | G | A | A | missense_variant | gacE | M5005_Spy0606 | c.282G>A | p.Met94Ile | 1 | | | | | | 0 |
| 4275 | G | A | A | missense_variant | gacE | M5005_Spy0606 | c.283G>A | p.Val95Ile | 1 | | | | | | 0 |
| 4276 | T | A | A | missense_variant | gacE | M5005_Spy0606 | c.284T>A | p.Val95Asp | 1 | | | | | | 0 |
| 4277 | T | C | C | synonymous_variant | gacE | M5005_Spy0606 | c.285T>C | p.Val95Val | 1 | | | | | | 0 |
| 4298 | G | A | A | synonymous_variant | gacE | M5005_Spy0606 | c.306G>A | p.Val102Val | 1 | | | | | | 0 |
| 4331 | C | T | T | synonymous_variant | gacE | M5005_Spy0606 | c.339C>T | p.Asn113Asn | 31 | | | | | | 0 |
| 4355 | G | A | A | synonymous_variant | gacE | M5005_Spy0606 | c.363G>A | p.Leu121Leu | 37 | | | | | | 0 |
| 4369 | A | G | G | missense_variant | gacE | M5005_Spy0606 | c.377A>G | p.Asp126Gly | 1 | | | | | | 0 |
| 4370 | C | T | T | synonymous_variant | gacE | M5005_Spy0606 | c.378C>T | p.Asp126Asp | 13 | | | | | | 0 |
| 4381 | A | C | C | missense_variant | gacE | M5005_Spy0606 | c.389A>C | p.Asp130Ala | 3 | | | | | | 0 |
| 4388 | C | T | T | synonymous_variant | gacE | M5005_Spy0606 | c.396C>T | p.Tyr132Tyr | 18 | | | | | | 0 |
| 4415 | G | A | A | synonymous_variant | gacE | M5005_Spy0606 | c.423G>A | p.Leu141Leu | 1 | | | | | | 0 |
| 4416 | C | A | A | missense_variant | gacE | M5005_Spy0606 | c.424C>A | p.His142Asn | 3 | | | | | | 0 |
| 4418 | T | C | C | synonymous_variant | gacE | M5005_Spy0606 | c.426T>C | p.His142His | 2 | | | | | | 0 |
| 4457 | C | T,G | G | synonymous_variant | gacE | M5005_Spy0606 | c.465C>G | p.Gly155Gly | 15 | T | synonymous_variant | gacE | c.465C>T | p.Gly155Gly | 2 |
| 4490 | T | C | C | synonymous_variant | gacE | M5005_Spy0606 | c.498T>C | p.Ile166Ile | 9 | | | | | | 0 |
| 4495 | C | T | T | missense_variant | gacE | M5005_Spy0606 | c.503C>T | p.Ala168Val | 4 | | | | | | 0 |
| 4496 | C | T | T | synonymous_variant | gacE | M5005_Spy0606 | c.504C>T | p.Ala168Ala | 4 | | | | | | 0 |
| 4499 | A | G | G | synonymous_variant | gacE | M5005_Spy0606 | c.507A>G | p.Gln169Gln | 1 | | | | | | 0 |
| 4505 | C | T | T | synonymous_variant | gacE | M5005_Spy0606 | c.513C>T | p.Asp171Asp | 6 | | | | | | 0 |
| 4526 | G | A | A | synonymous_variant | gacE | M5005_Spy0606 | c.534G>A | p.Val178Val | 12 | | | | | | 0 |
| 4571 | T | C | C | synonymous_variant | gacE | M5005_Spy0606 | c.579T>C | p.Tyr193Tyr | 1 | | | | | | 0 |
| 4576 | T | C | C | missense_variant | gacE | M5005_Spy0606 | c.584T>C | p.Met195Thr | 2 | | | | | | 0 |
| 4579 | A | G | G | missense_variant | gacE | M5005_Spy0606 | c.587A>G | p.Glu196Gly | 4 | | | | | | 0 |
| 4587 | G | T | T | missense_variant | gacE | M5005_Spy0606 | c.595G>T | p.Asp199Tyr | 3 | | | | | | 0 |
| 4643 | T | C | C | synonymous_variant | gacE | M5005_Spy0606 | c.651T>C | p.Tyr217Tyr | 15 | | | | | | 0 |
| 4655 | A | G | G | synonymous_variant | gacE | M5005_Spy0606 | c.663A>G | p.Ala221Ala | 45 | | | | | | 0 |
| 4659 | C | T | T | missense_variant | gacE | M5005_Spy0606 | c.667C>T | p.Leu223Phe | 1 | | | | | | 0 |
| 4673 | T | C | C | synonymous_variant | gacE | M5005_Spy0606 | c.681T>C | p.Gly227Gly | 3 | | | | | | 0 |
| 4692 | G | A | A | missense_variant | gacE | M5005_Spy0606 | c.700G>A | p.Glu234Lys | 1 | | | | | | 0 |
| 4700 | T | G | G | missense_variant | gacE | M5005_Spy0606 | c.708T>G | p.Phe236Leu | 1 | | | | | | 0 |
| 4702 | A | G | G | missense_variant | gacE | M5005_Spy0606 | c.710A>G | p.Asp237Gly | 2 | | | | | | 0 |
| 4703 | T | C | C | synonymous_variant | gacE | M5005_Spy0606 | c.711T>C | p.Asp237Asp | 31 | | | | | | 0 |
| 4721 | T | C | C | synonymous_variant | gacE | M5005_Spy0606 | c.729T>C | p.Ser243Ser | 7 | | | | | | 0 |
| 4722 | G | A | A | missense_variant | gacE | M5005_Spy0606 | c.730G>A | p.Val244Ile | 1 | | | | | | 0 |
| 4741 | C | T | T | missense_variant | gacE | M5005_Spy0606 | c.749C>T | p.Ala250Val | 2 | | | | | | 0 |
| 4746 | G | C | C | missense_variant | gacE | M5005_Spy0606 | c.754G>C | p.Asp252His | 1 | | | | | | 0 |
| 4749 | G | A | A | missense_variant | gacE | M5005_Spy0606 | c.757G>A | p.Ala253Thr | 1 | | | | | | 0 |
| 4753 | T | C | C | missense_variant | gacE | M5005_Spy0606 | c.761T>C | p.Met254Thr | 6 | | | | | | 0 |
| 4758 | G | A | A | missense_variant | gacE | M5005_Spy0606 | c.766G>A | p.Ala256Thr | 62 | | | | | | 0 |
| 4771 | C | T | T | missense_variant | gacE | M5005_Spy0606 | c.779C>T | p.Ser260Phe | 2 | | | | | | 0 |
| 4786 | C | T | T | missense_variant | gacE | M5005_Spy0606 | c.794C>T | p.Ala265Val | 1 | | | | | | 0 |
| 4802 | G | A | A | synonymous_variant | gacE | M5005_Spy0606 | c.810G>A | p.Val270Val | 49 | | | | | | 0 |
| 4811 | C | T | T | synonymous_variant | gacE | M5005_Spy0606 | c.819C>T | p.Ile273Ile | 8 | | | | | | 0 |
| 4817 | T | C | C | synonymous_variant | gacE | M5005_Spy0606 | c.825T>C | p.Asn275Asn | 5 | | | | | | 0 |
| 4818 | C | T | T | missense_variant | gacE | M5005_Spy0606 | c.826C>T | p.Pro276Ser | 2 | | | | | | 0 |
| 4819 | C | T | T | missense_variant | gacE | M5005_Spy0606 | c.827C>T | p.Pro276Leu | 2 | | | | | | 0 |
| 4828 | C | T | T | missense_variant | gacE | M5005_Spy0606 | c.836C>T | p.Thr279Ile | 10 | | | | | | 0 |
| 4855 | T | C | C | missense_variant | gacE | M5005_Spy0606 | c.863T>C | p.Val288Ala | 1 | | | | | | 0 |
| 4862 | T | C | C | synonymous_variant | gacE | M5005_Spy0606 | c.870T>C | p.Tyr290Tyr | 1 | | | | | | 0 |
| 4865 | A | G | G | synonymous_variant | gacE | M5005_Spy0606 | c.873A>G | p.Glu291Glu | 7 | | | | | | 0 |
| 4869 | T | C | C | synonymous_variant | gacE | M5005_Spy0606 | c.877T>C | p.Leu293Leu | 11 | | | | | | 0 |
| 4878 | G | A | A | missense_variant | gacE | M5005_Spy0606 | c.886G>A | p.Asp296Asn | 24 | | | | | | 0 |
| 4886 | C | T | T | synonymous_variant | gacE | M5005_Spy0606 | c.894C>T | p.Thr298Thr | 4 | | | | | | 0 |
| 4893 | G | A | A | missense_variant | gacE | M5005_Spy0606 | c.901G>A | p.Ala301Thr | 1 | | | | | | 0 |
| 4895 | C | T,A | A | synonymous_variant | gacE | M5005_Spy0606 | c.903C>A | p.Ala301Ala | 13 | T | synonymous_variant | gacE | c.903C>T | p.Ala301Ala | 1 |
| 4925 | T | C | C | synonymous_variant | gacE | M5005_Spy0606 | c.933T>C | p.Ile311Ile | 1 | | | | | | 0 |

| | | | | | | | | | | | | | | | |
|---|---|---|---|---|---|---|---|---|---|---|---|---|---|---|---|
| 4934 | C | T | T | synonymous_variant | gacE | M5005_Spy0606 | c.942C>T | p.Tyr314Tyr | 6 | | | | | | 0 |
| 4937 | C | T | T | synonymous_variant | gacE | M5005_Spy0606 | c.945C>T | p.Asn315Asn | 1 | | | | | | 0 |
| 4940 | C | T | T | synonymous_variant | gacE | M5005_Spy0606 | c.948C>T | p.Asp316Asp | 4 | | | | | | 0 |
| 4967 | T | C | C | synonymous_variant | gacE | M5005_Spy0606 | c.975T>C | p.Gly325Gly | 1 | | | | | | 0 |
| 4973 | T | G | G | synonymous_variant | gacE | M5005_Spy0606 | c.981T>G | p.Gly327Gly | 762 | | | | | | 0 |
| 4981 | G | A | A | missense_variant | gacE | M5005_Spy0606 | c.989G>A | p.Cys330Tyr | 1 | | | | | | 0 |
| 4987 | C | T | T | missense_variant | gacE | M5005_Spy0606 | c.995C>T | p.Ser332Phe | 1 | | | | | | 0 |
| 4988 | C | T | T | synonymous_variant | gacE | M5005_Spy0606 | c.996C>T | p.Ser332Ser | 263 | | | | | | 0 |
| 4991 | C | T | T | synonymous_variant | gacE | M5005_Spy0606 | c.999C>T | p.Tyr333Tyr | 1 | | | | | | 0 |
| 5000 | A | G | G | synonymous_variant | gacE | M5005_Spy0606 | c.1008A>G | p.Gln336Gln | 1 | | | | | | 0 |
| 5023 | A | C | C | missense_variant | gacE | M5005_Spy0606 | c.1031A>C | p.Lys344Thr | 1 | | | | | | 0 |
| 5025 | C | T | T | synonymous_variant | gacE | M5005_Spy0606 | c.1033C>T | p.Leu345Leu | 12 | | | | | | 0 |
| 5045 | C | T | T | synonymous_variant | gacE | M5005_Spy0606 | c.1053C>T | p.Val351Val | 7 | | | | | | 0 |
| 5057 | C | T | T | synonymous_variant | gacE | M5005_Spy0606 | c.1065C>T | p.Asn355Asn | 2 | | | | | | 0 |
| 5067 | C | T | T | synonymous_variant | gacE | M5005_Spy0606 | c.1075C>T | p.Leu359Leu | 2 | | | | | | 0 |
| 5072 | C | A | A | synonymous_variant | gacE | M5005_Spy0606 | c.1080C>A | p.Leu360Leu | 3 | | | | | | 0 |
| 5077 | C | T | T | missense_variant | gacE | M5005_Spy0606 | c.1085C>T | p.Ser362Leu | 3 | | | | | | 0 |
| 5080 | C | A | A | missense_variant | gacE | M5005_Spy0606 | c.1088C>A | p.Thr363Lys | 4 | | | | | | 0 |
| 5086 | A | G | G | missense_variant | gacE | M5005_Spy0606 | c.1094A>G | p.Glu365Gly | 1 | | | | | | 0 |
| 5094 | C | T | T | missense_variant | gacE | M5005_Spy0606 | c.1102C>T | p.Pro368Ser | 2 | | | | | | 0 |
| 5116 | A | G | G | missense_variant | gacE | M5005_Spy0606 | c.1124A>G | p.Asn375Ser | 38 | | | | | | 0 |
| 5122 | T | C | C | missense_variant | gacE | M5005_Spy0606 | c.1130T>C | p.Ile377Thr | 2 | | | | | | 0 |
| 5133 | G | A | A | missense_variant | gacE | M5005_Spy0606 | c.1141G>A | p.Asp381Asn | 12 | | | | | | 0 |
| 5138 | G | A | A | synonymous_variant | gacE | M5005_Spy0606 | c.1146G>A | p.Leu382Leu | 43 | | | | | | 0 |
| 5139 | T | C | C | missense_variant | gacE | M5005_Spy0606 | c.1147T>C | p.Ser383Pro | 1 | | | | | | 0 |
| 5153 | A | G | G | synonymous_variant | gacE | M5005_Spy0606 | c.1161A>G | p.Ser387Ser | 8 | | | | | | 0 |
| 5155 | C | T | T | missense_variant | gacE | M5005_Spy0606 | c.1163C>T | p.Ala388Val | 1 | | | | | | 0 |
| 5162 | C | T | T | synonymous_variant | gacE | M5005_Spy0606 | c.1170C>T | p.Gly390Gly | 220 | | | | | | 0 |
| 5166 | T | G | G | missense_variant | gacE | M5005_Spy0606 | c.1174T>G | p.Tyr392Asp | 1 | | | | | | 0 |
| 5172 | C | A | A | missense_variant | gacE | M5005_Spy0606 | c.1180C>A | p.Arg394Ser | 1 | | | | | | 0 |
| 5187 | G | T | T | missense_variant | gacE | M5005_Spy0606 | c.1195G>T | p.Val399Phe | 65 | | | | | | 0 |
| 5188 | T | A | A | missense_variant | gacE | M5005_Spy0606 | c.1196T>A | p.Val399Asp | 1 | | | | | | 0 |
| 5193 | C | T | T | missense_variant | gacE | M5005_Spy0606 | c.1201C>T | p.His401Tyr | 5 | | | | | | 0 |
| 5202 | G | C | C | upstream_gene_variant | intergenic | | | | 2 | | | | | | 0 |
| 5215 | G | A | A | upstream_gene_variant | intergenic | | | | 1 | | | | | | 0 |
| 5229 | A | C | C | missense_variant | gacF | M5005_Spy0607 | c.7A>C | p.Lys3Gln | 2 | | | | | | 0 |
| 5243 | T | C | C | synonymous_variant | gacF | M5005_Spy0607 | c.21T>C | p.Ile7Ile | 6 | | | | | | 0 |
| 5252 | T | C | C | synonymous_variant | gacF | M5005_Spy0607 | c.30T>C | p.Asn10Asn | 10 | | | | | | 0 |
| 5265 | C | A | A | missense_variant | gacF | M5005_Spy0607 | c.43C>A | p.Pro15Thr | 18 | | | | | | 0 |
| 5279 | T | G | G | missense_variant | gacF | M5005_Spy0607 | c.57T>G | p.Asp19Glu | 1 | | | | | | 0 |
| 5285 | T | C | C | synonymous_variant | gacF | M5005_Spy0607 | c.63T>C | p.Leu21Leu | 2 | | | | | | 0 |
| 5288 | C | T | T | synonymous_variant | gacF | M5005_Spy0607 | c.66C>T | p.Asp22Asp | 4 | | | | | | 0 |
| 5292 | T | C | C | missense_variant | gacF | M5005_Spy0607 | c.70T>C | p.Phe24Leu | 19 | | | | | | 0 |
| 5297 | A | T | T | missense_variant | gacF | M5005_Spy0607 | c.75A>T | p.Leu25Phe | 31 | | | | | | 0 |
| 5318 | G | A | A | synonymous_variant | gacF | M5005_Spy0607 | c.96G>A | p.Glu32Glu | 131 | | | | | | 0 |
| 5355 | G | A | A | missense_variant | gacF | M5005_Spy0607 | c.133G>A | p.Asp45Asn | 56 | | | | | | 0 |
| 5356 | A | G | G | missense_variant | gacF | M5005_Spy0607 | c.134A>G | p.Asp45Gly | 3 | | | | | | 0 |
| 5357 | C | T | T | synonymous_variant | gacF | M5005_Spy0607 | c.135C>T | p.Asp45Asp | 55 | | | | | | 0 |
| 5362 | G | T | T | missense_variant | gacF | M5005_Spy0607 | c.140G>T | p.Arg47Leu | 5 | | | | | | 0 |
| 5377 | G | C | C | missense_variant | gacF | M5005_Spy0607 | c.155G>C | p.Ser52Thr | 4 | | | | | | 0 |
| 5393 | T | C | C | synonymous_variant | gacF | M5005_Spy0607 | c.171T>C | p.Ser57Ser | 28 | | | | | | 0 |
| 5425 | C | T | T | missense_variant | gacF | M5005_Spy0607 | c.203C>T | p.Thr68Ile | 43 | | | | | | 0 |
| 5459 | A | G | G | synonymous_variant | gacF | M5005_Spy0607 | c.237A>G | p.Ala79Ala | 2 | | | | | | 0 |
| 5486 | T | C | C | synonymous_variant | gacF | M5005_Spy0607 | c.264T>C | p.Ile88Ile | 1 | | | | | | 0 |
| 5498 | T | C | C | synonymous_variant | gacF | M5005_Spy0607 | c.276T>C | p.Asp92Asp | 17 | | | | | | 0 |
| 5515 | C | T | T | missense_variant | gacF | M5005_Spy0607 | c.293C>T | p.Thr98Met | 12 | | | | | | 0 |
| 5516 | G | A | A | synonymous_variant | gacF | M5005_Spy0607 | c.294G>A | p.Thr98Thr | 39 | | | | | | 0 |
| 5537 | A | G | G | synonymous_variant | gacF | M5005_Spy0607 | c.315A>G | p.Lys105Lys | 7 | | | | | | 0 |
| 5562 | A | C | C | missense_variant | gacF | M5005_Spy0607 | c.340A>C | p.Lys114Gln | 18 | | | | | | 0 |
| 5563 | A | C | C | missense_variant | gacF | M5005_Spy0607 | c.341A>C | p.Lys114Thr | 6 | | | | | | 0 |
| 5566 | G | A | A | missense_variant | gacF | M5005_Spy0607 | c.344G>A | p.Arg115Gln | 10 | | | | | | 0 |
| 5597 | T | C | C | synonymous_variant | gacF | M5005_Spy0607 | c.375T>C | p.Phe125Phe | 1 | | | | | | 0 |

| | | | | | | | | | | | | | | | |
|---|---|---|---|---|---|---|---|---|---|---|---|---|---|---|---|
| 5598 | G | T | T | missense_variant | gacF | M5005_Spy0607 | c.376G>T | p.Val126Leu | 118 | | | | | | 0 |
| 5602 | A | C | C | missense_variant | gacF | M5005_Spy0607 | c.380A>C | p.Asn127Thr | 30 | | | | | | 0 |
| 5615 | C | T | T | synonymous_variant | gacF | M5005_Spy0607 | c.393C>T | p.Asp131Asp | 30 | | | | | | 0 |
| 5619 | T | C | C | synonymous_variant | gacF | M5005_Spy0607 | c.397T>C | p.Leu133Leu | 21 | | | | | | 0 |
| 5625 | G | A | A | missense_variant | gacF | M5005_Spy0607 | c.403G>A | p.Ala135Thr | 4 | | | | | | 0 |
| 5636 | T | C | C | synonymous_variant | gacF | M5005_Spy0607 | c.414T>C | p.Phe138Phe | 1 | | | | | | 0 |
| 5641 | C | T | T | missense_variant | gacF | M5005_Spy0607 | c.419C>T | p.Ser140Phe | 5 | | | | | | 0 |
| 5648 | T | C | C | synonymous_variant | gacF | M5005_Spy0607 | c.426T>C | p.His142His | 25 | | | | | | 0 |
| 5652 | C | T | T | missense_variant | gacF | M5005_Spy0607 | c.430C>T | p.Pro144Ser | 1 | | | | | | 0 |
| 5661 | G | A | A | missense_variant | gacF | M5005_Spy0607 | c.439G>A | p.Asp147Asn | 1 | | | | | | 0 |
| 5663 | T | C | C | synonymous_variant | gacF | M5005_Spy0607 | c.441T>C | p.Asp147Asp | 1 | | | | | | 0 |
| 5705 | A | T | T | synonymous_variant | gacF | M5005_Spy0607 | c.483A>T | p.Ala161Ala | 1 | | | | | | 0 |
| 5715 | T | C | C | synonymous_variant | gacF | M5005_Spy0607 | c.493T>C | p.Leu165Leu | 578 | | | | | | 0 |
| 5723 | T | C | C | synonymous_variant | gacF | M5005_Spy0607 | c.501T>C | p.Asp167Asp | 1 | | | | | | 0 |
| 5730 | T | C | C | synonymous_variant | gacF | M5005_Spy0607 | c.508T>C | p.Leu170Leu | 22 | | | | | | 0 |
| 5733 | A | G | G | missense_variant | gacF | M5005_Spy0607 | c.511A>G | p.Met171Val | 67 | | | | | | 0 |
| 5757 | A | G | G | missense_variant | gacF | M5005_Spy0607 | c.535A>G | p.Asn179Asp | 1 | | | | | | 0 |
| 5763 | G | A | A | missense_variant | gacF | M5005_Spy0607 | c.541G>A | p.Asp181Asn | 9 | | | | | | 0 |
| 5770 | C | T | T | missense_variant | gacF | M5005_Spy0607 | c.548C>T | p.Pro183Leu | 1 | | | | | | 0 |
| 5774 | C | T | T | synonymous_variant | gacF | M5005_Spy0607 | c.552C>T | p.Asp184Asp | 12 | | | | | | 0 |
| 5786 | T | C | C | synonymous_variant | gacF | M5005_Spy0607 | c.564T>C | p.Asp188Asp | 306 | | | | | | 0 |
| 5808 | C | T | T | synonymous_variant | gacF | M5005_Spy0607 | c.586C>T | p.Leu196Leu | 506 | | | | | | 0 |
| 5822 | C | T | T | synonymous_variant | gacF | M5005_Spy0607 | c.600C>T | p.Thr200Thr | 2 | | | | | | 0 |
| 5837 | A | C | C | synonymous_variant | gacF | M5005_Spy0607 | c.615A>C | p.Ser205Ser | 62 | | | | | | 0 |
| 5844 | A | G | G | missense_variant | gacF | M5005_Spy0607 | c.622A>G | p.Ile208Val | 1068 | | | | | | 0 |
| 5852 | C | T | T | synonymous_variant | gacF | M5005_Spy0607 | c.630C>T | p.Arg210Arg | 5 | | | | | | 0 |
| 5856 | A | G | G | missense_variant | gacF | M5005_Spy0607 | c.634A>G | p.Thr212Ala | 1 | | | | | | 0 |
| 5858 | A | G | G | synonymous_variant | gacF | M5005_Spy0607 | c.636A>G | p.Thr212Thr | 2 | | | | | | 0 |
| 5876 | A | G | G | synonymous_variant | gacF | M5005_Spy0607 | c.654A>G | p.Arg218Arg | 21 | | | | | | 0 |
| 5881 | C | T | T | missense_variant | gacF | M5005_Spy0607 | c.659C>T | p.Thr220Ile | 4 | | | | | | 0 |
| 5884 | A | G | G | missense_variant | gacF | M5005_Spy0607 | c.662A>G | p.Asp221Gly | 9 | | | | | | 0 |
| 5891 | A | C | C | synonymous_variant | gacF | M5005_Spy0607 | c.669A>C | p.Pro223Pro | 9 | | | | | | 0 |
| 5901 | T | C | C | missense_variant | gacF | M5005_Spy0607 | c.679T>C | p.Tyr227His | 1891 | | | | | | 0 |
| 5914 | A | G | G | missense_variant | gacF | M5005_Spy0607 | c.692A>G | p.Lys231Arg | 47 | | | | | | 0 |
| 5916 | C | T | T | synonymous_variant | gacF | M5005_Spy0607 | c.694C>T | p.Leu232Leu | 142 | | | | | | 0 |
| 5933 | A | T | T | missense_variant | gacF | M5005_Spy0607 | c.711A>T | p.Leu237Phe | 1 | | | | | | 0 |
| 5939 | C | T | T | synonymous_variant | gacF | M5005_Spy0607 | c.717C>T | p.Tyr239Tyr | 12 | | | | | | 0 |
| 5961 | G | A | A | missense_variant | gacF | M5005_Spy0607 | c.739G>A | p.Asp247Asn | 17 | | | | | | 0 |
| 5973 | G | A | A | missense_variant | gacF | M5005_Spy0607 | c.751G>A | p.Val251Met | 136 | | | | | | 0 |
| 5979 | G | A | A | missense_variant | gacF | M5005_Spy0607 | c.757G>A | p.Glu253Lys | 801 | | | | | | 0 |
| 5981 | A | G | G | synonymous_variant | gacF | M5005_Spy0607 | c.759A>G | p.Glu253Glu | 6 | | | | | | 0 |
| 5982 | C | T | T | synonymous_variant | gacF | M5005_Spy0607 | c.760C>T | p.Leu254Leu | 22 | | | | | | 0 |
| 5987 | A | G | G | synonymous_variant | gacF | M5005_Spy0607 | c.765A>G | p.Leu255Leu | 104 | | | | | | 0 |
| 6012 | A | G | G | missense_variant | gacF | M5005_Spy0607 | c.790A>G | p.Ile264Val | 1 | | | | | | 0 |
| 6058 | C | T | T | missense_variant | gacF | M5005_Spy0607 | c.836C>T | p.Thr279Ile | 4 | | | | | | 0 |
| 6060 | A | G | G | missense_variant | gacF | M5005_Spy0607 | c.838A>G | p.Ile280Val | 18 | | | | | | 0 |
| 6062 | C | T | T | synonymous_variant | gacF | M5005_Spy0607 | c.840C>T | p.Ile280Ile | 11 | | | | | | 0 |
| 6068 | T | C | C | synonymous_variant | gacF | M5005_Spy0607 | c.846T>C | p.Asp282Asp | 21 | | | | | | 0 |
| 6073 | C | T | T | missense_variant | gacF | M5005_Spy0607 | c.851C>T | p.Thr284Ile | 2 | | | | | | 0 |
| 6079 | A | G | G | missense_variant | gacF | M5005_Spy0607 | c.857A>G | p.Gln286Arg | 4 | | | | | | 0 |
| 6088 | C | T | T | missense_variant | gacF | M5005_Spy0607 | c.866C>T | p.Thr289Ile | 2 | | | | | | 0 |
| 6097 | T | C | C | missense_variant | gacF | M5005_Spy0607 | c.875T>C | p.Val292Ala | 2 | | | | | | 0 |
| 6098 | G | A,T | A | synonymous_variant | gacF | M5005_Spy0607 | c.876G>A | p.Val292Val | 51 | T | synonymous_variant | gacF | c.876G>T | p.Val292Val | 22 |
| 6104 | G | A | A | synonymous_variant | gacF | M5005_Spy0607 | c.882G>A | p.Leu294Leu | 40 | | | | | | 0 |
| 6155 | A | G | G | synonymous_variant | gacF | M5005_Spy0607 | c.933A>G | p.Gln311Gln | 40 | | | | | | 0 |
| 6157 | A | T | T | missense_variant | gacF | M5005_Spy0607 | c.935A>T | p.Tyr312Phe | 1 | | | | | | 0 |
| 6158 | C | T | T | synonymous_variant | gacF | M5005_Spy0607 | c.936C>T | p.Tyr312Tyr | 33 | | | | | | 0 |
| 6162 | G | A | A | missense_variant | gacF | M5005_Spy0607 | c.940G>A | p.Ala314Thr | 1 | | | | | | 0 |
| 6190 | A | G | G | missense_variant | gacF | M5005_Spy0607 | c.968A>G | p.Tyr323Cys | 20 | | | | | | 0 |
| 6191 | C | T | T | synonymous_variant | gacF | M5005_Spy0607 | c.969C>T | p.Tyr323Tyr | 1 | | | | | | 0 |
| 6198 | A | C | C | missense_variant | gacF | M5005_Spy0607 | c.976A>C | p.Lys326Gln | 2 | | | | | | 0 |
| 6254 | T | C | C | missense_variant | gacG | M5005_Spy0608 | c.28T>C | p.Phe10Leu | 1 | | | | | | 0 |

| | | | | | | | | | | | | | | | |
|---|---|---|---|---|---|---|---|---|---|---|---|---|---|---|---|
| 6278 | G | A | A | missense_variant | gacG | M5005_Spy0608 | c.52G>A | p.Ala18Thr | 5 | | | | | | 0 |
| 6291 | A | G | G | missense_variant | gacG | M5005_Spy0608 | c.65A>G | p.Tyr22Cys | 3 | | | | | | 0 |
| 6298 | G | A | A | synonymous_variant | gacG | M5005_Spy0608 | c.72G>A | p.Leu24Leu | 24 | | | | | | 0 |
| 6307 | G | T | T | missense_variant | gacG | M5005_Spy0608 | c.81G>T | p.Met27Ile | 7 | | | | | | 0 |
| 6310 | A | C | C | missense_variant | gacG | M5005_Spy0608 | c.84A>C | p.Arg28Ser | 5 | | | | | | 0 |
| 6311 | T | A | A | missense_variant | gacG | M5005_Spy0608 | c.85T>A | p.Ser29Thr | 1 | | | | | | 0 |
| 6329 | G | A | A | missense_variant | gacG | M5005_Spy0608 | c.103G>A | p.Val35Ile | 77 | | | | | | 0 |
| 6345 | G | A | A | missense_variant | gacG | M5005_Spy0608 | c.119G>A | p.Ser40Asn | 2 | | | | | | 0 |
| 6348 | A | G | G | missense_variant | gacG | M5005_Spy0608 | c.122A>G | p.Lys41Arg | 64 | | | | | | 0 |
| 6354 | G | A | A | missense_variant | gacG | M5005_Spy0608 | c.128G>A | p.Ser43Asn | 2 | | | | | | 0 |
| 6356 | C | T | T | missense_variant | gacG | M5005_Spy0608 | c.130C>T | p.His44Tyr | 5 | | | | | | 0 |
| 6358 | C | T | T | synonymous_variant | gacG | M5005_Spy0608 | c.132C>T | p.His44His | 537 | | | | | | 0 |
| 6361 | A | G | G | synonymous_variant | gacG | M5005_Spy0608 | c.135A>G | p.Glu45Glu | 2 | | | | | | 0 |
| 6362 | G | A | A | missense_variant | gacG | M5005_Spy0608 | c.136G>A | p.Asp46Asn | 18 | | | | | | 0 |
| 6372 | G | A | A | missense_variant | gacG | M5005_Spy0608 | c.146G>A | p.Arg49His | 149 | | | | | | 0 |
| 6376 | C | T | T | synonymous_variant | gacG | M5005_Spy0608 | c.150C>T | p.Leu50Leu | 1 | | | | | | 0 |
| 6382 | C | T | T | synonymous_variant | gacG | M5005_Spy0608 | c.156C>T | p.Asn52Asn | 302 | | | | | | 0 |
| 6383 | C | T | T | missense_variant | gacG | M5005_Spy0608 | c.157C>T | p.His53Tyr | 1 | | | | | | 0 |
| 6385 | C | T | T | synonymous_variant | gacG | M5005_Spy0608 | c.159C>T | p.His53His | 10 | | | | | | 0 |
| 6387 | G | T,A | A | missense_variant | gacG | M5005_Spy0608 | c.161G>A | p.Cys54Tyr | 38 | T | missense_variant | gacG | c.161G>T | p.Cys54Phe | 3 |
| 6392 | A | G | G | missense_variant | gacG | M5005_Spy0608 | c.166A>G | p.Ile56Val | 18 | | | | | | 0 |
| 6403 | T | C | C | synonymous_variant | gacG | M5005_Spy0608 | c.177T>C | p.Phe59Phe | 1 | | | | | | 0 |
| 6404 | T | C | C | synonymous_variant | gacG | M5005_Spy0608 | c.178T>C | p.Leu60Leu | 77 | | | | | | 0 |
| 6445 | C | T | T | synonymous_variant | gacG | M5005_Spy0608 | c.219C>T | p.His73His | 3 | | | | | | 0 |
| 6459 | T | C | C | missense_variant | gacG | M5005_Spy0608 | c.233T>C | p.Ile78Thr | 1938 | | | | | | 0 |
| 6461 | A | G | G | missense_variant | gacG | M5005_Spy0608 | c.235A>G | p.Met79Val | 24 | | | | | | 0 |
| 6464 | G | A | A | missense_variant | gacG | M5005_Spy0608 | c.238G>A | p.Gly80Arg | 1 | | | | | | 0 |
| 6465 | G | A | A | missense_variant | gacG | M5005_Spy0608 | c.239G>A | p.Gly80Glu | 1 | | | | | | 0 |
| 6472 | C | T | T | synonymous_variant | gacG | M5005_Spy0608 | c.246C>T | p.Asp82Asp | 2 | | | | | | 0 |
| 6475 | G | A | A | synonymous_variant | gacG | M5005_Spy0608 | c.249G>A | p.Lys83Lys | 730 | | | | | | 0 |
| 6476 | T | C | C | synonymous_variant | gacG | M5005_Spy0608 | c.250T>C | p.Leu84Leu | 4 | | | | | | 0 |
| 6484 | G | A | A | synonymous_variant | gacG | M5005_Spy0608 | c.258G>A | p.Glu86Glu | 2 | | | | | | 0 |
| 6490 | C | T | T | synonymous_variant | gacG | M5005_Spy0608 | c.264C>T | p.Asp88Asp | 6 | | | | | | 0 |
| 6492 | C | T | T | missense_variant | gacG | M5005_Spy0608 | c.266C>T | p.Ser89Leu | 3 | | | | | | 0 |
| 6493 | A | G | G | synonymous_variant | gacG | M5005_Spy0608 | c.267A>G | p.Ser89Ser | 3 | | | | | | 0 |
| 6533 | G | C | C | missense_variant | gacG | M5005_Spy0608 | c.307G>C | p.Glu103Gln | 1 | | | | | | 0 |
| 6540 | C | G | G | missense_variant | gacG | M5005_Spy0608 | c.314C>G | p.Ala105Gly | 738 | | | | | | 0 |
| 6542 | C | A | A | missense_variant | gacG | M5005_Spy0608 | c.316C>A | p.Pro106Thr | 3 | | | | | | 0 |
| 6556 | T | C | C | synonymous_variant | gacG | M5005_Spy0608 | c.330T>C | p.Asn110Asn | 30 | | | | | | 0 |
| 6569 | G | A | A | missense_variant | gacG | M5005_Spy0608 | c.343G>A | p.Glu115Lys | 6 | | | | | | 0 |
| 6571 | G | A | A | synonymous_variant | gacG | M5005_Spy0608 | c.345G>A | p.Glu115Glu | 9 | | | | | | 0 |
| 6572 | A | G | G | missense_variant | gacG | M5005_Spy0608 | c.346A>G | p.Thr116Ala | 1021 | | | | | | 0 |
| 6575 | G | A | A | missense_variant | gacG | M5005_Spy0608 | c.349G>A | p.Val117Ile | 7 | | | | | | 0 |
| 6580 | C | T | T | synonymous_variant | gacG | M5005_Spy0608 | c.354C>T | p.Asp118Asp | 39 | | | | | | 0 |
| 6603 | G | A | A | missense_variant | gacG | M5005_Spy0608 | c.377G>A | p.Arg126Lys | 6 | | | | | | 0 |
| 6605 | G | T | T | missense_variant | gacG | M5005_Spy0608 | c.379G>T | p.Gly127Cys | 1 | | | | | | 0 |
| 6607 | T | C | C | synonymous_variant | gacG | M5005_Spy0608 | c.381T>C | p.Gly127Gly | 1 | | | | | | 0 |
| 6614 | G | A | A | missense_variant | gacG | M5005_Spy0608 | c.388G>A | p.Ala130Thr | 130 | | | | | | 0 |
| 6667 | T | C | C | synonymous_variant | gacG | M5005_Spy0608 | c.441T>C | p.Ile147Ile | 1 | | | | | | 0 |
| 6713 | G | A | A | missense_variant | gacG | M5005_Spy0608 | c.487G>A | p.Glu163Lys | 14 | | | | | | 0 |
| 6724 | A | G | G | synonymous_variant | gacG | M5005_Spy0608 | c.498A>G | p.Gln166Gln | 5 | | | | | | 0 |
| 6736 | G | A | A | synonymous_variant | gacG | M5005_Spy0608 | c.510G>A | p.Gln170Gln | 885 | | | | | | 0 |
| 6738 | A | G | G | missense_variant | gacG | M5005_Spy0608 | c.512A>G | p.His171Arg | 6 | | | | | | 0 |
| 6742 | T | C | C | synonymous_variant | gacG | M5005_Spy0608 | c.516T>C | p.Tyr172Tyr | 276 | | | | | | 0 |
| 6752 | T | C | C | synonymous_variant | gacG | M5005_Spy0608 | c.526T>C | p.Leu176Leu | 1 | | | | | | 0 |
| 6761 | A | G | G | missense_variant | gacG | M5005_Spy0608 | c.535A>G | p.Ile179Val | 3 | | | | | | 0 |
| 6769 | A | G | G | synonymous_variant | gacG | M5005_Spy0608 | c.543A>G | p.Leu181Leu | 160 | | | | | | 0 |
| 6774 | A | G | G | missense_variant | gacG | M5005_Spy0608 | c.548A>G | p.Glu183Gly | 1 | | | | | | 0 |
| 6778 | C | A | A | synonymous_variant | gacG | M5005_Spy0608 | c.552C>A | p.Gly184Gly | 1 | | | | | | 0 |
| 6789 | A | G | G | missense_variant | gacG | M5005_Spy0608 | c.563A>G | p.Gln188Arg | 2 | | | | | | 0 |
| 6790 | G | A | A | synonymous_variant | gacG | M5005_Spy0608 | c.564G>A | p.Gln188Gln | 1 | | | | | | 0 |
| 6796 | C | A | A | synonymous_variant | gacG | M5005_Spy0608 | c.570C>A | p.Val190Val | 14 | | | | | | 0 |

| | | | | | | | | | | | | | | | |
|---|---|---|---|---|---|---|---|---|---|---|---|---|---|---|---|
| 6812 | G | A | A | missense_variant | gacG | M5005_Spy0608 | c.586G>A | p.Ala196Thr | 141 | | | | | | 0 |
| 6813 | C | T | T | missense_variant | gacG | M5005_Spy0608 | c.587C>T | p.Ala196Val | 110 | | | | | | 0 |
| 6831 | C | T | T | missense_variant | gacG | M5005_Spy0608 | c.605C>T | p.Pro202Leu | 8 | | | | | | 0 |
| 6833 | C | T | T | missense_variant | gacG | M5005_Spy0608 | c.607C>T | p.His203Tyr | 2 | | | | | | 0 |
| 6838 | G | A | A | synonymous_variant | gacG | M5005_Spy0608 | c.612G>A | p.Pro204Pro | 1 | | | | | | 0 |
| 6856 | T | C | C | synonymous_variant | gacG | M5005_Spy0608 | c.630T>C | p.Asn210Asn | 4 | | | | | | 0 |
| 6859 | A | G | G | synonymous_variant | gacG | M5005_Spy0608 | c.633A>G | p.Pro211Pro | 1 | | | | | | 0 |
| 6869 | T | C | C | synonymous_variant | gacG | M5005_Spy0608 | c.643T>C | p.Leu215Leu | 794 | | | | | | 0 |
| 6871 | A | G | G | synonymous_variant | gacG | M5005_Spy0608 | c.645A>G | p.Leu215Leu | 92 | | | | | | 0 |
| 6875 | C | T | T | missense_variant | gacG | M5005_Spy0608 | c.649C>T | p.His217Tyr | 1 | | | | | | 0 |
| 6919 | G | A | A | synonymous_variant | gacG | M5005_Spy0608 | c.693G>A | p.Gln231Gln | 1 | | | | | | 0 |
| 6920 | C | T,A | A | missense_variant | gacG | M5005_Spy0608 | c.694C>A | p.His232Asn | 18 | T | missense_variant | gacG | c.694C>T | p.His232Tyr | 4 |
| 6926 | G | A | A | missense_variant | gacG | M5005_Spy0608 | c.700G>A | p.Ala234Thr | 3 | | | | | | 0 |
| 6934 | C | T | T | synonymous_variant | gacG | M5005_Spy0608 | c.708C>T | p.Tyr236Tyr | 2 | | | | | | 0 |
| 6938 | T | C | C | synonymous_variant | gacG | M5005_Spy0608 | c.712T>C | p.Leu238Leu | 1688 | | | | | | 0 |
| 6950 | C | G | G | missense_variant | gacG | M5005_Spy0608 | c.724C>G | p.Arg242Gly | 35 | | | | | | 0 |
| 6952 | A | G | G | synonymous_variant | gacG | M5005_Spy0608 | c.726A>G | p.Arg242Arg | 2 | | | | | | 0 |
| 6955 | A | G | G | synonymous_variant | gacG | M5005_Spy0608 | c.729A>G | p.Glu243Glu | 2 | | | | | | 0 |
| 6960 | C | T | T | missense_variant | gacG | M5005_Spy0608 | c.734C>T | p.Thr245Ile | 1 | | | | | | 0 |
| 6961 | C | T | T | synonymous_variant | gacG | M5005_Spy0608 | c.735C>T | p.Thr245Thr | 9 | | | | | | 0 |
| 6962 | A | T | T | missense_variant | gacG | M5005_Spy0608 | c.736A>T | p.Asn246Tyr | 76 | | | | | | 0 |
| 6967 | T | C | C | synonymous_variant | gacG | M5005_Spy0608 | c.741T>C | p.Tyr247Tyr | 3 | | | | | | 0 |
| 6973 | A | G | G | missense_variant | gacG | M5005_Spy0608 | c.747A>G | p.Ile249Met | 25 | | | | | | 0 |
| 6974 | G | A | A | missense_variant | gacG | M5005_Spy0608 | c.748G>A | p.Asp250Asn | 1 | | | | | | 0 |
| 6979 | T | C | C | synonymous_variant | gacG | M5005_Spy0608 | c.753T>C | p.Leu251Leu | 1 | | | | | | 0 |
| 6983 | G | A | A | missense_variant | gacG | M5005_Spy0608 | c.757G>A | p.Val253Ile | 1 | | | | | | 0 |
| 7012 | A | G | G | synonymous_variant | gacG | M5005_Spy0608 | c.786A>G | p.Pro262Pro | 3 | | | | | | 0 |
| 7016 | A | G | G | missense_variant | gacG | M5005_Spy0608 | c.790A>G | p.Thr264Ala | 1938 | | | | | | 0 |
| 7017 | C | T | T | missense_variant | gacG | M5005_Spy0608 | c.791C>T | p.Thr264Ile | 2 | | | | | | 0 |
| 7020 | A | G | G | missense_variant | gacG | M5005_Spy0608 | c.794A>G | p.Lys265Arg | 31 | | | | | | 0 |
| 7041 | A | C | C | missense_variant | gacG | M5005_Spy0608 | c.815A>C | p.Tyr272Ser | 1 | | | | | | 0 |
| 7042 | C | T | T | synonymous_variant | gacG | M5005_Spy0608 | c.816C>T | p.Tyr272Tyr | 1 | | | | | | 0 |
| 7048 | C | T | T | synonymous_variant | gacG | M5005_Spy0608 | c.822C>T | p.Asn274Asn | 1 | | | | | | 0 |
| 7051 | C | T | T | synonymous_variant | gacG | M5005_Spy0608 | c.825C>T | p.Cys275Cys | 3 | | | | | | 0 |
| 7053 | A | G | G | missense_variant | gacG | M5005_Spy0608 | c.827A>G | p.Gln276Arg | 2 | | | | | | 0 |
| 7060 | A | G | G | synonymous_variant | gacG | M5005_Spy0608 | c.834A>G | p.Leu278Leu | 1 | | | | | | 0 |
| 7067 | C | A | A | missense_variant | gacG | M5005_Spy0608 | c.841C>A | p.Gln281Lys | 154 | | | | | | 0 |
| 7083 | T | C | C | missense_variant | gacG | M5005_Spy0608 | c.857T>C | p.Val286Ala | 4 | | | | | | 0 |
| 7112 | G | A | A | missense_variant | gacG | M5005_Spy0608 | c.886G>A | p.Asp296Asn | 2 | | | | | | 0 |
| 7118 | C | A | A | missense_variant | gacG | M5005_Spy0608 | c.892C>A | p.Leu298Ile | 1 | | | | | | 0 |
| 7132 | A | T | T | missense_variant | gacG | M5005_Spy0608 | c.906A>T | p.Leu302Phe | 1 | | | | | | 0 |
| 7136 | G | A | A | missense_variant | gacG | M5005_Spy0608 | c.910G>A | p.Ala304Thr | 17 | | | | | | 0 |
| 7137 | C | T | T | missense_variant | gacG | M5005_Spy0608 | c.911C>T | p.Ala304Val | 3 | | | | | | 0 |
| 7141 | C | T,A | A | missense_variant | gacG | M5005_Spy0608 | c.915C>A | p.Phe305Leu | 101 | T | synonymous_variant | gacG | c.915C>T | p.Phe305Phe | 17 |
| 7145 | A | G,T | G | missense_variant | gacG | M5005_Spy0608 | c.919A>G | p.Asn307Asp | 1907 | T | missense_variant | gacG | c.919A>T | p.Asn307Tyr | 18 |
| 7147 | C | T | T | synonymous_variant | gacG | M5005_Spy0608 | c.921C>T | p.Asn307Asn | 5 | | | | | | 0 |
| 7152 | A | G | G | missense_variant | gacG | M5005_Spy0608 | c.926A>G | p.Asn309Ser | 1 | | | | | | 0 |
| 7156 | C | T | T | synonymous_variant | gacG | M5005_Spy0608 | c.930C>T | p.Phe310Phe | 1 | | | | | | 0 |
| 7172 | A | G | G | missense_variant | gacG | M5005_Spy0608 | c.946A>G | p.Ile316Val | 3 | | | | | | 0 |
| 7177 | A | G | G | synonymous_variant | gacG | M5005_Spy0608 | c.951A>G | p.Thr317Thr | 21 | | | | | | 0 |
| 7199 | A | G | G | missense_variant | gacG | M5005_Spy0608 | c.973A>G | p.Lys325Glu | 15 | | | | | | 0 |
| 7202 | G | A | A | missense_variant | gacG | M5005_Spy0608 | c.976G>A | p.Glu326Lys | 19 | | | | | | 0 |
| 7221 | A | G | G | missense_variant | gacG | M5005_Spy0608 | c.995A>G | p.Gln332Arg | 2 | | | | | | 0 |
| 7225 | A | T,G | T | missense_variant | gacG | M5005_Spy0608 | c.999A>T | p.Arg333Ser | 17 | G | synonymous_variant | gacG | c.999A>G | p.Arg333Arg | 2 |
| 7230 | G | T | T | missense_variant | gacG | M5005_Spy0608 | c.1004G>T | p.Gly335Val | 18 | | | | | | 0 |
| 7233 | A | G | G | missense_variant | gacG | M5005_Spy0608 | c.1007A>G | p.Lys336Arg | 10 | | | | | | 0 |
| 7242 | A | G | G | missense_variant | gacG | M5005_Spy0608 | c.1016A>G | p.Asp339Gly | 1 | | | | | | 0 |
| 7249 | C | T | T | synonymous_variant | gacG | M5005_Spy0608 | c.1023C>T | p.Arg341Arg | 3 | | | | | | 0 |
| 7333 | C | T | T | synonymous_variant | gacG | M5005_Spy0608 | c.1107C>T | p.Phe369Phe | 1 | | | | | | 0 |
| 7393 | G | A | A | synonymous_variant | gacG | M5005_Spy0608 | c.1167G>A | p.Leu389Leu | 4 | | | | | | 0 |
| 7397 | G | A | A | missense_variant | gacG | M5005_Spy0608 | c.1171G>A | p.Asp391Asn | 4 | | | | | | 0 |
| 7405 | C | T | T | synonymous_variant | gacG | M5005_Spy0608 | c.1179C>T | p.Leu393Leu | 3 | | | | | | 0 |

| | | | | | | | | | | | | | | | |
|---|---|---|---|---|---|---|---|---|---|---|---|---|---|---|---|
| 7408 | G | T | T | synonymous_variant | gacG | M5005_Spy0608 | c.1182G>T | p.Val394Val | 33 | | | | | | 0 |
| 7413 | C | A | A | missense_variant | gacG | M5005_Spy0608 | c.1187C>A | p.Pro396Gln | 1 | | | | | | 0 |
| 7427 | C | A | A | missense_variant | gacG | M5005_Spy0608 | c.1201C>A | p.Leu401Ile | 1 | | | | | | 0 |
| 7433 | G | A | A | missense_variant | gacG | M5005_Spy0608 | c.1207G>A | p.Ala403Thr | 6 | | | | | | 0 |
| 7434 | C | T,G | G | missense_variant | gacG | M5005_Spy0608 | c.1208C>G | p.Ala403Gly | 129 | T | missense_variant | gacG | c.1208C>T | p.Ala403Val | 18 |
| 7445 | G | A,C | A | missense_variant | gacG | M5005_Spy0608 | c.1219G>A | p.Asp407Asn | 735 | C | missense_variant | gacG | c.1219G>C | p.Asp407His | 25 |
| 7457 | A | G | G | missense_variant | gacG | M5005_Spy0608 | c.1231A>G | p.Ile411Val | 9 | | | | | | 0 |
| 7459 | T | C | C | synonymous_variant | gacG | M5005_Spy0608 | c.1233T>C | p.Ile411Ile | 3 | | | | | | 0 |
| 7468 | A | G | G | synonymous_variant | gacG | M5005_Spy0608 | c.1242A>G | p.Ala414Ala | 1 | | | | | | 0 |
| 7504 | C | T | T | synonymous_variant | gacG | M5005_Spy0608 | c.1278C>T | p.Val426Val | 1 | | | | | | 0 |
| 7516 | T | C | C | synonymous_variant | gacG | M5005_Spy0608 | c.1290T>C | p.Asn430Asn | 4 | | | | | | 0 |
| 7528 | C | T | T | synonymous_variant | gacG | M5005_Spy0608 | c.1302C>T | p.Ile434Ile | 2 | | | | | | 0 |
| 7535 | G | A,C | A | missense_variant | gacG | M5005_Spy0608 | c.1309G>A | p.Glu437Lys | 17 | C | missense_variant | gacG | c.1309G>C | p.Glu437Gln | 5 |
| 7566 | T | C | C | missense_variant | gacG | M5005_Spy0608 | c.1340T>C | p.Val447Ala | 1 | | | | | | 0 |
| 7567 | T | A | A | synonymous_variant | gacG | M5005_Spy0608 | c.1341T>A | p.Val447Val | 1 | | | | | | 0 |
| 7573 | A | T | T | missense_variant | gacG | M5005_Spy0608 | c.1347A>T | p.Lys449Asn | 1 | | | | | | 0 |
| 7577 | A | T | T | missense_variant | gacG | M5005_Spy0608 | c.1351A>T | p.Ile451Phe | 1 | | | | | | 0 |
| 7581 | A | T | T | missense_variant | gacG | M5005_Spy0608 | c.1355A>T | p.Asp452Val | 1 | | | | | | 0 |
| 7585 | T | C | C | synonymous_variant | gacG | M5005_Spy0608 | c.1359T>C | p.Phe453Phe | 6 | | | | | | 0 |
| 7589 | G | A | A | missense_variant | gacG | M5005_Spy0608 | c.1363G>A | p.Ala455Thr | 118 | | | | | | 0 |
| 7597 | T | C | C | synonymous_variant | gacG | M5005_Spy0608 | c.1371T>C | p.Asp457Asp | 1 | | | | | | 0 |
| 7604 | G | A | A | missense_variant | gacG | M5005_Spy0608 | c.1378G>A | p.Val460Ile | 4 | | | | | | 0 |
| 7639 | C | T | T | synonymous_variant | gacG | M5005_Spy0608 | c.1413C>T | p.Tyr471Tyr | 12 | | | | | | 0 |
| 7663 | T | C | C | synonymous_variant | gacG | M5005_Spy0608 | c.1437T>C | p.Asp479Asp | 1 | | | | | | 0 |
| 7668 | A | G | G | missense_variant | gacG | M5005_Spy0608 | c.1442A>G | p.Glu481Gly | 1 | | | | | | 0 |
| 7675 | T | C | C | synonymous_variant | gacG | M5005_Spy0608 | c.1449T>C | p.Thr483Thr | 2 | | | | | | 0 |
| 7696 | A | G | G | synonymous_variant | gacG | M5005_Spy0608 | c.1470A>G | p.Glu490Glu | 473 | | | | | | 0 |
| 7724 | G | A | A | missense_variant | gacG | M5005_Spy0608 | c.1498G>A | p.Ala500Thr | 3 | | | | | | 0 |
| 7733 | C | A | A | synonymous_variant | gacG | M5005_Spy0608 | c.1507C>A | p.Arg503Arg | 12 | | | | | | 0 |
| 7739 | C | T | T | missense_variant | gacG | M5005_Spy0608 | c.1513C>T | p.Leu505Phe | 129 | | | | | | 0 |
| 7753 | G | A | A | synonymous_variant | gacG | M5005_Spy0608 | c.1527G>A | p.Ala509Ala | 1 | | | | | | 0 |
| 7760 | G | A | A | missense_variant | gacG | M5005_Spy0608 | c.1534G>A | p.Asp512Asn | 1949 | | | | | | 0 |
| 7761 | A | G | G | missense_variant | gacG | M5005_Spy0608 | c.1535A>G | p.Asp512Gly | 1 | | | | | | 0 |
| 7790 | C | T | T | missense_variant | gacG | M5005_Spy0608 | c.1564C>T | p.Pro522Ser | 94 | | | | | | 0 |
| 7805 | C | T | T | missense_variant | gacG | M5005_Spy0608 | c.1579C>T | p.Pro527Ser | 1 | | | | | | 0 |
| 7823 | C | T | T | synonymous_variant | gacG | M5005_Spy0608 | c.1597C>T | p.Leu533Leu | 2 | | | | | | 0 |
| 7852 | T | C | C | synonymous_variant | gacG | M5005_Spy0608 | c.1626T>C | p.Ala542Ala | 52 | | | | | | 0 |
| 7857 | C | G | G | missense_variant | gacG | M5005_Spy0608 | c.1631C>G | p.Thr544Ser | 11 | | | | | | 0 |
| 7861 | C | T | T | synonymous_variant | gacG | M5005_Spy0608 | c.1635C>T | p.Tyr545Tyr | 1 | | | | | | 0 |
| 7888 | C | A | A | synonymous_variant | gacG | M5005_Spy0608 | c.1662C>A | p.Ile554Ile | 1 | | | | | | 0 |
| 7891 | A | G | G | synonymous_variant | gacG | M5005_Spy0608 | c.1665A>G | p.Lys555Lys | 3 | | | | | | 0 |
| 7900 | G | A | A | synonymous_variant | gacG | M5005_Spy0608 | c.1674G>A | p.Leu558Leu | 1701 | | | | | | 0 |
| 7912 | C | T | T | synonymous_variant | gacG | M5005_Spy0608 | c.1686C>T | p.Ile562Ile | 856 | | | | | | 0 |
| 7918 | T | A | A | synonymous_variant | gacG | M5005_Spy0608 | c.1692T>A | p.Gly564Gly | 4 | | | | | | 0 |
| 7922 | G | A | A | missense_variant | gacG | M5005_Spy0608 | c.1696G>A | p.Ala566Thr | 2 | | | | | | 0 |
| 7927 | A | T,G | T | missense_variant | gacG | M5005_Spy0608 | c.1701A>T | p.Lys567Asn | 17 | G | synonymous_variant | gacG | c.1701A>G | p.Lys567Lys | 1 |
| 7936 | G | A | A | synonymous_variant | gacG | M5005_Spy0608 | c.1710G>A | p.Lys570Lys | 17 | | | | | | 0 |
| 7948 | G | T | T | missense_variant | gacG | M5005_Spy0608 | c.1722G>T | p.Leu574Phe | 2 | | | | | | 0 |
| 7964 | C | T | T | missense_variant | gacG | M5005_Spy0608 | c.1738C>T | p.Leu580Phe | 2 | | | | | | 0 |
| 7980 | C | T | T | synonymous_variant | gacH | M5005_Spy0609 | c.12C>T | p.Asp4Asp | 2 | | | | | | 0 |
| 7982 | C | T | T | missense_variant | gacH | M5005_Spy0609 | c.14C>T | p.Thr5Ile | 18 | | | | | | 0 |
| 7994 | C | T | T | missense_variant | gacH | M5005_Spy0609 | c.26C>T | p.Thr9Ile | 2 | | | | | | 0 |
| 8010 | T | C | C | synonymous_variant | gacH | M5005_Spy0609 | c.42T>C | p.Ile14Ile | 75 | | | | | | 0 |
| 8013 | T | C,A | A | missense_variant | gacH | M5005_Spy0609 | c.45T>A | p.Ser15Arg | 43 | C | synonymous_variant | gacH | c.45T>C | p.Ser15Ser | 57 |
| 8016 | C | T | T | synonymous_variant | gacH | M5005_Spy0609 | c.48C>T | p.His16His | 82 | | | | | | 0 |
| 8017 | C | T | T | missense_variant | gacH | M5005_Spy0609 | c.49C>T | p.His17Tyr | 4 | | | | | | 0 |
| 8024 | T | C | C | missense_variant | gacH | M5005_Spy0609 | c.56T>C | p.Ile19Thr | 442 | | | | | | 0 |
| 8028 | T | C | C | synonymous_variant | gacH | M5005_Spy0609 | c.60T>C | p.Leu20Leu | 2 | | | | | | 0 |
| 8038 | G | A | A | missense_variant | gacH | M5005_Spy0609 | c.70G>A | p.Gly24Ser | 68 | | | | | | 0 |
| 8058 | C | T | T | synonymous_variant | gacH | M5005_Spy0609 | c.90C>T | p.Tyr30Tyr | 391 | | | | | | 0 |
| 8067 | G | T | T | synonymous_variant | gacH | M5005_Spy0609 | c.99G>T | p.Ala33Ala | 6 | | | | | | 0 |
| 8077 | G | A | A | missense_variant | gacH | M5005_Spy0609 | c.109G>A | p.Val37Ile | 9 | | | | | | 0 |

| | | | | | | | | | | | | | | | |
|---|---|---|---|---|---|---|---|---|---|---|---|---|---|---|---|
| 8081 | G | A | A | missense_variant | gacH | M5005_Spy0609 | c.113G>A | p.Ser38Asn | 42 | | | | | | 0 |
| 8086 | A | G | G | missense_variant | gacH | M5005_Spy0609 | c.118A>G | p.Thr40Ala | 1 | | | | | | 0 |
| 8088 | G | A | A | synonymous_variant | gacH | M5005_Spy0609 | c.120G>A | p.Thr40Thr | 3 | | | | | | 0 |
| 8089 | G | A | A | missense_variant | gacH | M5005_Spy0609 | c.121G>A | p.Ala41Thr | 903 | | | | | | 0 |
| 8094 | A | G | G | synonymous_variant | gacH | M5005_Spy0609 | c.126A>G | p.Gln42Gln | 6 | | | | | | 0 |
| 8098 | G | A | A | missense_variant | gacH | M5005_Spy0609 | c.130G>A | p.Val44Ile | 6 | | | | | | 0 |
| 8105 | A | C | C | missense_variant | gacH | M5005_Spy0609 | c.137A>C | p.Tyr46Ser | 3 | | | | | | 0 |
| 8110 | G | A | A | missense_variant | gacH | M5005_Spy0609 | c.142G>A | p.Ala48Thr | 1794 | | | | | | 0 |
| 8111 | C | T | T | missense_variant | gacH | M5005_Spy0609 | c.143C>T | p.Ala48Val | 12 | | | | | | 0 |
| 8116 | T | C | C | synonymous_variant | gacH | M5005_Spy0609 | c.148T>C | p.Leu50Leu | 1 | | | | | | 0 |
| 8129 | C | T | T | missense_variant | gacH | M5005_Spy0609 | c.161C>T | p.Ser54Phe | 1 | | | | | | 0 |
| 8136 | G | T | T | synonymous_variant | gacH | M5005_Spy0609 | c.168G>T | p.Val56Val | 2 | | | | | | 0 |
| 8139 | A | C | C | synonymous_variant | gacH | M5005_Spy0609 | c.171A>C | p.Gly57Gly | 9 | | | | | | 0 |
| 8162 | T | C | C | missense_variant | gacH | M5005_Spy0609 | c.194T>C | p.Leu65Ser | 1 | | | | | | 0 |
| 8191 | C | A | A | synonymous_variant | gacH | M5005_Spy0609 | c.223C>A | p.Arg75Arg | 10 | | | | | | 0 |
| 8193 | A | C | C | synonymous_variant | gacH | M5005_Spy0609 | c.225A>C | p.Arg75Arg | 1 | | | | | | 0 |
| 8197 | T | C | C | synonymous_variant | gacH | M5005_Spy0609 | c.229T>C | p.Leu77Leu | 203 | | | | | | 0 |
| 8199 | G | A | A | synonymous_variant | gacH | M5005_Spy0609 | c.231G>A | p.Leu77Leu | 3 | | | | | | 0 |
| 8200 | T | C | C | missense_variant | gacH | M5005_Spy0609 | c.232T>C | p.Phe78Leu | 33 | | | | | | 0 |
| 8222 | T | C | C | missense_variant | gacH | M5005_Spy0609 | c.254T>C | p.Val85Ala | 20 | | | | | | 0 |
| 8239 | G | A | A | missense_variant | gacH | M5005_Spy0609 | c.271G>A | p.Val91Ile | 10 | | | | | | 0 |
| 8241 | C | T | T | synonymous_variant | gacH | M5005_Spy0609 | c.273C>T | p.Val91Val | 1 | | | | | | 0 |
| 8244 | C | T | T | synonymous_variant | gacH | M5005_Spy0609 | c.276C>T | p.Ser92Ser | 1 | | | | | | 0 |
| 8250 | G | C | C | missense_variant | gacH | M5005_Spy0609 | c.282G>C | p.Trp94Cys | 1 | | | | | | 0 |
| 8269 | C | T | T | synonymous_variant | gacH | M5005_Spy0609 | c.301C>T | p.Leu101Leu | 3 | | | | | | 0 |
| 8274 | T | C | C | synonymous_variant | gacH | M5005_Spy0609 | c.306T>C | p.Asn102Asn | 1 | | | | | | 0 |
| 8276 | A | T | T | missense_variant | gacH | M5005_Spy0609 | c.308A>T | p.Asp103Val | 6 | | | | | | 0 |
| 8280 | A | T | T | missense_variant | gacH | M5005_Spy0609 | c.312A>T | p.Lys104Asn | 4 | | | | | | 0 |
| 8281 | C | T | T | missense_variant | gacH | M5005_Spy0609 | c.313C>T | p.Arg105Trp | 1 | | | | | | 0 |
| 8283 | G | A | A | synonymous_variant | gacH | M5005_Spy0609 | c.315G>A | p.Arg105Arg | 1 | | | | | | 0 |
| 8303 | C | T | T | missense_variant | gacH | M5005_Spy0609 | c.335C>T | p.Thr112Ile | 1 | | | | | | 0 |
| 8319 | T | C | C | synonymous_variant | gacH | M5005_Spy0609 | c.351T>C | p.Tyr117Tyr | 48 | | | | | | 0 |
| 8330 | C | A | A | missense_variant | gacH | M5005_Spy0609 | c.362C>A | p.Ala121Asp | 1 | | | | | | 0 |
| 8334 | A | G | G | synonymous_variant | gacH | M5005_Spy0609 | c.366A>G | p.Leu122Leu | 1 | | | | | | 0 |
| 8339 | C | A,T | A | missense_variant | gacH | M5005_Spy0609 | c.371C>A | p.Ser124Tyr | 206 | T | missense_variant | gacH | c.371C>T | p.Ser124Phe | 4 |
| 8344 | C | T | T | missense_variant | gacH | M5005_Spy0609 | c.376C>T | p.Leu126Phe | 10 | | | | | | 0 |
| 8346 | C | T | T | synonymous_variant | gacH | M5005_Spy0609 | c.378C>T | p.Leu126Leu | 1 | | | | | | 0 |
| 8348 | T | G | G | missense_variant | gacH | M5005_Spy0609 | c.380T>G | p.Ile127Ser | 2 | | | | | | 0 |
| 8352 | T | C | C | synonymous_variant | gacH | M5005_Spy0609 | c.384T>C | p.Ile128Ile | 1 | | | | | | 0 |
| 8358 | G | A | A | synonymous_variant | gacH | M5005_Spy0609 | c.390G>A | p.Val130Val | 3 | | | | | | 0 |
| 8361 | G | A | A | missense_variant | gacH | M5005_Spy0609 | c.393G>A | p.Met131Ile | 43 | | | | | | 0 |
| 8363 | C | T | T | missense_variant | gacH | M5005_Spy0609 | c.395C>T | p.Ala132Val | 1 | | | | | | 0 |
| 8364 | C | T | T | synonymous_variant | gacH | M5005_Spy0609 | c.396C>T | p.Ala132Ala | 13 | | | | | | 0 |
| 8365 | A | G | G | missense_variant | gacH | M5005_Spy0609 | c.397A>G | p.Thr133Ala | 1 | | | | | | 0 |
| 8367 | T | C | C | synonymous_variant | gacH | M5005_Spy0609 | c.399T>C | p.Thr133Thr | 1 | | | | | | 0 |
| 8370 | A | G | G | synonymous_variant | gacH | M5005_Spy0609 | c.402A>G | p.Leu134Leu | 18 | | | | | | 0 |
| 8382 | G | A | A | synonymous_variant | gacH | M5005_Spy0609 | c.414G>A | p.Leu138Leu | 55 | | | | | | 0 |
| 8384 | T | C | C | missense_variant | gacH | M5005_Spy0609 | c.416T>C | p.Val139Ala | 1 | | | | | | 0 |
| 8385 | A | C | C | synonymous_variant | gacH | M5005_Spy0609 | c.417A>C | p.Val139Val | 2 | | | | | | 0 |
| 8387 | C | T | T | missense_variant | gacH | M5005_Spy0609 | c.419C>T | p.Ala140Val | 12 | | | | | | 0 |
| 8422 | T | C | C | synonymous_variant | gacH | M5005_Spy0609 | c.454T>C | p.Leu152Leu | 9 | | | | | | 0 |
| 8425 | G | T | T | missense_variant | gacH | M5005_Spy0609 | c.457G>T | p.Gly153Cys | 88 | | | | | | 0 |
| 8447 | C | T | T | missense_variant | gacH | M5005_Spy0609 | c.479C>T | p.Ser160Phe | 1 | | | | | | 0 |
| 8451 | A | G | G | synonymous_variant | gacH | M5005_Spy0609 | c.483A>G | p.Val161Val | 12 | | | | | | 0 |
| 8470 | G | A | A | missense_variant | gacH | M5005_Spy0609 | c.502G>A | p.Val168Met | 5 | | | | | | 0 |
| 8480 | A | G | G | missense_variant | gacH | M5005_Spy0609 | c.512A>G | p.Asn171Ser | 257 | | | | | | 0 |
| 8483 | A | G | G | missense_variant | gacH | M5005_Spy0609 | c.515A>G | p.Asp172Gly | 5 | | | | | | 0 |
| 8491 | C | T | T | synonymous_variant | gacH | M5005_Spy0609 | c.523C>T | p.Leu175Leu | 3 | | | | | | 0 |
| 8502 | A | G | G | synonymous_variant | gacH | M5005_Spy0609 | c.534A>G | p.Leu178Leu | 4 | | | | | | 0 |
| 8510 | C | A | A | missense_variant | gacH | M5005_Spy0609 | c.542C>A | p.Ser181Tyr | 1 | | | | | | 0 |
| 8530 | A | G | G | missense_variant | gacH | M5005_Spy0609 | c.562A>G | p.Thr188Ala | 3 | | | | | | 0 |
| 8531 | C | G | G | missense_variant | gacH | M5005_Spy0609 | c.563C>G | p.Thr188Arg | 502 | | | | | | 0 |

| | | | | | | | | | | | | | | | |
|---|---|---|---|---|---|---|---|---|---|---|---|---|---|---|---|
| 8534 | C | T | T | missense_variant | gacH | M5005_Spy0609 | c.566C>T | p.Ala189Val | 1 | | | | | | 0 |
| 8537 | G | A | A | missense_variant | gacH | M5005_Spy0609 | c.569G>A | p.Gly190Asp | 13 | | | | | | 0 |
| 8544 | C | A | A | missense_variant | gacH | M5005_Spy0609 | c.576C>A | p.Ser192Arg | 3 | | | | | | 0 |
| 8552 | C | T | T | missense_variant | gacH | M5005_Spy0609 | c.584C>T | p.Ala195Val | 16 | | | | | | 0 |
| 8556 | G | A | A | synonymous_variant | gacH | M5005_Spy0609 | c.588G>A | p.Leu196Leu | 6 | | | | | | 0 |
| 8570 | G | A | A | missense_variant | gacH | M5005_Spy0609 | c.602G>A | p.Gly201Glu | 2 | | | | | | 0 |
| 8571 | A | G | G | synonymous_variant | gacH | M5005_Spy0609 | c.603A>G | p.Gly201Gly | 1 | | | | | | 0 |
| 8572 | C | T | T | synonymous_variant | gacH | M5005_Spy0609 | c.604C>T | p.Leu202Leu | 1 | | | | | | 0 |
| 8574 | G | A | A | synonymous_variant | gacH | M5005_Spy0609 | c.606G>A | p.Leu202Leu | 4 | | | | | | 0 |
| 8576 | T | C,G | C | missense_variant | gacH | M5005_Spy0609 | c.608T>C | p.Leu203Ser | 2 | G | missense_variant | gacH | c.608T>G | p.Leu203Trp | 1 |
| 8578 | A | T | T | missense_variant | gacH | M5005_Spy0609 | c.610A>T | p.Ile204Leu | 30 | | | | | | 0 |
| 8579 | T | C | C | missense_variant | gacH | M5005_Spy0609 | c.611T>C | p.Ile204Thr | 1 | | | | | | 0 |
| 8580 | A | C | C | synonymous_variant | gacH | M5005_Spy0609 | c.612A>C | p.Ile204Ile | 2 | | | | | | 0 |
| 8584 | T | C,A | A | missense_variant | gacH | M5005_Spy0609 | c.616T>A | p.Leu206Ile | 11 | C | synonymous_variant | gacH | c.616T>C | p.Leu206Leu | 1 |
| 8587 | C | T | T | missense_variant | gacH | M5005_Spy0609 | c.619C>T | p.Leu207Phe | 1 | | | | | | 0 |
| 8589 | T | C | C | synonymous_variant | gacH | M5005_Spy0609 | c.621T>C | p.Leu207Leu | 11 | | | | | | 0 |
| 8592 | C | A | A | synonymous_variant | gacH | M5005_Spy0609 | c.624C>A | p.Val208Val | 2 | | | | | | 0 |
| 8603 | C | T | T | missense_variant | gacH | M5005_Spy0609 | c.635C>T | p.Thr212Ile | 56 | | | | | | 0 |
| 8611 | T | G | G | missense_variant | gacH | M5005_Spy0609 | c.643T>G | p.Tyr215Asp | 1 | | | | | | 0 |
| 8642 | A | G | G | missense_variant | gacH | M5005_Spy0609 | c.674A>G | p.Lys225Arg | 5 | | | | | | 0 |
| 8643 | G | C | C | missense_variant | gacH | M5005_Spy0609 | c.675G>C | p.Lys225Asn | 58 | | | | | | 0 |
| 8645 | C | T | T | missense_variant | gacH | M5005_Spy0609 | c.677C>T | p.Ser226Leu | 6 | | | | | | 0 |
| 8653 | C | T | T | missense_variant | gacH | M5005_Spy0609 | c.685C>T | p.Pro229Ser | 302 | | | | | | 0 |
| 8654 | C | T | T | missense_variant | gacH | M5005_Spy0609 | c.686C>T | p.Pro229Leu | 92 | | | | | | 0 |
| 8672 | C | T | T | missense_variant | gacH | M5005_Spy0609 | c.704C>T | p.Ala235Val | 9 | | | | | | 0 |
| 8674 | A | G | G | missense_variant | gacH | M5005_Spy0609 | c.706A>G | p.Thr236Ala | 1 | | | | | | 0 |
| 8680 | A | G | G | missense_variant | gacH | M5005_Spy0609 | c.712A>G | p.Ser238Gly | 12 | | | | | | 0 |
| 8683 | T | C | C | synonymous_variant | gacH | M5005_Spy0609 | c.715T>C | p.Leu239Leu | 35 | | | | | | 0 |
| 8700 | T | C | C | synonymous_variant | gacH | M5005_Spy0609 | c.732T>C | p.Val244Val | 13 | | | | | | 0 |
| 8706 | C | T | T | synonymous_variant | gacH | M5005_Spy0609 | c.738C>T | p.Asn246Asn | 7 | | | | | | 0 |
| 8715 | C | T | T | synonymous_variant | gacH | M5005_Spy0609 | c.747C>T | p.Phe249Phe | 156 | | | | | | 0 |
| 8724 | A | C | C | synonymous_variant | gacH | M5005_Spy0609 | c.756A>C | p.Gly252Gly | 374 | | | | | | 0 |
| 8745 | G | T | T | missense_variant | gacH | M5005_Spy0609 | c.777G>T | p.Leu259Phe | 1 | | | | | | 0 |
| 8751 | A | G | G | synonymous_variant | gacH | M5005_Spy0609 | c.783A>G | p.Gly261Gly | 2 | | | | | | 0 |
| 8759 | T | C | C | missense_variant | gacH | M5005_Spy0609 | c.791T>C | p.Val264Ala | 17 | | | | | | 0 |
| 8767 | G | A | A | missense_variant | gacH | M5005_Spy0609 | c.799G>A | p.Gly267Arg | 3 | | | | | | 0 |
| 8769 | A | G | G | synonymous_variant | gacH | M5005_Spy0609 | c.801A>G | p.Gly267Gly | 7 | | | | | | 0 |
| 8770 | G | A | A | missense_variant | gacH | M5005_Spy0609 | c.802G>A | p.Ala268Thr | 54 | | | | | | 0 |
| 8771 | C | T | T | missense_variant | gacH | M5005_Spy0609 | c.803C>T | p.Ala268Val | 2 | | | | | | 0 |
| 8772 | T | C | C | synonymous_variant | gacH | M5005_Spy0609 | c.804T>C | p.Ala268Ala | 1 | | | | | | 0 |
| 8797 | A | C | C | missense_variant | gacH | M5005_Spy0609 | c.829A>C | p.Thr277Pro | 1 | | | | | | 0 |
| 8799 | A | G | G | synonymous_variant | gacH | M5005_Spy0609 | c.831A>G | p.Thr277Thr | 7 | | | | | | 0 |
| 8806 | G | T | T | missense_variant | gacH | M5005_Spy0609 | c.838G>T | p.Ala280Ser | 1 | | | | | | 0 |
| 8809 | C | A | A | missense_variant | gacH | M5005_Spy0609 | c.841C>A | p.Leu281Ile | 92 | | | | | | 0 |
| 8820 | C | T | T | synonymous_variant | gacH | M5005_Spy0609 | c.852C>T | p.Tyr284Tyr | 4 | | | | | | 0 |
| 8825 | T | C | C | missense_variant | gacH | M5005_Spy0609 | c.857T>C | p.Ile286Thr | 1 | | | | | | 0 |
| 8845 | A | C | C | missense_variant | gacH | M5005_Spy0609 | c.877A>C | p.Thr293Pro | 30 | | | | | | 0 |
| 8852 | T | C | C | missense_variant | gacH | M5005_Spy0609 | c.884T>C | p.Phe295Ser | 16 | | | | | | 0 |
| 8863 | A | C | C | missense_variant | gacH | M5005_Spy0609 | c.895A>C | p.Ile299Leu | 11 | | | | | | 0 |
| 8866 | C | T | T | synonymous_variant | gacH | M5005_Spy0609 | c.898C>T | p.Leu300Leu | 1 | | | | | | 0 |
| 8867 | T | C | C | missense_variant | gacH | M5005_Spy0609 | c.899T>C | p.Leu300Pro | 6 | | | | | | 0 |
| 8873 | C | A | A | missense_variant | gacH | M5005_Spy0609 | c.905C>A | p.Thr302Lys | 1 | | | | | | 0 |
| 8877 | T | A | A | synonymous_variant | gacH | M5005_Spy0609 | c.909T>A | p.Ile303Ile | 3 | | | | | | 0 |
| 8878 | A | T | T | missense_variant | gacH | M5005_Spy0609 | c.910A>T | p.Ile304Phe | 1 | | | | | | 0 |
| 8883 | T | G | G | synonymous_variant | gacH | M5005_Spy0609 | c.915T>G | p.Ser305Ser | 2 | | | | | | 0 |
| 8887 | G | T | T | missense_variant | gacH | M5005_Spy0609 | c.919G>T | p.Val307Phe | 1 | | | | | | 0 |
| 8889 | T | A | A | synonymous_variant | gacH | M5005_Spy0609 | c.921T>A | p.Val307Val | 1 | | | | | | 0 |
| 8897 | T | A | A | stop_gained | gacH | M5005_Spy0609 | c.929T>A | p.Leu310* | 3 | | | | | | 0 |
| 8902 | G | A | A | missense_variant | gacH | M5005_Spy0609 | c.934G>A | p.Glu312Lys | 1 | | | | | | 0 |
| 8906 | C | A | A | stop_gained | gacH | M5005_Spy0609 | c.938C>A | p.Ser313* | 1 | | | | | | 0 |
| 8909 | T | G,* | G | missense_variant | gacH | M5005_Spy0609 | c.941T>G | p.Met314Arg | 3 | | | | | | 1 |
| 8914 | A | G | G | missense_variant | gacH | M5005_Spy0609 | c.946A>G | p.Ser316Gly | 2 | | | | | | 0 |

| | | | | | | | | | | | | | | | |
|---|---|---|---|---|---|---|---|---|---|---|---|---|---|---|---|
| 8916 | C | G | G | missense_variant | gacH | M5005_Spy0609 | c.948C>G | p.Ser316Arg | 2 | | | | | | 0 |
| 8922 | G | A | A | synonymous_variant | gacH | M5005_Spy0609 | c.954G>A | p.Pro318Pro | 510 | | | | | | 0 |
| 8933 | C | G | G | missense_variant | gacH | M5005_Spy0609 | c.965C>G | p.Thr322Ser | 7 | | | | | | 0 |
| 8947 | T | C | C | synonymous_variant | gacH | M5005_Spy0609 | c.979T>C | p.Leu327Leu | 1 | | | | | | 0 |
| 8973 | C | T | T | synonymous_variant | gacH | M5005_Spy0609 | c.1005C>T | p.Ser335Ser | 11 | | | | | | 0 |
| 8989 | G | T,A | A | missense_variant | gacH | M5005_Spy0609 | c.1021G>A | p.Val341Met | 39 | T | missense_variant | gacH | c.1021G>T | p.Val341Leu | 1 |
| 9004 | G | A | A | missense_variant | gacH | M5005_Spy0609 | c.1036G>A | p.Val346Ile | 7 | | | | | | 0 |
| 9007 | G | T | T | missense_variant | gacH | M5005_Spy0609 | c.1039G>T | p.Val347Leu | 2 | | | | | | 0 |
| 9033 | A | G | G | synonymous_variant | gacH | M5005_Spy0609 | c.1065A>G | p.Val355Val | 1383 | | | | | | 0 |
| 9046 | C | T | T | missense_variant | gacH | M5005_Spy0609 | c.1078C>T | p.His360Tyr | 1 | | | | | | 0 |
| 9050 | G | A | A | missense_variant | gacH | M5005_Spy0609 | c.1082G>A | p.Gly361Asp | 34 | | | | | | 0 |
| 9069 | A | T | T | missense_variant | gacH | M5005_Spy0609 | c.1101A>T | p.Lys367Asn | 1 | | | | | | 0 |
| 9078 | G | A | A | missense_variant | gacH | M5005_Spy0609 | c.1110G>A | p.Met370Ile | 1 | | | | | | 0 |
| 9082 | C | T | T | missense_variant | gacH | M5005_Spy0609 | c.1114C>T | p.Pro372Ser | 1 | | | | | | 0 |
| 9087 | C | T | T | synonymous_variant | gacH | M5005_Spy0609 | c.1119C>T | p.Val373Val | 1 | | | | | | 0 |
| 9095 | C | T | T | missense_variant | gacH | M5005_Spy0609 | c.1127C>T | p.Ala376Val | 52 | | | | | | 0 |
| 9100 | G | A | A | missense_variant | gacH | M5005_Spy0609 | c.1132G>A | p.Ala378Thr | 18 | | | | | | 0 |
| 9103 | G | C | C | missense_variant | gacH | M5005_Spy0609 | c.1135G>C | p.Val379Leu | 2 | | | | | | 0 |
| 9115 | T | G,A | A | missense_variant | gacH | M5005_Spy0609 | c.1147T>A | p.Phe383Ile | 197 | G | missense_variant | gacH | c.1147T>G | p.Phe383Val | 70 |
| 9126 | T | C | C | synonymous_variant | gacH | M5005_Spy0609 | c.1158T>C | p.Ser386Ser | 48 | | | | | | 0 |
| 9129 | T | C | C | synonymous_variant | gacH | M5005_Spy0609 | c.1161T>C | p.Cys387Cys | 2 | | | | | | 0 |
| 9136 | T | C | C | synonymous_variant | gacH | M5005_Spy0609 | c.1168T>C | p.Leu390Leu | 1 | | | | | | 0 |
| 9145 | T | C | C | missense_variant | gacH | M5005_Spy0609 | c.1177T>C | p.Phe393Leu | 6 | | | | | | 0 |
| 9166 | A | G | G | missense_variant | gacH | M5005_Spy0609 | c.1198A>G | p.Lys400Glu | 1 | | | | | | 0 |
| 9172 | T | C | C | synonymous_variant | gacH | M5005_Spy0609 | c.1204T>C | p.Leu402Leu | 18 | | | | | | 0 |
| 9179 | G | A | A | missense_variant | gacH | M5005_Spy0609 | c.1211G>A | p.Gly404Asp | 2 | | | | | | 0 |
| 9186 | G | T | T | synonymous_variant | gacH | M5005_Spy0609 | c.1218G>T | p.Pro406Pro | 1 | | | | | | 0 |
| 9197 | C | T | T | missense_variant | gacH | M5005_Spy0609 | c.1229C>T | p.Ala410Val | 6 | | | | | | 0 |
| 9216 | C | T | T | synonymous_variant | gacH | M5005_Spy0609 | c.1248C>T | p.Asp416Asp | 1240 | | | | | | 0 |
| 9222 | C | T | T | synonymous_variant | gacH | M5005_Spy0609 | c.1254C>T | p.Asn418Asn | 48 | | | | | | 0 |
| 9228 | A | G | G | synonymous_variant | gacH | M5005_Spy0609 | c.1260A>G | p.Leu420Leu | 2 | | | | | | 0 |
| 9244 | G | A | A | missense_variant | gacH | M5005_Spy0609 | c.1276G>A | p.Ala426Thr | 3 | | | | | | 0 |
| 9268 | G | A | A | missense_variant | gacH | M5005_Spy0609 | c.1300G>A | p.Val434Ile | 1 | | | | | | 0 |
| 9283 | G | A | A | missense_variant | gacH | M5005_Spy0609 | c.1315G>A | p.Val439Met | 23 | | | | | | 0 |
| 9291 | G | A | A | synonymous_variant | gacH | M5005_Spy0609 | c.1323G>A | p.Lys441Lys | 2 | | | | | | 0 |
| 9296 | T | A | A | missense_variant | gacH | M5005_Spy0609 | c.1328T>A | p.Ile443Lys | 1 | | | | | | 0 |
| 9301 | G | A | A | missense_variant | gacH | M5005_Spy0609 | c.1333G>A | p.Glu445Lys | 1 | | | | | | 0 |
| 9306 | A | G | G | synonymous_variant | gacH | M5005_Spy0609 | c.1338A>G | p.Lys446Lys | 9 | | | | | | 0 |
| 9310 | A | G | G | missense_variant | gacH | M5005_Spy0609 | c.1342A>G | p.Thr448Ala | 7 | | | | | | 0 |
| 9313 | A | G | G | missense_variant | gacH | M5005_Spy0609 | c.1345A>G | p.Asn449Asp | 6 | | | | | | 0 |
| 9321 | C | T | T | synonymous_variant | gacH | M5005_Spy0609 | c.1353C>T | p.Ser451Ser | 229 | | | | | | 0 |
| 9322 | C | A | A | missense_variant | gacH | M5005_Spy0609 | c.1354C>A | p.Gln452Lys | 3 | | | | | | 0 |
| 9327 | A | C | C | missense_variant | gacH | M5005_Spy0609 | c.1359A>C | p.Glu453Asp | 117 | | | | | | 0 |
| 9335 | C | T | T | missense_variant | gacH | M5005_Spy0609 | c.1367C>T | p.Ala456Val | 1 | | | | | | 0 |
| 9336 | G | A | A | synonymous_variant | gacH | M5005_Spy0609 | c.1368G>A | p.Ala456Ala | 7 | | | | | | 0 |
| 9344 | C | T | T | missense_variant | gacH | M5005_Spy0609 | c.1376C>T | p.Ala459Val | 1 | | | | | | 0 |
| 9384 | C | T | T | synonymous_variant | gacH | M5005_Spy0609 | c.1416C>T | p.Asp472Asp | 1 | | | | | | 0 |
| 9393 | T | C | C | synonymous_variant | gacH | M5005_Spy0609 | c.1425T>C | p.Asn475Asn | 13 | | | | | | 0 |
| 9395 | A | G | G | missense_variant | gacH | M5005_Spy0609 | c.1427A>G | p.Asn476Ser | 1 | | | | | | 0 |
| 9400 | G | A | A | missense_variant | gacH | M5005_Spy0609 | c.1432G>A | p.Ala478Thr | 61 | | | | | | 0 |
| 9435 | C | T | T | synonymous_variant | gacH | M5005_Spy0609 | c.1467C>T | p.Ser489Ser | 636 | | | | | | 0 |
| 9450 | T | G | G | missense_variant | gacH | M5005_Spy0609 | c.1482T>G | p.Asp494Glu | 22 | | | | | | 0 |
| 9461 | A | G | G | missense_variant | gacH | M5005_Spy0609 | c.1493A>G | p.Asn498Ser | 16 | | | | | | 0 |
| 9463 | G | A | A | missense_variant | gacH | M5005_Spy0609 | c.1495G>A | p.Val499Ile | 5 | | | | | | 0 |
| 9467 | C | T,A | A | missense_variant | gacH | M5005_Spy0609 | c.1499C>A | p.Thr500Asn | 7 | T | missense_variant | gacH | c.1499C>T | p.Thr500Ile | 1 |
| 9474 | C | T | T | synonymous_variant | gacH | M5005_Spy0609 | c.1506C>T | p.Ser502Ser | 2 | | | | | | 0 |
| 9475 | C | T | T | missense_variant | gacH | M5005_Spy0609 | c.1507C>T | p.His503Tyr | 1 | | | | | | 0 |
| 9477 | C | T | T | synonymous_variant | gacH | M5005_Spy0609 | c.1509C>T | p.His503His | 170 | | | | | | 0 |
| 9500 | C | T | T | missense_variant | gacH | M5005_Spy0609 | c.1532C>T | p.Ala511Val | 1 | | | | | | 0 |
| 9502 | A | G | G | missense_variant | gacH | M5005_Spy0609 | c.1534A>G | p.Ile512Val | 1 | | | | | | 0 |
| 9513 | C | T | T | synonymous_variant | gacH | M5005_Spy0609 | c.1545C>T | p.Ser515Ser | 4 | | | | | | 0 |
| 9522 | G | T,A | A | synonymous_variant | gacH | M5005_Spy0609 | c.1554G>A | p.Ala518Ala | 40 | T | synonymous_variant | gacH | c.1554G>T | p.Ala518Ala | 10 |

| | | | | | | | | | | | | | | | |
|---|---|---|---|---|---|---|---|---|---|---|---|---|---|---|---|
| 9525 | A | G | G | synonymous_variant | gacH | M5005_Spy0609 | c.1557A>G | p.Gly519Gly | 8 | | | | | | 0 |
| 9526 | C | T | T | missense_variant | gacH | M5005_Spy0609 | c.1558C>T | p.Leu520Phe | 27 | | | | | | 0 |
| 9528 | C | T | T | synonymous_variant | gacH | M5005_Spy0609 | c.1560C>T | p.Leu520Leu | 33 | | | | | | 0 |
| 9546 | C | T | T | synonymous_variant | gacH | M5005_Spy0609 | c.1578C>T | p.Tyr526Tyr | 2 | | | | | | 0 |
| 9549 | G | A | A | synonymous_variant | gacH | M5005_Spy0609 | c.1581G>A | p.Gly527Gly | 1 | | | | | | 0 |
| 9558 | G | T | T | synonymous_variant | gacH | M5005_Spy0609 | c.1590G>T | p.Thr530Thr | 1117 | | | | | | 0 |
| 9579 | G | A | A | synonymous_variant | gacH | M5005_Spy0609 | c.1611G>A | p.Thr537Thr | 6 | | | | | | 0 |
| 9586 | A | G | G | missense_variant | gacH | M5005_Spy0609 | c.1618A>G | p.Ser540Gly | 1 | | | | | | 0 |
| 9589 | T | G | G | missense_variant | gacH | M5005_Spy0609 | c.1621T>G | p.Leu541Val | 36 | | | | | | 0 |
| 9592 | C | T | T | missense_variant | gacH | M5005_Spy0609 | c.1624C>T | p.Pro542Ser | 5 | | | | | | 0 |
| 9594 | T | C | C | synonymous_variant | gacH | M5005_Spy0609 | c.1626T>C | p.Pro542Pro | 1 | | | | | | 0 |
| 9609 | T | G | G | synonymous_variant | gacH | M5005_Spy0609 | c.1641T>G | p.Ser547Ser | 1 | | | | | | 0 |
| 9614 | C | T | T | missense_variant | gacH | M5005_Spy0609 | c.1646C>T | p.Ser549Leu | 1 | | | | | | 0 |
| 9618 | A | G | G | synonymous_variant | gacH | M5005_Spy0609 | c.1650A>G | p.Val550Val | 3 | | | | | | 0 |
| 9624 | T | C | C | synonymous_variant | gacH | M5005_Spy0609 | c.1656T>C | p.Val552Val | 79 | | | | | | 0 |
| 9639 | C | T | T | synonymous_variant | gacH | M5005_Spy0609 | c.1671C>T | p.Val557Val | 2 | | | | | | 0 |
| 9645 | T | C | C | synonymous_variant | gacH | M5005_Spy0609 | c.1677T>C | p.Pro559Pro | 116 | | | | | | 0 |
| 9648 | A | C | C | missense_variant | gacH | M5005_Spy0609 | c.1680A>C | p.Lys560Asn | 7 | | | | | | 0 |
| 9654 | C | T | T | synonymous_variant | gacH | M5005_Spy0609 | c.1686C>T | p.Ala562Ala | 10 | | | | | | 0 |
| 9659 | C | A | A | missense_variant | gacH | M5005_Spy0609 | c.1691C>A | p.Pro564His | 1 | | | | | | 0 |
| 9712 | G | T | T | missense_variant | gacH | M5005_Spy0609 | c.1744G>T | p.Ala582Ser | 1 | | | | | | 0 |
| 9723 | C | T | T | synonymous_variant | gacH | M5005_Spy0609 | c.1755C>T | p.Asn585Asn | 4 | | | | | | 0 |
| 9765 | C | A,T | A | synonymous_variant | gacH | M5005_Spy0609 | c.1797C>A | p.Ser599Ser | 103 | T | synonymous_variant | gacH | c.1797C>T | p.Ser599Ser | 2 |
| 9787 | T | G | G | missense_variant | gacH | M5005_Spy0609 | c.1819T>G | p.Ser607Ala | 1 | | | | | | 0 |
| 9807 | C | T | T | synonymous_variant | gacH | M5005_Spy0609 | c.1839C>T | p.Asn613Asn | 2 | | | | | | 0 |
| 9819 | C | T | T | synonymous_variant | gacH | M5005_Spy0609 | c.1851C>T | p.Val617Val | 10 | | | | | | 0 |
| 9822 | T | C | C | synonymous_variant | gacH | M5005_Spy0609 | c.1854T>C | p.Gly618Gly | 9 | | | | | | 0 |
| 9834 | C | T | T | synonymous_variant | gacH | M5005_Spy0609 | c.1866C>T | p.Ser622Ser | 48 | | | | | | 0 |
| 9878 | G | T | T | missense_variant | gacH | M5005_Spy0609 | c.1910G>T | p.Ser637Ile | 1 | | | | | | 0 |
| 9879 | T | C | C | synonymous_variant | gacH | M5005_Spy0609 | c.1911T>C | p.Ser637Ser | 76 | | | | | | 0 |
| 9880 | G | C | C | missense_variant | gacH | M5005_Spy0609 | c.1912G>C | p.Glu638Gln | 4 | | | | | | 0 |
| 9885 | C | T | T | synonymous_variant | gacH | M5005_Spy0609 | c.1917C>T | p.Ser639Ser | 2 | | | | | | 0 |
| 9894 | C | T | T | synonymous_variant | gacH | M5005_Spy0609 | c.1926C>T | p.Phe642Phe | 4 | | | | | | 0 |
| 9942 | T | C | C | synonymous_variant | gacH | M5005_Spy0609 | c.1974T>C | p.Pro658Pro | 3 | | | | | | 0 |
| 9945 | A | G | G | synonymous_variant | gacH | M5005_Spy0609 | c.1977A>G | p.Glu659Glu | 2 | | | | | | 0 |
| 9952 | G | A | A | missense_variant | gacH | M5005_Spy0609 | c.1984G>A | p.Val662Ile | 43 | | | | | | 0 |
| 9975 | G | A | A | synonymous_variant | gacH | M5005_Spy0609 | c.2007G>A | p.Thr669Thr | 18 | | | | | | 0 |
| 9996 | A | T | T | synonymous_variant | gacH | M5005_Spy0609 | c.2028A>T | p.Leu676Leu | 1 | | | | | | 0 |
| 10011 | G | A | A | synonymous_variant | gacH | M5005_Spy0609 | c.2043G>A | p.Arg681Arg | 4 | | | | | | 0 |
| 10020 | G | A | A | synonymous_variant | gacH | M5005_Spy0609 | c.2052G>A | p.Ser684Ser | 2 | | | | | | 0 |
| 10033 | G | A | A | missense_variant | gacH | M5005_Spy0609 | c.2065G>A | p.Glu689Lys | 17 | | | | | | 0 |
| 10040 | G | A | A | missense_variant | gacH | M5005_Spy0609 | c.2072G>A | p.Arg691Lys | 37 | | | | | | 0 |
| 10043 | C | A | A | missense_variant | gacH | M5005_Spy0609 | c.2075C>A | p.Ala692Glu | 1 | | | | | | 0 |
| 10048 | T | C | C | synonymous_variant | gacH | M5005_Spy0609 | c.2080T>C | p.Leu694Leu | 22 | | | | | | 0 |
| 10061 | C | T | T | missense_variant | gacH | M5005_Spy0609 | c.2093C>T | p.Thr698Ile | 1 | | | | | | 0 |
| 10062 | A | C | C | synonymous_variant | gacH | M5005_Spy0609 | c.2094A>C | p.Thr698Thr | 1 | | | | | | 0 |
| 10071 | C | T | T | synonymous_variant | gacH | M5005_Spy0609 | c.2103C>T | p.Asn701Asn | 9 | | | | | | 0 |
| 10074 | G | A | A | synonymous_variant | gacH | M5005_Spy0609 | c.2106G>A | p.Lys702Lys | 18 | | | | | | 0 |
| 10075 | C | A | A | missense_variant | gacH | M5005_Spy0609 | c.2107C>A | p.Pro703Thr | 3 | | | | | | 0 |
| 10083 | A | G | G | synonymous_variant | gacH | M5005_Spy0609 | c.2115A>G | p.Thr705Thr | 1 | | | | | | 0 |
| 10085 | T | C | C | missense_variant | gacH | M5005_Spy0609 | c.2117T>C | p.Val706Ala | 18 | | | | | | 0 |
| 10089 | G | A,T | A | synonymous_variant | gacH | M5005_Spy0609 | c.2121G>A | p.Val707Val | 3 | T | synonymous_variant | gacH | c.2121G>T | p.Val707Val | 2 |
| 10113 | T | C | C | synonymous_variant | gacH | M5005_Spy0609 | c.2145T>C | p.Gly715Gly | 2 | | | | | | 0 |
| 10137 | C | T | T | synonymous_variant | gacH | M5005_Spy0609 | c.2169C>T | p.Asn723Asn | 1 | | | | | | 0 |
| 10181 | G | A | A | missense_variant | gacH | M5005_Spy0609 | c.2213G>A | p.Ser738Asn | 2 | | | | | | 0 |
| 10199 | A | G | G | missense_variant | gacH | M5005_Spy0609 | c.2231A>G | p.Lys744Arg | 289 | | | | | | 0 |
| 10226 | G | A | A | missense_variant | gacH | M5005_Spy0609 | c.2258G>A | p.Ser753Asn | 26 | | | | | | 0 |
| 10233 | T | C | C | synonymous_variant | gacH | M5005_Spy0609 | c.2265T>C | p.Phe755Phe | 1 | | | | | | 0 |
| 10245 | A | G | G | synonymous_variant | gacH | M5005_Spy0609 | c.2277A>G | p.Leu759Leu | 20 | | | | | | 0 |
| 10269 | A | G | G | synonymous_variant | gacH | M5005_Spy0609 | c.2301A>G | p.Val767Val | 4 | | | | | | 0 |
| 10272 | A | G | G | synonymous_variant | gacH | M5005_Spy0609 | c.2304A>G | p.Ser768Ser | 6 | | | | | | 0 |
| 10314 | T | C | C | synonymous_variant | gacH | M5005_Spy0609 | c.2346T>C | p.Ser782Ser | 11 | | | | | | 0 |

| | | | | | | | | | | | | | | | |
|---|---|---|---|---|---|---|---|---|---|---|---|---|---|---|---|
| 10315 | G | A | A | missense_variant | gacH | M5005_Spy0609 | c.2347G>A | p.Val783Ile | 1 | | | | | | 0 |
| 10317 | C | T | T | synonymous_variant | gacH | M5005_Spy0609 | c.2349C>T | p.Val783Val | 8 | | | | | | 0 |
| 10328 | C | T | T | missense_variant | gacH | M5005_Spy0609 | c.2360C>T | p.Pro787Leu | 22 | | | | | | 0 |
| 10335 | T | C | C | synonymous_variant | gacH | M5005_Spy0609 | c.2367T>C | p.Ser789Ser | 1 | | | | | | 0 |
| 10336 | C | T | T | missense_variant | gacH | M5005_Spy0609 | c.2368C>T | p.Pro790Ser | 4 | | | | | | 0 |
| 10339 | G | A | A | missense_variant | gacH | M5005_Spy0609 | c.2371G>A | p.Glu791Lys | 7 | | | | | | 0 |
| 10349 | C | T | T | missense_variant | gacH | M5005_Spy0609 | c.2381C>T | p.Ala794Val | 1 | | | | | | 0 |
| 10357 | A | G | G | missense_variant | gacH | M5005_Spy0609 | c.2389A>G | p.Asn797Asp | 19 | | | | | | 0 |
| 10363 | T | C | C | synonymous_variant | gacH | M5005_Spy0609 | c.2395T>C | p.Leu799Leu | 25 | | | | | | 0 |
| 10372 | A | G | G | missense_variant | gacH | M5005_Spy0609 | c.2404A>G | p.Ile802Val | 92 | | | | | | 0 |
| 10373 | T | C | C | missense_variant | gacH | M5005_Spy0609 | c.2405T>C | p.Ile802Thr | 1 | | | | | | 0 |
| 10377 | A | G | G | synonymous_variant | gacH | M5005_Spy0609 | c.2409A>G | p.Gln803Gln | 1 | | | | | | 0 |
| 10435 | T | C | C | missense_variant | gacH | M5005_Spy0609 | c.2467T>C | p.Ser823Pro | 31 | | | | | | 0 |
| 10455 | G | T | T | upstream_gene_variant | intergenic | | | | 6 | | | | | | 0 |
| 10456 | G | A | A | upstream_gene_variant | intergenic | | | | 1 | | | | | | 0 |
| 10459 | G | A | A | upstream_gene_variant | intergenic | | | | 4 | | | | | | 0 |
| 10463 | G | A | A | upstream_gene_variant | intergenic | | | | 32 | | | | | | 0 |
| 10465 | G | A,T | A | upstream_gene_variant | intergenic | | | | 4 | T | upstream_gene_variant | intergenic | c.-157G>T | | 20 |
| 10471 | G | A | A | upstream_gene_variant | intergenic | | | | 1225 | | | | | | 0 |
| 10473 | A | G | G | upstream_gene_variant | intergenic | | | | 1218 | | | | | | 0 |
| 10487 | G | T | T | upstream_gene_variant | intergenic | | | | 2 | | | | | | 0 |
| 10498 | C | T | T | upstream_gene_variant | intergenic | | | | 5 | | | | | | 0 |
| 10500 | A | G | G | upstream_gene_variant | intergenic | | | | 23 | | | | | | 0 |
| 10504 | A | G | G | upstream_gene_variant | intergenic | | | | 1 | | | | | | 0 |
| 10514 | G | A | A | upstream_gene_variant | intergenic | | | | 6 | | | | | | 0 |
| 10518 | G | A | A | upstream_gene_variant | intergenic | | | | 384 | | | | | | 0 |
| 10525 | A | G | G | upstream_gene_variant | intergenic | | | | 348 | | | | | | 0 |
| 10526 | C | T | T | upstream_gene_variant | intergenic | | | | 4 | | | | | | 0 |
| 10528 | A | C | C | upstream_gene_variant | intergenic | | | | 4 | | | | | | 0 |
| 10529 | G | A | A | upstream_gene_variant | intergenic | | | | 33 | | | | | | 0 |
| 10530 | G | A | A | upstream_gene_variant | intergenic | | | | 7 | | | | | | 0 |
| 10539 | A | T | T | upstream_gene_variant | intergenic | | | | 24 | | | | | | 0 |
| 10540 | T | C | C | upstream_gene_variant | intergenic | | | | 33 | | | | | | 0 |
| 10542 | A | G | G | upstream_gene_variant | intergenic | | | | 1 | | | | | | 0 |
| 10551 | A | G | G | upstream_gene_variant | intergenic | | | | 71 | | | | | | 0 |
| 10554 | G | A | A | upstream_gene_variant | intergenic | | | | 2 | | | | | | 0 |
| 10562 | C | T,A | A | upstream_gene_variant | intergenic | | | | 6 | T | upstream_gene_variant | intergenic | c.-60C>T | | 3 |
| 10563 | G | A | A | upstream_gene_variant | intergenic | | | | 1 | | | | | | 0 |
| 10564 | T | C | C | upstream_gene_variant | intergenic | | | | 11 | | | | | | 0 |
| 10570 | A | G | G | upstream_gene_variant | intergenic | | | | 2 | | | | | | 0 |
| 10583 | C | T,A | A | upstream_gene_variant | intergenic | | | | 1 | T | upstream_gene_variant | intergenic | c.-39C>T | | 37 |
| 10601 | A | C | C | upstream_gene_variant | intergenic | | | | 44 | | | | | | 0 |
| 10663 | T | G,C | G | missense_variant | gacI | M5005_Spy0610 | c.42T>G | p.Ser14Arg | 33 | C | synonymous_variant | gacI | c.42T>C | p.Ser14Ser | 5 |
| 10666 | T | C | C | synonymous_variant | gacI | M5005_Spy0610 | c.45T>C | p.Asn15Asn | 4 | | | | | | 0 |
| 10670 | G | A | A | missense_variant | gacI | M5005_Spy0610 | c.49G>A | p.Val17Ile | 12 | | | | | | 0 |
| 10682 | C | A,T | A | missense_variant | gacI | M5005_Spy0610 | c.61C>A | p.Arg21Ser | 3 | T | missense_variant | gacI | c.61C>T | p.Arg21Cys | 1 |
| 10695 | C | T | T | missense_variant | gacI | M5005_Spy0610 | c.74C>T | p.Ser25Leu | 15 | | | | | | 0 |
| 10702 | C | T | T | synonymous_variant | gacI | M5005_Spy0610 | c.81C>T | p.Ala27Ala | 14 | | | | | | 0 |
| 10703 | C | T | T | missense_variant | gacI | M5005_Spy0610 | c.82C>T | p.Pro28Ser | 109 | | | | | | 0 |
| 10741 | G | A | A | synonymous_variant | gacI | M5005_Spy0610 | c.120G>A | p.Thr40Thr | 6 | | | | | | 0 |
| 10750 | G | A | A | synonymous_variant | gacI | M5005_Spy0610 | c.129G>A | p.Thr43Thr | 25 | | | | | | 0 |
| 10754 | G | A | A | missense_variant | gacI | M5005_Spy0610 | c.133G>A | p.Ala45Thr | 16 | | | | | | 0 |
| 10765 | A | G | G | synonymous_variant | gacI | M5005_Spy0610 | c.144A>G | p.Gln48Gln | 3 | | | | | | 0 |
| 10766 | A | G | G | missense_variant | gacI | M5005_Spy0610 | c.145A>G | p.Lys49Glu | 144 | | | | | | 0 |
| 10768 | A | C | C | missense_variant | gacI | M5005_Spy0610 | c.147A>C | p.Lys49Asn | 26 | | | | | | 0 |
| 10777 | C | T,G | G | missense_variant | gacI | M5005_Spy0610 | c.156C>G | p.Phe52Leu | 24 | T | synonymous_variant | gacI | c.156C>T | p.Phe52Phe | 3 |
| 10783 | T | G | G | synonymous_variant | gacI | M5005_Spy0610 | c.162T>G | p.Val54Val | 29 | | | | | | 0 |
| 10796 | A | G | G | missense_variant | gacI | M5005_Spy0610 | c.175A>G | p.Ile59Val | 1 | | | | | | 0 |
| 10802 | C | T | T | synonymous_variant | gacI | M5005_Spy0610 | c.181C>T | p.Leu61Leu | 106 | | | | | | 0 |
| 10821 | T | A | A | missense_variant | gacI | M5005_Spy0610 | c.200T>A | p.Val67Glu | 1 | | | | | | 0 |
| 10836 | G | A | A | missense_variant | gacI | M5005_Spy0610 | c.215G>A | p.Arg72His | 49 | | | | | | 0 |
| 10848 | G | T | T | missense_variant | gacI | M5005_Spy0610 | c.227G>T | p.Arg76Ile | 3 | | | | | | 0 |

| | | | | | | | | | | | | | | | |
|---|---|---|---|---|---|---|---|---|---|---|---|---|---|---|---|
| 10861 | C | T | T | synonymous_variant | gacI | M5005_Spy0610 | c.240C>T | p.Asp80Asp | 28 | | | | | | 0 |
| 10868 | G | A | A | missense_variant | gacI | M5005_Spy0610 | c.247G>A | p.Val83Ile | 1 | | | | | | 0 |
| 10891 | G | A | A | synonymous_variant | gacI | M5005_Spy0610 | c.270G>A | p.Gln90Gln | 33 | | | | | | 0 |
| 10903 | C | T | T | synonymous_variant | gacI | M5005_Spy0610 | c.282C>T | p.Cys94Cys | 322 | | | | | | 0 |
| 10911 | A | G,T | G | missense_variant | gacI | M5005_Spy0610 | c.290A>G | p.Glu97Gly | 40 | T | missense_variant | gacI | c.290A>T | p.Glu97Val | 3 |
| 10912 | A | G | G | synonymous_variant | gacI | M5005_Spy0610 | c.291A>G | p.Glu97Glu | 1 | | | | | | 0 |
| 10914 | A | C | C | missense_variant | gacI | M5005_Spy0610 | c.293A>C | p.Lys98Thr | 10 | | | | | | 0 |
| 10924 | G | A | A | synonymous_variant | gacI | M5005_Spy0610 | c.303G>A | p.Glu101Glu | 55 | | | | | | 0 |
| 10966 | A | T,G | G | synonymous_variant | gacI | M5005_Spy0610 | c.345A>G | p.Arg115Arg | 63 | T | synonymous_variant | gacI | c.345A>T | p.Arg115Arg | 1 |
| 10978 | A | G | G | synonymous_variant | gacI | M5005_Spy0610 | c.357A>G | p.Lys119Lys | 23 | | | | | | 0 |
| 10993 | A | G | G | synonymous_variant | gacI | M5005_Spy0610 | c.372A>G | p.Ser124Ser | 3 | | | | | | 0 |
| 11008 | T | C | C | synonymous_variant | gacI | M5005_Spy0610 | c.387T>C | p.Arg129Arg | 1 | | | | | | 0 |
| 11011 | C | T | T | synonymous_variant | gacI | M5005_Spy0610 | c.390C>T | p.Ile130Ile | 1 | | | | | | 0 |
| 11014 | C | T | T | synonymous_variant | gacI | M5005_Spy0610 | c.393C>T | p.Gly131Gly | 716 | | | | | | 0 |
| 11027 | A | G | G | missense_variant | gacI | M5005_Spy0610 | c.406A>G | p.Thr136Ala | 21 | | | | | | 0 |
| 11029 | T | A,* | A | synonymous_variant | gacI | M5005_Spy0610 | c.408T>A | p.Thr136Thr | 125 | | | | | | 1 |
| 11041 | G | A | A | synonymous_variant | gacI | M5005_Spy0610 | c.420G>A | p.Ala140Ala | 1 | | | | | | 0 |
| 11052 | G | A | A | missense_variant | gacI | M5005_Spy0610 | c.431G>A | p.Gly144Glu | 120 | | | | | | 0 |
| 11074 | G | T | T | synonymous_variant | gacI | M5005_Spy0610 | c.453G>T | p.Thr151Thr | 1 | | | | | | 0 |
| 11080 | T | G | G | synonymous_variant | gacI | M5005_Spy0610 | c.459T>G | p.Gly153Gly | 1 | | | | | | 0 |
| 11081 | C | T | T | synonymous_variant | gacI | M5005_Spy0610 | c.460C>T | p.Leu154Leu | 8 | | | | | | 0 |
| 11086 | G | A | A | synonymous_variant | gacI | M5005_Spy0610 | c.465G>A | p.Arg155Arg | 18 | | | | | | 0 |
| 11091 | T | C | C | missense_variant | gacI | M5005_Spy0610 | c.470T>C | p.Ile157Thr | 11 | | | | | | 0 |
| 11095 | C | T | T | synonymous_variant | gacI | M5005_Spy0610 | c.474C>T | p.Asp158Asp | 29 | | | | | | 0 |
| 11100 | C | T | T | missense_variant | gacI | M5005_Spy0610 | c.479C>T | p.Ser160Leu | 1 | | | | | | 0 |
| 11110 | A | C | C | missense_variant | gacI | M5005_Spy0610 | c.489A>C | p.Glu163Asp | 1 | | | | | | 0 |
| 11123 | C | T | T | missense_variant | gacI | M5005_Spy0610 | c.502C>T | p.His168Tyr | 29 | | | | | | 0 |
| 11125 | T | G | G | missense_variant | gacI | M5005_Spy0610 | c.504T>G | p.His168Gln | 2 | | | | | | 0 |
| 11129 | C | T | T | missense_variant | gacI | M5005_Spy0610 | c.508C>T | p.Pro170Ser | 20 | | | | | | 0 |
| 11161 | T | C | C | synonymous_variant | gacI | M5005_Spy0610 | c.540T>C | p.Val180Val | 1 | | | | | | 0 |
| 11165 | G | A | A | missense_variant | gacI | M5005_Spy0610 | c.544G>A | p.Val182Ile | 1 | | | | | | 0 |
| 11175 | G | A | A | missense_variant | gacI | M5005_Spy0610 | c.554G>A | p.Ser185Asn | 17 | | | | | | 0 |
| 11196 | T | C | C | missense_variant | gacI | M5005_Spy0610 | c.575T>C | p.Ile192Thr | 38 | | | | | | 0 |
| 11224 | C | T,G | G | synonymous_variant | gacI | M5005_Spy0610 | c.603C>G | p.Gly201Gly | 25 | T | synonymous_variant | gacI | c.603C>T | p.Gly201Gly | 1 |
| 11233 | A | G | G | synonymous_variant | gacI | M5005_Spy0610 | c.612A>G | p.Ser204Ser | 38 | | | | | | 0 |
| 11242 | G | A | A | synonymous_variant | gacI | M5005_Spy0610 | c.621G>A | p.Ser207Ser | 261 | | | | | | 0 |
| 11247 | C | T | T | missense_variant | gacI | M5005_Spy0610 | c.626C>T | p.Thr209Met | 17 | | | | | | 0 |
| 11272 | A | G | G | synonymous_variant | gacI | M5005_Spy0610 | c.651A>G | p.Lys217Lys | 2 | | | | | | 0 |
| 11294 | G | T | T | missense_variant | gacI | M5005_Spy0610 | c.673G>T | p.Val225Phe | 3 | | | | | | 0 |
| 11298 | G | A | A | missense_variant | gacI | M5005_Spy0610 | c.677G>A | p.Arg226Lys | 2 | | | | | | 0 |
| 11299 | A | G | G | synonymous_variant | gacI | M5005_Spy0610 | c.678A>G | p.Arg226Arg | 90 | | | | | | 0 |
| 11310 | A | G | G | missense_variant | gacI | M5005_Spy0610 | c.689A>G | p.Asn230Ser | 2 | | | | | | 0 |
| 11311 | T | A | A | missense_variant | gacI | M5005_Spy0610 | c.690T>A | p.Asn230Lys | 2 | | | | | | 0 |
| 11360 | A | C | C | missense_variant | gacJ | M5005_Spy0611 | c.42A>C | p.Leu14Phe | 1 | | | | | | 0 |
| 11362 | C | T | T | missense_variant | gacJ | M5005_Spy0611 | c.44C>T | p.Thr15Ile | 6 | | | | | | 0 |
| 11367 | G | A | A | missense_variant | gacJ | M5005_Spy0611 | c.49G>A | p.Val17Ile | 17 | | | | | | 0 |
| 11387 | A | G | G | synonymous_variant | gacJ | M5005_Spy0611 | c.69A>G | p.Leu23Leu | 9 | | | | | | 0 |
| 11396 | A | G | G | synonymous_variant | gacJ | M5005_Spy0611 | c.78A>G | p.Lys26Lys | 1 | | | | | | 0 |
| 11399 | T | C | C | synonymous_variant | gacJ | M5005_Spy0611 | c.81T>C | p.Asp27Asp | 1778 | | | | | | 0 |
| 11426 | G | A | A | synonymous_variant | gacJ | M5005_Spy0611 | c.108G>A | p.Leu36Leu | 125 | | | | | | 0 |
| 11453 | A | G | G | synonymous_variant | gacJ | M5005_Spy0611 | c.135A>G | p.Leu45Leu | 22 | | | | | | 0 |
| 11469 | C | T | T | missense_variant | gacJ | M5005_Spy0611 | c.151C>T | p.Pro51Ser | 6 | | | | | | 0 |
| 11472 | A | G | G | missense_variant | gacJ | M5005_Spy0611 | c.154A>G | p.Asn52Asp | 20 | | | | | | 0 |
| 11478 | T | C | C | synonymous_variant | gacJ | M5005_Spy0611 | c.160T>C | p.Leu54Leu | 1 | | | | | | 0 |
| 11480 | A | G | G | synonymous_variant | gacJ | M5005_Spy0611 | c.162A>G | p.Leu54Leu | 2 | | | | | | 0 |
| 11482 | A | G | G | missense_variant | gacJ | M5005_Spy0611 | c.164A>G | p.Asn55Ser | 16 | | | | | | 0 |
| 11483 | T | A | A | missense_variant | gacJ | M5005_Spy0611 | c.165T>A | p.Asn55Lys | 2 | | | | | | 0 |
| 11484 | A | G | G | missense_variant | gacJ | M5005_Spy0611 | c.166A>G | p.Lys56Glu | 14 | | | | | | 0 |
| 11487 | A | T | T | missense_variant | gacJ | M5005_Spy0611 | c.169A>T | p.Met57Leu | 6 | | | | | | 0 |
| 11498 | C | T | T | synonymous_variant | gacJ | M5005_Spy0611 | c.180C>T | p.Phe60Phe | 115 | | | | | | 0 |
| 11507 | T | A,C | A | synonymous_variant | gacJ | M5005_Spy0611 | c.189T>A | p.Ile63Ile | 3 | C | synonymous_variant | gacJ | c.189T>C | p.Ile63Ile | 17 |
| 11514 | C | T | T | missense_variant | gacJ | M5005_Spy0611 | c.196C>T | p.Pro66Ser | 4 | | | | | | 0 |

| | | | | | | | | | | | | | | | |
|---|---|---|---|---|---|---|---|---|---|---|---|---|---|---|---|
| 11526 | T | C | C | synonymous_variant | gacJ | M5005_Spy0611 | c.208T>C | p.Leu70Leu | 92 | | | | | | 0 |
| 11552 | T | C | C | synonymous_variant | gacJ | M5005_Spy0611 | c.234T>C | p.Ser78Ser | 1 | | | | | | 0 |
| 11567 | T | C | C | synonymous_variant | gacJ | M5005_Spy0611 | c.249T>C | p.Phe83Phe | 1 | | | | | | 0 |
| 11571 | C | T | T | synonymous_variant | gacJ | M5005_Spy0611 | c.253C>T | p.Leu85Leu | 1 | | | | | | 0 |
| 11585 | T | C | C | synonymous_variant | gacJ | M5005_Spy0611 | c.267T>C | p.Ile89Ile | 727 | | | | | | 0 |
| 11610 | C | T | T | missense_variant | gacJ | M5005_Spy0611 | c.292C>T | p.Arg98Cys | 10 | | | | | | 0 |
| 11618 | C | T | T | synonymous_variant | gacJ | M5005_Spy0611 | c.300C>T | p.Thr100Thr | 2 | | | | | | 0 |
| 11624 | A | G | G | synonymous_variant | gacJ | M5005_Spy0611 | c.306A>G | p.Glu102Glu | 39 | | | | | | 0 |
| 11636 | G | T | T | missense_variant | gacJ | M5005_Spy0611 | c.318G>T | p.Met106Ile | 3 | | | | | | 0 |
| 11650 | C | A | A | missense_variant | gacJ | M5005_Spy0611 | c.332C>A | p.Thr111Asn | 1 | | | | | | 0 |
| 11661 | T | G | G | missense_variant | gacK | M5005_Spy0612 | c.9T>G | p.Asn3Lys | 4 | | | | | | 0 |
| 11663 | C | T | T | missense_variant | gacK | M5005_Spy0612 | c.11C>T | p.Pro4Leu | 4 | | | | | | 0 |
| 11664 | T | C | C | synonymous_variant | gacK | M5005_Spy0612 | c.12T>C | p.Pro4Pro | 30 | | | | | | 0 |
| 11666 | C | T | T | missense_variant | gacK | M5005_Spy0612 | c.14C>T | p.Ser5Leu | 51 | | | | | | 0 |
| 11681 | C | T | T | missense_variant | gacK | M5005_Spy0612 | c.29C>T | p.Thr10Met | 1 | | | | | | 0 |
| 11682 | G | A | A | synonymous_variant | gacK | M5005_Spy0612 | c.30G>A | p.Thr10Thr | 2 | | | | | | 0 |
| 11691 | A | G | G | synonymous_variant | gacK | M5005_Spy0612 | c.39A>G | p.Leu13Leu | 3 | | | | | | 0 |
| 11699 | T | C | C | missense_variant | gacK | M5005_Spy0612 | c.47T>C | p.Met16Thr | 33 | | | | | | 0 |
| 11711 | T | C | C | missense_variant | gacK | M5005_Spy0612 | c.59T>C | p.Leu20Ser | 6 | | | | | | 0 |
| 11721 | T | C | C | synonymous_variant | gacK | M5005_Spy0612 | c.69T>C | p.Ala23Ala | 1 | | | | | | 0 |
| 11767 | T | C | C | missense_variant | gacK | M5005_Spy0612 | c.115T>C | p.Ser39Pro | 22 | | | | | | 0 |
| 11769 | G | T | T | synonymous_variant | gacK | M5005_Spy0612 | c.117G>T | p.Ser39Ser | 1 | | | | | | 0 |
| 11770 | G | T | T | missense_variant | gacK | M5005_Spy0612 | c.118G>T | p.Ala40Ser | 181 | | | | | | 0 |
| 11790 | C | T | T | synonymous_variant | gacK | M5005_Spy0612 | c.138C>T | p.Ala46Ala | 30 | | | | | | 0 |
| 11863 | A | G | G | missense_variant | gacK | M5005_Spy0612 | c.211A>G | p.Ile71Val | 1 | | | | | | 0 |
| 11865 | T | G | G | missense_variant | gacK | M5005_Spy0612 | c.213T>G | p.Ile71Met | 5 | | | | | | 0 |
| 11886 | T | C | C | synonymous_variant | gacK | M5005_Spy0612 | c.234T>C | p.Ser78Ser | 1 | | | | | | 0 |
| 11889 | G | A | A | synonymous_variant | gacK | M5005_Spy0612 | c.237G>A | p.Gln79Gln | 12 | | | | | | 0 |
| 11898 | G | A | A | synonymous_variant | gacK | M5005_Spy0612 | c.246G>A | p.Val82Val | 107 | | | | | | 0 |
| 11899 | G | A | A | missense_variant | gacK | M5005_Spy0612 | c.247G>A | p.Ala83Thr | 14 | | | | | | 0 |
| 11901 | A | G | G | synonymous_variant | gacK | M5005_Spy0612 | c.249A>G | p.Ala83Ala | 16 | | | | | | 0 |
| 11916 | T | C | C | synonymous_variant | gacK | M5005_Spy0612 | c.264T>C | p.Cys88Cys | 1 | | | | | | 0 |
| 11931 | C | T | T | synonymous_variant | gacK | M5005_Spy0612 | c.279C>T | p.Ala93Ala | 67 | | | | | | 0 |
| 11934 | A | G | G | missense_variant | gacK | M5005_Spy0612 | c.282A>G | p.Ile94Met | 3 | | | | | | 0 |
| 11950 | A | G | G | missense_variant | gacK | M5005_Spy0612 | c.298A>G | p.Thr100Ala | 1 | | | | | | 0 |
| 11984 | T | C | C | missense_variant | gacK | M5005_Spy0612 | c.332T>C | p.Ile111Thr | 22 | | | | | | 0 |
| 11992 | C | T | T | synonymous_variant | gacK | M5005_Spy0612 | c.340C>T | p.Leu114Leu | 1 | | | | | | 0 |
| 12008 | G | A | A | missense_variant | gacK | M5005_Spy0612 | c.356G>A | p.Arg119Lys | 1 | | | | | | 0 |
| 12039 | G | A | A | synonymous_variant | gacK | M5005_Spy0612 | c.387G>A | p.Gln129Gln | 1 | | | | | | 0 |
| 12048 | C | T | T | synonymous_variant | gacK | M5005_Spy0612 | c.396C>T | p.Phe132Phe | 1 | | | | | | 0 |
| 12055 | C | T | T | missense_variant | gacK | M5005_Spy0612 | c.403C>T | p.His135Tyr | 2 | | | | | | 0 |
| 12063 | G | A | A | synonymous_variant | gacK | M5005_Spy0612 | c.411G>A | p.Arg137Arg | 88 | | | | | | 0 |
| 12075 | A | G | G | synonymous_variant | gacK | M5005_Spy0612 | c.423A>G | p.Ala141Ala | 1 | | | | | | 0 |
| 12090 | C | T | T | synonymous_variant | gacK | M5005_Spy0612 | c.438C>T | p.Ala146Ala | 93 | | | | | | 0 |
| 12100 | A | G | G | missense_variant | gacK | M5005_Spy0612 | c.448A>G | p.Thr150Ala | 8 | | | | | | 0 |
| 12102 | T | C | C | synonymous_variant | gacK | M5005_Spy0612 | c.450T>C | p.Thr150Thr | 3 | | | | | | 0 |
| 12114 | G | A | A | missense_variant | gacK | M5005_Spy0612 | c.462G>A | p.Met154Ile | 10 | | | | | | 0 |
| 12115 | G | A | A | missense_variant | gacK | M5005_Spy0612 | c.463G>A | p.Val155Met | 14 | | | | | | 0 |
| 12117 | G | T | T | synonymous_variant | gacK | M5005_Spy0612 | c.465G>T | p.Val155Val | 2 | | | | | | 0 |
| 12130 | A | T | T | missense_variant | gacK | M5005_Spy0612 | c.478A>T | p.Ile160Leu | 5 | | | | | | 0 |
| 12133 | T | C | C | synonymous_variant | gacK | M5005_Spy0612 | c.481T>C | p.Leu161Leu | 1 | | | | | | 0 |
| 12135 | A | G | G | synonymous_variant | gacK | M5005_Spy0612 | c.483A>G | p.Leu161Leu | 1 | | | | | | 0 |
| 12140 | C | A | A | missense_variant | gacK | M5005_Spy0612 | c.488C>A | p.Ser163Tyr | 17 | | | | | | 0 |
| 12144 | A | G | G | synonymous_variant | gacK | M5005_Spy0612 | c.492A>G | p.Lys164Lys | 10 | | | | | | 0 |
| 12150 | T | C | C | synonymous_variant | gacK | M5005_Spy0612 | c.498T>C | p.Leu166Leu | 6 | | | | | | 0 |
| 12164 | T | C | C | missense_variant | gacK | M5005_Spy0612 | c.512T>C | p.Val171Ala | 1 | | | | | | 0 |
| 12169 | G | A | A | missense_variant | gacK | M5005_Spy0612 | c.517G>A | p.Val173Ile | 2 | | | | | | 0 |
| 12170 | T | C | C | missense_variant | gacK | M5005_Spy0612 | c.518T>C | p.Val173Ala | 27 | | | | | | 0 |
| 12172 | T | G | G | missense_variant | gacK | M5005_Spy0612 | c.520T>G | p.Cys174Gly | 4 | | | | | | 0 |
| 12197 | T | C | C | missense_variant | gacK | M5005_Spy0612 | c.545T>C | p.Met182Thr | 4 | | | | | | 0 |
| 12207 | C | T | T | synonymous_variant | gacK | M5005_Spy0612 | c.555C>T | p.Asp185Asp | 4 | | | | | | 0 |
| 12213 | T | C | C | synonymous_variant | gacK | M5005_Spy0612 | c.561T>C | p.Gly187Gly | 85 | | | | | | 0 |

| | | | | | | | | | | | | | | | |
|---|---|---|---|---|---|---|---|---|---|---|---|---|---|---|---|
| 12214 | C | T | T | missense_variant | gacK | M5005_Spy0612 | c.562C>T | p.His188Tyr | 203 | | | | | | 0 |
| 12244 | A | G | G | missense_variant | gacK | M5005_Spy0612 | c.592A>G | p.Ser198Gly | 2 | | | | | | 0 |
| 12257 | G | A | A | missense_variant | gacK | M5005_Spy0612 | c.605G>A | p.Ser202Asn | 6 | | | | | | 0 |
| 12259 | A | G | G | missense_variant | gacK | M5005_Spy0612 | c.607A>G | p.Asn203Asp | 6 | | | | | | 0 |
| 12270 | T | C | C | synonymous_variant | gacK | M5005_Spy0612 | c.618T>C | p.Phe206Phe | 110 | | | | | | 0 |
| 12306 | A | G | G | synonymous_variant | gacK | M5005_Spy0612 | c.654A>G | p.Pro218Pro | 58 | | | | | | 0 |
| 12311 | T | C | C | missense_variant | gacK | M5005_Spy0612 | c.659T>C | p.Phe220Ser | 1 | | | | | | 0 |
| 12359 | C | T | T | missense_variant | gacK | M5005_Spy0612 | c.707C>T | p.Ala236Val | 3 | | | | | | 0 |
| 12367 | C | G | G | missense_variant | gacK | M5005_Spy0612 | c.715C>G | p.Leu239Val | 1 | | | | | | 0 |
| 12374 | C | T | T | missense_variant | gacK | M5005_Spy0612 | c.722C>T | p.Thr241Met | 50 | | | | | | 0 |
| 12377 | C | T | T | missense_variant | gacK | M5005_Spy0612 | c.725C>T | p.Thr242Ile | 2 | | | | | | 0 |
| 12383 | G | A | A | missense_variant | gacK | M5005_Spy0612 | c.731G>A | p.Gly244Asp | 1 | | | | | | 0 |
| 12388 | G | A | A | missense_variant | gacK | M5005_Spy0612 | c.736G>A | p.Val246Ile | 501 | | | | | | 0 |
| 12392 | C | T | T | missense_variant | gacK | M5005_Spy0612 | c.740C>T | p.Ala247Val | 9 | | | | | | 0 |
| 12396 | G | A | A | synonymous_variant | gacK | M5005_Spy0612 | c.744G>A | p.Leu248Leu | 3 | | | | | | 0 |
| 12399 | C | T | T | synonymous_variant | gacK | M5005_Spy0612 | c.747C>T | p.Gly249Gly | 4 | | | | | | 0 |
| 12435 | C | A | A | synonymous_variant | gacK | M5005_Spy0612 | c.783C>A | p.Ala261Ala | 2 | | | | | | 0 |
| 12450 | G | A | A | synonymous_variant | gacK | M5005_Spy0612 | c.798G>A | p.Leu266Leu | 1 | | | | | | 0 |
| 12453 | A | C | C | synonymous_variant | gacK | M5005_Spy0612 | c.801A>C | p.Leu267Leu | 22 | | | | | | 0 |
| 12472 | C | T | T | missense_variant | gacK | M5005_Spy0612 | c.820C>T | p.His274Tyr | 782 | | | | | | 0 |
| 12494 | C | T | T | missense_variant | gacK | M5005_Spy0612 | c.842C>T | p.Ala281Val | 4 | | | | | | 0 |
| 12501 | C | T | T | synonymous_variant | gacK | M5005_Spy0612 | c.849C>T | p.Ile283Ile | 2 | | | | | | 0 |
| 12511 | A | G | G | missense_variant | gacK | M5005_Spy0612 | c.859A>G | p.Ile287Val | 5 | | | | | | 0 |
| 12547 | G | A | A | missense_variant | gacK | M5005_Spy0612 | c.895G>A | p.Ala299Thr | 5 | | | | | | 0 |
| 12570 | G | A | A | synonymous_variant | gacK | M5005_Spy0612 | c.918G>A | p.Leu306Leu | 1 | | | | | | 0 |
| 12575 | C | T | T | missense_variant | gacK | M5005_Spy0612 | c.923C>T | p.Ala308Val | 64 | | | | | | 0 |
| 12592 | T | C | C | synonymous_variant | gacK | M5005_Spy0612 | c.940T>C | p.Leu314Leu | 5 | | | | | | 0 |
| 12604 | C | A | A | missense_variant | gacK | M5005_Spy0612 | c.952C>A | p.Pro318Thr | 7 | | | | | | 0 |
| 12605 | C | T | T | missense_variant | gacK | M5005_Spy0612 | c.953C>T | p.Pro318Leu | 2 | | | | | | 0 |
| 12618 | A | G | G | missense_variant | gacK | M5005_Spy0612 | c.966A>G | p.Ile322Met | 1188 | | | | | | 0 |
| 12619 | C | T | T | synonymous_variant | gacK | M5005_Spy0612 | c.967C>T | p.Leu323Leu | 4 | | | | | | 0 |
| 12627 | T | C | C | synonymous_variant | gacK | M5005_Spy0612 | c.975T>C | p.Gly325Gly | 34 | | | | | | 0 |
| 12633 | C | A | A | missense_variant | gacK | M5005_Spy0612 | c.981C>A | p.Asn327Lys | 2 | | | | | | 0 |
| 12639 | G | A | A | synonymous_variant | gacK | M5005_Spy0612 | c.987G>A | p.Thr329Thr | 14 | | | | | | 0 |
| 12654 | T | C | C | synonymous_variant | gacK | M5005_Spy0612 | c.1002T>C | p.Asp334Asp | 1287 | | | | | | 0 |
| 12670 | C | T | T | synonymous_variant | gacK | M5005_Spy0612 | c.1018C>T | p.Leu340Leu | 4 | | | | | | 0 |
| 12681 | G | A | A | synonymous_variant | gacK | M5005_Spy0612 | c.1029G>A | p.Ser343Ser | 5 | | | | | | 0 |
| 12689 | G | T | T | missense_variant | gacK | M5005_Spy0612 | c.1037G>T | p.Ser346Ile | 1 | | | | | | 0 |
| 12695 | C | T | T | missense_variant | gacK | M5005_Spy0612 | c.1043C>T | p.Ala348Val | 39 | | | | | | 0 |
| 12719 | C | T | T | missense_variant | gacK | M5005_Spy0612 | c.1067C>T | p.Thr356Ile | 1 | | | | | | 0 |
| 12730 | A | C | C | missense_variant | gacK | M5005_Spy0612 | c.1078A>C | p.Lys360Gln | 22 | | | | | | 0 |
| 12731 | A | C | C | missense_variant | gacK | M5005_Spy0612 | c.1079A>C | p.Lys360Thr | 33 | | | | | | 0 |
| 12745 | C | A | A | missense_variant | gacK | M5005_Spy0612 | c.1093C>A | p.Leu365Ile | 2 | | | | | | 0 |
| 12753 | A | G | G | synonymous_variant | gacK | M5005_Spy0612 | c.1101A>G | p.Pro367Pro | 33 | | | | | | 0 |
| 12762 | A | G | G | synonymous_variant | gacK | M5005_Spy0612 | c.1110A>G | p.Gly370Gly | 9 | | | | | | 0 |
| 12765 | A | G | G | synonymous_variant | gacK | M5005_Spy0612 | c.1113A>G | p.Gly371Gly | 12 | | | | | | 0 |
| 12777 | G | A | A | synonymous_variant | gacK | M5005_Spy0612 | c.1125G>A | p.Ser375Ser | 4 | | | | | | 0 |
| 12784 | A | G | G | missense_variant | gacK | M5005_Spy0612 | c.1132A>G | p.Ile378Val | 5 | | | | | | 0 |
| 12786 | T | C | C | synonymous_variant | gacK | M5005_Spy0612 | c.1134T>C | p.Ile378Ile | 3 | | | | | | 0 |
| 12787 | A | G | G | missense_variant | gacK | M5005_Spy0612 | c.1135A>G | p.Thr379Ala | 6 | | | | | | 0 |
| 12793 | C | T | T | missense_variant | gacK | M5005_Spy0612 | c.1141C>T | p.Leu381Phe | 1 | | | | | | 0 |
| 12803 | T | C | C | missense_variant | gacK | M5005_Spy0612 | c.1151T>C | p.Met384Thr | 33 | | | | | | 0 |
| 12809 | A | C | C | missense_variant | gacK | M5005_Spy0612 | c.1157A>C | p.Tyr386Ser | 1 | | | | | | 0 |
| 12813 | T | C | C | synonymous_variant | gacK | M5005_Spy0612 | c.1161T>C | p.His387His | 1445 | | | | | | 0 |
| 12823 | G | T | T | missense_variant | gacK | M5005_Spy0612 | c.1171G>T | p.Ala391Ser | 3 | | | | | | 0 |
| 12831 | G | A | A | synonymous_variant | gacK | M5005_Spy0612 | c.1179G>A | p.Leu393Leu | 1 | | | | | | 0 |
| 12859 | T | C | C | synonymous_variant | gacK | M5005_Spy0612 | c.1207T>C | p.Leu403Leu | 18 | | | | | | 0 |
| 12873 | T | C | C | synonymous_variant | gacK | M5005_Spy0612 | c.1221T>C | p.Ile407Ile | 17 | | | | | | 0 |
| 12909 | G | A | A | synonymous_variant | gacK | M5005_Spy0612 | c.1257G>A | p.Lys419Lys | 19 | | | | | | 0 |
| 12912 | A | G | G | synonymous_variant | gacK | M5005_Spy0612 | c.1260A>G | p.Gly420Gly | 143 | | | | | | 0 |
| 12927 | A | G | G | synonymous_variant | gacK_overlap | M5005_Spy0613 | | | 4 | | missense_variant | gacL_overlap | c.8A>G | p.Gln3Arg | 0 |
| 12930 | C | T | T | synonymous_variant | gacK_overlap | M5005_Spy0613 | | | 5 | | missense_variant | gacL_overlap | c.11C>T | p.Ser4Phe | 0 |

| | | | | | | | | | | | | | | | |
|---|---|---|---|---|---|---|---|---|---|---|---|---|---|---|---|
| 12932 | A | G | G | missense_variant | gacK_overlap | M5005_Spy0612 | c.1280A>G | p.Tyr427Cys | 9 | | missense_variant | gacL_overlap | | p.Met5Val | 0 |
| 12934 | G | A | A | missense_variant | gacK_overlap | M5005_Spy0612 | c.1282G>A | p.Asp428Asn | 5 | | missense_variant | gacL_overlap | | p.Met5Ile | 0 |
| 12938 | A | G | G | synonymous_variant | gacK_overlap | M5005_Spy0613 | | | 1 | | missense_variant | gacL_overlap | c.19A>G | p.Lys7Glu | 0 |
| 12950 | C | T | T | missense_variant | gacL | M5005_Spy0613 | c.31C>T | p.Leu11Phe | 4 | | | | | | 0 |
| 12991 | G | T | T | missense_variant | gacL | M5005_Spy0613 | c.72G>T | p.Leu24Phe | 20 | | | | | | 0 |
| 12998 | C | T | T | missense_variant | gacL | M5005_Spy0613 | c.79C>T | p.Leu27Phe | 581 | | | | | | 0 |
| 12999 | T | C | C | missense_variant | gacL | M5005_Spy0613 | c.80T>C | p.Leu27Pro | 5 | | | | | | 0 |
| 13001 | C | T,A | A | missense_variant | gacL | M5005_Spy0613 | c.82C>A | p.Leu28Met | 6 | T | synonymous_variant | gacL | c.82C>T | p.Leu28Leu | 17 |
| 13019 | G | A | A | missense_variant | gacL | M5005_Spy0613 | c.100G>A | p.Ala34Thr | 5 | | | | | | 0 |
| 13020 | C | T | T | missense_variant | gacL | M5005_Spy0613 | c.101C>T | p.Ala34Val | 7 | | | | | | 0 |
| 13021 | C | T | T | synonymous_variant | gacL | M5005_Spy0613 | c.102C>T | p.Ala34Ala | 7 | | | | | | 0 |
| 13032 | A | G | G | missense_variant | gacL | M5005_Spy0613 | c.113A>G | p.Asn38Ser | 11 | | | | | | 0 |
| 13036 | A | G | G | synonymous_variant | gacL | M5005_Spy0613 | c.117A>G | p.Leu39Leu | 6 | | | | | | 0 |
| 13067 | G | A | A | missense_variant | gacL | M5005_Spy0613 | c.148G>A | p.Val50Met | 22 | | | | | | 0 |
| 13072 | C | T | T | synonymous_variant | gacL | M5005_Spy0613 | c.153C>T | p.Ser51Ser | 138 | | | | | | 0 |
| 13084 | A | G | G | missense_variant | gacL | M5005_Spy0613 | c.165A>G | p.Ile55Met | 3 | | | | | | 0 |
| 13108 | G | A | A | synonymous_variant | gacL | M5005_Spy0613 | c.189G>A | p.Glu63Glu | 6 | | | | | | 0 |
| 13117 | G | A | A | synonymous_variant | gacL | M5005_Spy0613 | c.198G>A | p.Thr66Thr | 18 | | | | | | 0 |
| 13118 | A | C | C | missense_variant | gacL | M5005_Spy0613 | c.199A>C | p.Ile67Leu | 10 | | | | | | 0 |
| 13127 | A | C | C | missense_variant | gacL | M5005_Spy0613 | c.208A>C | p.Ile70Leu | 3 | | | | | | 0 |
| 13133 | G | A | A | missense_variant | gacL | M5005_Spy0613 | c.214G>A | p.Val72Ile | 6 | | | | | | 0 |
| 13134 | T | C | C | missense_variant | gacL | M5005_Spy0613 | c.215T>C | p.Val72Ala | 127 | | | | | | 0 |
| 13141 | A | G | G | synonymous_variant | gacL | M5005_Spy0613 | c.222A>G | p.Gly74Gly | 10 | | | | | | 0 |
| 13143 | C | T | T | missense_variant | gacL | M5005_Spy0613 | c.224C>T | p.Thr75Met | 1 | | | | | | 0 |
| 13145 | A | C,G | C | missense_variant | gacL | M5005_Spy0613 | c.226A>C | p.Ile76Leu | 403 | G | missense_variant | gacL | c.226A>G | p.Ile76Val | 19 |
| 13157 | A | G | G | missense_variant | gacL | M5005_Spy0613 | c.238A>G | p.Ile80Val | 9 | | | | | | 0 |
| 13165 | T | C | C | synonymous_variant | gacL | M5005_Spy0613 | c.246T>C | p.Pro82Pro | 88 | | | | | | 0 |
| 13180 | T | G | G | synonymous_variant | gacL | M5005_Spy0613 | c.261T>G | p.Pro87Pro | 6 | | | | | | 0 |
| 13189 | T | C | C | synonymous_variant | gacL | M5005_Spy0613 | c.270T>C | p.Tyr90Tyr | 1 | | | | | | 0 |
| 13249 | C | T | T | synonymous_variant | gacL | M5005_Spy0613 | c.330C>T | p.Asn110Asn | 3 | | | | | | 0 |
| 13282 | G | A | A | synonymous_variant | gacL | M5005_Spy0613 | c.363G>A | p.Lys121Lys | 26 | | | | | | 0 |
| 13288 | T | C | C | synonymous_variant | gacL | M5005_Spy0613 | c.369T>C | p.Ile123Ile | 80 | | | | | | 0 |
| 13309 | T | C | C | synonymous_variant | gacL | M5005_Spy0613 | c.390T>C | p.Phe130Phe | 4 | | | | | | 0 |
| 13310 | A | C | C | missense_variant | gacL | M5005_Spy0613 | c.391A>C | p.Ile131Leu | 5 | | | | | | 0 |
| 13319 | A | G | G | missense_variant | gacL | M5005_Spy0613 | c.400A>G | p.Asn134Asp | 1 | | | | | | 0 |
| 13327 | A | T | T | missense_variant | gacL | M5005_Spy0613 | c.408A>T | p.Lys136Asn | 259 | | | | | | 0 |
| 13329 | C | T | T | missense_variant | gacL | M5005_Spy0613 | c.410C>T | p.Ala137Val | 30 | | | | | | 0 |
| 13342 | C | T | T | synonymous_variant | gacL | M5005_Spy0613 | c.423C>T | p.Ser141Ser | 31 | | | | | | 0 |
| 13346 | A | G | G | missense_variant | gacL | M5005_Spy0613 | c.427A>G | p.Arg143Gly | 5 | | | | | | 0 |
| 13351 | G | A | A | synonymous_variant | gacL | M5005_Spy0613 | c.432G>A | p.Glu144Glu | 2 | | | | | | 0 |
| 13396 | T | C | C | synonymous_variant | gacL | M5005_Spy0613 | c.477T>C | p.Phe159Phe | 4 | | | | | | 0 |
| 13405 | A | G | G | missense_variant | gacL | M5005_Spy0613 | c.486A>G | p.Ile162Met | 9 | | | | | | 0 |
| 13436 | G | A | A | missense_variant | gacL | M5005_Spy0613 | c.517G>A | p.Ala173Thr | 38 | | | | | | 0 |
| 13437 | C | T | T | missense_variant | gacL | M5005_Spy0613 | c.518C>T | p.Ala173Val | 9 | | | | | | 0 |
| 13439 | C | T | T | synonymous_variant | gacL | M5005_Spy0613 | c.520C>T | p.Leu174Leu | 3 | | | | | | 0 |
| 13444 | T | C | C | synonymous_variant | gacL | M5005_Spy0613 | c.525T>C | p.Asp175Asp | 2 | | | | | | 0 |
| 13483 | C | T | T | synonymous_variant | gacL | M5005_Spy0613 | c.564C>T | p.Cys188Cys | 1 | | | | | | 0 |
| 13486 | C | T | T | synonymous_variant | gacL | M5005_Spy0613 | c.567C>T | p.Asn189Asn | 1 | | | | | | 0 |
| 13489 | T | C | C | synonymous_variant | gacL | M5005_Spy0613 | c.570T>C | p.Leu190Leu | 4 | | | | | | 0 |
| 13490 | A | G | G | missense_variant | gacL | M5005_Spy0613 | c.571A>G | p.Ile191Val | 2 | | | | | | 0 |
| 13495 | C | T | T | synonymous_variant | gacL | M5005_Spy0613 | c.576C>T | p.Ser192Ser | 20 | | | | | | 0 |
| 13504 | G | A | A | missense_variant | gacL | M5005_Spy0613 | c.585G>A | p.Met195Ile | 6 | | | | | | 0 |
| 13508 | G | T,A | A | missense_variant | gacL | M5005_Spy0613 | c.589G>A | p.Ala197Thr | 33 | T | missense_variant | gacL | c.589G>T | p.Ala197Ser | 2 |
| 13520 | A | G | G | missense_variant | gacL | M5005_Spy0613 | c.601A>G | p.Ile201Val | 8 | | | | | | 0 |
| 13528 | G | A | A | synonymous_variant | gacL | M5005_Spy0613 | c.609G>A | p.Leu203Leu | 28 | | | | | | 0 |
| 13530 | C | T | T | missense_variant | gacL | M5005_Spy0613 | c.611C>T | p.Ser204Phe | 1 | | | | | | 0 |
| 13543 | A | G | G | synonymous_variant | gacL | M5005_Spy0613 | c.624A>G | p.Lys208Lys | 16 | | | | | | 0 |
| 13579 | C | T | T | synonymous_variant | gacL | M5005_Spy0613 | c.660C>T | p.Asn220Asn | 2 | | | | | | 0 |
| 13597 | T | C | C | synonymous_variant | gacL | M5005_Spy0613 | c.678T>C | p.Ser226Ser | 13 | | | | | | 0 |
| 13639 | A | G | G | synonymous_variant | gacL | M5005_Spy0613 | c.720A>G | p.Thr240Thr | 1 | | | | | | 0 |
| 13647 | T | G | G | missense_variant | gacL | M5005_Spy0613 | c.728T>G | p.Leu243Trp | 1 | | | | | | 0 |
| 13665 | C | T | T | missense_variant | gacL | M5005_Spy0613 | c.746C>T | p.Ser249Leu | 3 | | | | | | 0 |

| | | | | | | | | | | | | | | | |
|---|---|---|---|---|---|---|---|---|---|---|---|---|---|---|---|
| 13668 | C | T | T | missense_variant | gacL | M5005_Spy0613 | c.749C>T | p.Ser250Leu | 34 | | | | | | 0 |
| 13670 | A | G | G | missense_variant | gacL | M5005_Spy0613 | c.751A>G | p.Lys251Glu | 33 | | | | | | 0 |
| 13686 | A | G | G | missense_variant | gacL | M5005_Spy0613 | c.767A>G | p.Tyr256Cys | 72 | | | | | | 0 |
| 13689 | A | G | G | missense_variant | gacL | M5005_Spy0613 | c.770A>G | p.Asn257Ser | 2 | | | | | | 0 |
| 13691 | A | G | G | missense_variant | gacL | M5005_Spy0613 | c.772A>G | p.Thr258Ala | 1 | | | | | | 0 |
| 13693 | C | T | T | synonymous_variant | gacL | M5005_Spy0613 | c.774C>T | p.Thr258Thr | 362 | | | | | | 0 |
| 13702 | C | T | T | synonymous_variant | gacL | M5005_Spy0613 | c.783C>T | p.Phe261Phe | 2 | | | | | | 0 |
| 13711 | A | G | G | synonymous_variant | gacL | M5005_Spy0613 | c.792A>G | p.Leu264Leu | 7 | | | | | | 0 |
| 13715 | C | T | T | synonymous_variant | gacL | M5005_Spy0613 | c.796C>T | p.Leu266Leu | 1 | | | | | | 0 |
| 13717 | A | G | G | synonymous_variant | gacL | M5005_Spy0613 | c.798A>G | p.Leu266Leu | 4 | | | | | | 0 |
| 13727 | C | T | T | synonymous_variant | gacL | M5005_Spy0613 | c.808C>T | p.Leu270Leu | 61 | | | | | | 0 |
| 13730 | G | A | A | missense_variant | gacL | M5005_Spy0613 | c.811G>A | p.Val271Ile | 1 | | | | | | 0 |
| 13761 | T | C | C | missense_variant | gacL | M5005_Spy0613 | c.842T>C | p.Val281Ala | 12 | | | | | | 0 |
| 13764 | G | A | A | missense_variant | gacL | M5005_Spy0613 | c.845G>A | p.Cys282Tyr | 2 | | | | | | 0 |
| 13768 | T | C | C | synonymous_variant | gacL | M5005_Spy0613 | c.849T>C | p.Leu283Leu | 144 | | | | | | 0 |
| 13770 | C | T | T | missense_variant | gacL | M5005_Spy0613 | c.851C>T | p.Pro284Leu | 1 | | | | | | 0 |
| 13787 | G | A | A | missense_variant | gacL | M5005_Spy0613 | c.868G>A | p.Glu290Lys | 25 | | | | | | 0 |
| 13791 | A | G | G | missense_variant | gacL | M5005_Spy0613 | c.872A>G | p.Lys291Arg | 1 | | | | | | 0 |
| 13793 | T | A | A | missense_variant | gacL | M5005_Spy0613 | c.874T>A | p.Phe292Ile | 1 | | | | | | 0 |
| 13795 | C | T,G | G | missense_variant | gacL | M5005_Spy0613 | c.876C>G | p.Phe292Leu | 3 | T | synonymous_variant | gacL | c.876C>T | p.Phe292Phe | 21 |
| 13806 | C | T | T | missense_variant | gacL | M5005_Spy0613 | c.887C>T | p.Thr296Ile | 7 | | | | | | 0 |
| 13822 | G | A | A | synonymous_variant | gacL | M5005_Spy0613 | c.903G>A | p.Leu301Leu | 31 | | | | | | 0 |
| 13838 | C | T | T | synonymous_variant | gacL | M5005_Spy0613 | c.919C>T | p.Leu307Leu | 8 | | | | | | 0 |
| 13840 | A | G | G | synonymous_variant | gacL | M5005_Spy0613 | c.921A>G | p.Leu307Leu | 1 | | | | | | 0 |
| 13882 | T | C | C | synonymous_variant | gacL | M5005_Spy0613 | c.963T>C | p.Tyr321Tyr | 35 | | | | | | 0 |
| 13904 | G | A | A | missense_variant | gacL | M5005_Spy0613 | c.985G>A | p.Val329Ile | 3 | | | | | | 0 |
| 13989 | T | C | C | missense_variant | gacL | M5005_Spy0613 | c.1070T>C | p.Ile357Thr | 2 | | | | | | 0 |
| 13993 | A | G | G | synonymous_variant | gacL | M5005_Spy0613 | c.1074A>G | p.Arg358Arg | 22 | | | | | | 0 |
| 13996 | T | C | C | synonymous_variant | gacL | M5005_Spy0613 | c.1077T>C | p.His359His | 24 | | | | | | 0 |
| 14010 | T | A | A | missense_variant | gacL | M5005_Spy0613 | c.1091T>A | p.Leu364Gln | 1 | | | | | | 0 |
| 14014 | T | C | C | synonymous_variant | gacL | M5005_Spy0613 | c.1095T>C | p.Ile365Ile | 32 | | | | | | 0 |
| 14026 | T | C | C | synonymous_variant | gacL | M5005_Spy0613 | c.1107T>C | p.Asn369Asn | 18 | | | | | | 0 |
| 14029 | T | A | A | synonymous_variant | gacL | M5005_Spy0613 | c.1110T>A | p.Ile370Ile | 22 | | | | | | 0 |
| 14035 | G | A | A | synonymous_variant | gacL | M5005_Spy0613 | c.1116G>A | p.Gln372Gln | 1 | | | | | | 0 |
| 14036 | T | G | G | missense_variant | gacL | M5005_Spy0613 | c.1117T>G | p.Phe373Val | 1 | | | | | | 0 |
| 14041 | C | T | T | synonymous_variant | gacL | M5005_Spy0613 | c.1122C>T | p.Gly374Gly | 41 | | | | | | 0 |
| 14056 | T | C | C | synonymous_variant | gacL | M5005_Spy0613 | c.1137T>C | p.Gly379Gly | 281 | | | | | | 0 |
| 14065 | C | T | T | synonymous_variant | gacL | M5005_Spy0613 | c.1146C>T | p.Asn382Asn | 6 | | | | | | 0 |
| 14066 | C | A | A | missense_variant | gacL | M5005_Spy0613 | c.1147C>A | p.Leu383Ile | 2 | | | | | | 0 |
| 14068 | T | A | A | synonymous_variant | gacL | M5005_Spy0613 | c.1149T>A | p.Leu383Leu | 418 | | | | | | 0 |
| 14072 | C | T | T | missense_variant | gacL | M5005_Spy0613 | c.1153C>T | p.Pro385Ser | 1 | | | | | | 0 |
| 14074 | G | C | C | synonymous_variant | gacL | M5005_Spy0613 | c.1155G>C | p.Pro385Pro | 17 | | | | | | 0 |
| 14085 | G | C | C | missense_variant | gacL | M5005_Spy0613 | c.1166G>C | p.Cys389Ser | 16 | | | | | | 0 |
| 14089 | C | T | T | synonymous_variant | gacL | M5005_Spy0613 | c.1170C>T | p.Phe390Phe | 1 | | | | | | 0 |
| 14099 | G | A | A | missense_variant | gacL | M5005_Spy0613 | c.1180G>A | p.Val394Ile | 2 | | | | | | 0 |
| 14106 | T | A | A | missense_variant | gacL | M5005_Spy0613 | c.1187T>A | p.Ile396Asn | 1 | | | | | | 0 |
| 14114 | G | A | A | missense_variant | gacL | M5005_Spy0613 | c.1195G>A | p.Ala399Thr | 7 | | | | | | 0 |
| 14118 | G | C | C | missense_variant | gacL | M5005_Spy0613 | c.1199G>C | p.Ser400Thr | 240 | | | | | | 0 |
| 14128 | T | C | C | synonymous_variant | gacL | M5005_Spy0613 | c.1209T>C | p.Thr403Thr | 1 | | | | | | 0 |
| 14133 | A | G | G | missense_variant | gacL | M5005_Spy0613 | c.1214A>G | p.Asn405Ser | 16 | | | | | | 0 |
| 14137 | T | C | C | synonymous_variant | gacL | M5005_Spy0613 | c.1218T>C | p.Ile406Ile | 3 | | | | | | 0 |
| 14138 | G | T | T | missense_variant | gacL | M5005_Spy0613 | c.1219G>T | p.Val407Leu | 1 | | | | | | 0 |
| 14147 | A | C | C | missense_variant | gacL | M5005_Spy0613 | c.1228A>C | p.Met410Leu | 45 | | | | | | 0 |
| 14158 | C | T | T | synonymous_variant | gacL | M5005_Spy0613 | c.1239C>T | p.Ile413Ile | 1 | | | | | | 0 |
| 14162 | G | A | A | missense_variant | gacL | M5005_Spy0613 | c.1243G>A | p.Val415Ile | 1 | | | | | | 0 |
| 14182 | T | C | C | synonymous_variant | gacL | M5005_Spy0613 | c.1263T>C | p.Gly421Gly | 3 | | | | | | 0 |
| 14204 | C | T,A | A | missense_variant | gacL | M5005_Spy0613 | c.1285C>A | p.Leu429Met | 2 | T | synonymous_variant | gacL | c.1285C>T | p.Leu429Leu | 1 |
| 14215 | A | G | G | synonymous_variant | gacL | M5005_Spy0613 | c.1296A>G | p.Thr432Thr | 8 | | | | | | 0 |
| 14224 | G | C | C | synonymous_variant | gacL | M5005_Spy0613 | c.1305G>C | p.Gly435Gly | 5 | | | | | | 0 |
| 14226 | C | T | T | missense_variant | gacL | M5005_Spy0613 | c.1307C>T | p.Ser436Phe | 2 | | | | | | 0 |
| 14234 | G | A | A | missense_variant | gacL | M5005_Spy0613 | c.1315G>A | p.Val439Ile | 1 | | | | | | 0 |
| 14236 | C | A,T | A | synonymous_variant | gacL | M5005_Spy0613 | c.1317C>A | p.Val439Val | 283 | T | synonymous_variant | gacL | c.1317C>T | p.Val439Val | 1 |

| | | | | | | | | | | | | | | | |
|---|---|---|---|---|---|---|---|---|---|---|---|---|---|---|---|
| 14261 | A | G | G | missense_variant | gacL | M5005_Spy0613 | c.1342A>G | p.Ile448Val | 26 | | | | | | 0 |
| 14266 | A | G | G | synonymous_variant | gacL | M5005_Spy0613 | c.1347A>G | p.Gly449Gly | 3 | | | | | | 0 |
| 14272 | C | T | T | synonymous_variant | gacL | M5005_Spy0613 | c.1353C>T | p.Ile451Ile | 14 | | | | | | 0 |
| 14273 | C | A | A | missense_variant | gacL | M5005_Spy0613 | c.1354C>A | p.Pro452Thr | 5 | | | | | | 0 |
| 14278 | A | G | G | synonymous_variant | gacL | M5005_Spy0613 | c.1359A>G | p.Leu453Leu | 39 | | | | | | 0 |
| 14284 | G | T,A | T | missense_variant | gacL | M5005_Spy0613 | c.1365G>T | p.Leu455Phe | 51 | A | synonymous_variant | gacL | c.1365G>A | p.Leu455Leu | 1 |
| 14307 | A | G | G | missense_variant | gacL | M5005_Spy0613 | c.1388A>G | p.Gln463Arg | 1 | | | | | | 0 |
| 14319 | A | C | C | missense_variant | gacL | M5005_Spy0613 | c.1400A>C | p.Gln467Pro | 1 | | | | | | 0 |
| 14327 | G | A | A | missense_variant | gacL | M5005_Spy0613 | c.1408G>A | p.Asp470Asn | 1 | | | | | | 0 |
| 14328 | A | G,C | C | missense_variant | gacL | M5005_Spy0613 | c.1409A>C | p.Asp470Ala | 1 | G | missense_variant | gacL | c.1409A>G | p.Asp470Gly | 2 |
| 14332 | C | A,T | A | synonymous_variant | gacL | M5005_Spy0613 | c.1413C>A | p.Ile471Ile | 10 | T | synonymous_variant | gacL | c.1413C>T | p.Ile471Ile | 1 |
| 14335 | T | G | G | synonymous_variant | gacL | M5005_Spy0613 | c.1416T>G | p.Leu472Leu | 1 | | | | | | 0 |
| 14347 | G | A | A | synonymous_variant | gacL | M5005_Spy0613 | c.1428G>A | p.Leu476Leu | 2 | | | | | | 0 |
| 14357 | G | A | A | missense_variant | gacL | M5005_Spy0613 | c.1438G>A | p.Val480Ile | 278 | | | | | | 0 |
| 14362 | G | T | T | synonymous_variant | gacL | M5005_Spy0613 | c.1443G>T | p.Ser481Ser | 6 | | | | | | 0 |
| 14363 | C | T | T | missense_variant | gacL | M5005_Spy0613 | c.1444C>T | p.Leu482Phe | 2 | | | | | | 0 |
| 14368 | G | T | T | missense_variant | gacL | M5005_Spy0613 | c.1449G>T | p.Leu483Phe | 77 | | | | | | 0 |
| 14377 | T | C | C | synonymous_variant | gacL | M5005_Spy0613 | c.1458T>C | p.Leu486Leu | 1947 | | | | | | 0 |
| 14387 | A | G | G | missense_variant | gacL | M5005_Spy0613 | c.1468A>G | p.Met490Val | 1 | | | | | | 0 |
| 14407 | T | A,C | A | stop_gained | gacL | M5005_Spy0613 | c.1488T>A | p.Tyr496* | 1 | C | synonymous_variant | gacL | c.1488T>C | p.Tyr496Tyr | 1 |
